# Thiazole substitution of a labile amide bond - a new option towards stable pantothenamide-mimics

**DOI:** 10.1101/2025.02.25.640258

**Authors:** Xiangning Liu, Annica Chu, Mina Nekouei, Chunling Blue Lan, Karine Auclair, Kevin J. Saliba

## Abstract

The emergence and spread of artemisinin-resistant, malaria-causing *P. falciparum* provide the impetus for the development of novel antimalarials. Pantothenamides are potent inhibitors of malaria parasite proliferation, however their clinical use is hindered by pantetheinase-mediated degradation in human serum. Here we report the synthesis and biological activity of a series of pantothenamide-mimics in which the labile amide bond is replaced by a thiazole ring with various orientations. Out of 23 novel compounds generated and tested in the presence of pantetheinase, several display sub-micromolar antiplasmodial activity *in vitro*. A selection of compounds was studied in more detail and CoA biosynthesis and/or utilisation pathways were confirmed to be the target. Toxicity to human cells was not observed. Kinetic studies identified the selected compounds as substrates of the *Hs*PanK3 enzyme, but with much lower affinity compared to that of the natural substrate pantothenate. The most potent thiazole-bearing antiplasmodial compound was found to bind to *Pf*PanK with a 120-fold higher affinity compared to *Hs*PanK, highlighting excellent selectivity, not only against the key first enzyme in the CoA biosynthesis pathway, but also at the whole-cell level. In conclusion, thiazole substitution of the labile amide bond represents a promising avenue for the development of antimalarial pantothenamide-mimics.

## 1. INTRODUCTION

Malaria is a lethal infectious disease caused by unicellular protozoan parasites of the genus *Plasmodium*. Despite remarkable progress towards malaria elimination, the 2024 World Malaria Report estimated 263 million new cases and 597,000 deaths worldwide attributed to malaria in 2023 (1). Of the six *Plasmodium* species that infect humans (2, 3), *Plasmodium falciparum* is responsible for most of the infections and deaths in the sub-Saharan African region (2, 4). Currently, artemisinin-based combination therapies (ACT) are the first-line treatment for uncomplicated *falciparum* malaria in all areas. However, the emergence of ACT-resistant parasites and their rapid spread across the globe (1, 5, 6), including in recent cases of severe malaria in Africa (7), threaten the efficacy of ACT, thus hampering the treatment of the disease. To increase the repertoire of available therapeutics, discovery of new antimalarials with unique mechanisms of action and no cross-resistance with existing drugs is essential (8, 9).

Water-soluble vitamin B_5_ (pantothenate) is an essential substance required by the intraerythrocytic stage parasites. Pantothenate is the precursor to coenzyme A (CoA), a cofactor involved in many metabolic processes (10, 11). Pantothenate utilisation and CoA biosynthesis in *P. falciparum* have been extensively studied (12–15) and have been identified as viable drug targets in the parasite (16, 17). First synthesized by Clifton *et al.* in 1970 (18), pantothenamides were initially reported to exhibit antibacterial activity (17, 19), with their antiplasmodial activity demonstrated by Spry *et al.* in 2013 (20). In *P. falciparum*, pantothenamides are first phosphorylated by pantothenate kinase (PanK) and further metabolised by phosphopantetheine adenylyltransferase (PPAT) and dephospho-CoA kinase (DPCK) to generate the corresponding antiplasmodial CoA antimetabolite (21, 22). Despite their high potency and low cytotoxicity *in vitro*, the clinical use of pantothenamides, such as the benchmark compound N5-Pan, is hindered due to their instability *in vivo* (**Figures 1A** and **1B**) (20, 23). The pantetheinases in human serum readily hydrolyze the amide depicted in red in Figure 1A, resulting in the rapid breakdown of pantothenamides. To overcome this issue, our research group and others have employed various structural modifications, including alterations at the *gem*-dimethyl group (24–26), the pantoyl primary alcohol (27), β-alanine (27–30), and the labile amide (27, 29, 31–35) moieties of pantothenamides. These findings motivated our search for new pantetheinase-resistant pantothenamides.

**Figure 1.**
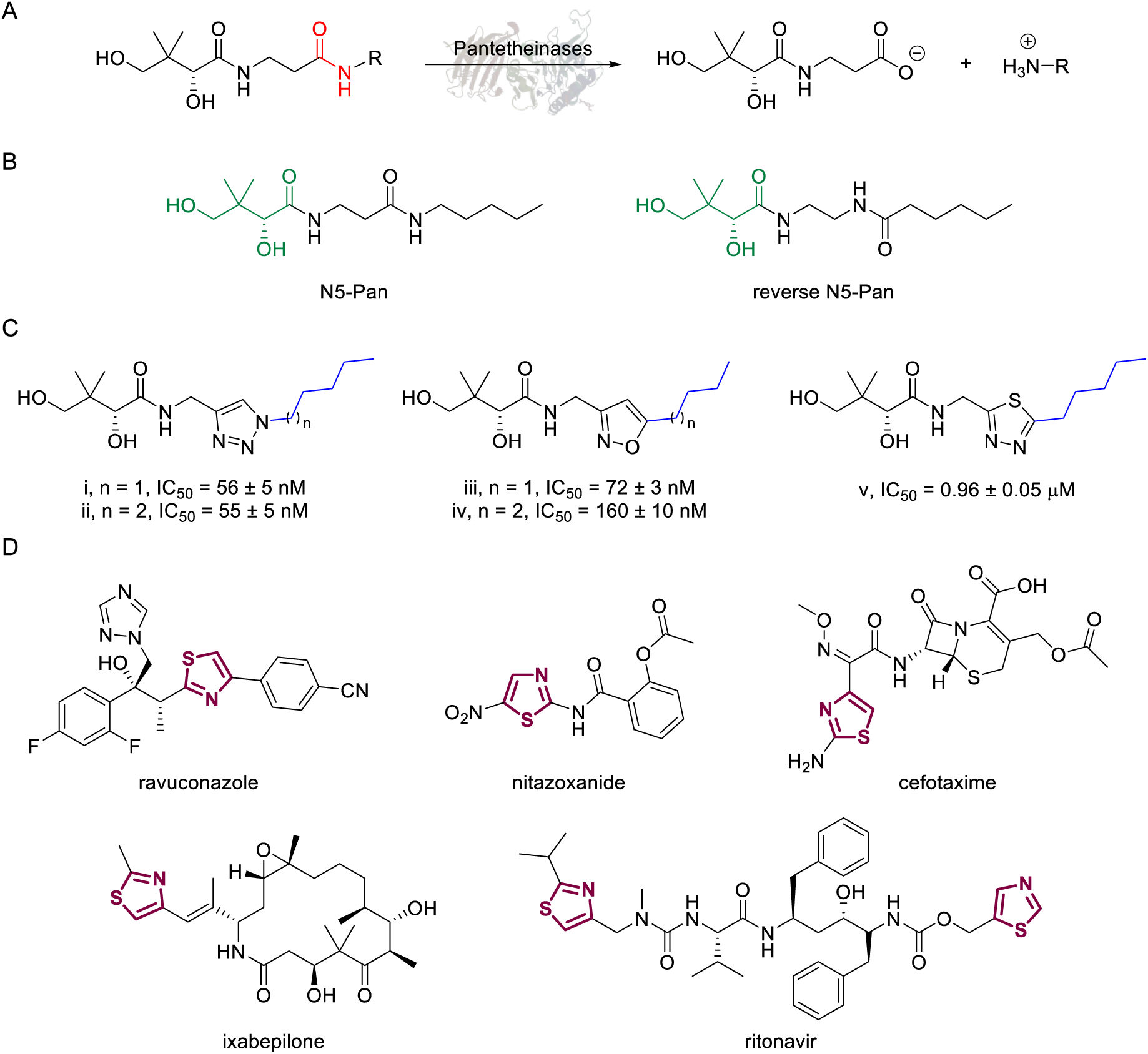
(**A**) Pantothenamides are readily hydrolysed by serum pantetheinases to yield pantothenate and an amine. The pantetheinase-sensitive amide bond is shown in red. R is commonly an aliphatic or aromatic moiety. **(B)** Structures of N5-Pan and reverse N5-Pan. The pantoyl moiety is drawn in green. **(C)** Structures of compounds **i** to **v** with their respective 50% inhibitory concentrations (IC_50_ values) measured against intraerythrocytic *P. falciparum* in the presence of pantetheinases. Side chains are shown in blue. **(D)** Examples of thiazole-containing pharmaceuticals. Thiazoles are highlighted in maroon.

We have recently focused on replacing the labile amide with heterocycles (32, 33, 35), which are privileged scaffolds in medicinal chemistry. More than 80% of marketed drugs contain at least one heterocyclic fragment in their structure (36, 37). They can participate in diverse secondary interactions due to the presence of the heteroatoms and nuanced aromaticity, and can significantly affect the physicochemical properties of the molecules (36, 38). Previously, we reported a series of analogues in which the labile amide group is replaced with diverse heterocycles, resulting in pantothenamide analogues that exhibited pantetheinase resistance and enhanced potency (32, 33, 35). In particular, the 1,4-substituted 1,2,3-triazole with a pentyl or hexyl side chain (compounds **i** and **ii**, **Figure 1C**), the 3,5-substituted 1,2-isoxazole with a butyl or pentyl chain (compounds **iii** and **iv**), as well as the 2,5-substituted 1,3,4-thiadiazole (compound **v**) were particularly potent.

Sulfur is an ubiquitous heteroatom in medicinal chemistry (39) and a variety of sulfur-containing scaffolds are employed in the pharmaceutical industry (40). Sulfur is known to serve as an excellent σ-hole to host electrons from adjacent donors, which has profound impact on conformational stabilization. This can greatly reduce entropic penalty during target engagement (38, 41), thus enhancing potency. One of the most prevalent scaffolds in this category is the thiazole (42). Examples of thiazole-containing pharmaceuticals (**Figure 1C**) include ravuconazole (antifungal) (43), nitazoxanide (antiparasitic) (44), cefotaxime (antibacterial) (45), ixabepilone (anticancer) (46), as well as ritonavir (antiviral) (47). Despite the high prevalence of sulfur in medicines, sulfur-containing pantothenamides are currently underdeveloped. To date, 11 such compounds have been reported (27, 30, 31, 34, 35), and only two analogues had a sulfur-containing heterocycle (35).

Herein, we describe the synthesis, antiplasmodial activity, target determination and toxicity profile of novel pantothenamide-mimics containing thiazoles. In addition, we report the kinetic analysis of a selection of these compounds using recombinantly-expressed and purified human PanK 3 (*Hs*PanK3) as well as the *P. falciparum* PanK (*Pf*PanK) in parasite lysates.

## 2. RESULTS

### 2.1. Synthesis

Considering that a one-carbon linker was reported to be preferred between the pantoyl (in green in **Figure 2B**) and the heterocyclic moieties in pantothenamide-mimics containing a triazole or isoxazole in place of the labile amide bond (32, 33, 35), we elected to synthesize thiazole analogues that harbor a one-carbon linker. **Scheme 1** illustrates the synthetic route for compounds **1a**-**1e** containing 2-aminomethyl-4-substituted thiazoles. Acid-catalyzed monobromination of the methyl ketones was first achieved to access the bromoketone fragments **1.1a**-**1.1e** for Hantzsch thiazole synthesis. The thioamide counterpart was synthesized through an amidation-thiation sequence from Boc-glycine methyl ester. Traditional Hantzsch thiazole synthesis between **1.3** and **1.1a**-**1.1e** generated the thiazole rings. Boc deprotection, followed by aminolysis of D-pantolactone yielded the target compounds **1a**-**1e**. Notably, the aminolysis conditions under microwave irradiation reported before (31) were not successful here, as decomposition was observed during the reaction. Instead, strictly thermal conditions were employed to avoid decomposition of the starting materials.

**Figure 2.**
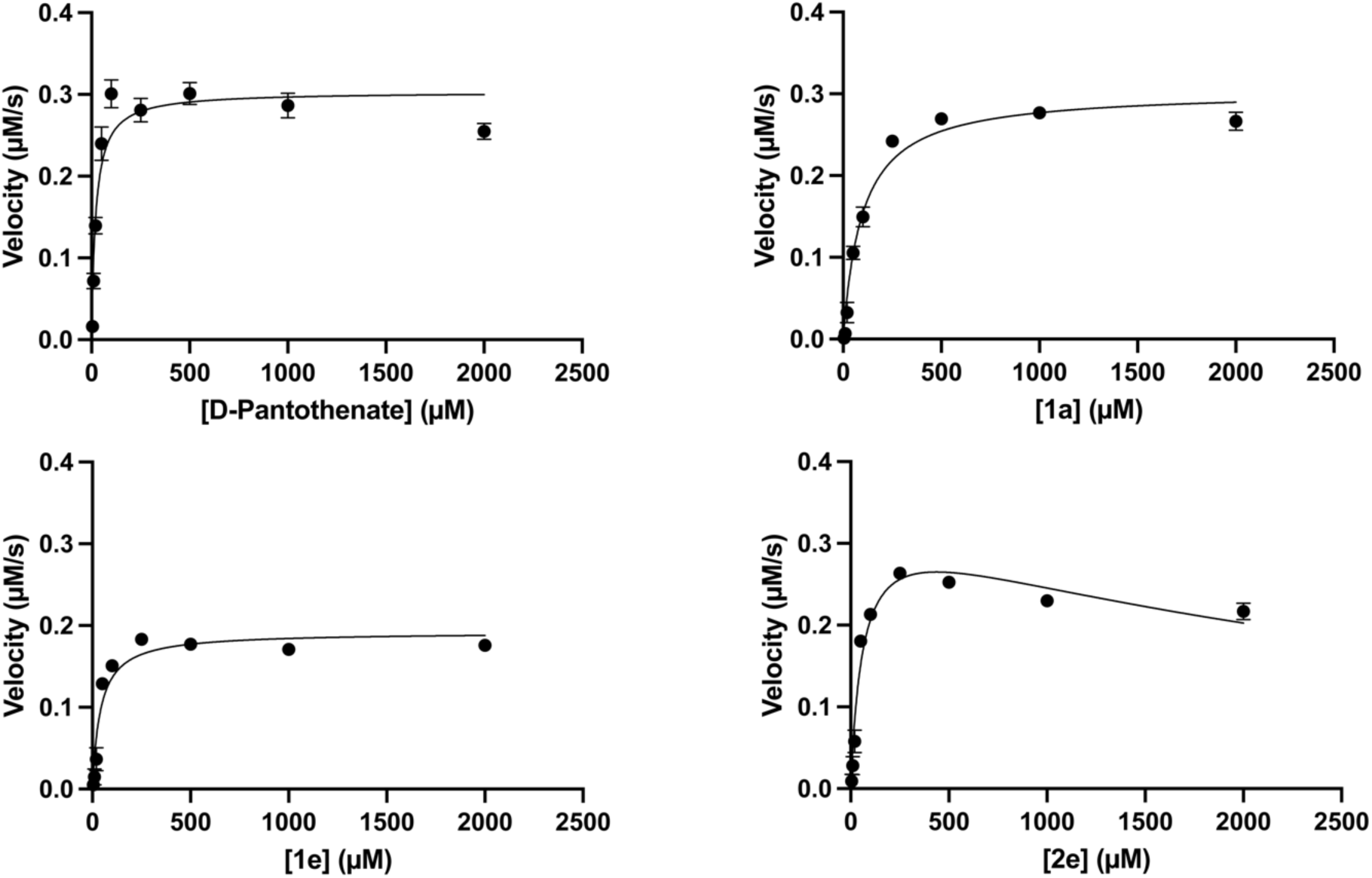
Examples of kinetics plots for *Hs*PanK3 in the presence of representative compounds. Velocities were determined as a function of pantothenate or compound concentration. The data points were fit to the Michaelis–Menten nonlinear regression equation to determine kinetic parameters. For **2e**, Equation 1, which accounts for substrate inhibition, was used to fit the data. Data are averaged from 3 independent experiments, each performed in triplicate. Error bars represent SEM and are not visible if smaller than the symbols. Kinetics of other compounds are shown in **Figure S4**.

**Scheme 1.**
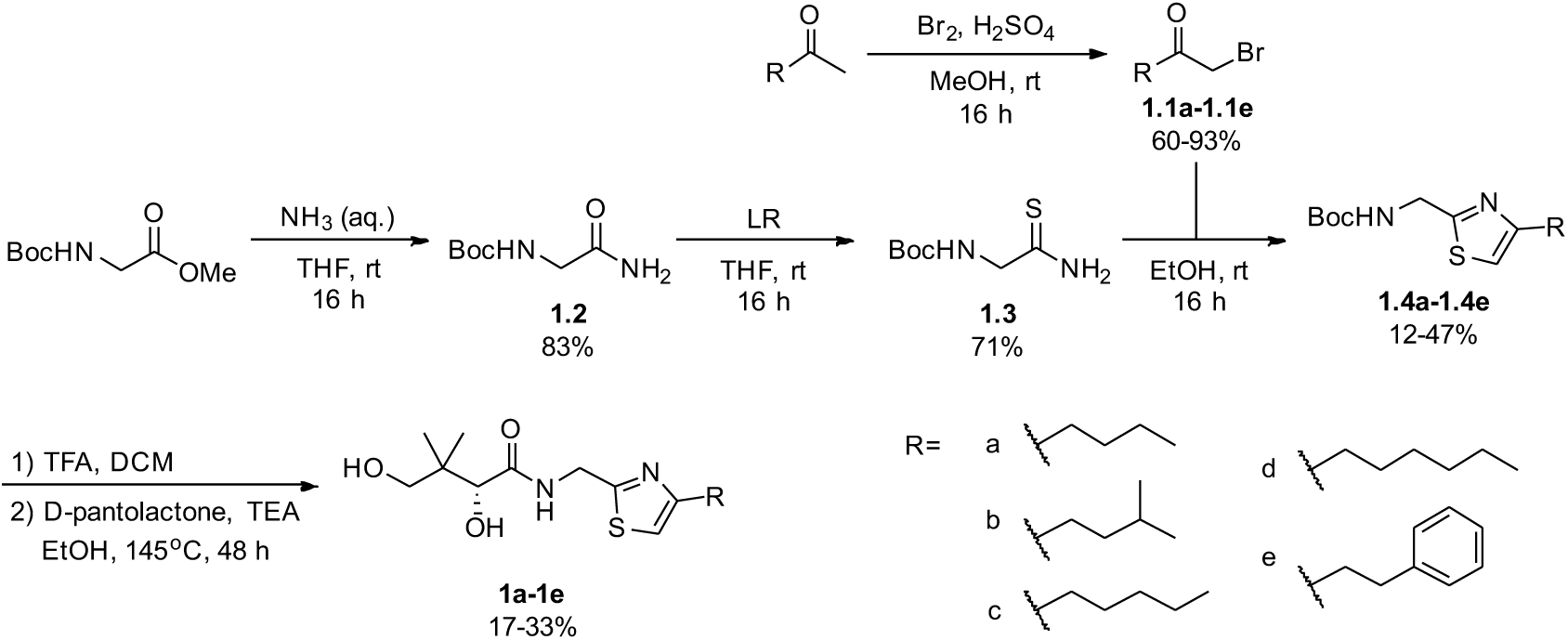
Synthetic route for compounds **1a**-**1e**. Boc: *tert*-butyloxycarbonyl; DCM: dichloromethane; LR, Lawesson’s reagent; rt: room temperature; TEA: triethylamine; TFA: trifluoroacetic acid; THF: tetrahydrofuran.

A similar approach was adopted for the synthesis of thiazole derivatives with the reverse ring orientation, *i.e.* 4-aminomethyl-2-substituted thiazoles **2a**-**2h** (**Scheme 2**). Substitution followed by monobromination afforded the bromoketones with a phthalimide-protected amino handle (**2.2**). The same amidation-thiation process as above was used to access the thioamides **2.4a**-**2.4h**, starting from various acyl chlorides. Likewise, Hantzsch thiazole synthesis led to the thiazole core. Finally, removal of the phthalimide protecting group, followed by aminolysis of D-pantolactone afforded the target compounds.

**Scheme 2.**
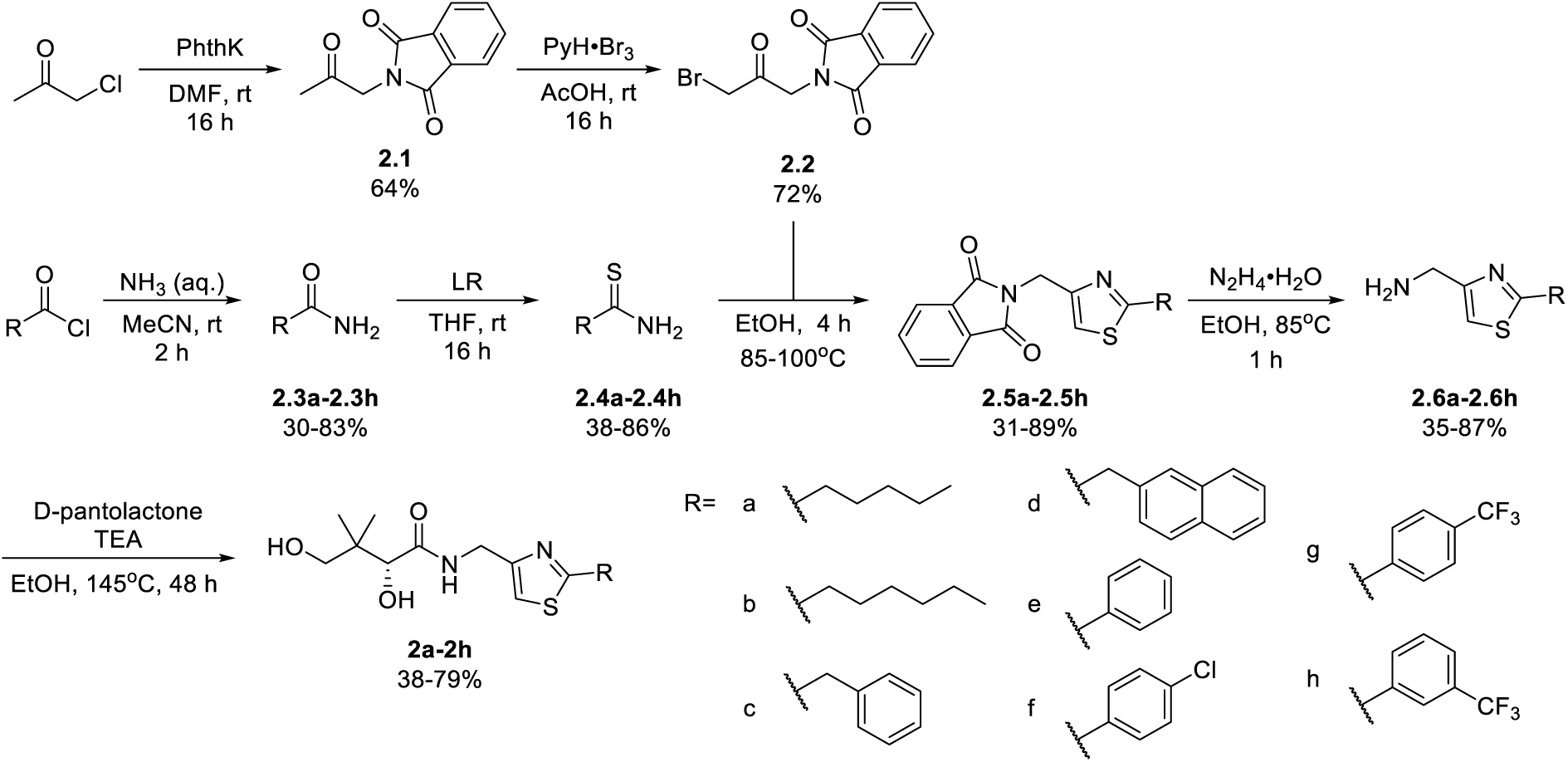
Synthetic route for compounds **2a**-**2h**. DMF: *N, N*-dimethylformamide; LR: Lawesson’s reagent; PhthK: potassium phthalimide; PyH: pyridine; rt: room temperature; TEA: triethylamine; THF: tetrahydrofuran.

For the synthesis of pantothenamide-mimics with 5-aminomethyl-2-substituted thiazoles (**Scheme 3**), two different routes were utilized. For synthetic targets harbouring a thiazole with alkyl substituents (**3a**-**3e**), decoration of the thiazole was adopted. Thus, Negishi coupling employing 2-bromo-5-cyanothiazole successfully furnished the alkyl substituted thiazole. Reduction of the nitrile afforded amines **3.2a**-**3.2e**, which were subjected to a recently published method (48) for catalytic D-pantolactone aminolysis to generate the corresponding products. To generate compounds with an aryl-substituted thiazole (**3f**-**3j**), a reported methodology was followed to access the ring (49). First, the amidation-thiation approach was again utilized to assemble the thioamides **3.4f**-**3.4j**, which underwent *N*-bromosuccinimide-promoted cyclization-oxidation to afford the thiazole core with a bromide handle. Gabriel synthesis converted the bromide handles to amines **3.7f**-**3.7j**, which were subjected to the same thermal aminolysis conditions as above to yield compound **3f**-**3j**.

**Scheme 3.**
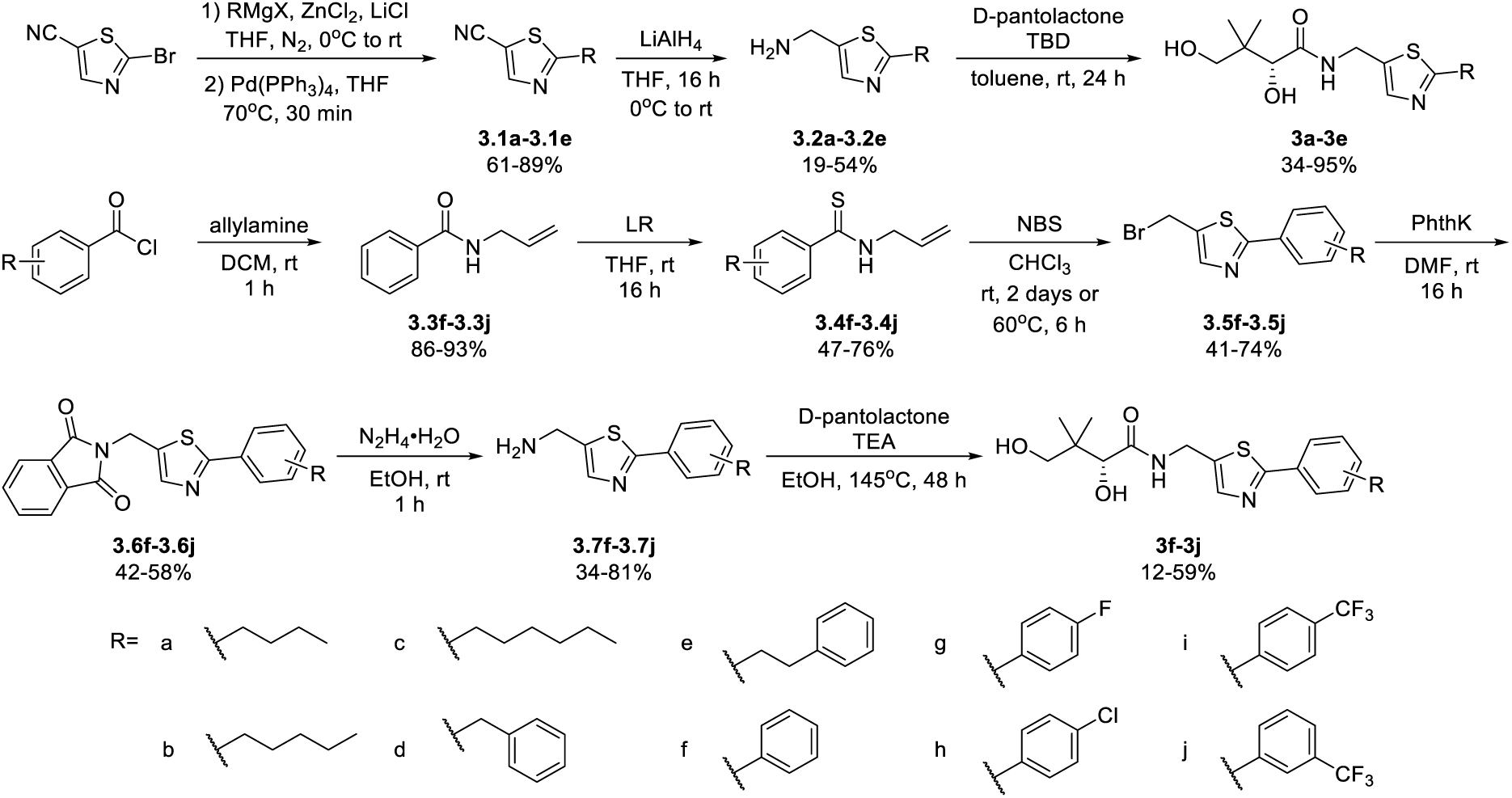
Synthetic route to compounds **3a**-**3j**. DCM: dichloromethane; DMF: *N, N*-dimethylformamide; LR: Lawesson’s reagent; NBS: *N*-bromosuccinimide; PhthK: potassium phthalimide; rt: room temperature; TBD: 1,5,7-triazabicyclo[4.4.0]dec-5-ene; TEA: triethylamine; THF: tetrahydrofuran.

### 2.2. *In Vitro* Antiplasmodial Activity

The antiplasmodial activity of all synthetic compounds was evaluated against *P. falciparum*. Thiazoles **1a**, **1e** and **2e** were found to inhibit intraerythrocytic proliferation of *P. falciparum* with IC_50_ values in the sub-micromolar range (**Table 1** and **Figure S1**). As evident from the overall higher activity of compound series **1** and series **2** relative to series **3**, it can be concluded that a nitrogen atom is preferred between the substituents of the ring, whereas a sulfur atom is beneficial at the other two ring positions. In series **1**, unlike what was previously reported by Howieson *et al*. (32) and Guan *et al*. (33, 35) for triazole– and isoxazole-containing analogues, phenethyl and butyl side chains are equally potent (see **1a** and **1e**), consistent with the length of a phenyl ring (2.8 Å) (50) roughly equalling that of a 3-carbon chain (2.5-2.6 Å) (51). The structure-activity relationships (SARs) reveal that a butyl chain is preferred over a pentyl or a hexyl group, and branching is detrimental. Interestingly, for the 4-aminomethyl-2-substituted thiazole series (**2a-2h**), alkyl chains were no longer preferred. Instead, the phenyl group is favoured (as in **2e**) with a slightly better IC_50_ (0.51 µM) than those of **1a** (0.77 µM) and **1e** (0.77 µM). Moreover, the addition of substituents to the phenyl ring was disadvantageous irrespective of the position, with larger substituents showing a stronger effect, as observed for *para*-chlorine (3.1 µM for **2f**), *para*-CF_3_ (57 µM for **2g**), and *meta*-trifluoromethyl (21 µM for **2h**). A similar trend was observed for the 5-aminomethyl-2-substitued thiazole series (**3a**-**3j**), with the preference for an unsubstituted phenyl group (**3f**) over alkyl groups, despite the overall lower potency of this series. In contrast to what was observed for **2f-2h**, **3j** is more potent than **3g-3i**, implying that a *meta*-trifluoromethyl group (11 µM for **3j**) is favored over any *para*-substituent group (14-46 µM for **3g-3i**) on the phenyl ring in this series.

**Table 1.**
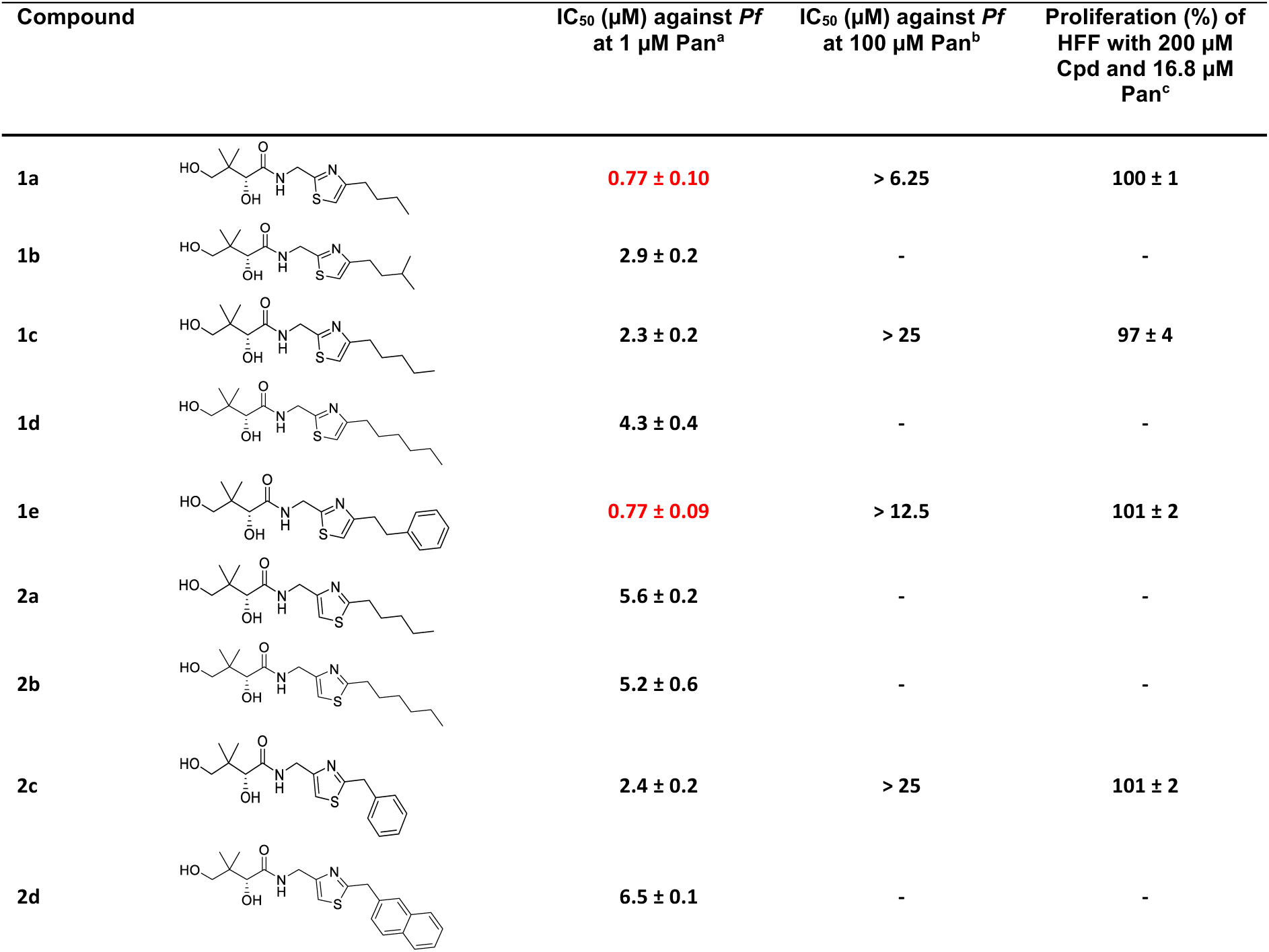
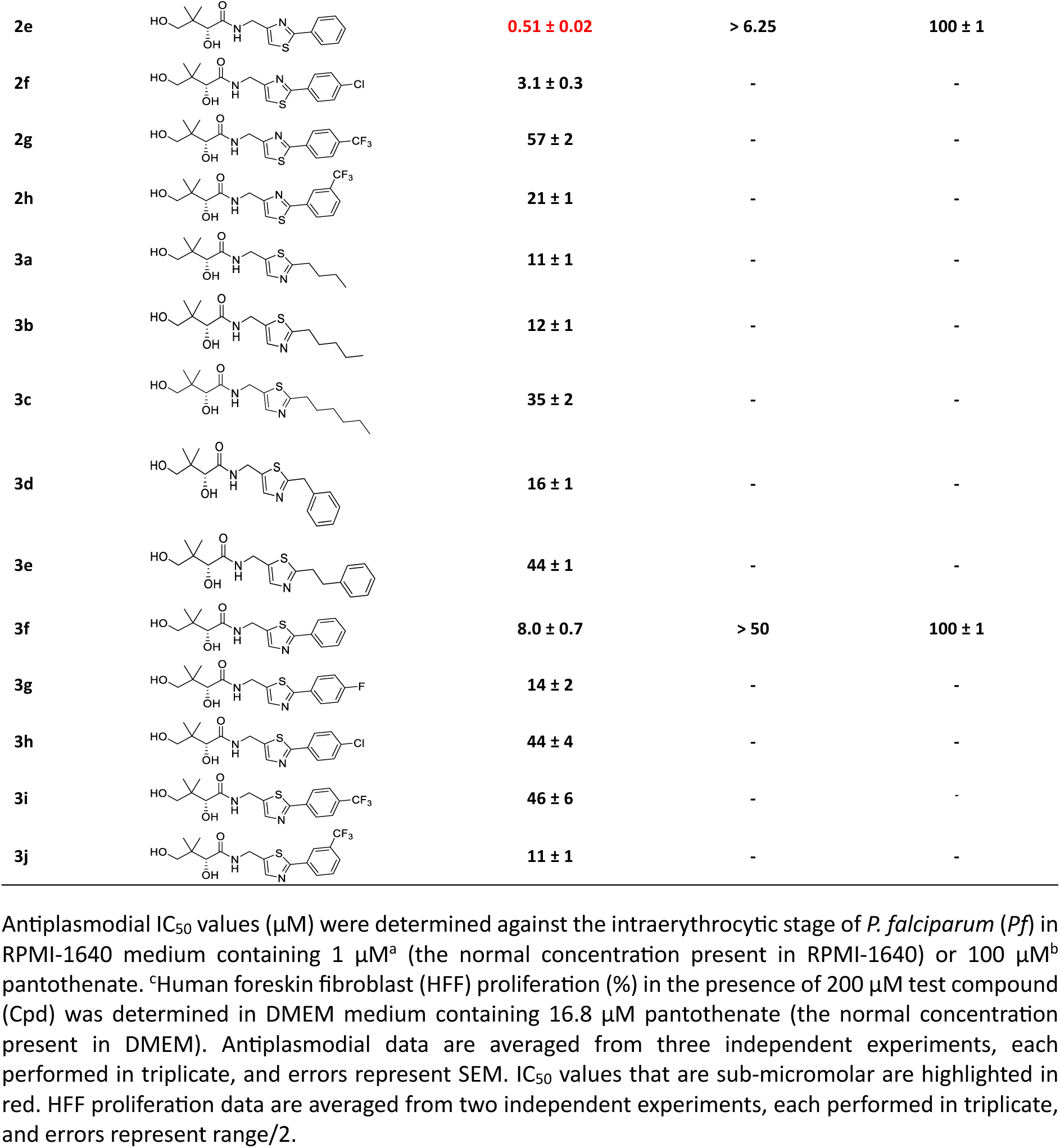
Effect of the compounds on the proliferation of intraerythrocytic *P. falciparum* and human foreskin fibroblasts (HFF) at different pantothenate concentrations.

| Compound | | IC<sub>50</sub> (μM) against *Pf*<br>at 1 μM Pan<sup>a</sup> | IC<sub>50</sub> (μM) against *Pf*<br>at 100 μM Pan<sup>b</sup> | Proliferation (%) of<br>HFF with 200 μM<br>Cpd and 16.8 μM<br>Pan<sup>c</sup> |
| --- | --- | --- | --- | --- |
| **1a** | | 0.77 ± 0.10 | > 6.25 | 100 ± 1 |
| **1b** | | 2.9 ± 0.2 | - | - |
| **1c** | | 2.3 ± 0.2 | > 25 | 97 ± 4 |
| **1d** | | 4.3 ± 0.4 | - | - |
| **1e** | | 0.77 ± 0.09 | > 12.5 | 101 ± 2 |
| **2a** | | 5.6 ± 0.2 | - | - |
| **2b** | | 5.2 ± 0.6 | - | - |
| **2c** | | 2.4 ± 0.2 | > 25 | 101 ± 2 |
| **2d** | | 6.5 ± 0.1 | - | - |
| 2e |  | 0.51 ± 0.02 | > 6.25 | 100 ± 1 |
| 2f |  | 3.1 ± 0.3 | - | - |
| 2g |  | 57 ± 2 | - | - |
| 2h |  | 21 ± 1 | - | - |
| 3a |  | 11 ± 1 | - | - |
| 3b |  | 12 ± 1 | - | - |
| 3c |  | 35 ± 2 | - | - |
| 3d |  | 16 ± 1 | - | - |
| 3e |  | 44 ± 1 | - | - |
| 3f |  | 8.0 ± 0.7 | > 50 | 100 ± 1 |
| 3g |  | 14 ± 2 | - | - |
| 3h |  | 44 ± 4 | - | - |
| 3i |  | 46 ± 6 | - | - |
| 3j |  | 11 ± 1 | - | - |

Antiplasmodial IC<sub>50</sub> values (μM) were determined against the intraerythrocytic stage of *P. falciparum* (Pf) in RPMI-1640 medium containing 1 μM<sup>a</sup> (the normal concentration present in RPMI-1640) or 100 μM<sup>b</sup> pantothenate. <sup>c</sup>Human foreskin fibroblast (HFF) proliferation (%) in the presence of 200 μM test compound (Cpd) was determined in DMEM medium containing 16.8 μM pantothenate (the normal concentration present in DMEM). Antiplasmodial data are averaged from three independent experiments, each performed in triplicate, and errors represent SEM. IC<sub>50</sub> values that are sub-micromolar are highlighted in red. HFF proliferation data are averaged from two independent experiments, each performed in triplicate, and errors represent range/2.

Previously, we have shown that the mechanism of action of pantothenamide-mimics against *P. falciparum* involves bioactivation by PanK, PPAT, and DPCK, resulting in the formation of the corresponding CoA antimetabolites that are hypothesized to inhibit CoA-utilizing enzymes (21, 35). To investigate whether the newly synthesised thiazole derivatives exert their antiplasmodial activity *via* a mechanism that is competitive with pantothenate, the substrate of PanK, we evaluated the antiplasmodial activity of a selection of compounds (**1a, 1c**, **1e**, **2c**, **2e** and **3f**) against *P. falciparum* in the presence of a higher concentration of extracellular pantothenate (100 µM, instead of the normal 1 µM that is present in RPMI-1640). Upon adding excess pantothenate, an increase (> 6-fold) in IC_50_ values was observed for all tested compounds (**Table 1** and **Figure S1**), similar to what was previously reported for triazole– and isoxazole-containing pantothenamide analogues (35), indicating that the thiazole derivatives are also competing with pantothenate, most likely as substrates of *Pf*PanK.

Subsequently, the effect of those compounds on the proliferation of human foreskin fibroblast (HFF) cells, a low-passage cell line, was assessed using the Incucyte® Live-Cell Analysis System, allowing automatic capture of cellular changes and assessment of cell proliferation in the presence of each compound. Cyclohexamide (10 µM), a potent protein synthesis inhibitor (52, 53), was used as a positive control, which completely inhibited the proliferation of HFF cells as expected (**Figure S2**). In contrast, none of the compounds inhibited the proliferation of the cells at 200 µM, the highest concentration tested, consistent with the compounds being nontoxic to HFF cells (**Table 1** and **Figure S2**).

### 2.3. Compound interaction with *Hs*PanK3 and *Pf*PanK

Considering the initial involvement of phosphorylation by *Pf*PanK in the mode of action of the novel thiazole derivatives, kinetic analyses were carried out for selected compounds to compare their transformation by *Pf*PanK and *Hs*PanK3. To date, three phylogenetically distinct types of PanK (type I, type II and type III) have been characterised, distinguished by their differences in structures, catalytic properties and inhibition profiles (11, 54). All PanKs characterised to date have been shown to exist as homodimers (55–62), with the exception of those from *P. falciparum* and *Toxoplasma gondii*, another apicomplexan parasite (63). *P. falciparum* expresses two type II PanKs (*Pf*PanK1 and *Pf*PanK2) which have been shown to form a heteromeric complex (63). To explore the difference in activity of the compounds against *P. falciparum* and HFF cells, we set out to perform kinetic analysis of *Hs*PanK3, a well-characterised homodimeric type II PanK expressed in humans, in the presence of selected compounds. The *Hs*PanK3 protein was expressed in *E. coli* as a His-tagged protein and purified by affinity chromatography (**Figure S3**). The purified *Hs*PanK3 protein was biochemically characterized and had a *K*_m_ for pantothenate of 24 ± 4 µM (n = 3; mean ± SEM; **Table 2** and **Figure 2**), which is consistent with previous reports (17 ± 1 µM) (64, 65). All tested compounds were found to be substrates of *Hs*PanK3. Substrate inhibition, which is a common phenomenon in enzyme kinetics (66–68), was observed for compounds **2c** and **2e** (**Figures 2** and **S4**). Such inhibition is typically caused by simultaneous binding of two or more substrate molecules to the enzyme’s active site, resulting in the formation of an unproductive enzyme– substrate complex (69). This increases with rising substrate concentration. Equation 1, which accounts for substrate inhibition, was used to determine the kinetic parameters of **2c** and **2e**, instead of the standard Michaelis-Menten equation that was used for the other compounds. We found that *Hs*PanK3 exhibited a 1.8-4.2 fold higher *K*_m_ (*i.e.* lower affinity) for all selected compounds compared to that of the natural substrate pantothenate. In contrast, the turnover numbers (*k*_cat_) of *Hs*PanK3 for **2e** and **3f** were slightly higher than that of pantothenate (*i.e.* higher efficiency). The *k*_cat_ for other compounds were the same as (**1a**, **1c**, **2c**) or slightly lower (**1e**) than the *k*_cat_ for pantothenate. The specificity of *Hs*PanK3 for each compound was then evaluated by computing the *k*_cat_/*K*_m_ values. In the presence of the compounds with sub-micromolar antiplasmodial activity (**1a**, **1e** or **2e**), *Hs*PanK3 exhibited a 2.5-4.5 fold lower *k*_cat_/*K*_m_ (*i.e.* lower specificity) compared to the natural substrate pantothenate (**Table 2**). Similar results were obtained for **1c**, **2c**, and **3f** (∼3-fold lower), consistent with *Hs*PanK3 having a much higher catalytic efficiency for pantothenate than for the thiazole analogues. Together, these data imply that the compounds are unlikely to be substantially metabolised by the human enzyme, as expected from the lack of toxicity observed on HFF cells.

**Table 2.**
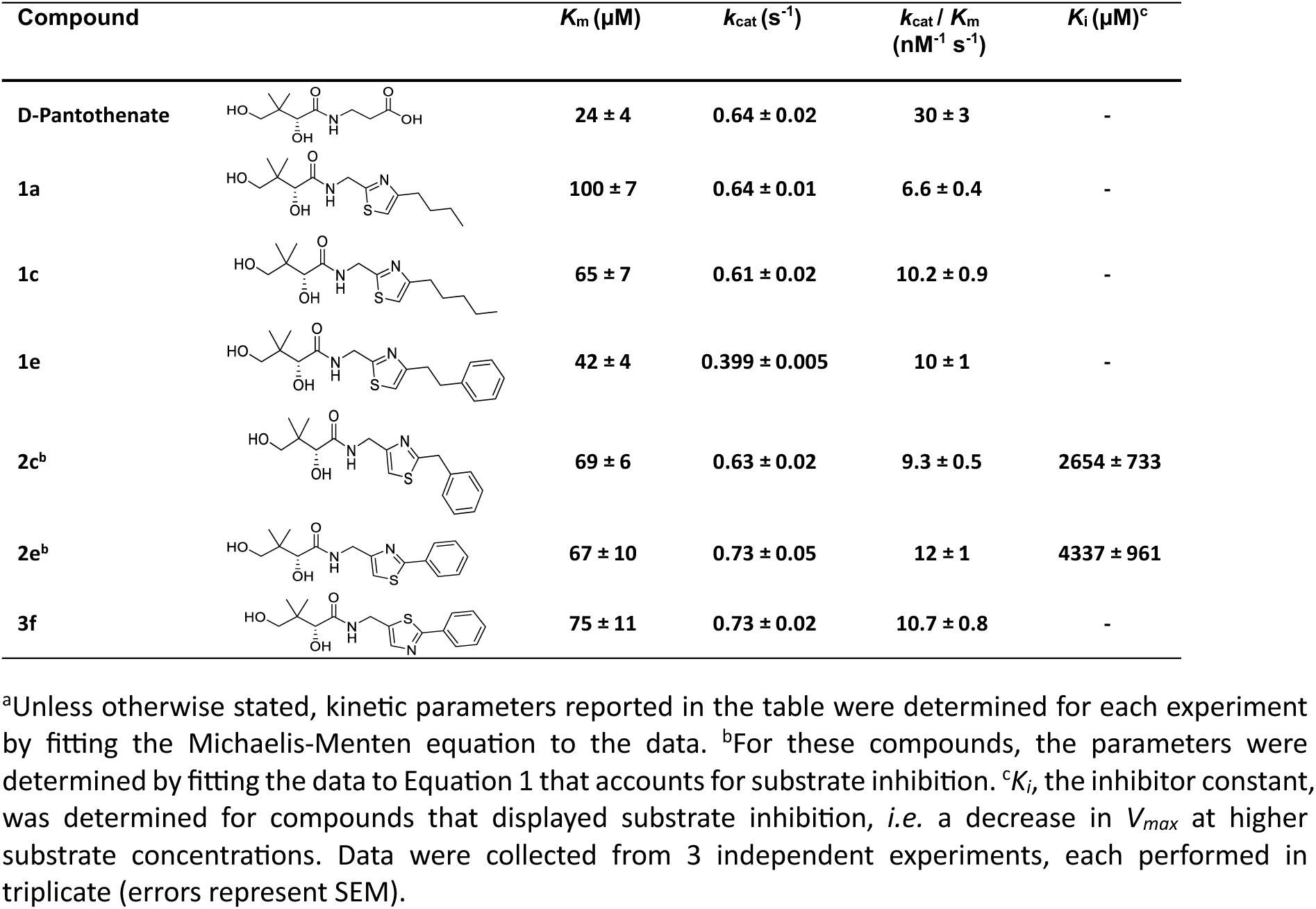
Kinetic data calculated for *Hs*PanK3 in the presence of various pantothenamide mimics^a^.

| Compound | | $K_m$ ($\mu\text{M}$) | $k_{\text{cat}}$ ($\text{s}^{-1}$) | $k_{\text{cat}} / K_m$<br>($\text{nM}^{-1} \text{s}^{-1}$) | $K_i$ ($\mu\text{M}$)<sup>c</sup> |
| --- | --- | --- | --- | --- | --- |
| D-Pantothenate | | $24 \pm 4$ | $0.64 \pm 0.02$ | $30 \pm 3$ | - |
| 1a | | $100 \pm 7$ | $0.64 \pm 0.01$ | $6.6 \pm 0.4$ | - |
| 1c | | $65 \pm 7$ | $0.61 \pm 0.02$ | $10.2 \pm 0.9$ | - |
| 1e | | $42 \pm 4$ | $0.399 \pm 0.005$ | $10 \pm 1$ | - |
| 2c<sup>b</sup> | | $69 \pm 6$ | $0.63 \pm 0.02$ | $9.3 \pm 0.5$ | $2654 \pm 733$ |
| 2e<sup>b</sup> | | $67 \pm 10$ | $0.73 \pm 0.05$ | $12 \pm 1$ | $4337 \pm 961$ |
| 3f | | $75 \pm 11$ | $0.73 \pm 0.02$ | $10.7 \pm 0.8$ | - |

<sup>a</sup>Unless otherwise stated, kinetic parameters reported in the table were determined for each experiment by fitting the Michaelis-Menten equation to the data. <sup>b</sup>For these compounds, the parameters were determined by fitting the data to Equation 1 that accounts for substrate inhibition. <sup>c</sup> $K_i$ , the inhibitor constant, was determined for compounds that displayed substrate inhibition, *i.e.* a decrease in $V_{\text{max}}$ at higher substrate concentrations. Data were collected from 3 independent experiments, each performed in triplicate (errors represent SEM).

Additional experiments were performed with the same selected compounds to further investigate their antiplasmodial mechanism(s) of action. We are specifically interested in determining whether there is a correlation between the antiplasmodial activity of the compounds and their interaction with *Pf*PanK, as we have shown that the thiazole derivatives compete with pantothenate (**Table 1**, **Figure S1**). Although we would like to directly interrogate the kinetic parameters of *Pf*PanK in the presence of each compound, the low yields achievable when purifying the enzyme from *P. falciparum* prohibited us from obtaining enough protein for conventional kinetic analysis. As an alternative, we tested the phosphorylation of [^14^C]pantothenate by *Pf*PanK in *P. falciparum* lysates in the presence of each compound at 10× their respective antiplasmodial IC_50_ values (**Table 1**, **Figure 3A**). Almost complete inhibition (>90%) of pantothenate phosphorylation was observed in the presence of 10× antiplasmodial IC_50_ values of **1a**, **1c**, **1e**, **2c** and **3f**, while approximately 80% inhibition was observed for compound **2e**, which adds another line of evidence that these compounds are competing with pantothenate for direct interaction with *Pf*PanK.

**Figure 3.**
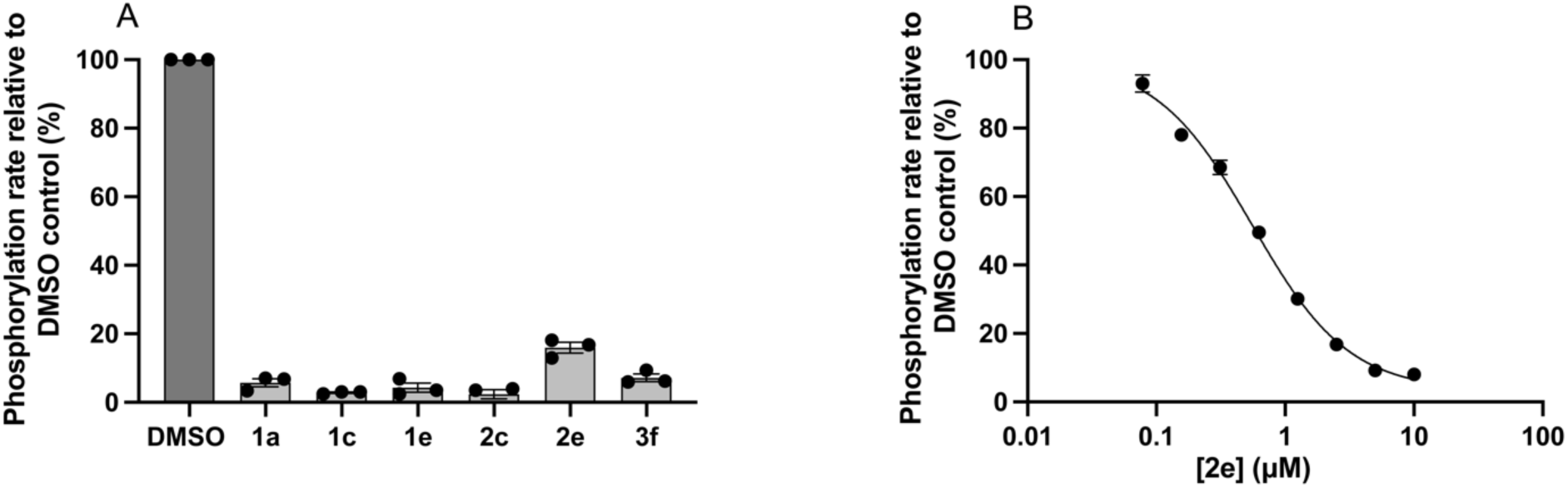
Effect of various compounds on [^14^C]pantothenate phosphorylation by *P. falciparum* lysates. [^14^C]pantothenate and Somogyi reagent were used to detect PanK activity in *P. falciparum* lysates. Compounds were tested at 10× their antiplasmodial IC_50_ values in (**A**). The effect of **2e** on [^14^C]pantothenate phosphorylation by *P. falciparum* lysates was tested at various concentrations in (**B**). Data are expressed as a percentage of the pantothenate phosphorylation rate in the DMSO (vehicle) control and are averaged from three independent experiments, each with a different batch of lysate and carried out in triplicate. Error bars represent SEM and are not visible if smaller than the symbols.

To examine the interaction of compound **2e**, the most potent antiplasmodial in this series, with *Pf*PanK in more detail, we tested the effect of various concentrations of **2e** on [^14^C]pantothenate phosphorylation. As expected, we observed a reduction in the pantothenate phosphorylation rate by *Pf*PanK (in lysates) as the concentration of **2e** was increased (**Figure 3B**). We found that **2e** was equally effective at inhibiting blood stage parasite proliferation (IC_50_ = 0.51 ± 0.02 µM; n = 3; mean ± SEM) and pantothenate phosphorylation (IC_50_ = 0.56 ± 0.03 µM; n = 3; mean ± SEM). The IC_50_ of **2e** at inhibiting pantothenate phosphorylation is approximately 2.1-fold higher than the *K*_m_ for pantothenate that has been reported previously for *Pf*PanK in parasite lysates (270 ± 95 nM; mean ± SEM) (21, 70, 71). In comparison, *Hs*PanK3 exhibits a 2.8-fold higher *K*_m_ for **2e** (67 ± 10 µM; n = 3; mean ± SEM); thus the IC_50_ of **2e** at inhibiting *Pf*PanK-mediated pantothenate phosphorylation is 120-fold lower than the *K*_m_ observed for *Hs*PanK3 (**Table 2**), consistent with **2e** interacting with much higher affinity with *Pf*PanK than *Hs*PanK3, and with the absence of toxicity to HFF cells.

### 2.4. Computational analysis of pantothenamides

The mode of action of pantothenamides may involve three bioactivation steps and several CoA-utilising enzymes, therefore small structural differences can affect permeability (across both erythrocyte and parasite membranes) as well as any of the many enzymes that are involved in bioactivation and/or serve as targets. It is therefore challenging to rationalize, for example, the preference for a phenethyl or phenyl group over the linear aliphatic chains observed for series **2** and **3** compounds. Further experiments specifically looking at cell permeability, bioactivation and target identification might help gain a better understanding of the SARs. As a complementary approach, ligand-based computational studies were used here to gain more insight into SARs. Conformational samplings and electronic charges were calculated using ORCA version 6.0 (72, 73). The results were visualized in ChimeraX (74–76) with SEQCROW (77, 78). Conformational samplings were obtained from the semiempirical tight binding (GFN2-xTB) method (79) and the analytical linearized Poisson-Boltzmann (ALPB) model (80) as an implicit solvent model of water. Electronic charge distributions were calculated based on charges from electrostatic potentials using a grid-based method (CHELPG) (81) at the B3LYP/def2-TZVPPD level of theory.

We focused on the different thiazole cores and simplified the shared structures of the compounds. Consequently, the pantoyl moiety was reduced to an acetyl group and the side chain (defined in **Figure 1**) was abbreviated with an ethyl or a phenyl substituent. Overall, both the position of the heteroatoms in the ring and the nature of the side chain were found to have a significant impact on the favored conformations and the electronic charges. For series **1** with an ethyl substituent, conformational sampling reveals that a 1,4-N···S interaction is key to the stabilization of the amide-thiazole backbone (**Figure 4a**). Taking a closer look at the global minimum of the structure, the distance between the nitrogen and sulfur atom is 2.9 Å, which is within the sum of the van der Waals radii of 3.35 Å (41). The torsion angle (φSCCN) is –13.5 degrees, which satisfies the requirement for 1,4-N···S interaction (41). Notably, the lesser negative charge on the amide nitrogen in the ethyl model (–0.41) than in the phenyl model (–0.59; **Figure 4b**) correlates with the proposal that this nitrogen serves as an electron donor in the 1,4-N···S interaction. Charge distribution analyses also show that the ring N−C−C−H moiety in series **1** with an ethyl substituent is polarized, similar to that of an amide O−C−N−H moiety. Conversely, a phenyl substituent (rather than ethyl) is detrimental in this series (**Figure 4b**). Despite slightly improving the conformational stability of the ensemble (see section 4 of the Supporting Information), installation of a phenyl group not only disrupts the 1,4-N···S interaction but also makes the N−C−C−H moiety less polar.

**Figure 4.**
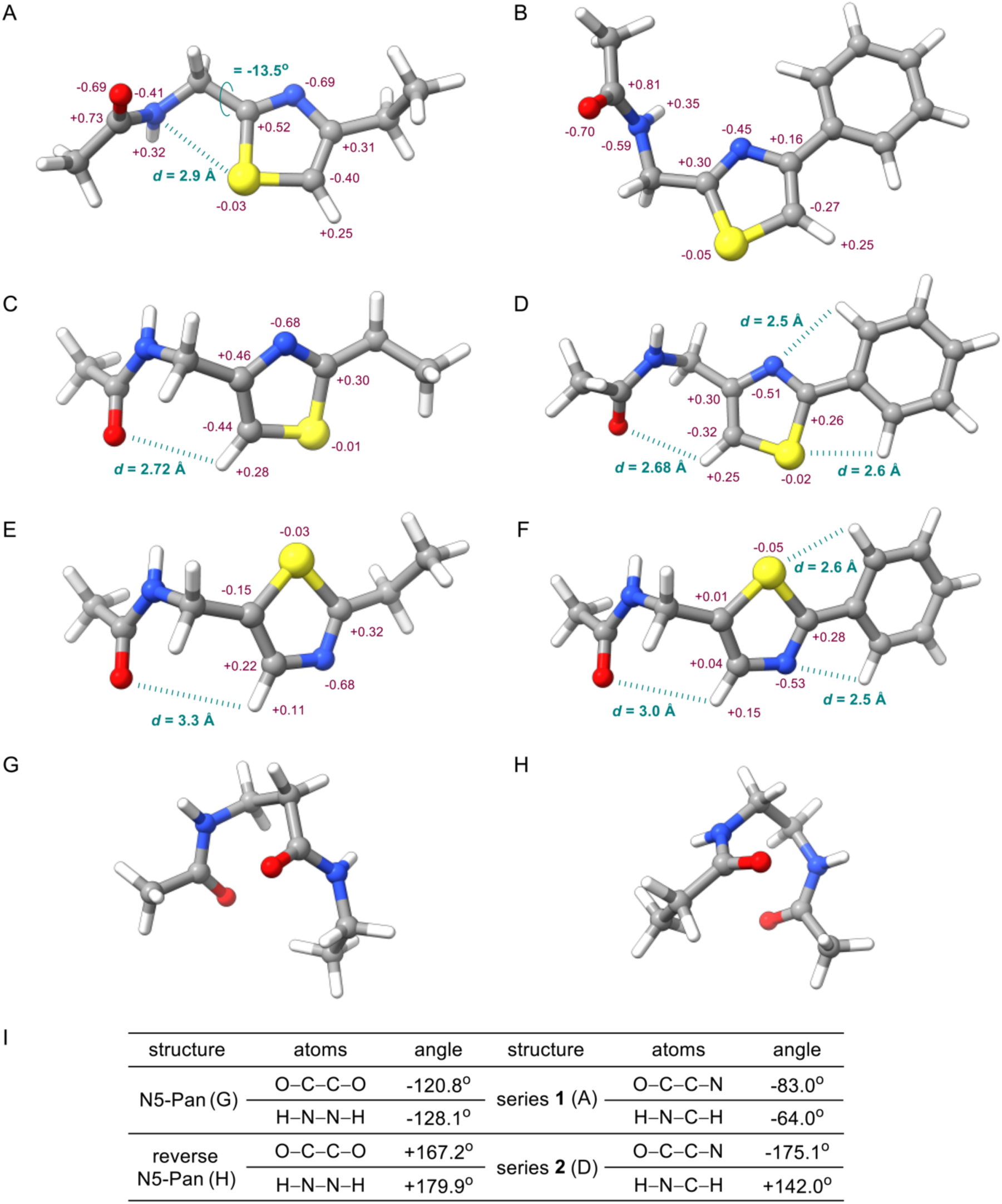
Computational results. Distances and torsion angles are highlighted in teal, and charge distributions are indicated in maroon. The global minimum conformations are shown, except for structure b, which is +0.43 kcal/mol above the global minimum. A) Truncated model structure of series **1** bearing an ethyl substituent. B) Truncated model structure of series **1** bearing a phenyl substituent. C) Truncated model structure of series **2** bearing an ethyl substituent. D) Truncated model structure of series **2** bearing a phenyl substituent. E) Truncated model structure of series **3** bearing an ethyl substituent. F) Truncated model structure of series **3** bearing a phenyl substituent. G) Truncated model structure of N5-Pan bearing an ethyl substituent. H) Truncated model structure of reverse N5-Pan bearing an ethyl substituent. I) Measured torsion angles.

As sulfur is positioned away from the amide bond in series **2**, no benefit can be gained from a 1,4-N···S interaction (**Figures 4c** and **4d**). Nevertheless, the different arrangement of the heteroatoms offers new favorable interactions. Unlike series **1**, in series **2** the introduction of a phenyl substituent (**Figure 4d**) significantly stabilizes the conformational ensemble (see section 4 of Supporting Information), consistent with the higher activity of **2e** over that of **2a**. In particular, the thiazole core and the phenyl ring are essentially coplanar across the whole ensemble, implying that they form an important interaction. Previous studies (82–84) have suggested that this phenomenon originates from unconventional hydrogen bonds between the heteroatoms and the aromatic C-H bond (C−H···N and C−H···S hydrogen bond). Indeed, the distance between N/S and H calculated here is 2.5 Å and 2.6 Å respectively, therefore within the sum of the van der Waals radii (2.75 Å and 3.0 Å) and confirming this beneficial interaction. A polarized N−C−C−H thiazole backbone is also present in series **2**, indicating its capability of electronically imitating the amide bond of pantothenamides. In addition, the distance between the amide oxygen and thiazole hydrogen is slightly below the sum of the van der Waals radii (2.72 Å), suggesting that extra factors may contribute to conformational stability. This may explain, at least in part, the better activity of series **1** and **2** relative to previously reported linear pantothenamides, such as N5-Pan for which the IC_50_ values are 124 ± 15 µM and 2.0 ± 0.5 µM in the presence and absence of pantetheinases, respectively (20).

Results with series **3** compounds somewhat resemble those of series **2**, albeit with less beneficial interactions (**Figure 4e** and **4f**), thus paralleling their lower biological activity. The 1,4-N···S interaction is absent in series **3**, and the amide oxygen and thiazole hydrogen are too far apart (3.3 Å and 3.0 Å for the ethyl and phenyl derivatives, respectively) to have any meaningful interactions. Remarkably, charge distribution analysis reveals that no polarized thiazole backbone exists in series **3**, making it electronically unfit to mimic an amide bond. These results correlate well with the fact that series **3** is generally inferior to series **1** and **2** for their antiplasmodial activity. They also tentatively offer a plausible explanation for different substituent preferences among these series of compounds. For example, the phenyl-thiazole interaction may rationalise the better activity of **3f** over that of **3a** in this series too.

Although both series **1** and **2** delivered potent antiplasmodial compounds, they had very distinctive SARs. To further explain the difference between the observed SARs, conformational samplings were also computed for the benchmark compounds N5-Pan and reverse N5-Pan. Similar reduction of molecular complexity was applied to facilitate the calculations. The results demonstrate that both compounds exhibit a U-shaped backbone to maximize amide carbonyl-carbonyl interactions, either through *n* → π^∗^ interactions or dipole-dipole interactions (85–87). Analyses of torsion angles were performed to compare amide bonds and the polarized thiazole backbones, where angles of O−C−C−O and H−N−N−H were measure for the two pantothenamides, and angles of O−C−C−N and H−N−C−H were measured in thiazole-containing pantothenamide-mimics. Gratifyingly, the results (**Figure 4i**) revealed that series **1** bears more resemblance with the normal amide, whereas series **2** with the reverse amide. This correlates well to previous findings that alkyl chains were preferred for compounds with a normal amide, whereas aryl substituents were favoured for those with a reverse amide (20, 29, 88, 89).

## 3. DISCUSSION

A goal of this study was to explore the antiplasmodial activity, specificity, and on-target activity of pantothenamide mimics which have the labile amide bond replaced with a thiazole to prevent degradation by pantetheinases. We previously had success at replacing this bond with a triazole or an isoxazole (**Figure 1B**). Preliminary results (35) suggested that a sulfur-containing heterocycle could be a promising replacement of the labile amide bond, hence we were encouraged to synthesise novel thiazole analogues. Three of the 23 compounds displayed sub-micromolar antiplasmodial activity against blood stage-*P. falciparum*.

In the pantothenamide-mimics previously reported by Spry *et al.* (30), Schalkwijk *et al.*(29), Guan *et al.* (33, 35), and de Vries *et al.* (88), a two-carbon linker is optimal between the pantoyl and the linear amide mimic, whereas a one-carbon linker was preferred between the pantoyl and the heterocyclic moieties. Therefore, the novel thiazole analogues reported here all harbor a one-carbon linker. The substituent preferences at the ring or amide nitrogen also differ between analogues containing a heterocycle and those with a linear amide. In the case of linear amide mimics, substituents such as phenyl or phenethyl are preferred over linear aliphatic groups (28–30). On the other hand, linear aliphatic chains with a length of C4 to C6 are preferred over other alkyl or aryl groups (*e.g.* 2-methyl-propyl, *p*-trifluoromethylphenyl, phenethyl groups, etc.) for molecules containing a triazole or isoxazole (32, 33, 35). In contrast, we found that phenethyl and butyl side chains give equally potent compounds in the thiazole series **1** (see **1a** and **1e**). Surprisingly, in the thiazole series **2** and **3**, the alkyl chains are no longer preferred, and a phenyl group is favoured instead (see **2a** and **2e**; **3a** and **3f**), yet substituents on the phenyl ring are detrimental. Preliminary computational studies were used to rationalize these SARs. A ligand-based approach was used considering the complex mode of action of these molecules and the several steps needed for their localisation into the parasite. The results confirm that a simple rotation of the ring or switching of two of its heteroatoms can have a dramatic effect on the preferred side chains and on the biological activity. This was rationalized by the effect of the ring on the conformational sampling of the molecule, as well as the polarity of the ring. Overall, this explains why the biological activity of pantothenamide mimics is so easily tuned, which is highly advantageous in drug development.

Kinetic analyses with *Hs*PanK3 revealed that the tested thiazole analogues are substrates of the enzyme, yet with lower specificity than pantothenate, as manifested by the higher *K*_m_ and lower *k*_cat_/*K*_m_ values. For comparison, we employed a [^14^C]pantothenate-based phosphorylation assay to investigate the effect of selected compounds on the activity of *Pf*PanK in parasite lysates. This radiolabelled assay is unable to distinguish whether a compound is a substrate or an inhibitor, but it can show whether the compound competes with pantothenate for binding to *Pf*PanK. However, based on the *Hs*PanK3 data presented above and a previous report that demonstrated that a similar triazole derivative (compound **i**, **Figure 1B**), is a *Pf*PanK substrate (21), we believe that it is likely that the compounds tested here are *Pf*PanK substrates, rather than inhibitors. We show that the most active compound (**2e**) is equally effective at inhibiting the phosphorylation of pantothenate by *Pf*PanK than at preventing the proliferation of intraerythrocytic *P. falciparum* parasites, yet we are not proposing that *Pf*PanK is the target of **2e**, or more generally, the target of this series of compounds. Instead, it is likely that the target is downstream in the CoA biosynthetic pathway and/or a CoA-utilising enzyme. This is consistent with previous reports on the triazole and reverse amide pantothenamide mimics (21, 29, 32, 88). The latter were reported to be metabolized into CoA antimetabolites, which target and inhibit acetyl-CoA synthetase (29, 88). Our data are consistent with the thiazole-containing pantothenamide-mimics sharing a similar mechanism of action as the previously reported ones.

Remarkably, our results with HFF cells suggest that the compounds may not be cytotoxic to mammalian cells. Considering the fact that the compounds are substrates for *Hs*PanK3, one explanation for the lack of toxicity against HFF cells may be that the compounds may not be able to permeate HFF cells, as previously hypothesised by Howieson *et al.* (32). Alternatively, the compounds may be phosphorylated by human PanK but not metabolised further by downstream enzymes of the CoA biosynthetic pathway in HFF cells (and/or may be rapidly hydrolyzed), as opposed to in *P. falciparum*, where their phosphorylated forms subsequently rejoin the pathway as substrates of *Pf*PPAT and *Pf*DPCK, generating antiplasmodial CoA antimetabolites (21, 22). If indeed these compounds exert their activity by inhibiting CoA-utilising enzymes, they would be ineffective in HFF cells for lack of production of the CoA antimetabolites.

We believe that replacing the labile amide group of pantothenamides with a thiazole ring is a promising strategy to identify blood-stable antiplasmodial agents targeting CoA metabolism/utilisation. Additional SARs studies, in particular at the side chain of both thiazole series **1** and **2**, and further biological testing are worthy of investigation.

## 4. MATERIALS and METHODS

### 4.1 Cell culture

Culturing of intraerythrocytic stage *P. falciparum* parasites (3D7 strain) was carried out with a method adapted from Trager and Jensen (90) and Allen and Kirk (91). Parasites were maintained within human, male or female, erythrocytes (typically Group O+, Rh+, 4% haematocrit) in RPMI-1640 medium (Thermo Fisher Scientific, catalogue number 72400120) supplemented with 25 mM HEPES, 11 mM glucose, 200 µM hypoxanthine, 24 µg/mL gentamicin and 6 g/L Albumax II on a horizontally rotating shaking incubator (New Brunswick Innova® 42/42R) at 37 °C under an atmosphere of 96% nitrogen, 3% carbon dioxide and 1% oxygen.

HFF cells were maintained *in vitro* in DMEM supplemented with 10% (v/v) bovine calf serum, 50 units/mL penicillin, 50 µg/mL streptomycin, 10 µg/mL gentamicin, 0.2 mM L-glutamine, and 0.25 µg/mL amphotericin B, as previously described (92).

### 4.2 *In vitro* parasite proliferation assay

To evaluate the effect of each compound on the *in vitro* proliferation of *P. falciparum* parasites, a SYBR Safe-based fluorescence assay was used as previously described (32). All pantothenamide-mimics tested were initially dissolved to a concentration of 200 mM in DMSO. The final DMSO concentration in the assay never exceeded 0.1% (v/v). Assays were performed using the same medium used to culture the parasites. Where specified, 100 µM sodium pantothenate (Yick-Vic Chemicals and Pharmaceuticals, Hong Kong) was added to the medium.

Assays were set up with synchronous ring-stage *P. falciparum*-infected erythrocytes in 96-well microtitre plates. Two-fold serial dilutions of the compound in complete medium were then added to the wells of each plate in triplicate (total volume of 100 µL per well), after which *P. falciparum*-infected erythrocytes were added to a final volume of 200 µL per well (0.5% parasitaemia and 1% haematocrit). *P. falciparum*-infected erythrocytes incubated in the presence of 0.5 µM chloroquine were used as a non-proliferation control and those incubated in the absence of compounds and chloroquine served as a 100% parasite proliferation control. To minimize ‘edge effects’ (93, 94), the outermost rows and columns of the microtitre plates were not used in the proliferation assays and were filled with 200 µL of medium. The plates were then incubated at 37°C for 96 h under a low-oxygen atmosphere. At the end of the incubation, plates were frozen at –80°C to lyse the cells. Following thawing, the content of each wells was resuspended in the plates by pipetting and then 100 µL from each well was transferred to a second 96-well microtitre plate containing 100 µL of lysis buffer (20 mM Tris, pH 7.5; 5 mM EDTA; 0.008% (w/v) saponin and 0.08% (v/v) Triton X-100) with SYBR Safe DNA gel stain (0.2 µL/mL) per well. The cells were gently mixed with the buffer, and the fluorescence signal in each well was then measured using a FLUOstar OPTIMA microplate reader with excitation and emission wavelengths of 490 and 520 nm, respectively, after setting the gain on a 100% parasite proliferation control well. The fluorescence in the non-proliferation control wells was subtracted from the fluorescence readings in the other wells prior to further analysis. Parasite proliferation, calculated as a percentage of the fluorescence measured in the test wells relative to the fluorescence measured in the 100% parasite proliferation control wells, was then plotted against the logarithm of the compound concentration. Using GraphPad Prism, each dataset was then fitted with a sigmoidal curve (Y=Bottom + (Top-Bottom)/(1+(IC_50_/X)^HillSlope), where Y represents percentage parasite proliferation, IC_50_ represents the concentration of the compound resulting in 50% parasite proliferation and X represents the compound concentration) using a nonlinear least squares regression. IC_50_ values (determined from the sigmoidal curves) were averaged from three independent experiments.

### 4.3 *In vitro* HFF proliferation assay

*In vitro* HFF proliferation assays were performed as previously described by Howieson *et al.* (32), with modifications. Confluent HFF cells (in a 75 cm^2^ flask) were harvested by treatment with 1× trypsin–EDTA (0.25% trypsin + 0.2 g/L EDTA) and diluted in 21 mL of DMEM supplemented with 10% newborn calf serum (32). Assays were set up in 96-well microtitre plates in a similar way to that described for the antiplasmodial activity assay. Cycloheximide (10 µM) was used as a non-proliferation control and 0.1% v/v DMSO served as a 100% proliferation control. The plates were incubated at 37°C in a CO_2_ incubator for 96 h and Incucyte® Live-Cell Analysis System (Essen BioScience) was used to capture cellular changes and assess cell growth in each well over time. Each well was scanned to capture bright field images, and confluency was assessed at the end of the 96 h incubation, which was then used to evaluate the effect of each compound on the proliferation of HFF cells by comparing to the 100% proliferation control.

### 4.4 Preparation of *P. falciparum* lysates

Parasite lysates were prepared from saponin-isolated mature trophozoite-stage parasites as described by Saliba *et al.* (14), with modifications. Briefly, a culture with predominantly trophozoite stage parasites was thoroughly mixed with saponin (0.05% (w/v) final concentration) and centrifuged at 2,000 × *g* for 8 min at 4°C. The pelleted parasites were then transferred to a 1.5 mL tube and washed three times (16,000 × *g*, 1 min each time) with cold malaria saline (125 mM NaCl, 5 mM KCl, 25 mM HEPES, 20 mM glucose, 1 mM MgCl_2_, pH 7.1). Subsequently, washed saponin-isolated parasites were suspended in 1 mL of 10 mM Tris/Cl, pH 7.4, followed by trituration (10 times) through a SafetyGlide injection needle (25-gauge). The lysates were then centrifuged at 16,000 × *g* for 30 min at 4°C and the supernatant was transferred to a new tube. The concentration of the lysates was then determined using Bradford reagent.

### 4.5 [^14^C]pantothenate phosphorylation assay

Phosphorylation of [^14^C]pantothenate was determined using a combination of zinc sulphate (ZnSO_4_) and barium hydroxide (Ba(OH)_2_) (Somogyi reagent) as previously described (71). In the 125 min time course, each reaction (650 µL) contained a final concentration of 50 mM Tris/Cl (pH 7.4), 5 mM ATP, 5 mM MgCl_2_, 2 µM (0.1 µCi/mL) [^14^C]pantothenate and 60 µg of total parasite lysate. The compounds were added at their 10× antiplasmodial IC_50_ values or at various concentrations to generate dose-response curves. Mixtures were incubated at 37°C throughout the experiment and reactions were initiated by the addition of the lysates. A reaction with identical components to the test reactions but lacking the parasite lysate was prepared to serve as a no-phosphorylation negative control and a reaction with DMSO vehicle control (0.2% v/v) served as a positive control. At pre-determined time points, 50 µL of the reaction mixture was transferred in triplicate to wells of a 96-well filter plate with 0.2 µm hydrophilic polyvinylidene fluoride (PVDF) membrane filter (Corning), which had been pre-loaded with 50 µL of 150 mM Ba(OH)_2_. The reaction mixtures were thoroughly mixed with the Ba(OH)_2_ in the wells to precipitate the enzyme and terminate the phosphorylation reaction. After all reactions had been terminated, phosphorylated compounds in each well were precipitated by the addition of 50 µL of 150 mM ZnSO_4_ to generate the Somogyi reagent. The plate was then washed twice with deionized H_2_O and once with 95% (v/v) ethanol using a vacuum manifold to remove the non-phosphorylated [^14^C]pantothenate and was subsequently dried overnight at 37°C in a non-humidified incubator. The following day, 30 µL of Microscint-O (PerkinElmer) was added to each well and the plate sealed on the top and bottom with TopSeal^TM^-A Plus sealing film (PerkinElmer). A Packard Microplate Scintillation and Luminescence Counter was used to measure the radioactivity in each well immediately after the plate had been sealed. To determine the total radioactivity in each phosphorylation assay, 50 µL of each reaction mixture was added in the wells of an OptiPlate-96 microplate (PerkinElmer) in triplicate and was then thoroughly mixed with 150 µL of Microscint-40 (PerkinElmer) and the radioactivity was measured in the same manner. Phosphorylation of [^14^C]pantothenate, calculated as a percentage of the radioactivity measured in the test wells relative to the total radioactivity in the phosphorylation assay, was then plotted against the reaction time. Each data set was analysed by linear regression, from which the phosphorylation rate of each reaction was then determined and expressed as a percentage of the phosphorylation rate of the DMSO positive control reaction.

### 4.6 *Hs*PanK protein cloning, expression and purification

The gene encoding the catalytic core domain of human PanK3 (amino acid residues 12–368) was synthesized and cloned into the pET28a(+) expression vector using the NdeI and BamHI restriction sites to produce an N-terminal His6-tagged protein (Bio Basic Inc). The plasmid was transformed into *E. coli* BL21(DE3) cells, which were cultured in Terrific Broth (TB) medium containing 50 µg/ml kanamycin at 37°C. When the culture reached an optical density at 600 nm (OD600) of 0.6, protein expression was induced by adding 1.0 mM isopropyl β-D-thiogalactoside (IPTG), followed by incubation at 18°C for 17–18 h. The cells were then harvested and resuspended in lysis buffer (20 mM Tris-HCl, pH 7.5, 500 mM NaCl, 5 mM imidazole, 5% glycerol) supplemented with 1 mM PMSF (phenylmethylsulfonyl fluoride) and lysozyme (5 mg/g of the cell pellet). The suspension was then sonicated on ice, and the resulting lysate was centrifuged twice at 54,000 × g for 30 min at 4°C.

The clarified supernatant was incubated with Ni-NTA resin pre-equilibrated with lysis buffer containing 5 mM imidazole at 4°C. After binding, the resin was washed with lysis buffer containing 30 mM imidazole, then eluted with lysis buffer containing 250 mM imidazole. The eluted *Hs*PanK3 protein was dialyzed against storage buffer (20 mM Tris-HCl, pH 7.5, 200 mM NaCl, 5% glycerol) and concentrated. The purified protein was stored at −80°C. Protein purity was assessed using SDS-PAGE and deemed >50% (**Figure S3**).

### 4.7 *Hs*PanK kinetic assays

The enzyme assay was adapted from a previously described method (95), which links the production of ADP to the consumption of NADH through the enzymatic activities of pyruvate kinase and lactic dehydrogenase. Reactions were conducted in 96-well microtiter plates, and the decrease in NADH concentration was measured at 340 nm at 30 s intervals after incubation at 30°C. Each reaction mixture (200 µL) contained the desired compound (substrate) at variable concentrations (0–2000 µM), ATP (1.5 mM), NADH (0.3 mM), phospho(enol)pyruvate (2.0 mM), MgCl_2_ (10 mM), KCl (20 mM), a commercial pyruvate kinase/lactic dehydrogenase mixture (12 units), and pantothenate kinase (0.48 µM) in 50 mM Tris–HCl buffer (pH 7.6). The reaction was initiated with the addition of ATP. The kinetic parameters were determined by fitting the rate data into the Michaelis–Menten equation using GraphPad Prism 8.0 with nonlinear regressions. Data were collected from 3 independent experiments, each performed in triplicate to verify the trends and optimize the selected concentrations. The kinetic parameters were calculated from the most complete triplicate set of data and errors represent SEM. For cases where substrate inhibition was suspected, the same conditions were used, but the data was fit using the built-in Equation 1 (Y = V_max_*X/(K_m_ + X*(1 + X/K_i_)) from GraphPad Prism, where X and Y represent substrate concentration and enzyme activity, respectively, V_max_ is the maximum enzyme activity in the same units as Y, and the K_m_ and K_i_ are the Michaelis-Menten and dissociation constants, respectively, both in the same units as X. Equation 1 is based on Equation 5.44, in RA Copeland, Enzymes, 2nd edition, Wiley, 2000.

### 4.8 Compound synthesis and characterisation

Details of chemical synthesis as well as compound characterisation are detailed in Section II (starting on page S11) of the Supporting Information. NMR spectra are also included starting on page S49.

## 5. AUTHOR CONTRIBUTIONS

All parasite, *Pf*PanK and HFF experiments were performed by X.L. Chemical compounds were synthesized by A.W.C. and C.B.L. *Hs*PanK3 was produced and assayed by M.N. Computational studies were performed by C.B.L. The manuscript was written by X.L. and C.B.L. and edited by A.W.C., M.N., K.A. and K.J.S. The project was overseen by K.A. and K.J.S.

## 6. ACKNOWLEDGEMENT

We are grateful to the Canberra Branch of the Australian Red Cross Lifeblood for the provision of red blood cells and members of the van Dooren Lab, Australian National University, for maintaining the HFF cultures. We also thank Compute Canada for providing a platform for computational studies, Dr. Kirill Levin for help with NMR at McGill, and Dr. Alexander S. Wahba and Mr. Nadim K. Saadeh for HRMS analyses at McGill. Free academic use of ORCA and ChimeraX is greatly appreciated. This work was supported by the Canadian Institute of Health Research (CIHR; grant PJT186220 to K.A. and K.J.S.).

## REFERENCES

1. World Health Organization. 2024. World malaria report 2024. Geneva.

2. Ashley EA, Pyae Phyo A, Woodrow CJ. 2018. Malaria. Lancet 391:1608–1621.

3. Sutherland CJ, Tanomsing N, Nolder D, Oguike M, Jennison C, Pukrittayakamee S, Dolecek C, Hien TT, do Rosario VE, Arez AP, Pinto J, Michon P, Escalante AA, Nosten F, Burke M, Lee R, Blaze M, Otto TD, Barnwell JW, Pain A, Williams J, White NJ, Day NP, Snounou G, Lockhart PJ, Chiodini PL, Imwong M, Polley SD. 2010. Two nonrecombining sympatric forms of the human malaria parasite Plasmodium ovale occur globally. J Infect Dis 201:1544–50.

4. Flannery EL, Chatterjee AK, Winzeler EA. 2017. Antimalarial drug discovery – approaches and progress towards new medicines. Nat Rev Microbiol 15:572.

5. Dondorp AM, Nosten F, Yi P, Das D, Phyo AP, Tarning J, Lwin KM, Ariey F, Hanpithakpong W, Lee SJ, Ringwald P, Silamut K, Imwong M, Chotivanich K, Lim P, Herdman T, An SS, Yeung S, Singhasivanon P, Day NP, Lindegardh N, Socheat D, White NJ. 2009. Artemisinin resistance in *Plasmodium falciparum* malaria. N Engl J Med 361:455–67.

6. Fairhurst RM, Dondorp AM. 2016. Artemisinin-resistant *Plasmodium falciparum* malaria. Microbiol Spectr 4.

7. Henrici RC, Namazzi R, Lima-Cooper G, Kato C, Aliwuya S, Dombrowski JG, Pratt S, Campino S, Conroy AL, Sutherland CJ, John CC, Opoka RO. 2024. Artemisinin partial resistance in Ugandan children with complicated malaria. JAMA doi:10.1001/jama.2024.22343.

8. Burrows JN, Duparc S, Gutteridge WE, van Huijsduijnen RH, Kaszubska W, Macintyre F, Mazzuri S, Mohrle JJ, Wells TNC. 2017. Erratum to: New developments in anti-malarial target candidate and product profiles. Malar J 16:151.

9. Wells TN, Hooft van Huijsduijnen R, Van Voorhis WC. 2015. Malaria medicines: a glass half full? Nat Rev Drug Discov 14:424–42.

10. Leonardi R, Zhang YM, Rock CO, Jackowski S. 2005. Coenzyme A: back in action. Prog Lipid Res 44:125–53.

11. Strauss E. 2010. Coenzyme A biosynthesis and enzymology, p 315-410. In Liu; H-W, Mander L (ed), Comprehensive natural products II – Chemistry and biology, vol 7. Elsevier.

12. Lehane AM, Marchetti RV, Spry C, van Schalkwyk DA, Teng R, Kirk K, Saliba KJ. 2007. Feedback inhibition of pantothenate kinase regulates pantothenol uptake by the malaria parasite. J Biol Chem 282:25395–405.

13. Saliba KJ, Ferru I, Kirk K. 2005. Provitamin B5 (pantothenol) inhibits growth of the intraerythrocytic malaria parasite. Antimicrob Agents Chemother 49:632–7.

14. Saliba KJ, Horner HA, Kirk K. 1998. Transport and metabolism of the essential vitamin pantothenic acid in human erythrocytes infected with the malaria parasite *Plasmodium falciparum*. J Biol Chem 273:10190–5.

15. Spry C, Saliba KJ. 2009. The human malaria parasite *Plasmodium falciparum* is not dependent on host coenzyme A biosynthesis. J Biol Chem 284:24904–13.

16. Spry C, van Schalkwyk DA, Strauss E, Saliba KJ. 2010. Pantothenate utilization by *Plasmodium* as a target for antimalarial chemotherapy. Infect Disord Drug Targets 10:200–16.

17. Spry C, Kirk K, Saliba KJ. 2008. Coenzyme A biosynthesis: an antimicrobial drug target. FEMS Microbiol Rev 32:56–106.

18. Clifton G, Bryant SR, Skinner CG. 1970. N’-(substituted) pantothenamides, antimetabolites of pantothenic acid. Arch Biochem Biophys 137:523–8.

19. Moolman WJ, de Villiers M, Strauss E. 2014. Recent advances in targeting coenzyme A biosynthesis and utilization for antimicrobial drug development. Biochem Soc Trans 42:1080–6.

20. Spry C, Macuamule C, Lin Z, Virga KG, Lee RE, Strauss E, Saliba KJ. 2013. Pantothenamides are potent, on-target inhibitors of *Plasmodium falciparum* growth when serum pantetheinase is inactivated. PLoS One 8:e54974.

21. Tjhin ET, Spry C, Sewell AL, Hoegl A, Barnard L, Sexton AE, Siddiqui G, Howieson VM, Maier AG, Creek DJ, Strauss E, Marquez R, Auclair K, Saliba KJ. 2018. Mutations in the pantothenate kinase of Plasmodium falciparum confer diverse sensitivity profiles to antiplasmodial pantothenate analogues. PLoS Pathog 14:e1006918.

22. de Villiers M, Spry C, Macuamule CJ, Barnard L, Wells G, Saliba KJ, Strauss E. 2017. Antiplasmodial mode of action of pantothenamides: pantothenate kinase serves as a metabolic activator not as a target. ACS Infect Dis 3:527–541.

23. Jansen PA, Hermkens PH, Zeeuwen PL, Botman PN, Blaauw RH, Burghout P, van Galen PM, Mouton JW, Rutjes FP, Schalkwijk J. 2013. Combination of pantothenamides with vanin inhibitors as a novel antibiotic strategy against gram-positive bacteria. Antimicrob Agents Chemother 57:4794–800.

24. Akinnusi TO, Vong K, Auclair K. 2011. Geminal dialkyl derivatives of N-substituted pantothenamides: synthesis and antibacterial activity. Bioorg Med Chem 19:2696–706.

25. Hoegl A, Darabi H, Tran E, Awuah E, Kerdo ES, Habib E, Saliba KJ, Auclair K. 2014. Stereochemical modification of geminal dialkyl substituents on pantothenamides alters antimicrobial activity. Bioorg Med Chem Lett 24:3274–7.

26. Guan J, Hachey M, Puri L, Howieson V, Saliba KJ, Auclair K. 2016. A cross-metathesis approach to novel pantothenamide derivatives. Beilstein J Org Chem 12:963–8.

27. Barnard L, Mostert KJ, van Otterlo WAL, Strauss E. 2018. Developing pantetheinase-resistant pantothenamide antibacterials: structural modification impacts on PanK Interaction and mode of action. ACS Infect Dis 4:736–743.

28. de Villiers M, Macuamule C, Spry C, Hyun YM, Strauss E, Saliba KJ. 2013. Structural modification of pantothenamides counteracts degradation by pantetheinase and improves antiplasmodial activity. ACS Med Chem Lett 4:784–9.

29. Schalkwijk J, Allman EL, Jansen PAM, de Vries LE, Verhoef JMJ, Jackowski S, Botman PNM, Beuckens-Schortinghuis CA, Koolen KMJ, Bolscher JM, Vos MW, Miller K, Reeves SA, Pett H, Trevitt G, Wittlin S, Scheurer C, Sax S, Fischli C, Angulo-Barturen I, Jimenez-Diaz MB, Josling G, Kooij TWA, Bonnert R, Campo B, Blaauw RH, Rutjes F, Sauerwein RW, Llinas M, Hermkens PHH, Dechering KJ. 2019. Antimalarial pantothenamide metabolites target acetyl-coenzyme A biosynthesis in *Plasmodium falciparum*. Sci Transl Med 11.

30. Spry C, Barnard L, Kok M, Powell AK, Mahesh D, Tjhin ET, Saliba KJ, Strauss E, de Villiers M. 2020. Toward a stable and potent coenzyme A-targeting antiplasmodial agent: structure-activity relationship studies of *N*-Phenethyl-α-methyl-pantothenamide. ACS Infect Dis 6:1844–1854.

31. Pett HE, Jansen PA, Hermkens PH, Botman PN, Beuckens-Schortinghuis CA, Blaauw RH, Graumans W, van de Vegte-Bolmer M, Koolen KM, Rutjes FP, Dechering KJ, Sauerwein RW, Schalkwijk J. 2015. Novel pantothenate derivatives for anti-malarial chemotherapy. Malar J 14:169.

32. Howieson VM, Tran E, Hoegl A, Fam HL, Fu J, Sivonen K, Li XX, Auclair K, Saliba KJ. 2016. Triazole substitution of a labile amide bond stabilizes pantothenamides and improves their antiplasmodial potency. Antimicrob Agents Chemother 60:7146–7152.

33. Guan J, Tjhin ET, Howieson VM, Kittikool T, Spry C, Saliba KJ, Auclair K. 2018. Structure-activity relationships of antiplasmodial pantothenamide analogues reveal a new way by which triazoles mimic amide bonds. ChemMedChem 13:2677–2683.

34. Jansen PAM, van der Krieken DA, Botman PNM, Blaauw RH, Cavina L, Raaijmakers EM, de Heuvel E, Sandrock J, Pennings LJ, Hermkens PHH, Zeeuwen P, Rutjes F, Schalkwijk J. 2019. Stable pantothenamide bioisosteres: novel antibiotics for Gram-positive bacteria. J Antibiot (Tokyo) 72:682–692.

35. Guan J, Spry C, Tjhin ET, Yang P, Kittikool T, Howieson VM, Ling H, Starrs L, Duncan D, Burgio G, Saliba KJ, Auclair K. 2021. Exploring heteroaromatic rings as a replacement for the labile amide of antiplasmodial pantothenamides. J Med Chem 64:4478–4497.

36. Gomtsyan A. 2012. Heterocycles in drugs and drug discovery. Chemistry of Heterocyclic Compounds 48:7–10.

37. Bhutani P, Joshi G, Raja N, Bachhav N, Rajanna PK, Bhutani H, Paul AT, Kumar R. 2021. U.S. FDA approved drugs from 2015-June 2020: a perspective. J Med Chem 64:2339–2381.

38. Meanwell NA. 2017. A synopsis of the properties and applications of heteroaromatic rings in medicinal chemistry, p 245-361, Advances in Heterocyclic Chemistry, vol 123. Academic Press.

39. Ilardi EA, Vitaku E, Njardarson JT. 2014. Data-mining for sulfur and fluorine: an evaluation of pharmaceuticals to reveal opportunities for drug design and discovery. J Med Chem 57:2832–42.

40. Feng M, Tang B, Liang SH, Jiang X. 2016. Sulfur containing scaffolds in drugs: synthesis and application in medicinal chemistry. Curr Top Med Chem 16:1200–16.

41. Beno BR, Yeung KS, Bartberger MD, Pennington LD, Meanwell NA. 2015. A survey of the role of noncovalent sulfur interactions in drug design. J Med Chem 58:4383–438.

42. Ayati A, Emami S, Asadipour A, Shafiee A, Foroumadi A. 2015. Recent applications of 1,3-thiazole core structure in the identification of new lead compounds and drug discovery. Eur J Med Chem 97:699–718.

43. Pasqualotto AC, Thiele KO, Goldani LZ. 2010. Novel triazole antifungal drugs: focus on isavuconazole, ravuconazole and albaconazole. Curr Opin Investig Drugs 11:165–74.

44. Fox LM, Saravolatz LD. 2005. Nitazoxanide: a new thiazolide antiparasitic agent. Clin Infect Dis 40:1173–80.

45. Jang HC, Choi SM, Kim HK, Kim SE, Kang SJ, Park KH, Ryu PY, Lee TH, Kim YR, Rhee JH, Jung SI, Choy HE. 2014. *In vivo* efficacy of the combination of ciprofloxacin and cefotaxime against *Vibrio vulnificus* sepsis. PLoS One 9:e101118.

46. Hunt JT. 2009. Discovery of ixabepilone. Mol Cancer Ther 8:275–81.

47. Eagling VA, Back DJ, Barry MG. 1997. Differential inhibition of cytochrome P450 isoforms by the protease inhibitors, ritonavir, saquinavir and indinavir. Br J Clin Pharmacol 44:190–4.

48. Lan CB, Auclair K. 2023. 1,5,7-Triazabicyclo[4.4.0]dec-5-ene: an effective catalyst for amide formation by lactone aminolysis. J Org Chem 88:10086–10095.

49. Zhou W, Ni SY, Mei HB, Han JL, Pan Y. 2015. Cyclization reaction of allylbenzothioamide for direct construction of thiazole and thiazoline. Tetrahedron Letters 56:4128–4130.

50. Mykhailiuk PK. 2019. Saturated bioisosteres of benzene: where to go next? Org Biomol Chem 17:2839–2849.

51. Lomer TR. 1963. Crystal and molecular structure of lauric acid (form A1). Acta Crystallographica 16:984-&.

52. Ennis HL, Lubin M. 1964. Cycloheximide: aspects of inhibition of protein synthesis in mammalian cells. Science 146:1474–6.

53. Schneider-Poetsch T, Ju J, Eyler DE, Dang Y, Bhat S, Merrick WC, Green R, Shen B, Liu JO. 2010. Inhibition of eukaryotic translation elongation by cycloheximide and lactimidomycin. Nat Chem Biol 6:209–217.

54. Brand LA, Strauss E. 2005. Characterization of a new pantothenate kinase isoform from *Helicobacter pylori*. J Biol Chem 280:20185–8.

55. Yun M, Park CG, Kim JY, Rock CO, Jackowski S, Park HW. 2000. Structural basis for the feedback regulation of *Escherichia coli* pantothenate kinase by coenzyme A. J Biol Chem 275:28093–9.

56. Das S, Kumar P, Bhor V, Surolia A, Vijayan M. 2006. Invariance and variability in bacterial PanK: a study based on the crystal structure of *Mycobacterium tuberculosis* PanK. Acta Crystallogr D Biol Crystallogr 62:628–38.

57. Yang K, Eyobo Y, Brand LA, Martynowski D, Tomchick D, Strauss E, Zhang H. 2006. Crystal structure of a type III pantothenate kinase: insight into the mechanism of an essential coenzyme A biosynthetic enzyme universally distributed in bacteria. J Bacteriol 188:5532–40.

58. Hong BS, Yun MK, Zhang YM, Chohnan S, Rock CO, White SW, Jackowski S, Park HW, Leonardi R. 2006. Prokaryotic type II and type III pantothenate kinases: The same monomer fold creates dimers with distinct catalytic properties. Structure 14:1251–61.

59. Nicely NI, Parsonage D, Paige C, Newton GL, Fahey RC, Leonardi R, Jackowski S, Mallett TC, Claiborne A. 2007. Structure of the type III pantothenate kinase from *Bacillus anthracis* at 2.0 A resolution: implications for coenzyme A-dependent redox biology. Biochemistry 46:3234–45.

60. Hong BS, Senisterra G, Rabeh WM, Vedadi M, Leonardi R, Zhang YM, Rock CO, Jackowski S, Park HW. 2007. Crystal structures of human pantothenate kinases. Insights into allosteric regulation and mutations linked to a neurodegeneration disorder. J Biol Chem 282:27984–93.

61. Li B, Tempel W, Smil D, Bolshan Y, Schapira M, Park HW. 2013. Crystal structures of *Klebsiella pneumoniae* pantothenate kinase in complex with N-substituted pantothenamides. Proteins 81:1466–72.

62. Franklin MC, Cheung J, Rudolph MJ, Burshteyn F, Cassidy M, Gary E, Hillerich B, Yao ZK, Carlier PR, Totrov M, Love JD. 2015. Structural genomics for drug design against the pathogen *Coxiella burnetii*. Proteins 83:2124–36.

63. Tjhin ET, Howieson VM, Spry C, van Dooren GG, Saliba KJ. 2021. A novel heteromeric pantothenate kinase complex in apicomplexan parasites. PLoS Pathog 17:e1009797.

64. Leonardi R, Zhang YM, Yun MK, Zhou R, Zeng FY, Lin W, Cui J, Chen T, Rock CO, White SW, Jackowski S. 2010. Modulation of pantothenate kinase 3 activity by small molecules that interact with the substrate/allosteric regulatory domain. Chem Biol 17:892–902.

65. Yao J, Subramanian C, Rock CO, Jackowski S. 2019. Human pantothenate kinase 4 is a pseudo-pantothenate kinase. Protein Sci 28:1031–1047.

66. Reed MC, Lieb A, Nijhout HF. 2010. The biological significance of substrate inhibition: a mechanism with diverse functions. Bioessays 32:422–9.

67. Wu B. 2011. Substrate inhibition kinetics in drug metabolism reactions. Drug Metab Rev 43:440–56.

68. Yoshino M, Murakami K. 2015. Analysis of the substrate inhibition of complete and partial types. Springerplus 4:292.

69. Kokkonen P, Beier A, Mazurenko S, Damborsky J, Bednar D, Prokop Z. 2021. Substrate inhibition by the blockage of product release and its control by tunnel engineering. RSC Chem Biol 2:645–655.

70. Saliba KJ, Kirk K. 2001. H+-coupled pantothenate transport in the intracellular malaria parasite. J Biol Chem 276:18115–21.

71. Spry C, Saliba KJ, Strauss E. 2014. A miniaturized assay for measuring small molecule phosphorylation in the presence of complex matrices. Anal Biochem 451:76–8.

72. Neese F. 2012. The ORCA program system. Wiley Interdisciplinary Reviews: Computational Molecular Science 2:73–78.

73. Neese F. 2022. Software update: The ORCA program system—Version 5.0. Wiley Interdisciplinary Reviews: Computational Molecular Science 12.

74. Pettersen EF, Goddard TD, Huang CC, Meng EC, Couch GS, Croll TI, Morris JH, Ferrin TE. 2021. UCSF ChimeraX: Structure visualization for researchers, educators, and developers. Protein Sci 30:70–82.

75. Goddard TD, Huang CC, Meng EC, Pettersen EF, Couch GS, Morris JH, Ferrin TE. 2018. UCSF ChimeraX: meeting modern challenges in visualization and analysis. Protein Sci 27:14–25.

76. Meng EC, Goddard TD, Pettersen EF, Couch GS, Pearson ZJ, Morris JH, Ferrin TE. 2023. UCSF ChimeraX: tools for structure building and analysis. Protein Sci 32:e4792.

77. Schaefer AJ, Ingman VM, Wheeler SE. 2021. SEQCROW: A ChimeraX bundle to facilitate quantum chemical applications to complex molecular systems. J Comput Chem 42:1750–1754.

78. Ingman VM, Schaefer AJ, Andreola LR, Wheeler SE. 2021. QChASM: Quantum chemistry automation and structure manipulation. Wiley Interdisciplinary Reviews: Computational Molecular Science 11.

79. Bannwarth C, Ehlert S, Grimme S. 2019. GFN2-xTB-an accurate and broadly parametrized self-consistent tight-binding quantum chemical method with multipole electrostatics and density-dependent dispersion contributions. J Chem Theory Comput 15:1652–1671.

80. Ehlert S, Stahn M, Spicher S, Grimme S. 2021. Robust and efficient implicit solvation model for fast semiempirical methods. J Chem Theory Comput 17:4250–4261.

81. CM B, KB. W. 1990. Determining atom-centered monopoles from molecular electrostatic potentials. The need for high sampling density in formamide conformational analysis. Journal of Computational Chemistry 11:361–373.

82. Grabowski SJ. 2004. Hydrogen bonding strength—measures based on geometric and topological parameters. Journal of Physical Organic Chemistry 17:18–31.

83. Domagala M, Grabowski SJ. 2005. C-H…N and C-H…S hydrogen bonds--influence of hybridization on their strength. J Phys Chem A 109:5683–8.

84. Castro M, Nicolas-Vazquez I, Zavala JI, Sanchez-Viesca F, Berros M. 2007. Theoretical study of intramolecular, CH [Formula: see text] X (X = N, O, Cl), hydrogen bonds in thiazole derivatives. J Chem Theory Comput 3:681–8.

85. Paulini R, Muller K, Diederich F. 2005. Orthogonal multipolar interactions in structural chemistry and biology. Angew Chem Int Ed Engl 44:1788–805.

86. Fischer FR, Wood PA, Allen FH, Diederich F. 2008. Orthogonal dipolar interactions between amide carbonyl groups. Proc Natl Acad Sci U S A 105:17290–4.

87. Choudhary A, Gandla D, Krow GR, Raines RT. 2009. Nature of amide carbonyl--carbonyl interactions in proteins. J Am Chem Soc 131:7244–6.

88. de Vries LE, Jansen PAM, Barcelo C, Munro J, Verhoef JMJ, Pasaje CFA, Rubiano K, Striepen J, Abla N, Berning L, Bolscher JM, Demarta-Gatsi C, Henderson RWM, Huijs T, Koolen KMJ, Tumwebaze PK, Yeo T, Aguiar ACC, Angulo-Barturen I, Churchyard A, Baum J, Fernandez BC, Fuchs A, Gamo FJ, Guido RVC, Jimenez-Diaz MB, Pereira DB, Rochford R, Roesch C, Sanz LM, Trevitt G, Witkowski B, Wittlin S, Cooper RA, Rosenthal PJ, Sauerwein RW, Schalkwijk J, Hermkens PHH, Bonnert RV, Campo B, Fidock DA, Llinas M, Niles JC, Kooij TWA, Dechering KJ. 2022. Preclinical characterization and target validation of the antimalarial pantothenamide MMV693183. Nat Commun 13:2158.

89. Josephus Schalkwijk, Pedro Harold Han Hermkens, Koen Jakob Dechering, Bonnert RV. 2020. Pantothenamide analoguesWO2020141155A1.

90. Trager W, Jensen JB. 1976. Human malaria parasites in continuous culture. Science 193:673–5.

91. Allen RJ, Kirk K. 2010. *Plasmodium falciparum* culture: the benefits of shaking. Mol Biochem Parasitol 169:63–5.

92. Jacot; D, Meissner; M, Sheiner; L, Soldati-Favre; D, Striepen B. 2014. Genetic manipulation of *toxoplasma gondii*, p 577-611. In Weiss; LM, Kim K (ed), Toxoplasma gondii (Second Edition) The Model Apicomplexan – Perspectives and Methods. Elsevier Academic Press, Burlington.

93. Lundholt BK, Scudder KM, Pagliaro L. 2003. A simple technique for reducing edge effect in cell-based assays. J Biomol Screen 8:566–70.

94. Johnson JD, Dennull RA, Gerena L, Lopez-Sanchez M, Roncal NE, Waters NC. 2007. Assessment and continued validation of the malaria SYBR green I-based fluorescence assay for use in malaria drug screening. Antimicrob Agents Chemother 51:1926–33.

95. Vong K, Tam IS, Yan X, Auclair K. 2012. Inhibitors of aminoglycoside resistance activated in cells. ACS Chem Biol 7:470–5.

