## Supplementary Information for "Thiazole substitution of a labile amide bond - a new option towards stable pantothenamide-mimics"

I. Biological Results ………………………………………………………………… .S2

II. Synthesis and compounds characterization …………………………………………..S11

### I. Biological Results

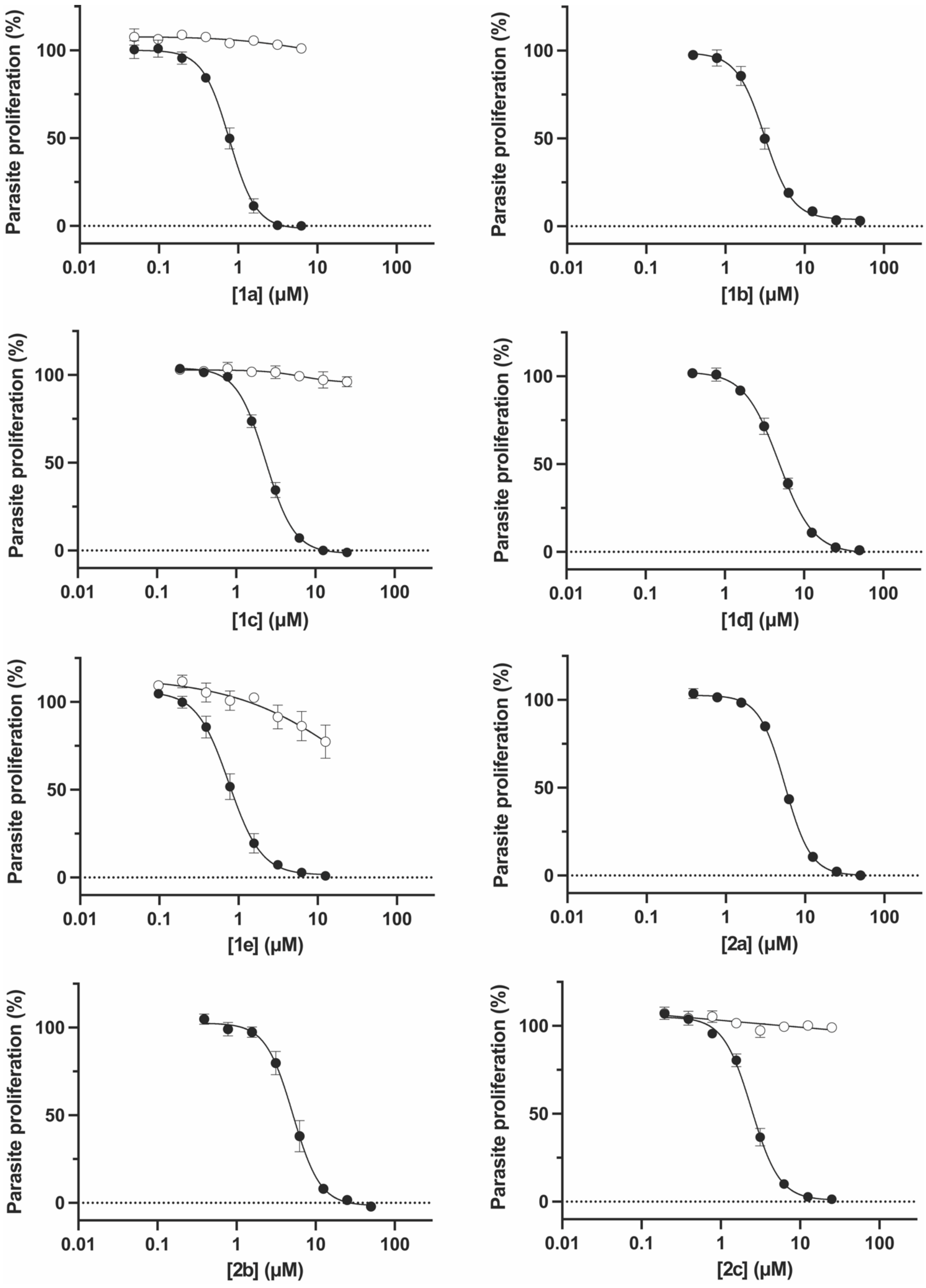

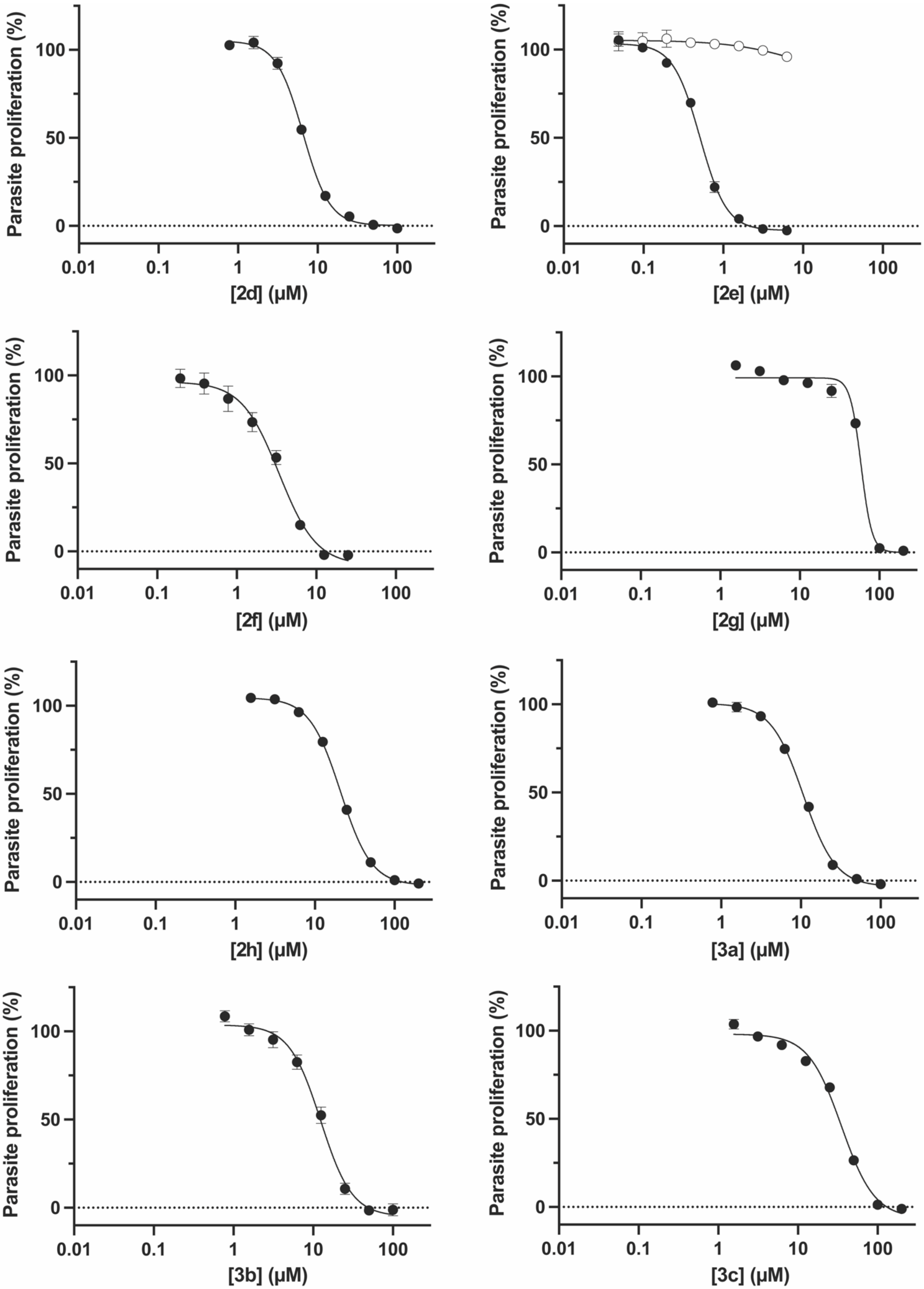
Parasite proliferation (%)
100
50
0
0.01
0.1
1
10
100
[2d] (µM)
[2e] (µM)
[2f] (µM)
[2g] (µM)
[2h] (µM)
[3a] (µM)
[3b] (µM)
[3c] (µM)

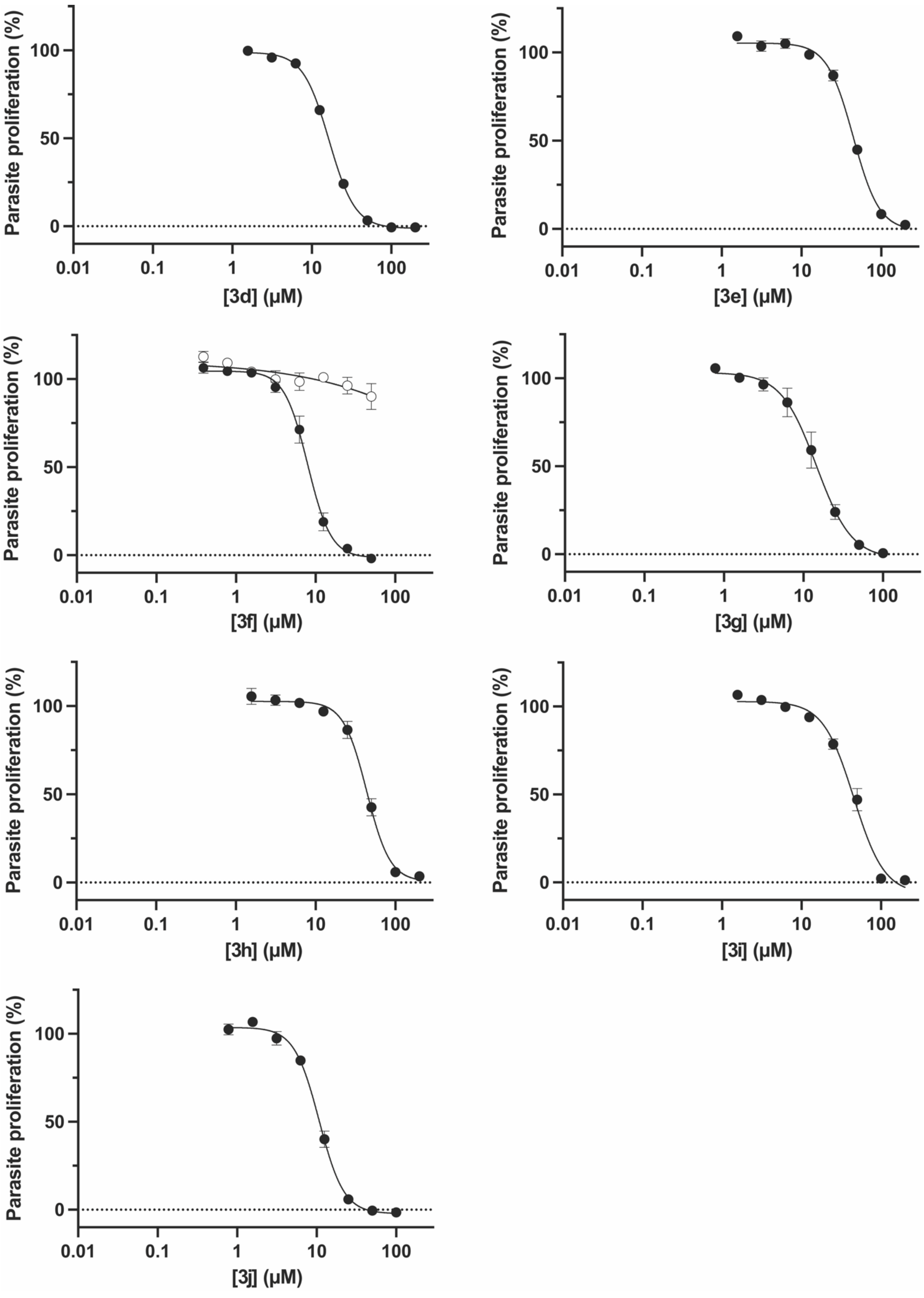
Parasite proliferation (%)
100
50
0
0.01
0.1
1
10
100
[3d] (μM)
[3e] (μM)
[3f] (μM)
[3g] (μM)
[3h] (μM)
[3i] (μM)
[3j] (μM)

**Figure S1. The effect of thiazole-bearing pantothenamide mimics on the proliferation of *P. falciparum*.** The effect of the compounds on the proliferation of *P. falciparum* 3D7 parasites in the presence of 1 μM (black circles) or, if carried out, 100 μM pantothenate (white circles). Values are averaged from three independent experiments, each carried out in triplicate. Error bars represent SEM and where not visible, are smaller than the symbols.

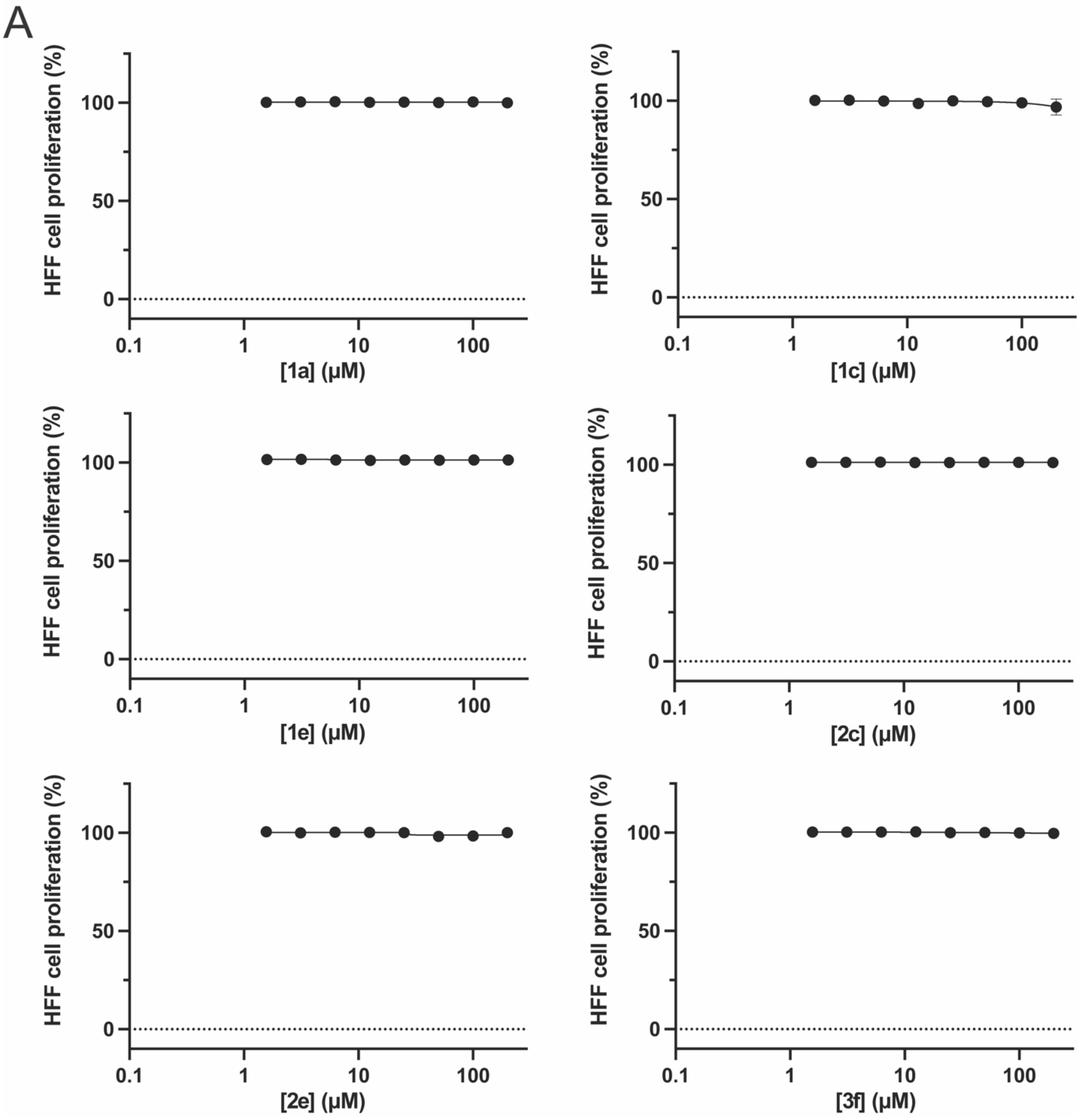
A
HFF cell proliferation (%)
100
50
0
0.1
1
10
100
[1a] (µM)
[1c] (µM)
[1e] (µM)
[2c] (µM)
[2e] (µM)
[3f] (µM)

B

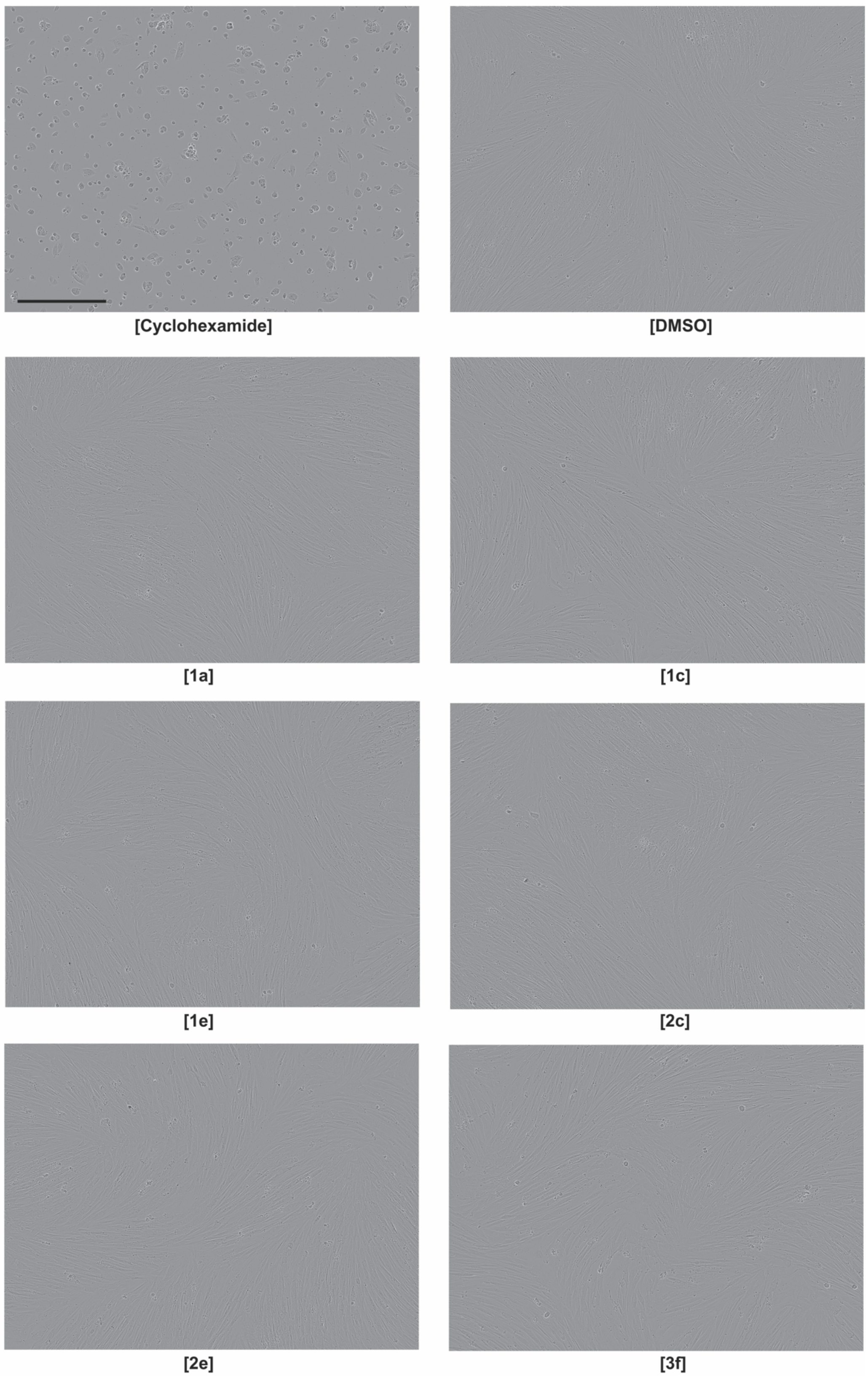

**Figure S2. The effect of thiazole-bearing pantothenamide mimics on the proliferation of HFF cells**. The effect of the compounds on the proliferation of HFF cells are shown in A. Values are averaged from 2 independent experiments, each carried out in triplicate. Error bars represent range/2 and where not visible, are smaller than the symbols. Microscopy images of HFF cells incubated with a concentration of 200 μM compounds are shown in B. The effect of 10 μM cyclohexamide and DMSO (vehicle control) on the proliferation of HFF cells is also shown (top two images). Images are captured by Incucyte® Live-Cell Analysis System and are representative of 2 independent experiments. Scale bar represents 400 μm.

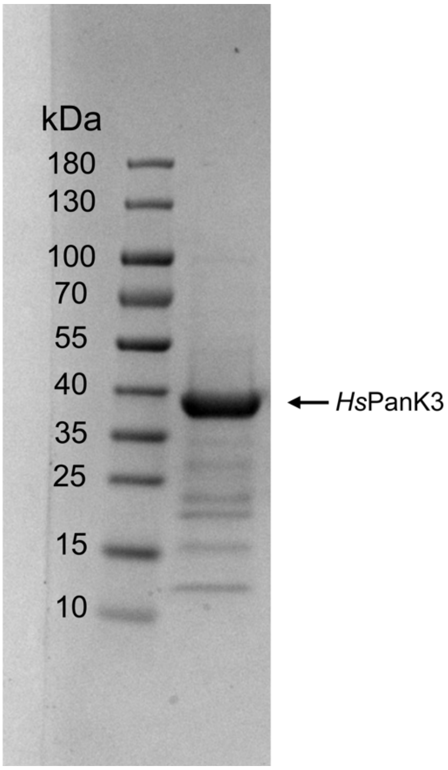


**Figure S3. SDS-PAGE gel analysis of eluted *Hs*PanK protein following Ni-NTA purification.** The right lane contains an aliquot of the purified protein (predicted weight of 41.6 kDa including the N-terminal $His_6$-tag and a thrombin cleavage site that have not been removed). The protein ladder is shown on the left, with molecular weights indicated.

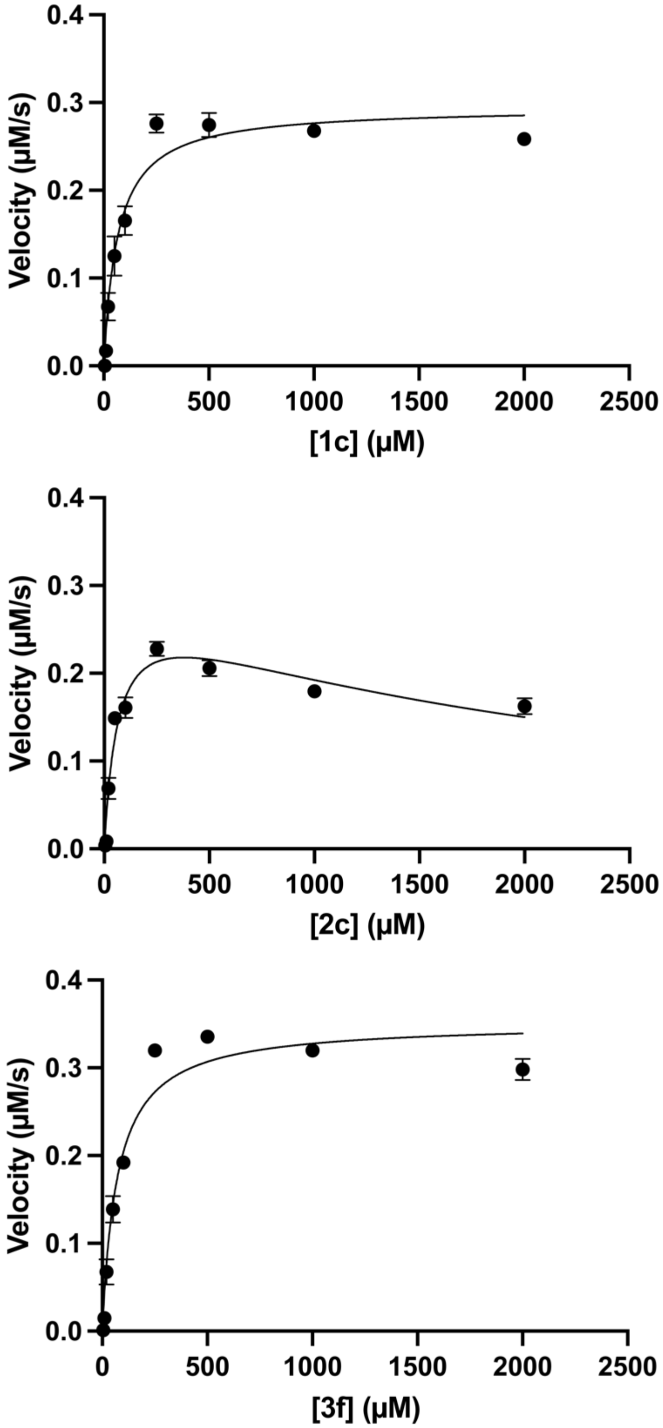


**Figure S4. Kinetics plots for *Hs*PanK3 in the presence of 1c, 2c or 3f.** Velocities were determined as a function of compound concentration. For **1c** and **3f**, the data points were fit to the Michaelis–Menten nonlinear regression equation to determine kinetic parameters. For **2c**, Equation 1, which accounts for substrate inhibition, was used to fit the data. Data are averaged from 3 independent experiments, each performed in triplicate. Error bars represent SEM and are not visible if smaller than the symbols.

#### II. Synthesis and compounds characterization

##### 1. List of abbreviations

| | |
|---|---|
| DCM | dichloromethane |
| DMF | *N,N*-dimethylformamide |
| DMSO | dimethyl sulfoxide |
| EA | ethyl acetate |
| Hex | hexane |
| NBS | *N*-bromosuccinimide |
| Phth | phthalimide |
| Rf | TLC retention factor |
| TBD | 1,5,7-triazabicyclo[4.4.0]-dec-5-ene |
| TEA | triethylamine |
| TFA | trifluoroacetic acid |
| THF | tetrahydrofuran |
| rt | room temperature |

##### 2. General Considerations

For reactions that required heating, the reaction vessel was immersed in an oil bath with digital temperature control. Room temperature is between 22 and 23$^{o}$C. Analytical thin-layer chromatography (TLC) was carried out using 200 μm aluminum-backed silica gel plates with the F-254 indicator, purchased from Silicycle (TLA-R10011B-323). Visualization of the compounds on TLC plates was accomplished under UV light (254 nm) and/or with a potassium permanganate stain. Product purification was achieved by flash chromatography on a Biotage Isolera One using 40-63 μm silica gel purchased from Silicycle (R12030B). Unless otherwise noted, reagents were purchased from commercial sources (Sigma-Aldrich, Thermo Scientific, Oakwood, Combi-Blocks), and used as received. Proton nuclear magnetic resonance ($^{1}$H-NMR), proton-decoupled carbon nuclear magnetic resonance ($^{13}$C-NMR), and proton-decoupled fluorine nuclear magnetic resonance ($^{19}$F-NMR) spectra were recorded on a Bruker AVIIIHD 500 MHz spectrometer at 298 K. Chemical shifts ($\delta$) for $^{1}$H-NMR are reported in parts per million (ppm) and referenced to residual protons in the solvent, $CDCl_3$ ($\delta$ = 7.26 ppm), DMSO-$d_6$ ($\delta$ = 2.50 ppm), or $CD_3OD$ ($\delta$ = 3.31 ppm). Chemical shifts ($\delta$) for $^{13}$C-NMR are reported in ppm and referenced to the carbon signal of the solvent, $CDCl_3$ ($\delta$ = 77.16 ppm), DMSO-$d_6$ ($\delta$ = 39.52 ppm), $CD_3OD$ ($\delta$ = 49.00 ppm). NMR splitting patterns are reported as singlet (s), broad singlet (bs), doublet (d), triplet (t), quartet (q), and quintet (p). High resolution mass spectra (HR-MS) were obtained by electrospray ionization (ESI) on a Bruker Maxis atmospheric pressure ionization (API) Quadruple Time-of-

Flight (QTOF) mass spectrometer. Purity of all final compounds was determined on an Agilent 1100 series HPLC system using a Phenomenex Luna® LC column (C18, 200 × 4.6 mm, 100 Å, 5 µm), with water as mobile phase A and acetonitrile as mobile phase B. The detector was set to 214 nm and the flow rate was 1 mL/min. All changes in solvent ratio during elution were linear. Elution method A: 0-5 min: 1% B in A, 5-15 min: 1% to 50% B, 15-20 min: 50% B, 20-25 min: 50% to 99% B, 30-32 min: 99% to 1% B, and 32-35 min: 1% B. Elution method B: 0-10 min: 1% to 99% B in A, 10-20 min: 99% B, 20-23 min: 99% to 1% B, and 35-25 min: 1% B. Elution method C: 0-3 min: 1% B in A, 3-5 min: 1% to 10% B, 5-10 min: 10% to 15% B, 10-13 min: 15% to 30% B, 13-26 min: 30% B, 26-30 min: 30% to 99% B, 30-32 min: 99% B, 32-35 min: 99% B to 1% B, and 35-38 min: 1% B. Elution method D: 0-8 min: 1% B in A, 8-15 min: 1% to 50% B, 15-25 min: 50% B, 25-30 min: 50% to 99% B, 30-38 min: 99% B, 38-40 min: 99% to 1% B, and 40-43 min: 1% B.

#### 3. Synthesis and characterization of intermediates and final compounds

General procedure A: Synthesis of a-bromo ketone

To a 50 mL round bottom flask were added the corresponding ketone (1.0 eq.) and MeOH (0.7 mL/mmol). The solution was cooled to 0°C, and $Br_2$ (1.0 eq.) was added dropwise. The reaction mixture was stirred at 0°C for 45 minutes, and then at rt for 45 minutes. Water (0.25 mL/mmol) was added followed by conc. $H_2SO_4$ (0.52 mL/mmol). The mixture was stirred at rt overnight. Water (1 mL/mmol) was then added, and the product was extracted in diethyl ether (4 × 0.8 mL/mmol). The combined organic layers were washed with sat. $NaHCO_3$ (1 mL/mmol), water (2 × 1 mL/mmol), and dried over anhydrous $MgSO_4$. The solvent was evaporated under vacuum to give the desired product, which was used in the next step without purification.

General procedure B: Hantzsch thiazole synthesis

General procedure $B_1$: Hantzsch thiazole synthesis with *tert*-butyl (2-amino-2-thioxoethyl) carbamate

To a 25 mL round bottom flask were added *tert*-butyl (2-amino-2-thioxoethyl)carbamate (1.0 eq), the α-bromo ketone (1.0 eq.), and EtOH (3 mL/mmol). The reaction mixture was stirred at rt under $N_2$ overnight. Water (4 mL/mmol) was then added to the mixture, followed by dropwise addition of aqueous $NH_3$ (28%) to reach pH ~ 8. The product was extracted in DCM (3 × 7 mL/mmol). The combined organic layers were dried over anhydrous $MgSO_4$ and the solvent was evaporated under vacuum. The residue was purified by flash chromatography to give the desired product.

General procedure $B_2$: Hantzsch thiazole synthesis with 2-(3-bromo-2-oxopropyl)isoindoline-1,3-dione

To a 25 mL round bottom flask were added 2-(3-bromo-2-oxopropyl)isoindoline-1,3-dione (1.0 eq), the thioamide (1.0 eq.), and EtOH (1.5 mL/mmol). The reaction mixture was heated to reflux under $N_2$ for 4 hours. The mixture was cooled to rt, and water (2 mL/mmol) was then added to the mixture, followed by dropwise addition of aqueous $NH_3$ (28%) to reach pH ~ 8. The precipitate was collected by filtration and dried under vacuum before purification by flash chromatography to give the desired product.

General procedure C: Deprotection of Boc groups

To a 25 mL round bottom flask were added the Boc-protected amine (1.0 eq.) in DCM (4 mL/mmol) and TFA (10.0 eq). The reaction mixture was stirred at rt for 3 hours. Then, 15% aqueous NaOH was added dropwise to reach pH ~11. The product was extracted in ethyl acetate (3 × 10 mL/mmol) and the combined organic layers were dried over $MgSO_4$. The solvent was evaporated under vacuum to give the desired product, which was used in the next step without purification.

General procedure D: Thermal ring-opening of D-pantolactone

To a Biotage® 5 mL microwave vial were added the corresponding amine (1.0 eq.), D-pantolactone (3.0 eq.), TEA (3.0 eq.), and EtOH (10 ml/mmol). The reaction mixture was stirred at 145°C for 2 days. The solvent was then evaporated under vacuum and the desired product was purified from the residue by flash chromatography.

General procedure E: Synthesis of primary amides

To a 50 mL or 100 mL round bottom flask were added the acyl chloride (1.0 eq) and MeCN (0.5 mL/mmol). The mixture was cooled to 0°C and aqueous $NH_3$ (28%, 10.0 eq.) was added. The reaction mixture was stirred at room temperature for 2 hours before extracting the product in ethyl acetate (3 × 0.8 mL/mmol). The combined organic layers were washed with 2 M NaOH (2 mL/mmol), dried over anhydrous $MgSO_4$, and the solvent was evaporated under vacuum. The product was used directly in next step without further purification.

General procedure F: Synthesis of thioamides

To a 100 mL round bottom flask were added the amide (1.0 eq.) and THF (20 mL/mmol). The atmosphere was changed to $N_2$ before addition of Lawesson's reagent (0.5 eq.). The reaction mixture was stirred at rt overnight. The solvent was then evaporated under vacuum and the desired product was purified from the residue by flash chromatography.

General procedure G: Gabriel synthesis

General procedure $G_1$: Synthesis of phthalimides

To a 50 mL round bottom flask were added bromomethylthiazole (1.0 eq), DMF (2.4 mL/mmol), and potassium phthalimide (1.0 eq). The reaction mixture was stirred at rt overnight. Water (3.5 mL/mmol) was then added, and the mixture was stirred at rt for two hours. The precipitate was collected by filtration and dried under vacuum before purification by flash chromatography to give the desired product.

General procedure $G_2$: Hydrazinolysis of phthalimides

To a 50 mL round bottom flask were added phthalimide (1.0 eq.), $N_2H_4 \cdot H_2O$ (3.0 eq.), and EtOH (15 mL/mmol). The mixture was heated to reflux for 1 hour. Next, 2 M HCl (3 mL/mmol) was added and the mixture was heated to reflux for an additional 5 minutes until all the precipitate dissolved. The solution was cooled down to 25 °C and sat. $NaHCO_3$ was added dropwise until neutralization. The solvent was then evaporated under vacuum. Water (15 mL/mmol) was added to the residue, and the product was extracted in DCM (3 × 25 mL/mmol). The combined organic layers were dried over anhydrous $MgSO_4$ and the solvent was evaporated under vacuum. The desired product was purified from the residue by flash chromatography.

General procedure H: Synthesis of organozinc reagents

Mg turnings were pretreated prior to use. Thus, Mg turnings (5 g) were suspended in 1 M HCl (25 mL). When bubbling ceased, the mixture was filtered under vacuum. This process was repeated again, and the Mg turnings collected by filtration were washed with acetone and diethyl ether. These pretreated Mg turnings were dried under vacuum for one hour and stored under nitrogen.

To a 50 mL three-neck round bottom flask equipped with a condenser and a pressure equalizing addition funnel, were added Mg turnings (401.0 mg, 16.5 mmol, 1.1 eq.) and a tiny crystal of $I_2$. Anhydrous THF (2.5 mL) was added and the system was purged with nitrogen and heated to 65$^{o}$C. The alkyl halide (15 mmol, 1.0 eq.) was dissolved in THF (12.5 mL), transferred to the addition funnel, and added dropwise to the mixture. The addition rate was adjusted to maintain a gentle reflux. The reaction mixture was kept refluxing for an hour, then cooled down to rt before use. The concentration of this generated Grignard reagent was determined by titration as reported by Knochel *et al*.[1]

To a 50 mL round bottom flask were added anhydrous $ZnCl_2$ (736.0 mg, 5.4 mmol, 1.2 eq.) and anhydrous LiCl (229.0 mg, 5.4 mmol, 1.2 eq.). The mixture was cooled to 0$^{o}$C and the Grignard reagent or commercial organolithium reagent (4.5 mmol. 1.0 eq.) was added dropwise. The reaction mixture was stirred at rt for 30 minutes and directly used in the next step.

General procedure I: Negishi coupling

To a 50 mL round bottom flask were added 2-bromo-5-cyanothiazole (567.0 mg, 3 mmol, 1.0 eq.) and $Pd(PPh_3)_4$ (173.3 mg, 0.15 mmol, 0.05 eq.). The system was purged with $N_2$, before the organozinc reagent was added. The reaction mixture was stirred at 70$^{o}$C for 30 minutes, before cooling to rt and quenching with sat. $NH_4Cl$ (20 mL). The product was extracted in EA (3 × 25 mL). The combined organic layers were washed with brine (2 × 20 mL), dried over anhydrous $Na_2SO_4$, and the solvent was evaporated under vacuum. The mixture was purified by flash chromatography to give the 2-alkyl-5-cyanothiazole.

General procedure J: Nitrile reduction

To a 50 mL round bottom flask were added the 2-alkyl-5-cyanothiazole (1.0 eq.) and THF (5 mL). The system was purged with $N_2$ and the mixture was cooled to 0$^{o}$C before $LiAlH_4$ (2 M in THF, 3.0 eq.) was added dropwise. The reaction mixture was stirred at rt for 3 hours, before cooling to 0$^{o}$C and quenching with $H_2O$ (3 mL), 1 M NaOH (3 mL), and $H_2O$ (9 mL) (caution: violent gas generation and extremely exothermic!). The mixture was filtered over Celite and the filter cake was washed with EA. The filtrate was extracted with EA (3 × 25 mL), and the combined organic layers were washed with 1 M HCl (3 × 25 mL). The pH of the combined aqueous layer was adjusted to 12 with solid $Na_2CO_3$ before extraction of the product in EA (3 × 25 mL). The combined organic layers were dried over anhydrous $Na_2SO_4$, and the solvent was evaporated under vacuum. The mixture was purified by flash chromatography to give the (2-alkylthiazol-5-yl)methanamine.

General procedure K: Catalytic ring-opening of D-pantolactone

Following a published method,[2] to a 25 mL round bottom flask were added the (2-alkylthiazol-5-yl)methanamine (1.0 eq.), D-pantolactone (2.0 eq.), 1,5,7-triazabicyclo[4.4.0]dec-5-ene (TBD, 0.1 eq.), and toluene. The reaction mixture was stirred at rt for 24 hours. The mixture was purified by flash chromatography to give the desired product.

General procedure L: Acylation of amine

To a 25 mL round bottom flask were added allylamine (2.67 mmol, 1.0 eq), TEA (2.58 mmol, 1.0 eq), and DCM (3 mL). The mixture was cooled to 0°C and the acyl chloride (2.67 mmol, 1.0 eq) was added dropwise. The reaction mixture was stirred at room temperature for 1 hour. Water (2 mL) was then added, and the product was extracted in DCM (3 × 3 mL). The combined organic layers were dried over anhydrous $MgSO_4$ and the solvent was evaporated under vacuum. The product was used without further purification.

General procedure G: Thiazole synthesis via bromocyclization

Following a published method,[3] to a 50 mL round bottom flask were added the *N*-allyl thioamide (1.0 eq) and $CHCl_3$ (17 mL/mmol). The atmosphere was changed to $N_2$ before addition of NBS (2.5 eq). The reaction mixture was stirred at rt overnight. The reaction was then quenched with sat. aqueous $Na_2SO_3$ (15 mL/mmol) and the product was extracted in DCM (3 × 17 mL/mmol). The combined organic layers were washed with brine (1 × 46 mL/mmol), dried over anhydrous $Na_2SO_4$, and the solvent was evaporated under vacuum. The residue was purified by flash chromatography to give the desired product.

Synthesis of **1.3**

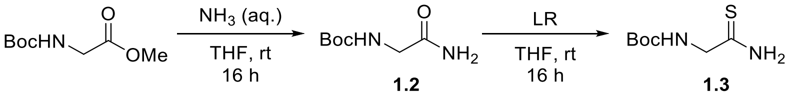


To a 50 mL round bottom flask were added N-Boc-glycine methyl ester (2.0 mL, 11.4 mmol, 1.0 equiv.) and THF (5 mL). An $NH_3$ aqueous solution (4.44 mL, 114 mmol, 10.0 equiv.) was then added. The reaction mixture was stirred at rt for 24 hours. The solvent was evaporated under vacuum to give **1.2** (1.73g, 87%) as a white solid. $R_f = 0.11$ (40% EA in Hex). $^1H$ NMR (500 MHz, $CDCl_3$) $\delta$ 6.44 (bs, 1H), 6.19 (bs, 1H), 5.47 (bs, 1H), 3.79 (d, $J$ = 5.7 Hz, 2H), 1.43 (s, 9H). Characterization matched previously reported data.[4]

Following general procedure F, **1.2** (615.2 mg, 3.53 mmol) was used to afford **1.3** (390 mg, 58%) as a yellow solid solid. $R_f = 0.30$ (Hex/EA = 1:1). $^1H$ NMR (500 MHz, $CDCl_3$) $\delta$ 7.86 (bs, 1H), 7.53 (bs, 1H), 5.27 (bs, 1H), 4.16 (d, $J$ = 4.9 Hz, 2H), 1.46 (s, 9H). Characterization matched previously reported data.[4]

Synthesis of **1a**

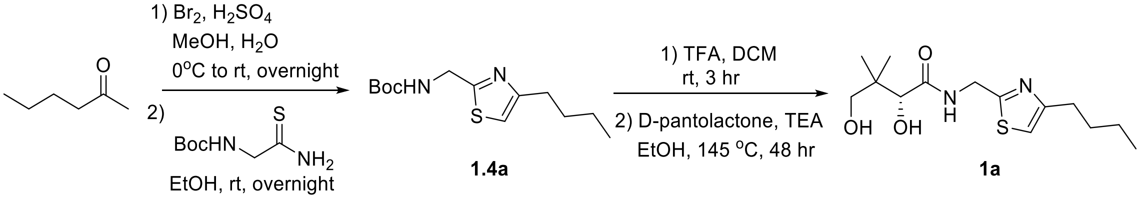


**1.4a** **1a**

Following general procedure A, 2-hexanone (1.0 mL, 8.1 mmol), MeOH (5 mL), $Br_2$ (0.42 mL, 8.1 mmol), $H_2O$ (2 mL), and $H_2SO_4$ (4.2 mL) were reacted to synthesize **1.1a** (1.23 g, 85%), which was used without further purification. Following general procedure $B_1$, **1.1a** (1.23 g, 6.9 mmol), **1.3** (1.31 g, 6.9 mmol), and EtOH (20 mL) were used. Purification by flash chromatography gave **1.4a** (1.03 g, 55%) as a colorless oil. $R_f$ = 0.65 (Hex/EA = 1:1); $^1$H NMR (500 MHz, $CDCl_3$): $\delta$ 6.77 (s, 1H), 5.35 (s, 1H), 4.56 (m, 2H), 2.70 (t, $J$ = 7.6 Hz, 2H), 1.64 (p, $J$ = 7.5 Hz, 2H), 1.44 (s, 9H), 1.35 (sext, $J$ = 7.3 Hz, 2H), 0.91 (t, $J$ = 7.4 Hz, 3H); $^{13}$C NMR (125 MHz, $CDCl_3$): $\delta$ 167.95, 157.61, 155.63, 112.81, 80.04, 42.42, 31.33, 31.21, 28.35, 22.36, 13.87; HRMS (ESI$^+$) *m/z* $[C_{13}H_{22}O_2N_2SNa]^+$ calcd: 293.1294; found: 293.1303.

Following general procedure C, **1.4a** (193.0 mg, 0.71 mmol), TFA (0.55 mL, 7.1 mmol), and DCM (3.5 mL) were reacted to give the corresponding amine, which was used directly in the next step without purification. Following general procedure D, the corresponding amine (80.0 mg, 0.47 mmol), D-pantolactone (183.5 mg, 1.41 mmol), TEA (142.5 mg, 1.41 mmol), and EtOH (3.5 mL) were used. Purification by flash chromatography gave **1a** (32 mg, 25%) as a yellow oil. $R_f$ = 0.25 (EA); $^1$H NMR (500 MHz, $CDCl_3$): $\delta$ 7.69 (m, 1H), 6.80 (s, 1H), 4.80 (dd, $J$ = 16.2, 6.4 Hz, 1H), 4.68 (bs, 1H), 4.63 (dd, $J$ = 16.1, 6.0 Hz, 1H), 4.08 (s, 1H), 3.49 (s, 2H), 2.70 (t, $J$ = 7.6 Hz, 2H), 1.63 (p, $J$ = 7.6 Hz, 2H), 1.34 (sext, $J$ = 7.5 Hz, 2H), 1.01 (s, 3H), 0.95 (s, 3H), 0.91 (t, $J$ = 7.4 Hz, 3H); $^{13}$C NMR (125 MHz, $CDCl_3$): $\delta$ 173.57, 166.75, 157.35, 113.32, 77.75, 70.92, 40.29, 39.51, 31.24, 31.03, 22.31, 21.50, 20.75, 13.86; HRMS (ESI$^+$) *m/z* $[C_{14}H_{24}O_3N_2SNa]^+$ calcd: 323.1400; found: 323.1397; HPLC purity: 99%; method A: $t_R$ = 18.46 min; method B $t_R$ = 9.24 min.

Synthesis of **1b**

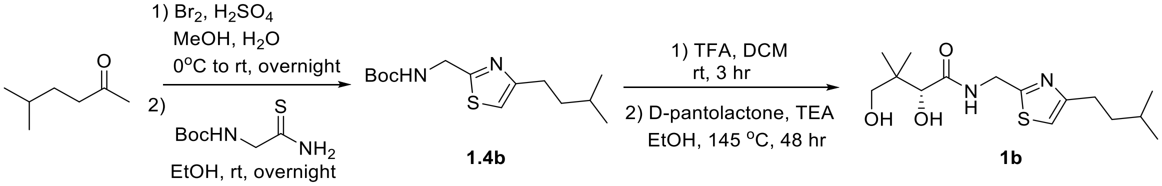


**1.4b** **1b**

Following general procedure A, 5-methyl-2-hexanone (1.0 mL, 7.1 mmol), MeOH (4.5 mL), $Br_2$ (0.36 mL, 7.1 mmol), $H_2O$ (2.2 mL), and $H_2SO_4$ (3.6 mL) were reacted to synthesize **1.1b** (1.37 g, quant.), which was used without further purification. Following general procedure $B_1$, **1.1b** (110.0 mg, 0.57 mmol), **1.3** (743.2 g, 3.9 mmol), and EtOH (1 mL) were used. No purification was needed to give **1.4b** (146 mg, 90%) as a colorless oil. $^1$H NMR (500 MHz, $CDCl_3$) $\delta$ 6.76 (s, 1H), 5.39 (s, 1H), 4.55 (m, 2H), 2.70 (m, 2H), 1.62-1.52 (m, 3H), 1.43 (s, 9H), 0.91 (d, $J$ = 6.2 Hz, 6H); $^{13}$C NMR (125 MHz, $CDCl_3$) $\delta$ 168.01, 157.78, 155.63, 112.66, 80.02, 42.40, 38.24, 29.47, 28.34, 27.76, 22.45. HRMS (ESI$^+$) *m/z* $[C_{14}H_{24}O_2N_2SNa]^+$ calcd: 307.1451; found: 307.1461.

Following general procedure C, **1.4b** (99.7 mg, 0.35 mmol), TFA (0.27 mL, 3.5 mmol), and DCM (1.7 mL) were reacted to give the corresponding amine, which was employed directly in the next step without purification. Following general procedure D, the corresponding amine (64.5 mg, 0.35 mmol), D-pantolactone (470.0 mg, 3.61 mmol), TEA (363.0 mg, 3.56 mmol), and EtOH (1.9 mL) were reacted. Purification by flash chromatography gave **1b** (14 mg, 9%) as an orange oil. $R_f$ = 0.26 (EA); $^1$H NMR (500 MHz, $CDCl_3$) $\delta$ 7.67 (t, $J$ = 6.2 Hz, 1H), 6.80 (s, 1H), 4.80 (m, 1H), 4.64 (m, 1H), 4.08 (s, 1H), 3.50 (s, 2H), 2.72-2.68 (m, 2H), 1.59-1.51 (m, 3H), 1.02 (s, 3H), 0.95 (s, 3H), 0.91 (d, $J$ = 6.2 Hz, 6H); $^{13}$C NMR: (125 MHz, $CDCl_3$) $\delta$ 173.53, 166.75, 157.55, 113.16, 77.79, 70.95, 40.33, 39.51, 38.19, 29.30, 27.72, 22.45, 21.51, 20.76. HRMS (ESI$^+$) *m/z* $[C_{15}H_{26}O_3N_2SNa]^+$ calcd: 337.1556; found: 337.1573; HPLC purity: 77%; method A: $t_R$ = 19.98 min; method B $t_R$ = 9.90 min.

<u>Synthesis of **1c**</u>

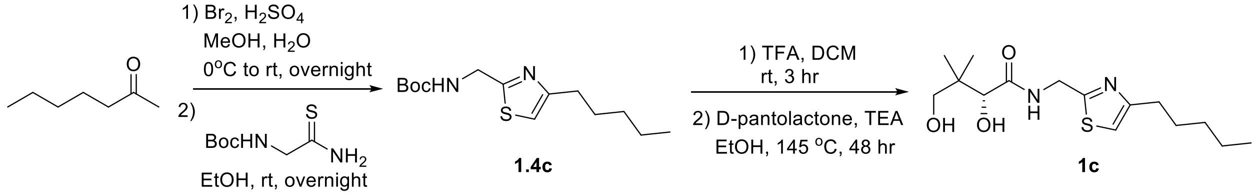


Following general procedure A, 2-heptanone (2.0 mL, 14.3 mmol), MeOH (9 mL), $Br_2$ (0.74 mL, 14.3 mmol), $H_2O$ (4.5 mL), and $H_2SO_4$ (7.4 mL) were reacted to synthesize **1.1c** (1.65 g, 60%), which was used without further purification. Following general procedure $K_2$, **1.1c** (180.0 mg, 0.93 mmol), **1.3** (176.5 mg, 0.93 mmol), and EtOH (2.5 mL) were reacted. Purification by flash chromatography gave **1.4c** (140 mg, 53%) as an orange oil.

Following general procedure C, **1.4c** (76.2 mg, 0.27 mmol), TFA (0.21 mL, 2.7 mmol), and DCM (1.3 mL) were reacted to give the corresponding amine, which was used directly in the next step without purification. This step was repeated again. Following general procedure D, the corresponding amine (66.3 mg, 0.36 mmol), D-pantolactone (140.5 mg, 1.08 mmol), TEA (109.1 mg, 1.08 mmol), and EtOH (3 mL) were reacted. Purification by flash chromatography gave **1c** (28 mg, 25%) as an orange oil. $R_f$ = 0.25 (EA); $^1$H NMR (500 MHz, $CDCl_3$): $\delta$ 7.71 (m, 1H), 6.79 (s, 1H), 4.78 (dd, $J$ = 16.0, 6.5 Hz, 1H), 4.63 (dd, $J$ = 16.0, 5.9 Hz, 1H), 4.08 (s, 1H), 3.48 (s, 2H), 2.69 (t, $J$ = 7.7 Hz, 2H), 1.63 (p, $J$ = 7.6 Hz, 2H), 1.33-1.26 (m, 4H), 1.00 (s, 3H), 0.94 (s, 3H), 0.87 (t, $J$ = 6.8 Hz, 3H); $^{13}$C NMR (125 MHz, $CDCl_3$): $\delta$ 173.64, 166.79, 157.38, 113.31, 77.68, 70.90, 40.29, 39.49, 31.42, 31.31, 28.82, 22.45, 21.44, 20.74, 14.02; HRMS (ESI$^+$) *m/z* $[C_{15}H_{26}O_3N_2SNa]^+$ calcd: 337.1556; found: 337.1552; HPLC purity: 97%; method A: $t_R$ = 20.27 min; method B $t_R$ = 9.99 min.

<u>Synthesis of **1d**</u>

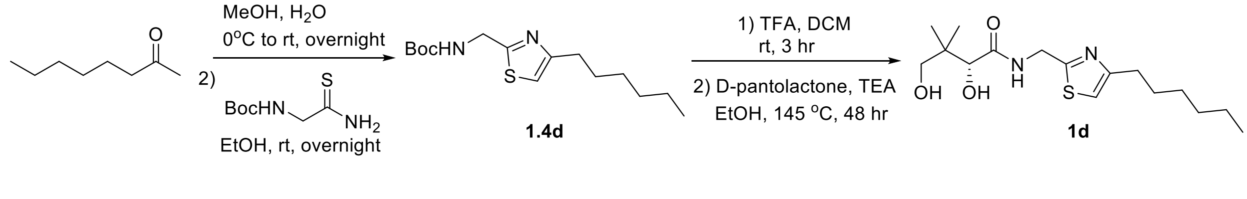

Following general procedure A, 2-octanone (1.0 mL, 6.4 mmol), MeOH (4 mL), $Br_2$ (0.33 mL, 6.4 mmol), $H_2O$ (2 mL), and $H_2SO_4$ (3.3 mL) were reacted to synthesize **1.1d** (1.14 g, 86%), which was used without further purification. Following general procedure $B_1$, **1.1d** (1.14 g, 5.5 mmol), **1.3** (1.04 g, 5.5 mmol), and EtOH (16 mL) were reacted. Purification by flash chromatography gave **1.4d** (725 mg, 44%) as a yellow oil. $R_f$ = 0.66 (Hex/EA = 1:1); $^1$H NMR (500 MHz, $CDCl_3$): $\delta$ 6.76 (s, 1H), 5.38 (s, 1H), 4.55 (m, 2H), 2.69 (t, $J$ = 7.6 Hz, 2H), 1.65 (p, $J$ = 7.6 Hz, 2H), 1.44 (s, 9H), 1.35-1.24 (m, 6H), 0.86 (t, $J$ = 7.1 Hz, 3H); $^{13}$C NMR (125 MHz, $CDCl_3$): $\delta$ 167.96, 157.64, 155.63, 112.79, 80.02, 42.41, 31.62, 31.52, 29.18, 28.96, 28.34, 22.57, 14.07; HRMS (ESI$^+$) *m/z* $[C_{15}H_{26}O_2N_2SNa]^+$ calcd: 321.1607; found: 321.1601.

Following general procedure C, **1.4d** (129.0 mg, 0.43 mmol), TFA (0.33 mL, 4.3 mmol), and DCM (2 mL) were reacted to give the corresponding amine, which was used directly in the next step without purification. Following general procedure D, the corresponding amine (75.4 mg, 0.38 mmol), D-pantolactone (148.4 mg, 1.14 mmol), TEA (115.2 mg, 1.14 mmol), and EtOH (3 mL) were reacted. Purification by flash chromatography gave **1d** (41 mg, 33%) as an orange oil. $R_f$ = 0.24 (EA); $^1$H NMR (500 MHz, $CDCl_3$): $\delta$ 7.71 (t, $J$ = 6.3 Hz, 1H), 6.79 (s, 1H), 4.78 (m, 1H), 4.63 (m, 1H), 4.07 (s, 1H), 3.48 (s, 2H), 2.68 (t, $J$ = 7.8 Hz, 2H), 1.63 (p, $J$ = 7.9 Hz, 2H), 1.30-1.26 (m, 6H), 1.00 (s, 3H), 0.94 (s, 3H), 0.86 (t, $J$ = 6.7 Hz, 3H); $^{13}$C NMR (125 MHz, $CDCl_3$): $\delta$ 173.67, 166.79, 157.39, 113.30, 77.65, 70.90, 40.30, 39.48, 31.61, 31.35, 29.09, 28.91, 22.57, 21.42, 20.75, 14.07; HRMS (ESI$^+$) *m/z* $[C_{16}H_{28}O_3N_2SNa]^+$ calcd: 351.1713; found: 351.1712; HPLC purity: 90%; method A: $t_R$ = 22.99 min; method B $t_R$ = 10.76 min.

Synthesis of **1e**

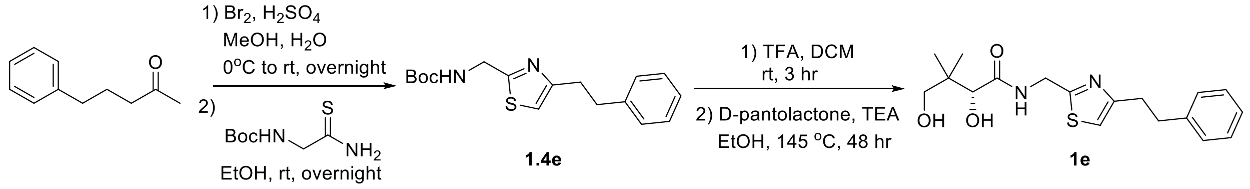


Following general procedure A, 4-phenyl-2-butanone (1.5 mL, 10.0 mmol), MeOH (9 mL), $Br_2$ (0.51 mL, 10.0 mmol), $H_2O$ (4.4 mL), and $H_2SO_4$ (7.4 mL) were reacted to synthesize **1.1e** (2.11 g, 93%), which was used without further purification. Following general procedure $B_1$, **1.1e** (65.2 mg, 0.29 mmol), **1.3** (54.6 mg, 0.29 mmol), and EtOH (0.7 mL) were reacted. No purification was needed to give **1.4e** (67 mg, 73%) as a yellow oil. $R_f$ = 0.50 (DCM/EA = 5:1); $^1$H NMR (500 MHz, $CDCl_3$) $\delta$ 7.31-7.24 (m, 2H), 7.22-7.15 (m, 3H), 6.75 (s, 1H), 5.32 (s, 1H), 4.60 (m, 2H), 3.06-2.98 (m, 4H), 1.47 (s, 9H); $^{13}$C NMR (125 MHz, $CDCl_3$) $\delta$ 168.17, 156.43, 155.63, 141.39, 128.43, 128.38, 126.02, 113.60, 80.14, 42.46, 35.52, 33.34, 28.38. HRMS (ESI$^+$) *m/z* $[C_{17}H_{22}O_2N_2SNa]^+$ calcd: 341.1294; found: 341.1282.

Following general procedure C, **1.4e** (67.0 mg, 0.21 mmol), TFA (0.16 mL, 2.1 mmol), and DCM (1.2 mL) were reacted to give the corresponding amine, which was used directly in the next step without purification. Following general procedure D, the corresponding amine (50.0 mg, 0.23 mmol), D-pantolactone (60.0 mg, 0.46 mmol), TEA (46.5 mg, 0.46 mmol), and EtOH (0.5 mL) were reacted. Purification by flash chromatography gave **1e** (15 mg, 19%) as an orange oil. $R_f$ =

0.20 (EA); $^1$H NMR (500 MHz, $CDCl_3$) $\delta$ 7.56 (m, 1H), 7.29-7.15 (m, 5H), 6.77 (s, 1H), 4.88 (dd, $J$ = 16.2, 6.6 Hz, 1H), 4.71 (dd, $J$ = 16.2, 5.7 Hz, 1H), 4.10 (s, 1H), 3.55 (s, 2H), 3.06-3.02 (m, 4H), 1.07 (s, 3H), 1.00 (s, 3H); $^{13}$C NMR (125 MHz, $CDCl_3$) $\delta$ 173.18, 166.87, 156.15, 141.17, 128.42, 128.42, 126.10, 114.03, 78.15, 71.03, 40.41, 39.59, 35.45, 33.17, 21.71, 20.91. HRMS ($ESI^+$) *m/z* $[C_{18}H_{24}O_3N_2SNa]^+$ calcd: 371.1400; found: 371.1406; HPLC purity: 77%; method A: $t_R$ = 19.49 min; method B $t_R$ = 9.66 min.

<u>Synthesis of **2.2**</u>

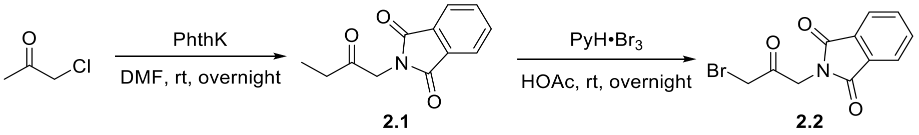


To a 50 mL round bottom flask were added potassium phthalimide (2.00 g, 10.8 mmol, 1.0 eq.) and DMF (6 mL). Chloroacetone (0.95 mL, 11.9 mmol, 1.1 eq.) was added dropwise and the mixture was stirred at rt overnight. Water (10 mL) was then added and the mixture was stirred for another hour. The mixture was cooled to 0 °C and stirred for 10 minutes before collecting the product by filtration. The filter cake was washed with water and dried to give **2.1** (1.47 g ,67%) as a white solid. $^1$H NMR (500 MHz, $CDCl_3$): $\delta$ 7.88 (m, 2H), 7.74 (m, 2H), 4.50 (s, 2H), 2.27 (s, 3H). Characterization matched previously reported data.[5]

To a 25 mL round bottom flask were added **2.1** (330.0 mg, 1.62 mmol, 1.0 eq.), pyridinium tribromide (570.0 mg, 1.78 mmol, 1.1 eq.), and HOAc (1.3 mL). The mixture was stirred at rt overnight. Water (1.3 mL) was then added and the mixture was stirred for another 2 hours. The precipitate was collected by filtration, washed with water, and dried to give 2-(3-bromo-2-oxopropyl)isoindoline-1,3-dione (375 mg, 82%) as a beige solid. $^1$H NMR (500 MHz, $CDCl_3$): $\delta$ 7.89 (m, 2H), 7.76 (m, 2H), 4.78 (s, 2H), 4.01 (s, 2H). Characterization matched previously reported data.[6]

<u>Synthesis of **2a**</u>

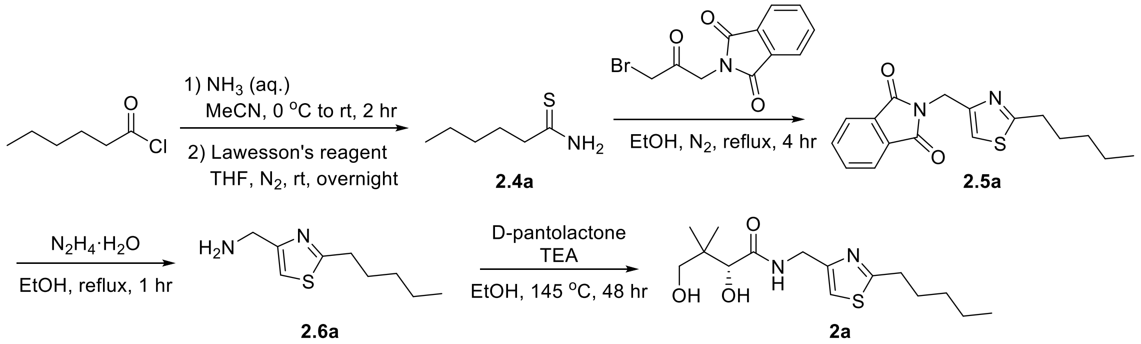


Following general procedure E, hexanoyl chloride (1.92 g, 14.3 mmol), aqueous $NH_3$ (9.6 mL, 143 mmol), and MeCN (7 mL) were reacted to synthesize **2.3a** (264 mg, 16%), which was used without further purification. Following general procedure F, **2.3a** (230 mg, 2.0 mmol), Lawesson's reagent (404.5 mg, 1.0 mmol), and THF (40 mL) were reacted. Purification by flash chromatography gave **2.4a** (173 mg, 66%) as a white solid. $R_f$ = 0.29 (DCM); $^1$H NMR (500 MHz,

$CDCl_3$): $\delta$ 7.77 (s, 1H), 6.93 (bs, 1H), 2.64 (t, $J$ = 7.6, 2H), 1.75 (p, $J$ =7.5 Hz, 2H), 1.39-1.27 (m, 4H), 0.89 (t, $J$ = 7.0 Hz, 3H). Characterization matched previously reported data.[7]

Following general procedure $B_2$, **2.4a** (173 mg, 1.32 mmol), **2.2** (372.4 mg, 1.32 mmol), and EtOH (2 mL) were reacted. Purification by flash chromatography gave **2.5a** (245 mg, 59%) as a white solid. $R_f$ = 0.66 (Hex/EA = 1:1); $^1$H NMR (500 MHz, $CDCl_3$): $\delta$ 7.85 (m, 2H), 7.71 (m, 2H), 6.97 (m, 1H), 4.95 (d, $J$ = 0.9 Hz, 2H), 2.92 (t, $J$ = 7.7 Hz, 2H), 1.72 (p, $J$ = 7.8 Hz, 2H), 1.34- 1.29 (m, 4H), 0.86 (t, $J$ = 7.1 Hz, 3H); $^{13}$C NMR (125 MHz, $CDCl_3$): $\delta$ 172.08, 167.80, 150.47, 134.04, 132.14, 123.44, 114.76, 37.81, 33.45, 31.27, 29.71, 22.32, 13.93; HRMS (ESI$^+$) *m/z* $[C_{17}H_{18}O_2N_2SNa]^+$ calcd: 337.0981; found: 337.0977.

Following general procedure $G_2$, **2.5a** (210 mg, 0.67 mmol), $N_2H_4{\cdot}H_2O$ (100.2 mg, 2.01 mmol), and EtOH (10 mL) were reacted. Purification by flash chromatography gave **2.6a** (101 mg, 76%) as a yellow oil. $R_f$ = 0.09 (DCM/MeOH = 9:1); $^1$H NMR (500 MHz, $CDCl_3$): $\delta$ 6.92 (s, 1H), 3.94 (s, 2H), 2.95 (t, $J$ = 7.7 Hz, 2H), 2.07 (s, 2H), 1.76 (p, $J$ = 7.6 Hz, 2H), 1.40-1.32 (m, 4H), 0.89 (t, $J$ = 7.0 Hz, 3H); $^{13}$C NMR (125 MHz, $CDCl_3$): $\delta$ 172.09, 157.54, 112.36, 42.46, 33.52, 31.31, 29.85, 22.35, 13.95; HRMS (ESI$^+$) *m/z* $[C_9H_{17}N_2S]^+$ calcd:185.1107; found: 185.1104.

Following general procedure D, **2.6a** (89.2 mg, 0.45 mmol), D-pantolactone (175.7 mg, 1.35 mmol), TEA (136.6 mg, 1.35 mmol), and EtOH (4.5 mL) were reacted. Purification by flash chromatography gave **2a** (56 mg, 38%) as a yellow oil. $R_f$ = 0.20 (EA); $^1$H NMR (500 MHz, $CDCl_3$): $\delta$ 7.55 (m, 1H), 7.01 (s, 1H), 4.58-4.39 (m, 2H), 4.05 (s, 1H), 3.46 (s, 2H), 2.92 (t, $J$ = 7.7 Hz, 2H), 1.73 (m, 2H), 1.36-1.31 (m, 4H), 1.00 (s, 3H), 0.90-0.87 (m, 6H); $^{13}$C NMR (125 MHz, $CDCl_3$): $\delta$ 173.41, 172.74, 152.01, 114.65, 77.56, 70.86, 39.50, 38.84, 33.32, 31.20, 29.80, 22.32, 21.71, 20.39, 13.92; HRMS (ESI$^+$) *m/z* $[C_{15}H_{26}O_3N_2SNa]^+$ calcd: 337.1556; found: 337.1554.

Synthesis of **2b**

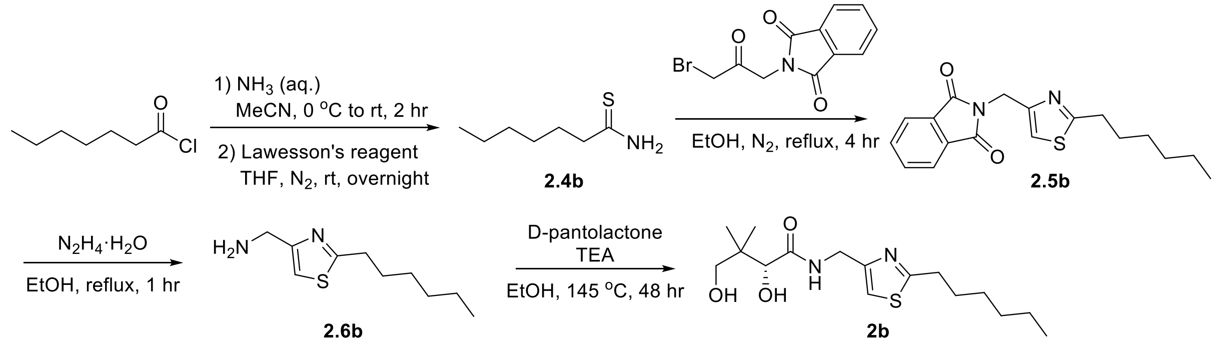


Following general procedure E, heptanoyl chloride (1.92 g, 12.9 mmol), aqueous $NH_3$ (8.7 mL, 129 mmol), and MeCN (6 mL) were reacted to synthesize the **2.3b** (500 mg, 30%), which was used without further purification. Following general procedure F, **2.3b** (400.5 mg, 3.1 mmol), Lawesson's reagent (627.0 mg, 1.55 mmol), and THF (62 mL) were reacted. Purification by flash chromatography gave **2.4b** (266 mg, 59%) as a white solid. $R_f$ = 0.29 (DCM); $^1$H NMR (500 MHz, $CDCl_3$): $\delta$ 7.44 (bs, 1H), 6.79 (bs, 1H), 2.66 (t, $J$ = 7.7, 2H), 1.77 (p, $J$ =7.8 Hz, 2H), 1.39-1.27 (m, 6H), 0.89 (t, $J$ = 6.6 Hz, 3H). Characterization matched previously reported data.[8]

Following general procedure $B_2$, **2.4b** (257.1 mg, 1.77 mmol), **2.2** (500.0 mg, 1.77 mmol), and EtOH (3 mL) were reacted. Purification by flash chromatography gave **2.5b** (215 mg, 37%) as a white solid. $R_f$ = 0.21 (Hex/DCM = 1:4); $^1$H NMR (500 MHz, $CDCl_3$): $\delta$ 7.86 (m, 2H), 7.72 (m, 2H), 6.98 (m, 1H), 4.96 (m, 2H), 2.92 (t, $J$ = 7.8 Hz, 2H), 1.72 (p, $J$ = 7.8 Hz, 2H), 1.35 (m, 2H), 1.28-1.25 (m, 4H), 0.85 (t, $J$ = 7.1 Hz, 3H); $^{13}$C NMR (125 MHz, $CDCl_3$): $\delta$ 172.10, 167.81, 150.46, 134.04, 132.15, 123.45, 114.77, 37.82, 33.50, 31.45, 29.99, 28.77, 22.48, 14.02; HRMS (ESI$^+$) *m/z* $[C_{18}H_{20}O_2N_2SNa]^+$ calcd: 351.1138; found: 351.1140.

Following general procedure $G_2$, **2.5b** (194.0 mg, 0.59 mmol), $N_2H_4 \cdot H_2O$ (88.2 mg, 1.77 mmol), and EtOH (9 mL) were reacted. No chromatography was needed to give **2.6b** (109 mg, 87%) as a yellow oil. $^1$H NMR (500 MHz, $CDCl_3$): $\delta$ 6.89 (m, 1H), 3.92 (s, 2H), 2.95 (t, $J$ = 7.6 Hz, 2H), 1.79-1.72 (m, 4H), 1.38 (m, 2H), 1.32-1.27 (m, 4H), 0.86 (t, $J$ = 7.1 Hz, 3H); $^{13}$C NMR (125 MHz, $CDCl_3$): $\delta$ 172.05, 158.00, 112.12, 42.63, 33.56, 31.47, 30.13, 28.80, 22.50, 14.03. HRMS (ESI$^+$) *m/z* $[C_{10}H_{18}N_2SNa]^+$ calcd: 221.1083; found: 221.1079.

Following general procedure D, **2.6b** (106 mg, 0.50 mmol), D-pantolactone (195.2 mg, 1.50 mmol), TEA (151.8 mg, 1.5 mmol), and EtOH (5 mL) were reacted. Purification by flash chromatography gave **2b** (94 mg, 55%) as a yellow oil. $R_f$ = 0.20 (EA); $^1$H NMR (500 MHz, $CDCl_3$): $\delta$ 7.52 (t, $J$ = 6.0 Hz, 1H), 7.01 (m, 1H), 4.61 (bs, 1H), 4.53 (dd, $J$ = 15.1, 5.9 Hz, 1H), 4.47 (dd, $J$ = 15.7, 5.8 Hz, 1H), 4.06 (s, 1H), 3.95 (bs, 1H), 3.47 (s, 2H), 2.93 (t, $J$ = 7.8 Hz, 2H), 1.72 (p, $J$ = 7.9 Hz, 2H), 1.36 (m, 2H), 1.31-1.27 (m, 4H), 1.01 (s, 3H), 0.91 (s, 3H), 0.87 (t, $J$ = 7.2 Hz); $^{13}$C NMR (125 MHz, $CDCl_3$): $\delta$ 173.31, 172.74, 152.01, 114.61, 77.64, 70.90, 39.53, 38.85, 33.37, 31.44, 30.09, 28.73, 22.48, 21.78, 20.38, 14.03. HRMS (ESI$^+$) *m/z* $[C_{16}H_{28}O_3N_2SNa]^+$ calcd: 351.1713; found: 351.1724; HPLC purity: 91%; method A: $t_R$ = 22.52 min; method B $t_R$ = 10.68 min.

Synthesis of **2c**

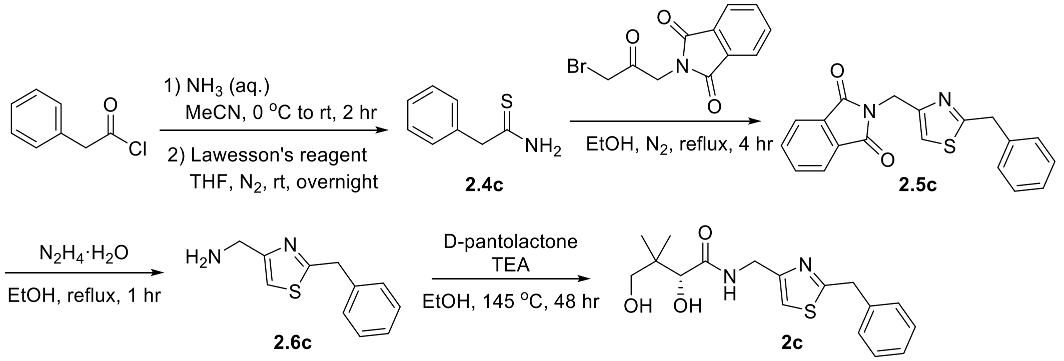


Following general procedure E, phenylacetyl chloride (1.17 g, 7.56 mmol), aqueous $NH_3$ (5.1 mL, 75.6 mmol), and MeCN (3.5 mL) were reacted to synthesize the **2.3c** (419 mg, 41%), which was used without further purification. Following general procedure F, 2-phenylacetamide (300.0 mg, 2.22 mmol), Lawesson's reagent (449.0 mg, 1.11 mmol), and THF (44 mL) were reacted. Purification by flash chromatography gave **2.4c** (161 mg, 48%) as a white solid. $R_f$ = 0.29 (DCM); $^1$H NMR (500 MHz, $CDCl_3$): $\delta$ 7.44 (bs, 1H), 6.79 (bs, 1H), 2.66 (t, $J$ = 7.7, 2H), 1.77 (p, $J$ =7.8 Hz, 2H), 1.39-1.27 (m, 6H), 0.89 (t, $J$ = 6.6 Hz, 3H). Characterization matched previously reported data.[9]

Following general procedure $B_2$, **2.4c** (161.0 mg, 1.06 mmol), **2.2** (299.4 mg, 1.06 mmol), and EtOH (2.5 mL) were reacted. Purification by flash chromatography gave **2.5c** (259 mg, 73%) as a white solid. $R_f$ = 0.76 (Hex/EA = 1:1); $^1$H NMR (500 MHz, $CDCl_3$): $\delta$ 7.88 (m, 2H), 7.73 (m, 2H), 7.33-7.26 (m, 5H), 7.02 (m, 1H), 4.99 (d, $J$ = 0.9 Hz, 2H), 4.28 (s, 2H); $^{13}$C NMR (125 MHz, $CDCl_3$): $\delta$ 171.14, 167.83, 150.79, 137.07, 134.07, 132.16, 129.13, 128.76, 127.17, 123.49, 116.02, 39.73, 37.77. HRMS (ESI$^+$) *m/z* $[C_{19}H_{14}O_2N_2SNa]^+$ calcd: 357.0668, found: 357.0667.

Following general procedure $G_2$, **2.5c** (254.0 mg, 0.76 mmol), $N_2H_4{\cdot}H_2O$ (133.6 mg, 2.28 mmol), and EtOH (12 mL) were reacted. No chromatography was needed to give **2.6c** (78 mg, 50%) as an orange oil. $^1$H NMR (500 MHz, $CDCl_3$): $\delta$ 7.34-7.26 (m, 5H), 6.93 (s, 1H), 4.30 (s, 2H), 3.95 (s, 2H), 1.87 (bs, 2H); $^{13}$C NMR (125 MHz, $CDCl_3$): $\delta$ 170.82, 158.28, 137.85, 129.04, 128.78, 127.14, 113.33, 42.60, 39.75; HRMS (ESI$^+$) *m/z* $[C_{11}H_{13}N_2S]^+$ calcd: 205.0794, found: 205.0794.

Following general procedure D, **2.6c** (65.4 mg, 0.32 mmol), D-pantolactone (125.0 mg, 0.96 mmol), TEA (97.1 mg, 0.96 mmol), and EtOH (3 mL) were reacted. Purification by flash chromatography ($SiO_2$) gave **2c** (64 mg, 60%) as a yellow oil. $R_f$ = 0.14 (EA); $^1$H NMR (500 MHz, $CDCl_3$): $\delta$ 7.49 (m, 1H), 7.33-7.24 (m, 5H), 7.02 (s, 1H), 4.52 (m, 2H), 4.25 (s, 2H), 4.04 (s, 1H), 3.45 (s, 2H), 1.00 (s, 3H), 0.90 (s, 3H); $^{13}$C NMR (125 MHz, $CDCl_3$): $\delta$ 173.27, 171.42, 152.46, 137.45, 128.98, 128.86, 127.30, 115.62, 77.64, 70.91, 39.50, 39.50, 38.90, 21.72, 20.42; HRMS (ESI$^+$) *m/z* $[C_{17}H_{22}O_3N_2SNa]^+$ calcd: 357.1243; found: 357.1240; HPLC purity: 79%; method A: $t_R$ = 18.26 min; method B $t_R$ = 9.08 min.

Synthesis of **2d**

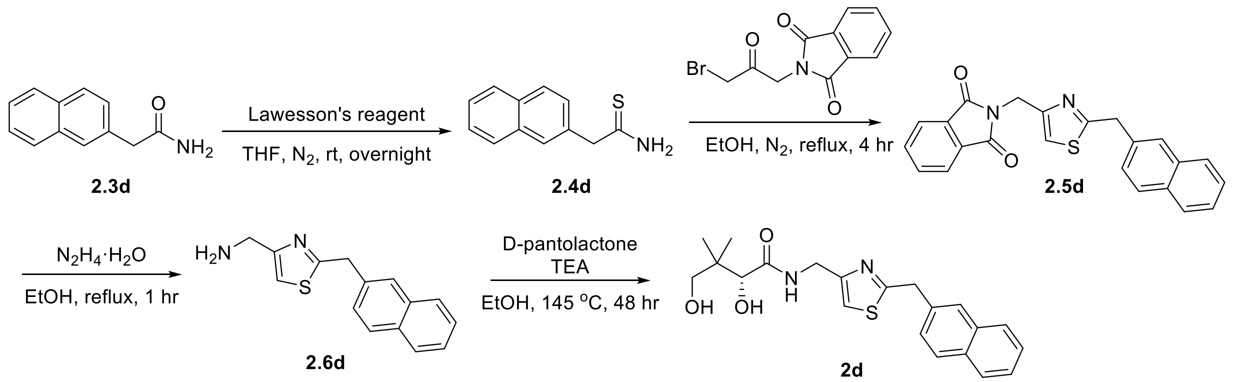


Following general procedure F, **2.3d** (400.0 mg, 2.16 mmol), Lawesson's reagent (436.8 mg, 1.08 mmol), and THF (43 mL) were reacted. Purification by flash chromatography gave **2.4d** (165 mg, 38%) as a white solid. $R_f$ = 0.29 (DCM); $^1$H NMR (500 MHz, $CDCl_3$): $\delta$ 8.01 (d, $J$ = 8.3 Hz, 1H), 7.91-7.85 (m, 2H), 7.60-7.52 (m, 2H), 7.50-7.46 (m, 1H), 7.41 (d, $J$ = 6.6 Hz, 1H), 6.57 (bs, 1H), 4.56 (s, 2H). Characterization matched previously reported data.[10]

Following general procedure $B_2$, **2.4d** (157.0 mg, 0.78 mmol), **2.2** (200.3 mg, 0.78 mmol), and EtOH (2 mL) were reacted. Purification by flash chromatography gave **2.5d** (177 mg, 59%) as a white solid. $R_f$ = 0.45 (Hex/EA = 1:1); $^1$H NMR (500 MHz, $CDCl_3$): $\delta$ 8.01 (m, 1H), 7.88 (m, 2H), 7.84 (m, 1H), 7.80 (m, 1H), 7.73 (m, 2H), 7.46-7.42 (m, 4H), 6.96 (s, 1H), 5.00 (s, 2H), 4.73 (s, 2H); $^{13}$C NMR (125 MHz, $CDCl_3$): $\delta$ 171.45, 167.83, 150.56, 134.06, 133.99, 133.82, 132.16,

131.77, 128.70, 128.32, 127.71, 126.41, 125.87, 125.55, 124.11, 123.49, 116.10, 37.78, 37.49. HRMS (ESI$^+$) *m/z* [$C_{23}H_{16}O_2N_2SNa$]$^+$ calcd: 407.0825; found: 407.0813.

Following general procedure G$_2$, **2.5d** (169.0 mg, 0.44 mmol), $N_2H_4{\cdot}H_2O$ (77.3 mg, 1.32 mmol), and EtOH (7 mL) were reacted. Purification by flash chromatography gave **2.6d** (50 mg, 45%) as a light yellow oil. $R_f$ = 0.26 (DCM/MeOH = 9:1); $^1$H NMR (500 MHz, $CD_3OD$): $\delta$ 8.02 (m, 1H), 7.88 (m, 1H), 7.83 (m, 1H), 7.50-7.44 (m, 4H), 7.15 (s, 1H), 4.76 (s, 2H), 3.90 (s, 2H). $^{13}$C NMR (125 MHz, $CD_3OD$): $\delta$ 172.21, 155.88, 134.21, 133.69, 131.66, 128.45, 127.99, 127.42, 126.02, 125.54, 125.25, 123.50, 114.34, 40.83, 36.43; HRMS (ESI$^+$) *m/z* [$C_{15}H_{14}N_2SNa$]$^+$ calcd: 277.0770; found: 277.0766.

Following general procedure D, **2.6d** (50 mg, 0.19 mmol), D-pantolactone (74.2 mg, 0.57 mmol), TEA (57.7 mg, 0.57 mmol), and EtOH (2 mL) were reacted. Purification by flash chromatography gave **2d** (38 mg, 52%) as a colorless oil. $R_f$ = 0.13 (EA); $^1$H NMR (500 MHz, $CDCl_3$): $\delta$ 7.96 (m, 1H), 7.86 (m, 1H), 7.81 (m, 1H), 7.51 (m, 1H), 7.49-7.44 (m, 2H), 7.44-7.39 (m, 2H), 6.96 (s, 1H), 4.70 (d, $J$ = 2.0 Hz, 2H), 4.53 (m, 2H), 4.05 (s, 1H), 3.45 (s, 2H), 1.00 (s, 3H), 0.90 (s, 3H); $^{13}$C NMR (125 MHz, $CDCl_3$): $\delta$ 173.28, 171.87, 152.18, 134.03, 133.51, 131.67, 128.86, 128.45, 127.70, 126.52, 125.96, 125.59, 123.81, 115.69, 77.60, 70.91, 39.48, 38.94, 37.23, 21.67, 20.41; HRMS (ESI$^+$) *m/z* $C_{21}H_{24}O_3N_2SNa$ [M+Na]$^+$ calcd: 407.1400, found: 407.1390; HPLC purity: 86%; method A: $t_R$ = 20.58 min; method B $t_R$ = 10.01 min.

<u>Synthesis of **2e**</u>

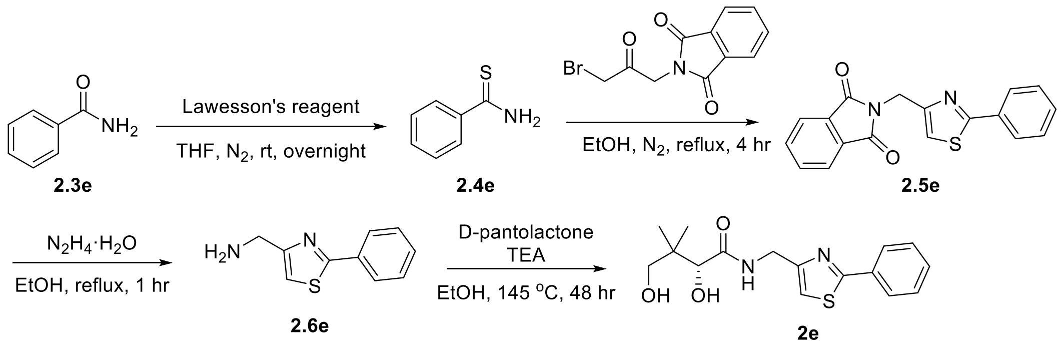


Following general procedure F, **2.3e** (200.0 mg, 1.65 mmol), Lawesson's reagent (335.7 mg, 0.83 mmol), and THF (33 mL) were reacted. Purification by flash chromatography gave **2.4e** (134 mg, 59%) as a yellow solid. $R_f$ = 0.26 (DCM); $^1$H NMR (500 MHz, $CDCl_3$): $\delta$ 7.89-7.86 (m, 2H), 7.71 (bs, 1H), 7.51 (tt, $J$ = 7.5, 1.2 Hz, 1H), 7.43-7.39 (m, 2H), 7.21 (bs, 1H). Characterization matched previously reported data.[11]

Following general procedure B$_2$, **2.4e** (127.6 mg, 0.93 mmol), **2.2** (238.8 mg, 0.93 mmol), and EtOH (2.5 mL) were reacted. Purification by flash chromatography gave **2.5e** (92 mg, 31%) as a white solid. $R_f$ = 0.47 (Hex/EA = 4:1); $^1$H NMR (500 MHz, $CDCl_3$): $\delta$ 7.91-7.88 (m, 4H), 7.73 (m, 2H), 7.40-7.38 (m, 3H), 7.16 (m, 1H), 5.06 (d, $J$ = 0.9 Hz, 2H); $^{13}$C NMR (125 MHz, $CDCl_3$): $\delta$ 168.46, 167.84, 152.09, 134.08, 133.44, 132.17, 130.07, 128.86, 126.59, 123.49, 115.76, 37.89. HRMS (ESI$^+$) *m/z* [$C_{18}H_{12}O_2N_2SNa$]$^+$ calcd: 343.0512; found: 343.0521.

Following general procedure $G_2$, **2.5e** (73.7 mg, 0.23 mmol), $N_2H_4\cdot H_2O$ (40.4 mg, 0.69 mmol), and EtOH (3.5 mL) were reacted. No chromatography was needed to give **2.6e** (18 mg, 40%) as a yellow oil. $^1H$ NMR (500 MHz, $CDCl_3$): $\delta$ 7.94 (m, 2H), 7.44-7.41 (m, 3H), 7.07 (s, 1H), 4.03 (s, 2H), 1.73 (bs, 2H); $^{13}C$ NMR (125 MHz, $CDCl_3$): $\delta$ 168.52, 159.22, 133.73, 129.99, 128.94, 126.54, 113.11, 42.80. HRMS ($ESI^+$) *m/z* $[C_{10}H_{11}N_2S]^+$ calcd: 191.0637; found: 191.0634.

Following general procedure D, **2.6e** (11.4 mg, 0.06 mmol), D-pantolactone (23.4 mg, 0.18 mmol), TEA (18.2 mg, 0.18 mmol), and EtOH (0.6 mL) were reacted. Purification by flash chromatography gave **2e** (7 mg, 38%) as a colorless oil. $R_f$ = 0.32 (EA); $^1H$ NMR (500 MHz, $CDCl_3$): $\delta$ 7.88 (m, 2H), 7.49 (m, 1H), 7.44-7.40 (m, 3H), 7.15 (s, 1H), 4.61 (d, $J$ = 5.9 Hz, 2H), 4.08 (s, 1H), 3.52 (d, $J$ = 11.2 Hz, 1H), 3.48 (d, $J$ = 11.2 Hz, 1H), 1.03 (s, 3H), 0.93 (s, 3H). $^{13}C$ NMR (125 MHz, $CDCl_3$): $\delta$ 173.12, 168.93, 153.67, 133.30, 130.27, 129.03, 126.53, 115.39, 77.79, 71.13, 39.51, 39.18, 21.56, 20.33; HRMS ($ESI^+$) *m/z* $[C_{16}H_{20}O_3N_2SNa]^+$ calcd: 343.1087; found: 343.1089; HPLC purity: 76%; method A: $t_R$ = 18.43 min; method B $t_R$ = 9.21 min.

<u>Synthesis of **2f**</u>

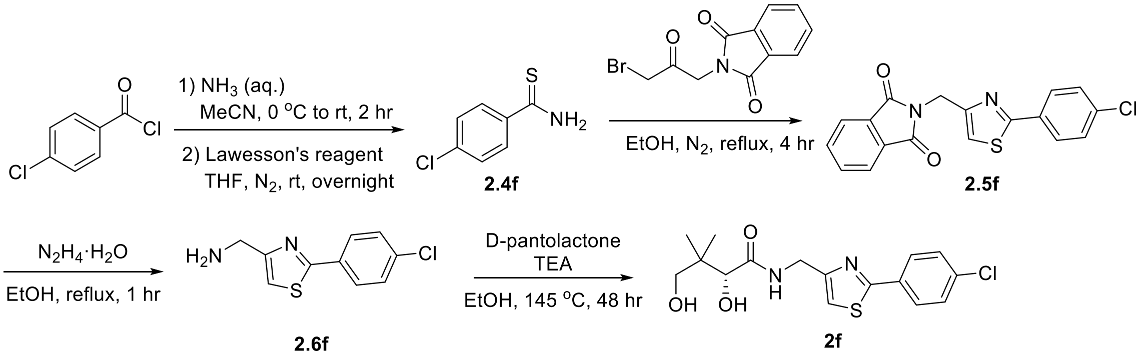


Following general procedure E, 4-chlorobenzoyl chloride (1.37 g, 7.80 mmol), aqueous $NH_3$ (5.2 mL, 78 mmol), and MeCN (3.5 mL) were reacted to synthesize the **2.3f** (607 mg, 50%), which was used without further purification. Following general procedure F, **2.3f** (398.0 mg, 2.56 mmol), Lawesson's reagent (517.7 mg, 1.28 mmol), and THF (50 mL) were reacted. Purification by flash chromatography gave **2.4f** (347 mg, 79%) as a yellow solid. $R_f$ = 0.26 (DCM); $^1H$ NMR (500 MHz, DMSO-$d_6$): $\delta$ 9.95 (bs, 1H), 9.56 (bs, 1H), 7.92-7.88 (m, 2H), 7.51-7.47 (m, 2H). Characterization matched previously reported data.[12]

Following general procedure $B_2$, **2.4f** (347.0 mg, 2.02 mmol), **2.2** (570.6 mg, 2.02 mmol), and EtOH (5 mL) were reacted. Purification by flash chromatography gave **2.5f** (300 mg, 42%) as a light yellow solid. $R_f$ = 0.22 (Hex/EA = 4:1); $^1H$ NMR (500 MHz, $CDCl_3$): $\delta$ 7.89 (m, 2H), 7.84 (d, $J$ = 8.5 Hz, 2H), 7.73 (m, 2H), 7.36 (d, $J$ = 8.6 Hz, 2H), 7.18 (s, 1H), 5.04 (s, 2H); $^{13}C$ NMR (125 MHz, $CDCl_3$): $\delta$ 167.82, 167.07, 152.29, 136.01, 134.12, 132.13, 131.93, 129.09, 127.78, 123.51, 116.14, 37.78; HRMS ($ESI^+$) *m/z* $[C_{18}H_{11}O_2N_2SClNa]^+$ calcd: 377.0122; found: 377.0129.

Following general procedure $G_2$, **2.5f** (291.0 mg, 0.82 mmol), $N_2H_4\cdot H_2O$ (144.0 mg, 2.46 mmol), and EtOH (13 mL) were reacted. Purification by flash chromatography gave **2.6f** (64 mg, 35%) as a yellow oil. $R_f$ = 0.15 (DCM/MeOH = 9:1); $^1H$ NMR (500 MHz, $CD_3OD$): $\delta$ 7.94 (d, $J$ = 8.7 Hz, 2H), 7.48 (d, $J$ = 8.7 Hz, 2H), 7.38 (s, 1H), 3.97 (s, 2H); $^{13}C$ NMR (125 MHz, $CD_3OD$): $\delta$ 167.09,

158.02, 135.72, 132.02, 128.91, 127.49, 114.52, 41.12; HRMS (ESI$^+$) *m/z* $[C_{10}H_{10}N_2SCl]^+$ calcd: 225.0248; found: 225.0239.

Following general procedure D, **2.6f** (64 mg, 0.28 mmol), D-pantolactone (109.3 mg, 0.84 mmol), TEA (85.0 mg, 0.84 mmol), and EtOH (2.5 mL) were reacted. Purification by flash chromatography gave **2f** (57 mg, 57%) as a yellow oil. $R_f$ = 0.26 (EA); $^1$H NMR (500 MHz, $CDCl_3$): $\delta$ 7.79 (d, $J$ = 8.6 Hz, 2H), 7.55 (t, $J$ = 5.9 Hz, 1H), 7.37 (d, $J$ = 8.5 Hz, 2H), 7.14 (s, 1H), 4.58-4.56 (m, 2H), 4.07 (s, 1H), 3.48 (s, 2H), 0.99 (s, 3H), 0.92 (s, 3H); $^{13}$C NMR (125 MHz, $CDCl_3$): $\delta$ 173.45, 167.48, 153.92, 136.22, 131.75, 129.23, 127.68, 115.61, 77.60, 71.05, 39.45, 39.13, 21.33, 20.43; HRMS (ESI$^+$) *m/z* $[C_{16}H_{19}O_3N_2SClNa]^+$ calcd: 377.0697; found: 377.0697; HPLC purity: 86%; method A: $t_R$ = 20.84 min; method B $t_R$ = 10.17 min.

Synthesis of **2g**

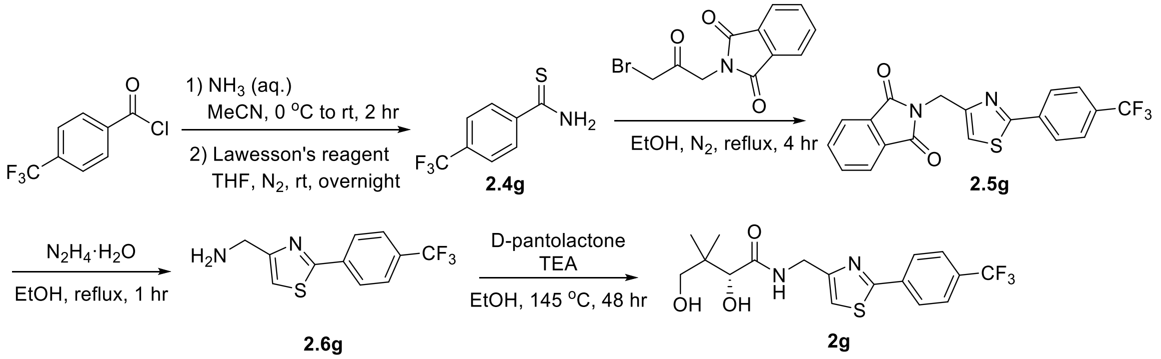


Following general procedure E, 4-trifluoromethylbenzoyl chloride (1.40 g, 6.73 mmol), aqueous $NH_3$ (4.5 mL, 67 mmol), and MeCN (3 mL) were reacted to synthesize the **2.3g** (1.06 g, 83%), which was used without further purification. Following general procedure F, **2.3g** (500.0 mg, 2.64 mmol), Lawesson's reagent (533.8 mg, 1.32 mmol), and THF (52 mL) were reacted. Purification by flash chromatography gave **2.4g** (450 mg, 83%) as a yellow solid. $R_f$ = 0.10 (Hex/DCM = 1:1); $^1$H NMR (500 MHz, DMSO-$d_6$): $\delta$ 10.12 (bs, 1H), 9.72 (bs, 1H), 8.01 (d, $J$ = 8.2 Hz, 2H), 7.79 (d, $J$ = 8.3 Hz, 2H). Characterization matched previously reported data.[13]

Following general procedure $B_2$, **2.4g** (360.0 mg, 1.75 mmol), **2.2** (494.3 mg, 1.75 mmol), and EtOH (4.5 mL) were reacted. No chromatography was needed to give **2.5g** (605 mg, 89%) as a light yellow solid. $^1$H NMR (500 MHz, $CDCl_3$): $\delta$ 8.02 (d, $J$ = 8.1 Hz, 2H), 7.90 (m, 2H), 7.74 (m, 2H), 7.65 (d, $J$ = 8.2 Hz, 2H), 7.26 (s, 1H), 5.07 (s, 2H); $^{19}$F NMR (471 MHz, $CDCl_3$): $\delta$ -62.81; $^{13}$C NMR (125 MHz, $CDCl_3$): $\delta$ 167.81, 166.49, 152.68, 136.47, 134.15, 132.11, 131.63 (q, $J$ = 32.5 Hz), 126.78, 125.88 (q, $J$ = 3.8 Hz), 123.88 (q, $J$ = 271.3 Hz), 123.53, 117.01, 37.74; HRMS (ESI$^+$) *m/z* $[C_{19}H_{11}F_3O_2N_2SNa]^+$ calcd: 411.0386; found: 411.0384.

Following general procedure $G_2$, **2.5g** (400.0 mg, 1.03 mmol), $N_2H_4 \cdot H_2O$ (181.0 mg, 3.10 mmol), and EtOH (16 mL) were reacted. Purification by flash chromatography gave **2.6g** (109 mg, 41%) as a yellow solid. $R_f$ = 0.22 (DCM/MeOH = 9:1); $^1$H NMR (500 MHz, $CD_3OD$): $\delta$ 8.12 (d, $J$ = 8.1 Hz, 2H), 7.75 (d, $J$ = 8.2 Hz, 2H), 7.43 (s, 1H), 4.84 (s, 2H); $^{19}$F NMR (471 MHz, $CD_3OD$): $\delta$ -64.33; $^{13}$C NMR (125 MHz, $CD_3OD$): $\delta$ 166.40, 159.00, 136.77, 131.22, (q, $J$ = 32.5 Hz), 126.53,

125.72 (q, $J$ = 3.8 Hz), 124.04 (q, $J$ = 270 Hz), 115.18, 41.24; HRMS (ESI$^+$) $m/z$ [$C_{11}H_{10}F_3N_2S$]$^+$ calcd: 259.0511; found: 259.0507.

Following general procedure D, **2.6g** (77.5 mg, 0.30 mmol), D-pantolactone (117.1 mg, 0.90 mmol), TEA (91.0 mg, 0.90 mmol), and EtOH (2.5 mL) were reacted. Purification by flash chromatography gave **2g** (63 mg, 54%) as a colorless oil. $R_f$ = 0.33 (EA); $^1$H NMR (500 MHz, $CDCl_3$): $\delta$ 8.00 (d, $J$ = 8.2 Hz, 2H), 7.67 (d, $J$ = 8.2 Hz, 2H), 7.49 (m, 1H), 7.23 (s, 1H), 4.63 (m, 2H), 4.10 (s, 2H), 4.04 (bs, 1H), 3.53 (d, $J$ = 11.2 Hz, 1H), 3.50 (d, $J$ = 11.2 Hz, 1H), 3.39 (bs, 1H), 1.03 (s, 3H), 0.94 (s, 3H); $^{19}$F NMR (471 MHz, $CDCl_3$) $\delta$ -62.83; $^{13}$C NMR (125 MHz, $CDCl_3$): $\delta$ 173.36, 166.83, 154.36, 136.37, 131.79 (q, $J$ = 32.5 Hz), 126.70, 126.01 (q, $J$ = 3.7 Hz), 123.83 (q, $J$ = 270 Hz), 116.39, 77.70, 71.14, 39.46, 39.15, 21.30, 20.41; HRMS (ESI$^+$) $m/z$ [$C_{17}H_{19}F_3O_3N_2SNa$]$^+$ calcd: 411.0961; found: 411.0965; HPLC purity: 96%; method A: $t_R$ = 22.40 min; method B $t_R$ = 10.51 min.

<u>Synthesis of **2h**</u>

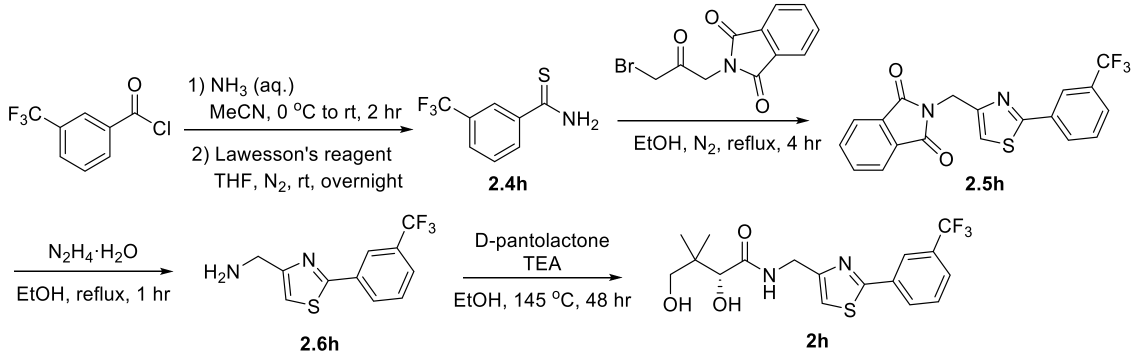


Following general procedure E, 3-trifluoromethylbenzoyl chloride (1.38 g, 6.63 mmol), aqueous $NH_3$ (4.4 mL, 66 mmol), and MeCN (3 mL) were reacted to synthesize the **2.3h** (1.27 g, quant.), which was used without further purification. Following general procedure F, **2.3h** (400.0 mg, 2.11 mmol), Lawesson's reagent (426.6 mg, 1.06 mmol), and THF (42 mL) were reacted. Purification by flash chromatography gave **2.4h** (372 mg, 86%) as a yellow solid. $R_f$ = 0.79 (Hex/EA = 1:1); $^1$H NMR (500 MHz, $CDCl_3$) $\delta$ 8.10 (s, 1H), 8.04 (d, $J$ = 8.0 Hz), 7.85 (bs, 1H), 7.76 (d, $J$ = 7.8 Hz, 1H), 7.55 (t, $J$ = 7.8 Hz, 1H), 7.26 (bs, 1H). Characterization matched previously reported data.[14]

Following general procedure $B_2$, **2.4h** (350.0 mg, 1.71 mmol), **2.2** (483.0 mg, 1.71 mmol), and EtOH (4.5 mL) were reacted. Purification by flash chromatography gave **2.5h** (405 mg, 61%) as a white solid. $^1$H NMR (500 MHz, $CDCl_3$): $\delta$ 8.14 (s, 1H), 8.07 (d, $J$ = 7.8 Hz, 1H), 7.89 (m, 2H), 7.74 (m, 2H), 7.63 (d, $J$ = 7.9 Hz, 1H), 7.52 (t, $J$ = 7.9 Hz, 1H), 7.24 (m, 1H), 5.07 (d, $J$ = 0.9 Hz, 2H); $^{19}$F NMR (471 MHz, $CDCl_3$): $\delta$ -62.79; $^{13}$C NMR (125 MHz, $CDCl_3$): $\delta$ 167.81, 166.54, 152.59, 134.15, 134.12, 132.12, 131.42 (q, $J$ = 32.5 Hz), 129.72, 129.44, 126.51 (q, $J$ = 3.8 Hz), 123.53, 123.30 (q, $J$ = 3.8 Hz), 123.78 (q, $J$ = 270 Hz), 116.68, 37.78; HRMS (ESI$^+$) $m/z$ [$C_{19}H_{11}F_3O_2N_2SNa$]$^+$ calcd: 411.0386; found: 411.0367.

Following general procedure $G_2$, **2.5h** (384.5 mg, 0.99 mmol), $N_2H_4 \cdot H_2O$ (175.2 mg, 3.00 mmol), and EtOH (15 mL) were reacted. Purification by flash chromatography gave **2.6h** (105 mg, 41%) as a white solid. $R_f$ = 0.25 (DCM/MeOH = 9:1); $^1$H NMR (500 MHz, $CD_3OD$): $\delta$ 8.25 (s, 1H), 8.16

(d, $J$ = 7.7 Hz, 1H), 7.74 (d, $J$ = 7.7 Hz, 1H), 7.66 (t, $J$ = 7.8 Hz, 1H), 7.42 (t, $J$ = 0.7 Hz, 1H), 3.97 (s, 2H); $^{19}$F NMR (471 MHz, $CD_3OD$): $\delta$ -64.36; $^{13}$C NMR (125 MHz, $CD_3OD$): $\delta$ 166.43, 158.71, 134.28, 131.14 (q, $J$ = 32.5 Hz), 129.80, 129.64, 126.18 (q, $J$ = 3.8 Hz), 123.95 (q, $J$ = 270 Hz), 122.37 (q, $J$ = 3.8 Hz), 114.96, 41.17; HRMS (ESI$^+$) *m/z* $[C_{11}H_{10}F_3N_2S]^+$ calcd: 259.0511; found: 259.0504.

Following general procedure D, **2.6h** (87.8 mg, 0.34 mmol), D-pantolactone (132.7 mg, 1.02 mmol), TEA (103.1 mg, 1.02 mmol), and EtOH (3 mL) were reacted. Purification by flash chromatography gave **2h** (104 mg, 79%) as a colorless oil. $R_f$ = 0.22 (EA); $^1$H NMR (500 MHz, $CDCl_3$): $\delta$ 8.16 (s, 1H), 8.04 (d, $J$ = 7.9 Hz, 1H), 7.66 (d, $J$ = 7.9 Hz, 1H), 7.54 (m, 1H), 7.51 (m, 1H), 7.21 (s, 1H), 4.64 (dd, $J$ = 15.4, 5.8 Hz, 1H), 4.58 (dd, $J$ = 15.5, 5.7 Hz, 1H), 4.09 (s, 1H), 3.51 (d, $J$ = 11.2 Hz, 1H), 3.50 (d, $J$ = 11.2 Hz, 1H), 1.01 (s, 3H), 0.93 (s, 3H); $^{19}$F NMR (471 MHz, $CDCl_3$) $\delta$ -62.81; $^{13}$C NMR (125 MHz, $CDCl_3$): $\delta$ 173.34, 166.84, 154.25, 134.04, 131.55 (q, $J$ = 32.5 Hz), 129.63, 129.58, 126.62 (q, $J$ = 3.8 Hz), 123.76 (q, $J$ = 271.3 Hz), 123.23 (q, $J$ = 3.8 Hz), 116.09, 77.68, 71.14, 39.46, 39.14, 21.27, 20.38; HRMS (ESI$^+$) *m/z* $[C_{17}H_{19}F_3O_3N_2SNa]^+$ calcd: 411.0961; found: 411.0951; HPLC purity: 97% method A: $t_R$ = 21.95 min; method B $t_R$ = 10.41 min.

Synthesis of **3a**

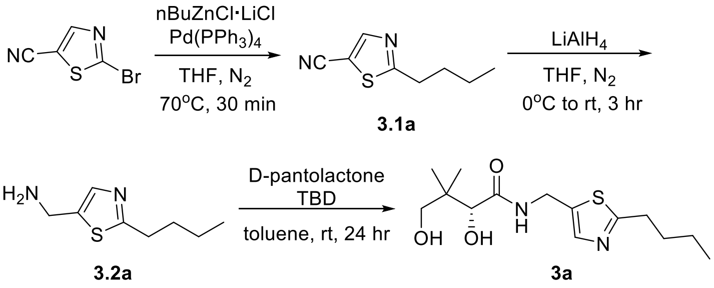


Following general procedure H and I, a commercial nBuLi solution (1.10 M titration in hexanes, 4.1 mL) was used for generation of the organozinc reagent. Purification by flash chromatography gave **3.1a** (350 mg, 70%) as a yellow oil. $R_f$ = 0.30 (Hex/EA = 5:1); $^1$H NMR ($CDCl_3$, 500 MHz): $\delta$ 8.13 (s, 1H), 3.05 (t, $J$ = 7.5 Hz, 2H), 1.79 (p, $J$ = 7.5 Hz, 2H), 1.42 (h, $J$ = 7.3 Hz, 2H), 0.95 (t, $J$ = 7.4 Hz, 3H); $^{13}$C NMR ($CDCl_3$, 125 MHz): $\delta$ 178.08, 151.89, 112.27, 105.31, 33.53, 31.91, 22.29, 13.84; HRMS (ESI$^+$) *m/z* $[C_8H_{10}N_2SNa]^+$ calcd: 189.0457; found: 189.0459.

Following general procedure J, **3.1a** (350.0 mg, 2.1 mmol) and $LiAlH_4$ (3.2 mL) were reacted. Purification by flash chromatography gave **3.2a** (194 mg, 54%) as a brown oil, which solidifies upon standing. $R_f$ = 0.13 (DCM/MeOH = 10:1); $^1$H NMR ($CDCl_3$, 500 MHz): $\delta$ 7.40 (s, 1H), 4.00 (s, 2H), 2.93 (t, $J$ = 7.5 Hz, 2H), 1.73 (p, $J$ = 7.6 Hz, 2H), 1.68 (bs, 2H), 1.40 (h, $J$ = 7.4 Hz, 2H), 0.92 (t, $J$ = 7.4 Hz, 3H); $^{13}$C NMR ($CDCl_3$, 125 MHz): $\delta$ 171.30, 140.59, 138.61, 39.13, 33.52, 32.25, 22.37, 13.95; HRMS (ESI$^+$) *m/z* $[C_8H_{15}N_2S]^+$ calcd: 171.0950; found: 171.0945.

Following general procedure K, **3.1a** (194.0 mg, 1.14 mmol), D-pantolactone (296.7 mg, 2.28 mmol), TBD (15.9 mg, 0.11 mmol), and toluene (1.2 mL) were reacted. Purification by flash chromatography gave **3a** (273 mg, 80%) as a sticky yellow oil. $R_f$ = 0.42 (DCM/MeOH = 10:1);

$^{1}H$ NMR ($CDCl_3$, 500 MHz): $\delta$ 7.45 (t, $J$ = 6.1 Hz, 1H), 7.40 (s, 1H), 4.58 (dd, $J$ = 15.4, 6.0 Hz, 1H), 4.52 (dd, $J$ = 15.4, 6.0 Hz, 1H), 4.03 (s, 1H), 3.47 (d, $J$ = 11.3 Hz, 1H), 3.44 (d, $J$ = 11.2 Hz, 1H), 2.90 (t, $J$ = 7.5 Hz, 2H), 1.69 (p, $J$ = 7.6 Hz, 2H), 1.36 (h, $J$ = 7.4 Hz, 2H), 0.96 (s, 3H), 0.90 (t, $J$ = 7.4 Hz, 3H), 0.89 (s, 3H); $^{13}C$ NMR ($CDCl_3$, 125 MHz): $\delta$ 173.68, 172.90, 140.21, 134.75, 77.60, 71.38, 39.54, 35.41, 33.31, 32.19, 22.31, 21.35, 20.64, 13.91; HRMS ($ESI^+$) *m/z* $[C_{14}H_{24}N_2O_3SNa]^+$ calcd: 323.1400; found: 323.1404. HPLC: method A $t_R$ = 16.38 min (99%); method B $t_R$ = 15.70 min (99%).

Synthesis of **3b**

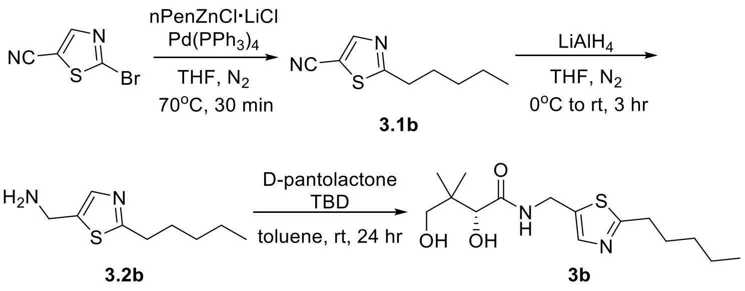


Following procedure H and I, 1-bromopentane (1.86 mL, 15 mmol) was used to generate the Grignard reagent (0.390 M titration, 11.5 mL used for generation of the organozinc reagent). Purification by flash chromatography gave **3.1b** (365 mg, 67%) as a yellow oil. $R_f$ = 0.29 (Hex/EA = 5:1); $^{1}H$ NMR ($CDCl_3$, 500 MHz): $\delta$ 8.12 (s, 1H), 3.03 (m, 2H), 1.80 (m, 2H), 1.41-1.29 (m, 4H), 0.89 (t, $J$ = 7.2 Hz, 3H); $^{13}C$ NMR ($CDCl_3$, 125 MHz): $\delta$ 177.98, 151.77, 112.15, 105.18, 33.68, 31.14, 29.44, 22.32, 13.93; HRMS ($ESI^+$) *m/z* $[C_9H_{12}N_2SNa]^+$ calcd: 203.0613; found: 203.0611.

Following procedure J, **3.1b** (365.0 mg, 2.0 mmol) and $LiAlH_4$ (3.0 mL) were reacted. Purification by flash chromatography gave **3.2b** (69 mg, 19%) as a brown oil. $R_f$ = 0.20 (DCM/MeOH = 10:1); $^{1}H$ NMR ($CDCl_3$, 500 MHz): $\delta$ 7.41 (s, 1H), 4.01 (s, 2H), 2.93 (t, $J$ = 7.5 Hz, 2H), 1.82 (bs, 2H), 1.75 (p, $J$ = 7.5 Hz, 2H), 1.38-1.30 (m, 4H), 0.88 (t, $J$ = 7.1 Hz, 3H); $^{13}C$ NMR ($CDCl_3$, 125 MHz): $\delta$ 171.30, 140.28, 138.61, 38.97, 33.71, 31.33, 29.78, 22.46, 14.03; HRMS ($ESI^+$) *m/z* $[C_9H_{17}N_2S]^+$ calcd: 185.1107; found: 185.1104.

Following procedure K, **3.2b** (69.0 mg, 0.37 mmol), D-pantolactone (97.6 mg, 0.75 mmol), TBD (5.6 mg, 0.04 mmol), and toluene (0.4 mL) were reacted. Purification by flash chromatography ($SiO_2$, DCM/EA = 4:1 to EA) gave **3b** (40 mg, 34%) as a sticky light brown oil. $R_f$ = 0.18 (EA); $^{1}H$ NMR ($CDCl_3$, 500 MHz): $\delta$ 7.45 (s, 1H), 7.33 (t, $J$ = 6.0 Hz, 1H), 4.61 (dd, $J$ = 15.4, 6.0 Hz, 1H), 4.55 (dd, $J$ = 15.4, 6.0 Hz, 1H), 4.07 (s, 1H), 3.52 (d, $J$ = 11.1 Hz, 1H), 3.48 (d, $J$ = 11.1 Hz, 1H), 2.92 (t, $J$ = 7.7 Hz, 2H), 1.74 (m, 2H), 1.37-1.31 (m, 4H), 1.00 (s, 3H), 0.92 (s, 3H), 0.89 (t, $J$ = 7.0 Hz, 3H); $^{13}C$ NMR ($CDCl_3$, 125 MHz): $\delta$ 173.22, 172.83, 140.20, 134.63, 77.76, 71.53, 39.52, 35.39, 33.56, 31.30, 29.75, 22.45, 21.35, 20.54, 14.05; HRMS ($ESI^+$) *m/z* $[C_{15}H_{27}N_2O_3S]^+$ calcd: 315.1737; found: 315.1738; HPLC: method A $t_R$ = 21.41 min (96%); method B $t_R$ = 17.38 min (99%).

Synthesis of **3c**

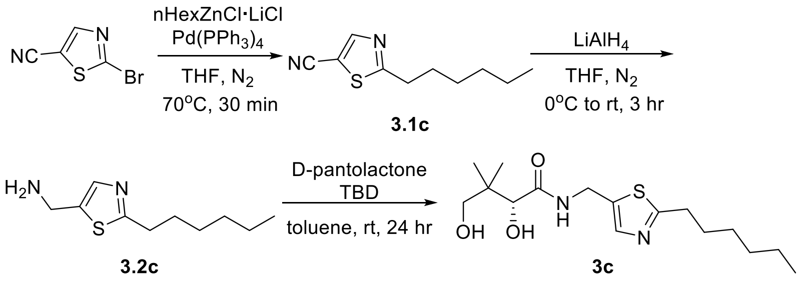


Following procedure H and I, 1-bromohexane (2.11 mL, 15 mmol) was used to generate the Grignard reagent (0.448 M titration, 10 mL used for generation of the organozinc reagent). Purification by flash chromatography gave **3.1c** (520 mg, 89%) as a yellow oil. $R_f$ = 0.26 (Hex/EA = 5:1); $^1$H NMR ($CDCl_3$, 500 MHz): $\delta$ 8.13 (s, 1H), 3.04 (t, $J$ = 7.6 Hz, 2H), 1.79 (p, $J$ = 7.5 Hz, 2H), 1.38 (m, 2H), 1.33-1.27 (m, 4H), 0.88 (t, $J$ = 7.0 Hz, 3H); $^{13}$C NMR ($CDCl_3$, 125 MHz): $\delta$ 178.00, 151.78, 112.16, 105.19, 33.73, 31.42, 29.74, 28.69, 22.51, 14.07; HRMS (ESI$^+$) *m/z* $[C_{10}H_{14}N_2SNa]^+$ calcd: 217.0770; found: 217.0771.

Following procedure J, **3.1c** (520.0 mg, 2.67 mmol) and $LiAlH_4$ (4.0 mL) were reacted. Purification by flash chromatography gave **3.2c** (158 mg, 30%) as a light brown oil. $R_f$ = 0.20 (DCM/MeOH = 10:1); $^1$H NMR ($CDCl_3$, 500 MHz): $\delta$ 7.40 (s, 1H), 4.00 (s, 2H), 2.92 (t, $J$ = 7.6 Hz, 2H), 1.74 (p, $J$ = 7.6 Hz, 2H), 1.65 (bs, 2H), 1.37 (m, 2H), 1.31-1.27 (m, 4H), 0.86 (t, $J$ = 7.0 Hz, 3H); $^{13}$C NMR ($CDCl_3$, 125 MHz): $\delta$ 171.24, 140.44, 138.52, 39.02, 33.74, 31.58, 30.04, 28.82, 22.58, 14.12; HRMS (ESI$^+$) *m/z* $[C_{10}H_{19}N_2S]^+$ calcd: 199.1263; found: 199.1268.

Following procedure K, **3.2c** (158.0 mg, 0.8 mmol), D-pantolactone (208.2 mg, 1.6 mmol), TBD (11.1 mg, 0.08 mmol), and toluene (0.8 mL) were reacted. Purification by flash chromatography gave **3c** (250 mg, 95%) as a sticky yellow oil. $R_f$ = 0.41 (DCM/MeOH = 10:1); $^1$H NMR ($CDCl_3$, 500 MHz): $\delta$ 7.45 (t, $J$ = 6.0 Hz, 1H), 7.39 (s, 1H), 5.06 (bs, 1H), 4.58 (dd, $J$ = 15.5, 5.9 Hz, 1H), 4.51 (dd, $J$ = 15.5, 5.9 Hz, 1H), 4.14 (bs, 1H), 4.03 (s, 1H), 3.46 (d, $J$ = 16.0 Hz, 1H), 3.44 (d, $J$ = 16.0 Hz, 1H), 2.89 (t, $J$ = 7.7 Hz, 2H), 1.70 (m, 2H), 1.34 (m, 2H), 1.30-1.24 (m, 4H), 0.96 (s, 3H), 0.88 (s, 3H), 0.86 (t, $J$ = 7.0 Hz, 3H); $^{13}$C NMR ($CDCl_3$, 125 MHz): $\delta$ 173.58, 172.82, 140.13, 134.61, 77.49, 71.27, 39.43, 35.30, 33.53, 31.51, 29.99, 28.76, 22.56, 21.25, 20.52, 14.10; HRMS (ESI$^+$) *m/z* $[C_{16}H_{29}N_2O_3S]^+$ calcd: 329.1893; found: 329.1894. HPLC: method A $t_R$ = 30.89 min (99%); method B $t_R$ = 19.16 min (99%).

Synthesis of **3d**

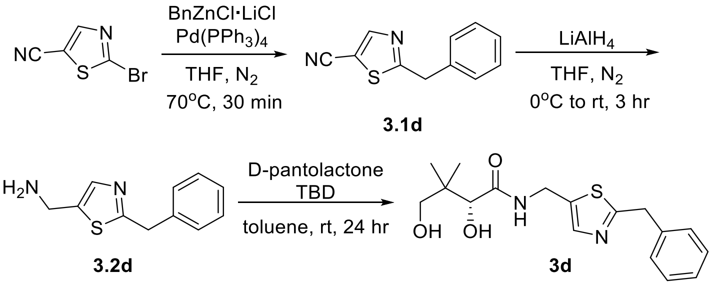

Following procedure H and I, a commercial BnMgCl solution (0.667 M titration, 6.75 mL) was used for generation of the organozinc reagent. Purification by flash chromatography gave **3.1d** (366 mg, 61%) as a yellow solid. $R_f$ = 0.20 (Hex/EA = 5:1); $^1$H NMR ($CDCl_3$, 500 MHz): $\delta$ 8.14 (s, 1H), 7.38-7.29 (m, 5H), 4.35 (s, 2H); $^{13}$C NMR ($CDCl_3$, 125 MHz): $\delta$ 177.22, 151.85, 136.12, 129.11, 129.03, 127.84, 111.87, 106.07, 39.76; HRMS (ESI$^+$) *m/z* $[C_{11}H_8N_2SNa]^+$ calcd: 223.0300; found: 223.0297.

Following procedure J, **3.1d** (366.0 mg, 1.83 mmol) and $LiAlH_4$ (2.74 mL) were reacted. Purification by flash chromatography gave **3.2d** (81 mg, 22%) as a brown solid. $R_f$ = 0.15 (DCM/MeOH = 10:1); $^1$H NMR ($CDCl_3$, 500 MHz): $\delta$ 7.44 (s, 1H), 7.33-7.23 (m, 5H), 4.26 (s, 2H), 3.96 (s, 2H), 1.58 (bs, 2H); $^{13}$C NMR ($CDCl_3$, 125 MHz): $\delta$ 169.93, 141.79, 138.83, 138.09, 129.12, 128.91, 127.21, 40.07, 39.04; HRMS (ESI$^+$) *m/z* $[C_{11}H_{13}N_2S]^+$ calcd: 205.0794; found: 205.0794.

Following procedure K, **3.2d** (81.0 mg, 0.4 mmol), D-pantolactone (104.1 mg, 0.8 mmol), TBD (5.6 mg, 0.04 mmol), and toluene (0.4 mL) were reacted. Purification by flash chromatography gave **3d** (96 mg, 72%) as a sticky yellow oil. $R_f$ = 0.14 (EA); $^1$H NMR ($CDCl_3$, 500 MHz): $\delta$ 7.45 (s, 1H), 7.38 (t, $J$ = 6.2 Hz, 1H), 7.31-7.28 (m, 2H), 7.26-7.23 (m, 3H), 4.52 (dd, $J$ = 15.6, 6.2 Hz, 1H), 4.47 (dd, $J$ = 15.6, 6.2 Hz, 1H), 4.35 (bs, 2H), 4.22 (s, 2H), 4.01 (s, 1H), 3.45 (d, $J$ = 11.4 Hz, 1H), 3.42 (d, $J$ = 11.4 Hz, 1H), 0.94 (s, 3H), 0.87 (s, 3H); $^{13}$C NMR ($CDCl_3$, 125 MHz): $\delta$ 173.46, 171.47, 140.37, 137.52, 135.80, 129.01, 128.95, 127.36, 77.56, 71.30, 39.70, 39.41, 35.26, 21.23, 20.55; HRMS (ESI$^+$) *m/z* $[C_{17}H_{23}N_2O_3S]^+$ calcd: 335.1424; found: 335.1418; HPLC: method A $t_R$ = 15.46 min (95%); method B $t_R$ = 15.80 min (95%).

Synthesis of **3e**

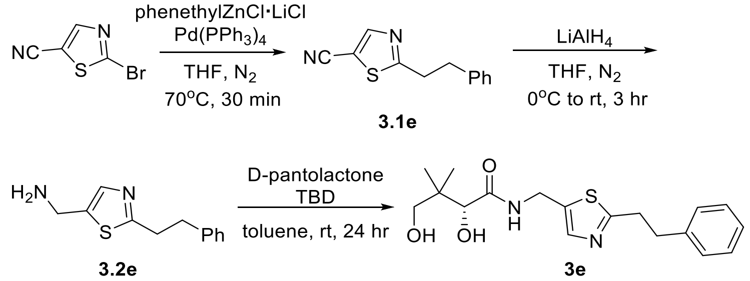


Following procedure H and I, 2-phenethyl bromide (2.05 mL, 15 mmol) was used to generate the Grignard reagent (0.376 M titration, 12 mL used for generation of the organozinc reagent). Purification by flash chromatography gave **3.1e** (421 mg, 65%) as a yellow oil. $R_f$ = 0.63 (Hex/EA = 2:1); $^1$H NMR ($CDCl_3$, 500 MHz): $\delta$ 8.15 (s, 1H), 7.31 (m, 2H), 1.25 (m, 1H), 7.19 (m, 2H), 3.38 (t, $J$ = 7.7 Hz, 2H), 3.14 (t, $J$ = 7.7 Hz, 2H); $^{13}$C NMR ($CDCl_3$, 125 MHz): $\delta$ 176.26, 151.74, 139.25, 128.81, 128.49, 126.87, 112.06, 105.47, 35.40, 35.27; HRMS (ESI$^+$) *m/z* $[C_{12}H_{10}N_2SNa]^+$ calcd: 237.0457; found: 237.0452.

Following procedure J, **3.1e** (421.0 mg, 1.96 mmol) and $LiAlH_4$ (2.9 mL) were reacted. Purification by flash chromatography gave **3.2e** (198 mg, 46%) as a light brown oil. $R_f$ = 0.20 (DCM/MeOH = 10:1); $^1$H NMR ($CDCl_3$, 500 MHz): $\delta$ 7.45 (s, 1H), 7.30-7.27 (m, 2H), 7.22-7.19

(m, 3H), 4.01 (s, 2H), 3.27 (m, 2H), 3.10 (m, 2H), 1.55 (bs, 2H); $^{13}C$ NMR ($CDCl_3$, 125 MHz): $\delta$ 169.64, 140.88, 140.50, 138.59, 128.59, 128.49, 126.40, 39.02, 35.96, 35.36; HRMS ($ESI^+$) *m/z* $[C_{12}H_{15}N_2S]^+$ calcd: 219.0951; found: 219.0954.

Following procedure K, **3.2e** (198.0 mg, 0.9 mmol), D-pantolactone (234.3 mg, 1.8 mmol), TBD (12.5 mg, 0.09 mmol), and toluene (0.9 mL) were reacted. Purification by flash chromatography gave **3e** (262 mg, 84%) as a light yellow solid. $R_f$ = 0.42 (DCM/MeOH = 10:1); $^1H$ NMR ($CDCl_3$, 500 MHz): $\delta$ 7.45-7.43 (m, 2H), 7.26 (m, 2H), 7.20-7.15 (m, 3H), 4.57 (dd, *J* = 15.4, 6.1 Hz, 1H), 4.48 (dd, *J* = 15.4, 6.0 Hz, 1H), 4.36 (bs, 2H), 4.04 (s, 1H), 3.48 (d, *J* = 11.1 Hz, 1H), 3.45 (d, *J* = 11.1 Hz, 1H), 3.22 (m, 2H), 3.04 (m, 2H), 0.97 (s, 3H), 0.90 (s, 3H); $^{13}C$ NMR ($CDCl_3$, 125 MHz): $\delta$ 173.53, 171.22, 140.25, 140.07, 134.94, 128.63, 128.43, 126.52, 77.54, 71.30, 39.41, 35.86, 35.25, 35.11, 21.25, 20.56; HRMS ($ESI^+$) *m/z* $[C_{18}H_{24}N_2O_3SNa]^+$ calcd: 371.1340; found: 371.1392; HPLC method A: $t_R$ = 18.87 min (99%); method B $t_R$ = 16.84 min (99%).

Synthesis of **3f**

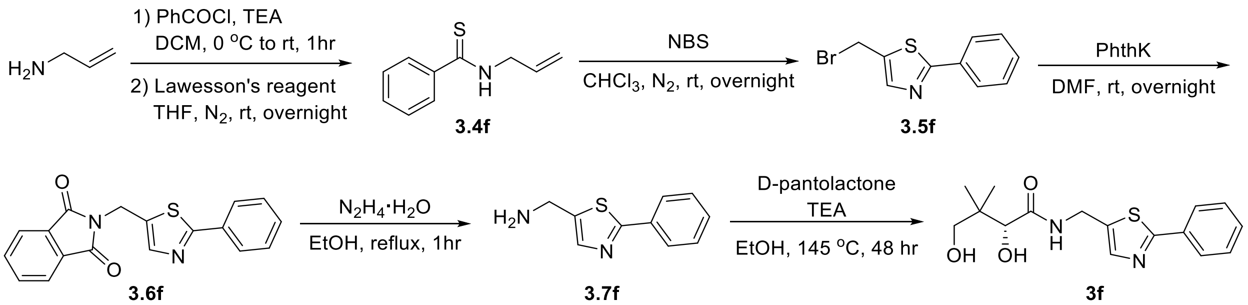


Following general procedure L, benzoyl chloride (375.3 mg, 2.67 mmol) was used to synthesize **3.3f** (368 mg, 86%), which was used without further purification. Following general procedure F, **3.3f** (220.0 mg, 1.24 mmol), Lawesson's reagent (250.8 mg, 0.62 mmol), and THF (25 mL) were reacted. Purification by flash chromatography gave **3.4f** (152 mg, 69%) as a yellow oil. $R_f$ = 0.89 (Hex/EA = 1:1); $^1H$ NMR (500 MHz, $CDCl_3$): $\delta$ 7.74 (m, 2H), 7.62 (bs, 1H), 7.46 (m, 1H), 7.38 (m, 2H), 6.00 (ddt, *J* = 17.1, 10.3, 6.0 Hz, 1H), 5.35 (dq, *J* = 17.2, 1.2 Hz, 1H), 5.29 (dq, *J* = 10.2, 1.2 Hz, 1H), 4.46 (tt, *J* = 5.9, 1.5 Hz, 2H). Characterization matched previously reported data.[15]

Following general procedure M, **3.4f** (126.0 mg, 0.71 mmol), NBS (316.0 mg, 1.78 mmol), and $CHCl_3$ (12 mL) were reacted. Purification by flash chromatography gave **3.5f** (96 mg, 53%) as a white solid. $R_f$ = 0.86 (Hex/EA = 1:1); $^1H$ NMR (500 MHz, $CDCl_3$): $\delta$ 7.92 (m, 2H), 7.78 (s, 1H), 7.46-7.42 (m, 3H), 4.75 (s, 2H). Characterization matched previously reported data.[3]

Following general procedure $G_1$, **3.5f** (96 mg, 0.37 mmol), potassium phthalimide (68.5 mg, 0.37 mmol), and DMF (1 mL) were reacted. No chromatography was needed to give **3.6f** (63 mg, 53%) as a yellow solid. $^1H$ NMR (500 MHz, $CDCl_3$): $\delta$ 7.90-7.86 (m, 4H), 7.86 (s, 1H), 7.73 (m, 2H), 7.42-7.39 (m, 3H), 5.07 (s, 2H); $^{13}C$ NMR (125 MHz, $CDCl_3$): $\delta$ 169.30, 167.34, 143.76, 134.27, 133.49, 132.42, 131.94, 130.16, 128.93, 126.47, 123.60, 33.33; HRMS ($ESI^+$) *m/z* $[C_{18}H_{12}O_2N_2SNa]^+$ calcd: 343.0512; found :343.0499.

Following general procedure $G_2$, **3.6f** (63 mg, 0.2 mmol), $N_2H_4{\cdot}H_2O$ (30.0 mg, 0.6 mmol), and EtOH (3 mL) were reacted. No chromatography was needed to give **3.7f** (19 mg, 50%) as a yellow solid. $^1$H NMR (500 MHz, $CDCl_3$): $\delta$ 7.91 (m, 2H), 7.63 (s, 1H), 7.45-7.39 (m, 3H), 4.10 (s, 2H), 1.68 (bs, 2H); $^{13}$C NMR (125 MHz, $CDCl_3$): $\delta$ 167.65, 141.67, 140.03, 133.84, 129.88, 128.94, 126.32, 39.03; HRMS (ESI$^+$) *m/z* $[C_{10}H_{11}N_2S]^+$ calcd: 191.0637; found: 191.0628.

Following general procedure D, **3.7f** (43.8 mg, 0.23 mmol), D-pantolactone (89.8 mg, 0.69 mmol), TEA (70.0 mg, 0.69 mmol), and EtOH (2.5 mL) were reacted. Purification by flash chromatography gave **3f** (35 mg, 48%) as a yellow oil. $R_f$ = 0.21 (EA); $^1$H NMR (500 MHz, $CDCl_3$): $\delta$ 7.89 (m, 2H, H-12a,b), 7.67 (s, 1H, H-9), 7.44-7.41 (m, 3H), 7.28 (m, 1H), 4.69 (dd, $J$ = 15.5, 6.2 Hz, 1H), 4.64 (dd, $J$ =15.4, 6.3 Hz, 1H), 4.10 (s, 1H), 3.57 (d, $J$ = 11.1 Hz, 1H), 3.50 (d, $J$ = 11.1 Hz, 1H), 1.04 (s, 3H), 0.94 (s, 3H); $^{13}$C NMR (125 MHz, $CDCl_3$): $\delta$ 172.84, 169.00, 142.03, 135.40, 133.46, 130.22, 129.01, 126.44, 77.90, 71.60, 39.43, 35.34, 21.23, 20.37; HRMS (ESI$^+$) *m/z* $[C_{16}H_{20}O_3N_2SNa]^+$ calcd: 343.1087; found: 343.1079; HPLC purity: 81%; method A: $t_R$ = 17.78 min; method B $t_R$ = 8.85 min.

Synthesis of **3g**

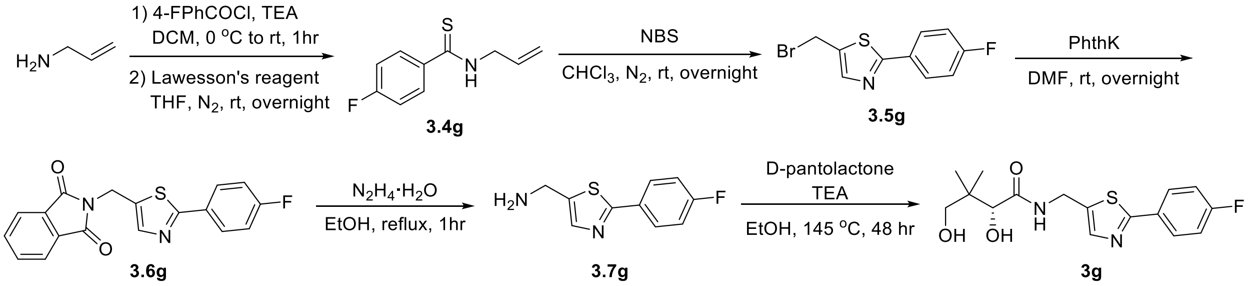


Following general procedure L, 4-fluorobenzoyl chloride (415.0 mg, 2.62 mmol) was used to synthesize **3.3g** (413 mg, 88%), which was used without further purification. Following general procedure F, **3.3g** (200.0 mg, 1.11 mmol), Lawesson's reagent (224.5 mg, 0.56 mmol), and THF (22 mL) were reacted. Purification by flash chromatography gave **3.4g** (102 mg, 47%) as a yellow oil. $R_f$ = 0.85 (Hex/EA = 1:1); $^1$H NMR (500 MHz, $CDCl_3$): $\delta$ 7.75 (m, 2H), 7.62 (bs, 1H), 7.04 (m, 2H), 5.99 (ddt, $J$ = 17.2, 10.2, 6.0 Hz, 1H), 5.34 (dq, $J$ = 17.1, 1.5 Hz, 1H), 5.28 (dq, $J$ = 10.1, 1.4 Hz, 1H), 4.43 (tt, $J$ = 6.1, 1.5 Hz, 2H); $^{19}$F NMR (471 MHz, $CDCl_3$): $\delta$ -108.87; $^{13}$C NMR (125 MHz, $CDCl_3$): $\delta$ 197.87, 164.55 (d, $J$ = 251.3 Hz), 137.87 (d, $J$ =2.5 Hz), 131.77, 128.90 (d, $J$ = 7.5 Hz), 118.92, 115.47 (d, $J$ = 22.5 Hz), 49.20; HRMS (ESI$^+$) *m/z* $[C_{10}H_{10}FNSNa]^+$ calcd: 218.0410; found: 218.0404.

Following general procedure M, **3.4g** (102 mg, 0.53 mmol), NBS (236.7 mg, 1.33 mmol), and $CHCl_3$ (9 mL) were reacted. Purification by flash chromatography gave **3.5g** (22 mg, 15%) as a white solid. $R_f$ = 0.94 (Hex/EA = 1:1); $^1$H NMR (500 MHz, $CDCl_3$): $\delta$ 7.91 (m, 2H), 7.76 (s, 1H), 7.13 (m, 2H), 4.74 (d, $J$ = 0.8 Hz, 2H). Characterization matched previously reported data.[3]

Following general procedure $G_1$, **3.5g** (84.5 mg, 0.31 mmol), potassium phthalimide (57.4 mg, 0.31 mmol), and DMF (1 mL) were reacted. No chromatography was needed to give **3.6g** (53 mg, 51%) as a white solid. $^1$H NMR (500 MHz, $CDCl_3$): $\delta$ 7.89-7.86 (m, 4H), 7.83 (s, 1H), 7.73 (m,

2H), 7.09 (m, 2H), 5.06 (s, 2H); $^{19}$F NMR (471 MHz, $CDCl_3$): $\delta$ -110.28; $^{13}$C NMR (125 MHz, $CDCl_3$): $\delta$ 168.06, 167.43, 163.93 (d, $J$ = 250 Hz), 143.74, 134.29, 132.48, 131.92, 129.86 (d, $J$ = 3.8 Hz), 128.38 (d, $J$ = 8.8 Hz), 123.61, 116.04 (d, $J$ = 22.5 Hz), 33.28; HRMS (ESI$^+$) *m/z* $[C_{18}H_{11}F\text{-}O_2N_2SNa]^+$ calcd: 361.0417; found: 361.0409.

Following general procedure $G_2$, **3.6g** (53.0 mg, 0.16 mmol), $N_2H_4{\cdot}H_2O$ (24.0 mg, 0.48 mmol), and EtOH (2.5 mL) were reacted. No chromatography was needed to give **3.7g** (27 mg, 81%) as a yellow solid. $^1$H NMR (500 MHz, CDCl3): $\delta$ 7.89 (m, 2H), 7.61 (s, 1H), 7.12 (m, 2H), 4.11 (s, 2H), 1.60 (bs, 2H); $^{19}$F NMR (471 MHz, $CDCl_3$): $\delta$ -110.85; $^{13}$C NMR (125 MHz, $CDCl_3$): $\delta$ 166.44, 163.77 (d, $J$ = 248.8 Hz), 141.78, 140.00, 130.23 (d, $J$ = 3.8 Hz), 128.20 (d, $J$ = 8.8 Hz), 116.01 (d, $J$ = 22.5 Hz), 39.02; HRMS (ESI$^+$) *m/z* $[C_{10}H_{10}FN_2S]^+$ calcd: 209.0543; found: 209.0538.

Following general procedure D, **3.7g** (44.0 mg, 0.21 mmol), D-pantolactone (81.7 mg, 0.63 mmol), TEA (63.7 mg, 0.63 mmol), and EtOH (2.5 mL) were reacted. Purification by flash chromatography gave **3g** (35 mg, 48%) as a yellow oil. $R_f$ = 0.26 (EA); $^1$H NMR (500 MHz, $CDCl_3$): $\delta$ 7.86 (m, 2H), 7.63 (s, 1H), 7.33 (t, $J$ = 6.2 Hz, 1H), 7.10 (t, $J$ = 8.6 Hz, 2H), 4.67 (dd, $J$ = 15.5, 6.2 Hz, 1H), 4.61 (dd, $J$ = 15.4, 6.2 Hz, 1H), 4.09 (s, 1H), 3.56 (d, $J$ = 11.2 Hz, 1H), 3.50 (d, $J$ = 11.1 Hz, 1H), 1.02 (s, 3H), 0.93 (s, 3H); $^{19}$F NMR (471 MHz, $CDCl_3$): $\delta$ -110.10; $^{13}$C NMR (125 MHz, $CDCl_3$): $\delta$ 173.00, 167.78, 163.95 (d, $J$ = 248.8 Hz), 141.98, 135.47, 129.8 (d, $J$ = 2.5 Hz), 128.35 (d, $J$ = 8.8 Hz), 116.11 (d, $J$ = 21.3 Hz), 77.85, 71.56, 39.40, 35.30, 21.16, 20.41; HRMS (ESI$^+$) *m/z* $[C_{16}H_{19}FO_3N_2SNa]^+$ calcd: 361.0993; found: 361.1001; HPLC purity: 82%; method A: $t_R$ = 18.28 min; method B $t_R$ = 9.10 min.

Synthesis of **3h**

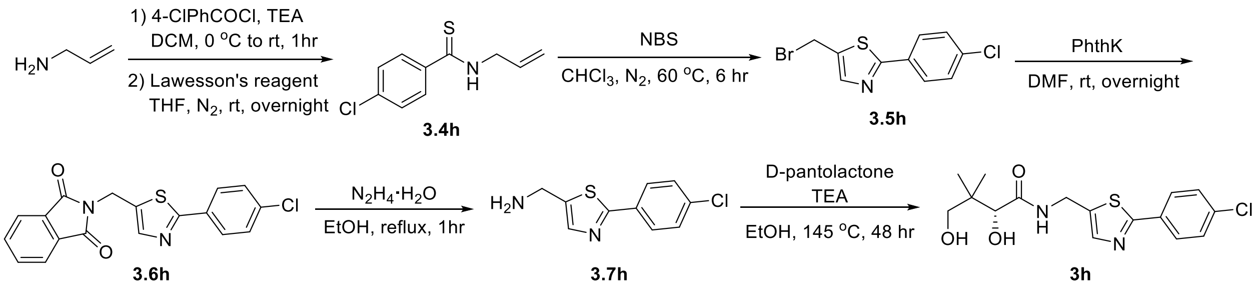


Following general procedure L, 4-chlorobenzoyl chloride (463.8 mg, 2.65 mmol) was reacted to synthesize the **3.3h** (420 mg, 81%), which was used without further purification. Following general procedure F, **3.3h** (220 mg, 1.12 mmol), Lawesson's reagent (224.5 mg, 0.56 mmol), and THF (22 mL) were reacted. Purification by flash chromatography gave **3.4h** (133 mg, 56%) as a yellow oil. $R_f$ = 0.89 (Hex/EA = 1:1); $^1$H NMR (500 MHz, $CDCl_3$): $\delta$ 7.70 (m, 2H), 7.52 (bs, 1H), 7.36 (m, 2H), 6.01 (m, 1H), 5.35 (dq, $J$ = 17.0, 1.5 Hz, 1H), 5.31 (dq, $J$ = 10.2, 1.4 Hz, 1H), 4.46 (tt, $J$ = 5.6, 1.4 Hz, 2H); $^{13}$C NMR (125 MHz, $CDCl_3$): $\delta$ 197.86, 140.02, 137.43, 131.71, 128.72, 128.00, 119.07, 49.21; HRMS (ESI$^+$) *m/z* $[C_{10}H_{10}ClNSNa]^+$ calcd: 234.0115; found: 234.0109.

Following general procedure M, **3.4h** (133 mg, 0.63 mmol), NBS (281.3 mg, 1.58 mmol), and $CHCl_3$ (11 mL) were reacted. The reaction mixture was heated to 60°C for 6 hours. Purification by flash chromatography gave **3.5h** (115 mg, 63%) as a light yellow solid. $R_f$ = 0.92 (Hex/EA =

1:1); $^{1}$H NMR (500 MHz, $CDCl_3$): $\delta$ 7.85 (d, $J$ = 8.7 Hz, 2H), 7.78 (s, 1H), 7.41 (d, $J$ = 8.6 Hz, 2H), 4.74 (d, $J$ = 0.9 Hz, 2H). Characterization matched previously reported data.[3]

Following general procedure $G_1$, **3.5h** (308.0 mg, 1.07 mmol), potassium phthalimide (198.1 mg, 1.07 mmol), and DMF (4 mL) were reacted. Purification by flash chromatography gave **3.6h** (224 mg, 59%) as a yellow solid. $R_f$ = 0.74 (Hex/EA = 1:1); $^{1}$H NMR (500 MHz, $CDCl_3$): $\delta$ 7.87 (m, 2H), 7.85 (s, 1H), 7.82 (d, $J$ = 8.6 Hz, 2H), 7.73 (m, 2H), 7.38 (d, $J$ = 8.5 Hz, 2H), 5.06 (s, 2H); $^{13}$C NMR (125 MHz, $CDCl_3$): $\delta$ 167.90, 167.41, 143.87, 136.14, 134.30, 132.81, 131.99, 131.91, 129.18, 127.64, 123.62, 33.27; HRMS (ESI$^+$) $m/z$ $[C_{18}H_{11}ClO_2N_2SNa]^+$ calcd: 377.0122; found: 377.0115.

Following general procedure $G_2$, **3.6h** (78.0 mg, 0.22 mmol), $N_2H_4 \cdot H_2O$ (33.0 mg, 0.66 mmol), and EtOH (3.5 mL) were reacted. Purification by flash chromatography gave **3.7h** (20 mg, 39%) as light yellow solid. $R_f$ = 0.40 (DCM/MeOH = 9:1); $^{1}$H NMR (500 MHz, $CDCl_3$): $\delta$ 7.85 (d, $J$ = 8.7 Hz, 2H), 7.64 (s, 1H), 7.40 (d, $J$ = 8.7 Hz, 2H), 4.12 (s, 2H), 1.63 (bs, 2H); $^{13}$C NMR (125 MHz, $CDCl_3$): $\delta$ 166.28, 142.11, 140.16, 135.80, 132.36, 129.17, 127.51, 39.01. HRMS (ESI$^+$) $m/z$ $[C_{10}H_{10}ClN_2S]^+$ calcd: 225.0248; found: 225.0241.

Following general procedure D, **3.7h** (20.0 mg, 0.08 mmol), D-pantolactone (31.0 mg, 0.24 mmol), TEA (24.2 mg, 0.24 mmol), and EtOH (1 mL) were reacted. Purification by flash chromatography ($SiO_2$) gave **3h** (10 mg, 35%) as a yellow solid. $R_f$ = 0.13 (EA); $^{1}$H NMR (500 MHz, $CDCl_3$): $\delta$ 7.82 (d, $J$ = 8.7 Hz, 2H), 7.67 (s, 1H), 7.40 (d, $J$ = 8.6 Hz, 2H), 7.30 (t, $J$ = 6.1 Hz, 1H), 4.69 (dd, $J$ = 15.6, 6.1 Hz, 1H), 4.63 (dd, $J$ = 15.6, 6.1 Hz, 1H), 4.10 (s, 1H), 3.56 (d, $J$ = 11.1 Hz, 1H), 3.51 (d, $J$ = 11.1 Hz, 1H), 1.03 (s, 3H), 0.94 (s, 3H); $^{13}$C NMR (125 MHz, $CDCl_3$): $\delta$ 172.86, 167.58, 142.17, 136.16, 135.81, 131.98,129.24, 127.61, 77.91, 71.63, 39.41, 35.32, 21.13, 20.41; HRMS (ESI$^+$) $m/z$ $[C_{16}H_{19}ClO_3N_2SNa]^+$ calcd: 377.0697; found: 377.0683; HPLC purity: 80%; method A: $t_R$ = 19.81 min; method B $t_R$ = 9.83 min.

Synthesis of **3i**

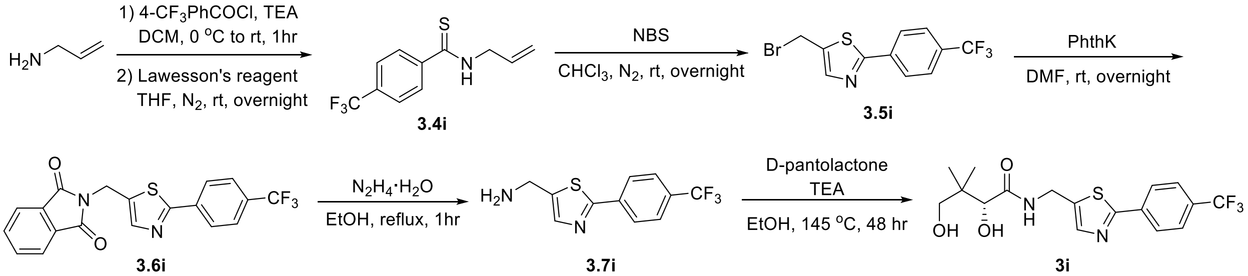


Following general procedure L, 4-trifluoromethylbenzoyl chloride (561.0 mg, 2.69 mmol) was reacted to synthesize the **3.3i** (573 mg, 93%), which was used without further purification. Following general procedure F, **3.3i** (566 mg, 2.47 mmol), Lawesson's reagent (495.1 mg, 1.24 mmol), and THF (48 mL) were reacted. Purification by flash chromatography gave **3.4i** (339 mg, 56%) as a yellow oil. $R_f$ = 0.90 (Hex/EA = 1:1); $^{1}$H NMR (500 MHz, $CDCl_3$): $\delta$ 7.81 (d, $J$ = 8.0 Hz, 2H), 7.69 (bs, 1H), 7.62 (d, $J$ = 8.0 Hz, 2H), 6.00 (ddt, $J$ = 17.1, 10.2, 5.8 Hz, 1H), 5.36 (dq, $J$ = 17.2, 1.5 Hz, 1H), 5.31 (dq, $J$ = 10.2, 1.3 Hz, 1H), 4.45 (tt, $J$ = 5.7, 1.4 Hz, 2H); $^{19}$F NMR (471 MHz, $CDCl_3$) $\delta$ -62.93; $^{13}$C NMR (125 MHz, $CDCl_3$): $\delta$ 197.76, 144.72, 132.55 (q, $J$ = 32.5 Hz),

131.43, 127.09, 125.50 (q, $J$ = 3.8 Hz), 123.67 (q, $J$ = 271.3 Hz), 119.07, 49.20; HRMS (ESI$^+$) $m/z$ $[C_{11}H_{10}F_3NSNa]^+$ calcd: 268.0378; found: 268.0379.

Following general procedure M, **3.4i** (117.7 mg, 0.48 mmol), NBS (214.4 mg, 1.20 mmol), and $CHCl_3$ (8 mL) were reacted. Purification by flash chromatography gave **3.5i** (107 mg, 69%) as a white solid. $R_f$ = 0.89 (Hex/EA = 1:1); $^1$H NMR (500 MHz, $CDCl_3$): $\delta$ 8.03 (d, $J$ = 8.2 Hz, 2H), 7.83 (s, 1H), 7.70 (d, $J$ = 8.2 Hz, 2H), 4.76 (d, $J$ = 0.8 Hz, 2H). Characterization matched previously reported data.[3]

Following general procedure $G_1$, **3.5i** (405.9 mg, 1.26 mmol), potassium phthalimide (233.3 mg, 1.26 mmol), and DMF (5 mL) were reacted. Purification by flash chromatography ($SiO_2$) gave **3.6i** (210 mg, 43%) as a yellow solid. $R_f$ = 0.18 (DCM); $^1$H NMR (500 MHz, $CDCl_3$): $\delta$ 7.99 (d, $J$ = 8.1 Hz, 2H), 7.90 (s, 1H), 7.87 (m, 2H), 7.73 (m, 2H), 7.66 (d, $J$ = 8.2 Hz, 2H), 5.07 (s, 2H); $^{19}$F NMR (470 MHz, $CDCl_3$): $\delta$ -62.83; $^{13}$C NMR (125 MHz, $CDCl_3$): $\delta$ 167.38, 167.25, 144.17, 136.55, 134.33, 133.72 131.87, 131.71 (q, $J$ = 32.5 Hz), 126.64, 125.96 (q, $J$ = 3.8 Hz), 123.85 (q, $J$ = 270 Hz), 123.63, 33.22; HRMS (ESI$^+$) $m/z$ $[C_{19}H_{11}F_3O_2N_2SNa]^+$ calcd: 411.0386; found: 411.0372.

Following general procedure $G_2$, **3.6i** (143.7 mg, 0.37 mmol), $N_2H_4{\cdot}H_2O$ (55.5 mg, 1.11 mmol), and EtOH (6 mL) were reacted. No chromatography was needed to give **3.7i** (76 mg, 80%) as a light yellow solid. $^1$H NMR (500 MHz, $CD_3OD$): $\delta$ 8.04 (d, $J$ = 8.2 Hz, 2H), 7.74 (s, 1H), 7.72 (d, $J$ = 8.3 Hz, 2H), 4.05 (s, 2H); $^{19}$F NMR (471 MHz, $CD_3OD$) $\delta$ -64.24; $^{13}$C NMR (125 MHz, $CD_3OD$) $\delta$ 165.76, 142.55, 140.76, 136.78, 131.13 (q, $J$ = 32.5 Hz), 126.34, 125.71 (q, $J$ = 3.8 Hz), 124.02 (q, $J$ = 270 Hz), 37.52. HRMS (ESI$^+$) $m/z$ $[C_{11}H_{10}F_3N_2S]^+$ calcd: 259.0511; found: 259.0504.

Following general procedure D, **3.7i** (64.6 mg, 0.25 mmol), D-pantolactone (96.9 mg, 0.75 mmol), TEA (75.6 mg, 0.75 mmol), and EtOH (3 mL) were reacted. Purification by flash chromatography gave **3i** (40 mg, 41%) as a colorless oil. $R_f$ = 0.26 (EA); $^1$H NMR (500 MHz, $CD_3OD$): $\delta$ 8.08 (d, $J$ = 8.3 Hz, 2H), 7.79 (s, 1H), 7.76 (d, $J$ = 8.3 Hz, 2H), 4.66 (d, $J$ = 15.3 Hz, 1H), 4.61 (d, $J$ = 15.3 Hz, 1H), 3.96 (s, 1H), 3.49 (d, $J$ = 10.9 Hz, 1H), 3.39 (d, $J$ = 11.0 Hz, 1H), 0.94 (s, 3H), 0.92 (s, 3H); $^{19}$F NMR (471 MHz, $CD_3OD$): $\delta$ -64.33; $^{13}$C NMR (125 MHz, $CD_3OD$): $\delta$ 174.89, 166.45, 142.08, 138.25, 136.73, 131.27 (q, $J$ = 32.5 Hz), 126.43, 125. 76 (q, $J$ = 3.8 Hz), 124.02 (q, $J$ = 270 Hz), 75.93, 68.93, 39.17, 34.47, 19.99, 19.43; HRMS (ESI$^+$) $m/z$ $[C_{17}H_{19}F_3O_3N_2SNa]^+$ calcd: 411.0961; found: 411.0952; HPLC purity: 94%; method A: $t_R$ = 21.36 min; method B $t_R$ = 10.25 min.

Synthesis of **3j**

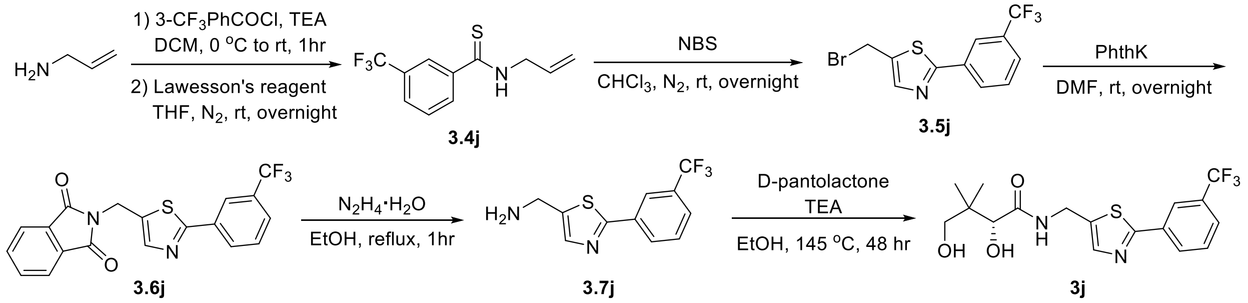

Following general procedure L, 3-trifluoromethylbenzoyl chloride (552.0 mg, 2.65 mmol) was reacted to synthesize the **3.3j** (607 mg, quant.), which was used without further purification. Following general procedure F, **3.3j** (360 mg, 1.57 mmol), Lawesson's reagent (314.7 mg, 0.79 mmol), and THF (30 mL) were reacted. Purification by flash chromatography gave **3.4j** (254 mg, 66%) as a yellow oil. $R_f$ = 0.77 (Hex/EA = 1:1); $^1$H NMR (500 MHz, $CDCl_3$): $\delta$ 7.97 (s, 1H), 7.93 (d, $J$ = 7.9 Hz, 1H), 7.71 (d, $J$ = 7.9 Hz, 1H), 7.60 (bs, 1H), 7.52 (t, $J$ = 7.8 Hz, 1H), 6.02 (ddt, $J$ = 17.1, 10.2, 6.1 Hz, 1H), 5.38 (dq, $J$ = 17.2, 1.6 Hz, 1H), 5.33 (dq, $J$ = 10.2, 1.6 Hz, 1H), 4.47 (tt, $J$ = 6.0, 1.5 Hz, 2H); $^{19}$F NMR (471 MHz, $CDCl_3$): $\delta$ -62.70; $^{13}$C NMR (125 MHz, $CDCl_3$): $\delta$ 197.59, 142.32, 131.45, 130.93 (q, $J$ = 32.5 Hz), 129.91, 129.15, 127.54 (q, $J$ = 3.8 Hz), 123.64 (q, $J$ = 271.3 Hz), 123.61 (q, $J$ = 3.8 Hz), 119.19, 49.28. HRMS (ESI$^+$) *m/z* $[C_{11}H_{11}F_3NS]^+$ calcd: 246.0559; found: 246.0560.

Following general procedure M, **3.4j** (117.7 mg, 0.48 mmol), NBS (214.4 mg, 1.20 mmol), and $CHCl_3$ (8 mL) were reacted. Purification by flash chromatography gave **3.5j** (107 mg, 69%) as a white solid. $^1$H NMR (500 MHz, $CDCl_3$): $\delta$ 8.03 (d, $J$ = 8.2 Hz, 2H), 7.83 (s, 1H), 7.70 (d, $J$ = 8.2 Hz, 2H), 4.76 (d, $J$ = 0.8 Hz, 2H). Characterization matched previously reported data.[3]

Following general procedure $G_1$, **3.5j** (235.0 mg, 0.73 mmol), potassium phthalimide (135.1 mg, 0.73 mmol), and DMF (3 mL) were reacted. Purification by flash chromatography gave **3.6j** (173 mg, 61%) as a yellow solid. $R_f$ = 0.19 (Hex/EA = 4:1); $^1$H NMR (500 MHz, $CDCl_3$): $\delta$ 8.13 (s, 1H), 8.02 (d, $J$ = 8.0 Hz, 1H), 7.87 (s, 1H), 7.84 (m, 2H), 7.70 (m, 2H), 7.61 (d, $J$ = 7.9 Hz, 1H), 7.50 (t, $J$ = 8.0 Hz, 1H), 5.06 (s, 2H); $^{19}$F NMR (471 MHz, $CDCl_3$): $\delta$ -62.86; $^{13}$C NMR (125 MHz, $CDCl_3$): $\delta$ 167.36, 167.27, 144.00, 134.30, 134.16, 133.48, 131.85, 131.46 (q, $J$ = 32.5 Hz), 129.52, 129.50, 126.55 (q, $J$ = 3.8 Hz), 123.74 (q, $J$ = 270 Hz), 123.59, 123.18 (q, $J$ = 3.8 Hz), 33.22; HRMS (ESI$^+$) *m/z* $[C_{19}H_{11}F_3O_2N_2SNa]^+$ calcd: 411.0386; found: 411.0382.

Following general procedure $G_2$, **3.6j** (101.0 mg, 0.26 mmol), $N_2H_4 \cdot H_2O$ (38.9 mg, 0.78 mmol), and EtOH (4 mL) were reacted. No chromatography was needed to give **3.7j** (46 mg, 69%) as a yellow oil. $^1$H NMR (500 MHz, $CDCl_3$): $\delta$ 8.20 (s, 1H), 8.06 (d, $J$ = 7.9 Hz, 1H), 7.67 (s, 1H), 7.64 (d, $J$ = 7.8 Hz, 1H), 7.55 (t, $J$ = 7.8 Hz, 1H), 4.13 (s, 2H), 1.60 (bs, 2H); $^{19}$F NMR (471 MHz, $CDCl_3$): $\delta$ -62.84; $^{13}$C NMR (125 MHz, $CDCl_3$): $\delta$ 165.67, 142.83, 140.28, 134.56, 131.49 (q, $J$ = 31.3 Hz), 129.49, 129.42, 126.26 (q, $J$ = 3.8 Hz), 123.83 (q, $J$ = 271.3 Hz), 123.04 (q, $J$ = 3.8 Hz), 38.99; HRMS (ESI$^+$) *m/z* $[C_{11}H_{10}F_3N_2S]^+$ calcd: 259.0511; found: 259.0500.

Following general procedure D, **3.7j** (25.8 mg, 0.10 mmol), D-pantolactone (38.8 mg, 0.3 mmol), TEA (30.3 mg, 0.3 mmol), and EtOH (1.5 mL) were reacted. Purification by flash chromatography gave **3j** (23 mg, 59%) as a colorless oil. $R_f$ = 0.26 (EA); NMR (500 MHz, $CDCl_3$): $\delta$ 8.16 (s, 1H), 8.02 (d, $J$ = 7.8 Hz, 1H), 7.69 (s, 1H), 7.65 (d, $J$ = 7.8 Hz, 1H), 7.54 (t, $J$ = 7.8 Hz, 1H), 7.42 (t, $J$ = 6.2 Hz, 1H), 4.68 (dd, $J$ = 15.5, 6.2 Hz), 4.63 (dd, $J$ = 15.4, 6.1 Hz, 1H), 4.30 (bs, 1H), 4.10 (s, 1H), 3.54 (d, $J$ = 11.1 Hz, 1H), 3.50 (d, $J$ = 11.1 Hz, 1H), 3.38 (bs, 1H), 1.01 (s, 3H), 0.93 (s, 3H); $^{19}$F NMR (471 MHz, $CDCl_3$): $\delta$ -62.83; $^{13}$C NMR (125 MHz, $CDCl_3$): $\delta$ 173.14, 167.02, 142.38, 136.47, 134.12, 131.56 (q, $J$ = 32.5 Hz), 129.58, 129.55, 126.61 (q, $J$ = 3.8 Hz), 123.75 (q, $J$ = 271.3 Hz), 123.13 (q, $J$ = 3.8 Hz), 77.78, 71.49, 39.38, 35.28, 21.06, 20.46; HRMS (ESI$^+$) *m/z* $[C_{17}H_{19}F_3O_3N_2SNa]^+$ calcd: 411.0961, found: 411.0945; HPLC purity: 90%; method A: $t_R$ = 21.06 min; method B $t_R$ = 10.18 min.

#### 4. Computational Studies

The conformational ensemble summary is given below. For detailed output files and conformational ensembles, please refer to the additional uploaded files.

Ensemble summary for Figure **4a**

| Conformer | Energy (kcal/mol) | Degen. | % total | % cumul. |
|---|---|---|---|---|
| 0 | 0.000 | 1 | 3.03 | 3.03 |
| 1 | 0.001 | 1 | 3.03 | 6.06 |
| 2 | 0.002 | 1 | 3.02 | 9.08 |
| 3 | 0.027 | 1 | 2.90 | 11.98 |
| 4 | 0.054 | 1 | 2.77 | 14.74 |
| 5 | 0.424 | 1 | 1.48 | 16.23 |
| 6 | 0.427 | 1 | 1.48 | 17.70 |
| 7 | 0.430 | 1 | 1.47 | 19.17 |
| 8 | 0.430 | 1 | 1.47 | 20.64 |
| 9 | 0.430 | 1 | 1.47 | 22.11 |
| 10 | 0.435 | 1 | 1.45 | 23.56 |
| 11 | 0.435 | 1 | 1.45 | 25.01 |
| 12 | 0.436 | 1 | 1.45 | 26.47 |
| 13 | 0.443 | 1 | 1.44 | 27.90 |
| 14 | 0.456 | 1 | 1.40 | 29.31 |
| 15 | 0.465 | 1 | 1.38 | 30.69 |
| 16 | 0.492 | 1 | 1.32 | 32.01 |
| 17 | 0.503 | 1 | 1.30 | 33.31 |
| 18 | 0.505 | 1 | 1.29 | 34.60 |
| 19 | 0.512 | 1 | 1.28 | 35.88 |
| 20 | 0.512 | 1 | 1.28 | 37.16 |
| 21 | 0.517 | 1 | 1.27 | 38.43 |
| 22 | 0.520 | 1 | 1.26 | 39.69 |
| 23 | 0.522 | 1 | 1.26 | 40.94 |
| 24 | 0.525 | 1 | 1.25 | 42.19 |
| 25 | 0.527 | 1 | 1.25 | 43.44 |
| 26 | 0.527 | 1 | 1.25 | 44.69 |
| 27 | 0.528 | 1 | 1.24 | 45.93 |
| 28 | 0.531 | 1 | 1.24 | 47.17 |
| 29 | 0.535 | 1 | 1.23 | 48.40 |
| 30 | 0.547 | 1 | 1.20 | 49.60 |
| 31 | 0.548 | 1 | 1.20 | 50.80 |
| 32 | 0.550 | 1 | 1.20 | 52.00 |
| 33 | 0.551 | 1 | 1.20 | 53.20 |

| | | | | |
|---|---|---|---|---|
| 34 | 0.554 | 1 | 1.19 | 54.39 |
| 35 | 0.559 | 1 | 1.18 | 55.57 |
| 36 | 0.560 | 1 | 1.18 | 56.75 |
| 37 | 0.561 | 1 | 1.18 | 57.92 |
| 38 | 0.563 | 1 | 1.17 | 59.10 |
| 39 | 0.567 | 1 | 1.17 | 60.26 |
| 40 | 0.573 | 1 | 1.15 | 61.41 |
| 41 | 0.573 | 1 | 1.15 | 62.57 |
| 42 | 0.576 | 1 | 1.15 | 63.71 |
| 43 | 0.577 | 1 | 1.15 | 64.86 |
| 44 | 0.585 | 1 | 1.13 | 65.99 |
| 45 | 0.588 | 1 | 1.12 | 67.11 |
| 46 | 0.590 | 1 | 1.12 | 68.23 |
| 47 | 0.594 | 1 | 1.11 | 69.35 |
| 48 | 0.599 | 1 | 1.10 | 70.45 |
| 49 | 0.600 | 1 | 1.10 | 71.55 |
| 50 | 0.601 | 1 | 1.10 | 72.65 |
| 51 | 0.601 | 1 | 1.10 | 73.75 |
| 52 | 0.604 | 1 | 1.09 | 74.84 |
| 53 | 0.615 | 1 | 1.07 | 75.91 |
| 54 | 0.708 | 1 | 0.92 | 76.83 |
| 55 | 0.807 | 1 | 0.78 | 77.61 |
| 56 | 0.840 | 1 | 0.73 | 78.34 |
| 57 | 0.841 | 1 | 0.73 | 79.08 |
| 58 | 0.842 | 1 | 0.73 | 79.81 |
| 59 | 0.845 | 1 | 0.73 | 80.54 |
| 60 | 0.848 | 1 | 0.72 | 81.26 |
| 61 | 0.848 | 1 | 0.72 | 81.99 |
| 62 | 0.851 | 1 | 0.72 | 82.71 |
| 63 | 0.854 | 1 | 0.72 | 83.43 |
| 64 | 0.883 | 1 | 0.68 | 84.11 |
| 65 | 0.886 | 1 | 0.68 | 84.79 |
| 66 | 0.889 | 1 | 0.68 | 85.46 |
| 67 | 0.889 | 1 | 0.68 | 86.14 |
| 68 | 0.889 | 1 | 0.68 | 86.82 |
| 69 | 0.894 | 1 | 0.67 | 87.49 |
| 70 | 0.896 | 1 | 0.67 | 88.15 |
| 71 | 0.906 | 1 | 0.66 | 88.81 |
| 72 | 0.908 | 1 | 0.65 | 89.47 |
| 73 | 0.912 | 1 | 0.65 | 90.12 |
| 74 | 0.912 | 1 | 0.65 | 90.77 |
| 75 | 0.913 | 1 | 0.65 | 91.42 |
| 76 | 0.913 | 1 | 0.65 | 92.07 |

| | | | | |
|---|---|---|---|---|
| 77 | 0.914 | 1 | 0.65 | 92.72 |
| 78 | 0.919 | 1 | 0.64 | 93.36 |
| 79 | 0.922 | 1 | 0.64 | 94.00 |
| 80 | 0.923 | 1 | 0.64 | 94.64 |
| 81 | 0.940 | 1 | 0.62 | 95.26 |
| 82 | 0.994 | 1 | 0.57 | 95.82 |
| 83 | 0.996 | 1 | 0.56 | 96.39 |
| 84 | 0.998 | 1 | 0.56 | 96.95 |
| 85 | 1.006 | 1 | 0.55 | 97.51 |
| 86 | 1.014 | 1 | 0.55 | 98.05 |
| 87 | 1.048 | 1 | 0.52 | 98.57 |
| 88 | 1.162 | 1 | 0.43 | 99.00 |
| 89 | 1.253 | 1 | 0.37 | 99.36 |
| 90 | 1.289 | 1 | 0.34 | 99.71 |
| 91 | 1.738 | 1 | 0.16 | 99.87 |
| 92 | 1.864 | 1 | 0.13 | 100.00 |

Conformers below 3 kcal/mol: 93
Lowest energy conformer : -36.435573 Eh

Ensemble summary for Figure **4b**

| Conformer | Energy (kcal/mol) | Degen. | % total | % cumul. |
|---|---|---|---|---|
| 0 | 0.000 | 1 | 6.68 | 6.68 |
| 1 | 0.428 | 1 | 3.24 | 9.93 |
| 2 | 0.441 | 1 | 3.18 | 13.11 |
| 3 | 0.444 | 1 | 3.16 | 16.27 |
| 4 | 0.451 | 1 | 3.12 | 19.39 |
| 5 | 0.454 | 1 | 3.11 | 22.50 |
| 6 | 0.545 | 1 | 2.67 | 25.16 |
| 7 | 0.547 | 1 | 2.66 | 27.82 |
| 8 | 0.548 | 1 | 2.65 | 30.47 |
| 9 | 0.548 | 1 | 2.65 | 33.12 |
| 10 | 0.548 | 1 | 2.65 | 35.77 |
| 11 | 0.551 | 1 | 2.64 | 38.41 |
| 12 | 0.552 | 1 | 2.63 | 41.04 |
| 13 | 0.553 | 1 | 2.63 | 43.67 |
| 14 | 0.554 | 1 | 2.62 | 46.30 |
| 15 | 0.555 | 1 | 2.62 | 48.92 |
| 16 | 0.563 | 1 | 2.59 | 51.51 |
| 17 | 0.565 | 1 | 2.57 | 54.08 |
| 18 | 0.566 | 1 | 2.57 | 56.65 |
| 19 | 0.575 | 1 | 2.53 | 59.18 |

| | | | | |
|---|---|---|---|---|
| 20 | 0.578 | 1 | 2.52 | 61.70 |
| 21 | 0.588 | 1 | 2.48 | 64.18 |
| 22 | 0.589 | 1 | 2.47 | 66.65 |
| 23 | 0.592 | 1 | 2.46 | 69.11 |
| 24 | 0.599 | 1 | 2.43 | 71.55 |
| 25 | 0.603 | 1 | 2.42 | 73.97 |
| 26 | 0.787 | 1 | 1.77 | 75.74 |
| 27 | 0.801 | 1 | 1.73 | 77.47 |
| 28 | 0.834 | 1 | 1.64 | 79.10 |
| 29 | 0.843 | 1 | 1.61 | 80.72 |
| 30 | 0.950 | 1 | 1.34 | 82.06 |
| 31 | 0.955 | 1 | 1.33 | 83.39 |
| 32 | 0.957 | 1 | 1.33 | 84.72 |
| 33 | 0.958 | 1 | 1.33 | 86.05 |
| 34 | 0.961 | 1 | 1.32 | 87.37 |
| 35 | 0.962 | 1 | 1.32 | 88.69 |
| 36 | 0.963 | 1 | 1.32 | 90.00 |
| 37 | 0.963 | 1 | 1.31 | 91.32 |
| 38 | 0.968 | 1 | 1.31 | 92.62 |
| 39 | 0.974 | 1 | 1.29 | 93.91 |
| 40 | 0.975 | 1 | 1.29 | 95.20 |
| 41 | 0.984 | 1 | 1.27 | 96.47 |
| 42 | 0.991 | 1 | 1.25 | 97.73 |
| 43 | 1.063 | 1 | 1.11 | 98.84 |
| 44 | 1.650 | 1 | 0.41 | 99.25 |
| 45 | 1.655 | 1 | 0.41 | 99.66 |
| 46 | 1.980 | 1 | 0.24 | 99.90 |
| 47 | 2.482 | 1 | 0.10 | 100.00 |

Conformers below 3 kcal/mol: 48
Lowest energy conformer : -44.989032 Eh

Ensemble summary for Figure **4c**

| Conformer | Energy (kcal/mol) | Degen. | % total | % cumul. |
|---|---|---|---|---|
| 0 | 0.000 | 1 | 6.00 | 6.00 |
| 1 | 0.023 | 1 | 5.77 | 11.77 |
| 2 | 0.026 | 1 | 5.74 | 17.51 |
| 3 | 0.052 | 1 | 5.50 | 23.01 |
| 4 | 0.054 | 1 | 5.48 | 28.49 |
| 5 | 0.059 | 1 | 5.43 | 33.92 |
| 6 | 0.072 | 1 | 5.31 | 39.23 |
| 7 | 0.082 | 1 | 5.23 | 44.46 |

| | | | | |
|---|---|---|---|---|
| 8 | 0.108 | 1 | 5.00 | 49.46 |
| 9 | 0.114 | 1 | 4.95 | 54.41 |
| 10 | 0.127 | 1 | 4.84 | 59.26 |
| 11 | 0.136 | 1 | 4.77 | 64.02 |
| 12 | 0.149 | 1 | 4.66 | 68.69 |
| 13 | 0.387 | 1 | 3.12 | 71.81 |
| 14 | 0.453 | 1 | 2.79 | 74.60 |
| 15 | 0.870 | 1 | 1.38 | 75.98 |
| 16 | 0.877 | 1 | 1.36 | 77.35 |
| 17 | 0.878 | 1 | 1.36 | 78.71 |
| 18 | 0.884 | 1 | 1.35 | 80.06 |
| 19 | 0.894 | 1 | 1.33 | 81.39 |
| 20 | 0.898 | 1 | 1.32 | 82.71 |
| 21 | 0.958 | 1 | 1.19 | 83.90 |
| 22 | 0.985 | 1 | 1.14 | 85.03 |
| 23 | 1.028 | 1 | 1.06 | 86.09 |
| 24 | 1.036 | 1 | 1.04 | 87.14 |
| 25 | 1.041 | 1 | 1.04 | 88.17 |
| 26 | 1.046 | 1 | 1.03 | 89.20 |
| 27 | 1.049 | 1 | 1.02 | 90.22 |
| 28 | 1.051 | 1 | 1.02 | 91.24 |
| 29 | 1.257 | 1 | 0.72 | 91.96 |
| 30 | 1.272 | 1 | 0.70 | 92.66 |
| 31 | 1.289 | 1 | 0.68 | 93.34 |
| 32 | 1.292 | 1 | 0.68 | 94.02 |
| 33 | 1.303 | 1 | 0.67 | 94.68 |
| 34 | 1.312 | 1 | 0.66 | 95.34 |
| 35 | 1.333 | 1 | 0.63 | 95.97 |
| 36 | 1.337 | 1 | 0.63 | 96.60 |
| 37 | 1.340 | 1 | 0.63 | 97.23 |
| 38 | 1.457 | 1 | 0.51 | 97.74 |
| 39 | 1.729 | 1 | 0.32 | 98.06 |
| 40 | 1.846 | 1 | 0.27 | 98.33 |
| 41 | 1.860 | 1 | 0.26 | 98.59 |
| 42 | 1.867 | 1 | 0.26 | 98.84 |
| 43 | 2.131 | 1 | 0.16 | 99.01 |
| 44 | 2.133 | 1 | 0.16 | 99.17 |
| 45 | 2.156 | 1 | 0.16 | 99.33 |
| 46 | 2.217 | 1 | 0.14 | 99.47 |
| 47 | 2.241 | 1 | 0.14 | 99.61 |
| 48 | 2.244 | 1 | 0.14 | 99.75 |
| 49 | 2.294 | 1 | 0.12 | 99.87 |
| 50 | 2.422 | 1 | 0.10 | 99.97 |

| | | | | |
|---|---|---|---|---|
| 51 | 3.153 | 1 | 0.03 | 100.00 |

Conformers below 3 kcal/mol: 51
Lowest energy conformer    : -36.438128 Eh

Ensemble summary for Figure **4d**

### Final ensemble info #

| Conformer | Energy (kcal/mol) | Degen. | % total | % cumul. |
|---|---|---|---|---|
| 0 | 0.000 | 1 | 15.99 | 15.99 |
| 1 | 0.001 | 1 | 15.96 | 31.95 |
| 2 | 0.013 | 1 | 15.63 | 47.58 |
| 3 | 0.025 | 1 | 15.33 | 62.91 |
| 4 | 0.362 | 1 | 8.67 | 71.58 |
| 5 | 0.367 | 1 | 8.61 | 80.19 |
| 6 | 0.721 | 1 | 4.73 | 84.93 |
| 7 | 0.870 | 1 | 3.68 | 88.61 |
| 8 | 0.875 | 1 | 3.65 | 92.25 |
| 9 | 1.273 | 1 | 1.86 | 94.12 |
| 10 | 1.339 | 1 | 1.67 | 95.78 |
| 11 | 1.346 | 1 | 1.65 | 97.43 |
| 12 | 1.869 | 1 | 0.68 | 98.12 |
| 13 | 2.120 | 1 | 0.45 | 98.56 |
| 14 | 2.154 | 1 | 0.42 | 98.98 |
| 15 | 2.240 | 1 | 0.36 | 99.35 |
| 16 | 2.250 | 1 | 0.36 | 99.71 |
| 17 | 2.614 | 1 | 0.19 | 99.90 |
| 18 | 3.018 | 1 | 0.10 | 100.00 |

Conformers below 3 kcal/mol: 18
Lowest energy conformer    : -44.993184 Eh

Ensemble summary for Figure **4e**

| Conformer | Energy (kcal/mol) | Degen. | % total | % cumul. |
|---|---|---|---|---|
| 0 | 0.000 | 1 | 4.98 | 4.98 |
| 1 | 0.006 | 1 | 4.93 | 9.91 |
| 2 | 0.019 | 1 | 4.82 | 14.73 |
| 3 | 0.022 | 1 | 4.80 | 19.53 |
| 4 | 0.024 | 1 | 4.79 | 24.32 |
| 5 | 0.027 | 1 | 4.76 | 29.08 |
| 6 | 0.088 | 1 | 4.29 | 33.37 |
| 7 | 0.142 | 1 | 3.92 | 37.29 |

| | | | | |
|---|---|---|---|---|
| 8 | 0.149 | 1 | 3.87 | 41.17 |
| 9 | 0.177 | 1 | 3.70 | 44.86 |
| 10 | 0.182 | 1 | 3.67 | 48.53 |
| 11 | 0.192 | 1 | 3.60 | 52.13 |
| 12 | 0.193 | 1 | 3.60 | 55.72 |
| 13 | 0.362 | 1 | 2.70 | 58.43 |
| 14 | 0.379 | 1 | 2.63 | 61.05 |
| 15 | 0.418 | 1 | 2.46 | 63.51 |
| 16 | 0.468 | 1 | 2.26 | 65.78 |
| 17 | 0.547 | 1 | 1.98 | 67.75 |
| 18 | 0.574 | 1 | 1.89 | 69.64 |
| 19 | 0.575 | 1 | 1.89 | 71.53 |
| 20 | 0.629 | 1 | 1.72 | 73.25 |
| 21 | 0.687 | 1 | 1.56 | 74.82 |
| 22 | 0.758 | 1 | 1.38 | 76.20 |
| 23 | 0.760 | 1 | 1.38 | 77.58 |
| 24 | 0.767 | 1 | 1.36 | 78.95 |
| 25 | 0.789 | 1 | 1.32 | 80.26 |
| 26 | 0.912 | 1 | 1.07 | 81.33 |
| 27 | 0.915 | 1 | 1.06 | 82.39 |
| 28 | 0.918 | 1 | 1.06 | 83.45 |
| 29 | 0.954 | 1 | 1.00 | 84.45 |
| 30 | 1.013 | 1 | 0.90 | 85.35 |
| 31 | 1.022 | 1 | 0.89 | 86.24 |
| 32 | 1.027 | 1 | 0.88 | 87.12 |
| 33 | 1.092 | 1 | 0.79 | 87.90 |
| 34 | 1.096 | 1 | 0.78 | 88.69 |
| 35 | 1.097 | 1 | 0.78 | 89.47 |
| 36 | 1.100 | 1 | 0.78 | 90.25 |
| 37 | 1.103 | 1 | 0.77 | 91.02 |
| 38 | 1.176 | 1 | 0.68 | 91.71 |
| 39 | 1.177 | 1 | 0.68 | 92.39 |
| 40 | 1.178 | 1 | 0.68 | 93.07 |
| 41 | 1.178 | 1 | 0.68 | 93.75 |
| 42 | 1.193 | 1 | 0.67 | 94.42 |
| 43 | 1.194 | 1 | 0.66 | 95.08 |
| 44 | 1.197 | 1 | 0.66 | 95.75 |
| 45 | 1.427 | 1 | 0.45 | 96.19 |
| 46 | 1.430 | 1 | 0.45 | 96.64 |
| 47 | 1.440 | 1 | 0.44 | 97.08 |
| 48 | 1.469 | 1 | 0.42 | 97.49 |
| 49 | 1.473 | 1 | 0.41 | 97.91 |
| 50 | 1.651 | 1 | 0.31 | 98.22 |

| | | | | |
|---|---|---|---|---|
| 51 | 1.668 | 1 | 0.30 | 98.51 |
| 52 | 1.686 | 1 | 0.29 | 98.80 |
| 53 | 1.853 | 1 | 0.22 | 99.02 |
| 54 | 1.865 | 1 | 0.21 | 99.24 |
| 55 | 1.866 | 1 | 0.21 | 99.45 |
| 56 | 1.869 | 1 | 0.21 | 99.66 |
| 57 | 1.979 | 1 | 0.18 | 99.84 |
| 58 | 2.031 | 1 | 0.16 | 100.00 |

Conformers below 3 kcal/mol: 59
Lowest energy conformer : -36.436518 Eh

Ensemble summary for Figure **4f**

| Conformer | Energy (kcal/mol) | Degen. | % total | % cumul. |
|---|---|---|---|---|
| 0 | 0.000 | 1 | 18.42 | 18.42 |
| 1 | 0.005 | 1 | 18.29 | 36.71 |
| 2 | 0.169 | 1 | 13.84 | 50.55 |
| 3 | 0.412 | 1 | 9.20 | 59.75 |
| 4 | 0.482 | 1 | 8.17 | 67.92 |
| 5 | 0.512 | 1 | 7.77 | 75.69 |
| 6 | 0.700 | 1 | 5.65 | 81.34 |
| 7 | 0.776 | 1 | 4.98 | 86.32 |
| 8 | 0.937 | 1 | 3.79 | 90.11 |
| 9 | 0.945 | 1 | 3.74 | 93.84 |
| 10 | 0.949 | 1 | 3.71 | 97.55 |
| 11 | 1.700 | 1 | 1.05 | 98.60 |
| 12 | 1.929 | 1 | 0.71 | 99.31 |
| 13 | 1.946 | 1 | 0.69 | 100.00 |

Conformers below 3 kcal/mol: 14
Lowest energy conformer : -44.991719 Eh

Ensemble summary for Figure **4g**

| Conformer | Energy (kcal/mol) | Degen. | % total | % cumul. |
|---|---|---|---|---|
| 0 | 0.000 | 1 | 40.62 | 40.62 |
| 1 | 0.769 | 1 | 11.09 | 51.71 |
| 2 | 0.821 | 1 | 10.16 | 61.87 |
| 3 | 1.223 | 1 | 5.15 | 67.02 |
| 4 | 1.446 | 1 | 3.54 | 70.56 |
| 5 | 1.829 | 1 | 1.85 | 72.42 |
| 6 | 1.927 | 1 | 1.57 | 73.99 |

| | | | | |
|---|---|---|---|---|
| 7 | 1.937 | 1 | 1.55 | 75.53 |
| 8 | 1.967 | 1 | 1.47 | 77.00 |
| 9 | 1.973 | 1 | 1.45 | 78.46 |
| 10 | 2.025 | 1 | 1.33 | 79.79 |
| 11 | 2.162 | 1 | 1.06 | 80.85 |
| 12 | 2.165 | 1 | 1.05 | 81.90 |
| 13 | 2.169 | 1 | 1.04 | 82.94 |
| 14 | 2.173 | 1 | 1.04 | 83.98 |
| 15 | 2.181 | 1 | 1.02 | 85.00 |
| 16 | 2.182 | 1 | 1.02 | 86.02 |
| 17 | 2.183 | 1 | 1.02 | 87.04 |
| 18 | 2.185 | 1 | 1.02 | 88.06 |
| 19 | 2.192 | 1 | 1.00 | 89.06 |
| 20 | 2.205 | 1 | 0.98 | 90.05 |
| 21 | 2.207 | 1 | 0.98 | 91.03 |
| 22 | 2.305 | 1 | 0.83 | 91.86 |
| 23 | 2.391 | 1 | 0.72 | 92.57 |
| 24 | 2.410 | 1 | 0.70 | 93.27 |
| 25 | 2.639 | 1 | 0.47 | 93.74 |
| 26 | 2.648 | 1 | 0.47 | 94.21 |
| 27 | 2.661 | 1 | 0.46 | 94.66 |
| 28 | 2.793 | 1 | 0.36 | 95.03 |
| 29 | 2.795 | 1 | 0.36 | 95.39 |
| 30 | 2.796 | 1 | 0.36 | 95.75 |
| 31 | 3.054 | 1 | 0.23 | 95.99 |
| 32 | 3.060 | 1 | 0.23 | 96.22 |
| 33 | 3.315 | 1 | 0.15 | 96.37 |
| 34 | 3.315 | 1 | 0.15 | 96.52 |
| 35 | 3.324 | 1 | 0.15 | 96.67 |
| 36 | 3.361 | 1 | 0.14 | 96.81 |
| 37 | 3.431 | 1 | 0.12 | 96.93 |
| 38 | 3.445 | 1 | 0.12 | 97.06 |
| 39 | 3.455 | 1 | 0.12 | 97.18 |
| 40 | 3.456 | 1 | 0.12 | 97.29 |
| 41 | 3.458 | 1 | 0.12 | 97.41 |
| 42 | 3.459 | 1 | 0.12 | 97.53 |
| 43 | 3.461 | 1 | 0.12 | 97.65 |
| 44 | 3.470 | 1 | 0.12 | 97.77 |
| 45 | 3.548 | 1 | 0.10 | 97.87 |
| 46 | 3.550 | 1 | 0.10 | 97.97 |
| 47 | 3.600 | 1 | 0.09 | 98.06 |
| 48 | 3.606 | 1 | 0.09 | 98.15 |
| 49 | 3.612 | 1 | 0.09 | 98.25 |

| | | | | |
|---|---|---|---|---|
| 50 | 3.616 | 1 | 0.09 | 98.34 |
| 51 | 3.620 | 1 | 0.09 | 98.43 |
| 52 | 3.664 | 1 | 0.08 | 98.51 |
| 53 | 3.683 | 1 | 0.08 | 98.59 |
| 54 | 3.817 | 1 | 0.06 | 98.66 |
| 55 | 3.900 | 1 | 0.06 | 98.71 |
| 56 | 3.917 | 1 | 0.05 | 98.77 |
| 57 | 3.927 | 1 | 0.05 | 98.82 |
| 58 | 3.933 | 1 | 0.05 | 98.87 |
| 59 | 3.935 | 1 | 0.05 | 98.93 |
| 60 | 3.955 | 1 | 0.05 | 98.98 |
| 61 | 3.955 | 1 | 0.05 | 99.03 |
| 62 | 3.961 | 1 | 0.05 | 99.08 |
| 63 | 3.968 | 1 | 0.05 | 99.13 |
| 64 | 4.049 | 1 | 0.04 | 99.17 |
| 65 | 4.056 | 1 | 0.04 | 99.22 |
| 66 | 4.085 | 1 | 0.04 | 99.26 |
| 67 | 4.175 | 1 | 0.04 | 99.29 |
| 68 | 4.179 | 1 | 0.04 | 99.33 |
| 69 | 4.236 | 1 | 0.03 | 99.36 |
| 70 | 4.296 | 1 | 0.03 | 99.39 |
| 71 | 4.299 | 1 | 0.03 | 99.42 |
| 72 | 4.302 | 1 | 0.03 | 99.45 |
| 73 | 4.330 | 1 | 0.03 | 99.47 |
| 74 | 4.334 | 1 | 0.03 | 99.50 |
| 75 | 4.343 | 1 | 0.03 | 99.53 |
| 76 | 4.353 | 1 | 0.03 | 99.55 |
| 77 | 4.354 | 1 | 0.03 | 99.58 |
| 78 | 4.363 | 1 | 0.03 | 99.61 |
| 79 | 4.376 | 1 | 0.03 | 99.63 |
| 80 | 4.404 | 1 | 0.02 | 99.65 |
| 81 | 4.407 | 1 | 0.02 | 99.68 |
| 82 | 4.412 | 1 | 0.02 | 99.70 |
| 83 | 4.425 | 1 | 0.02 | 99.73 |
| 84 | 4.429 | 1 | 0.02 | 99.75 |
| 85 | 4.439 | 1 | 0.02 | 99.77 |
| 86 | 4.453 | 1 | 0.02 | 99.79 |
| 87 | 4.493 | 1 | 0.02 | 99.81 |
| 88 | 4.544 | 1 | 0.02 | 99.83 |
| 89 | 4.580 | 1 | 0.02 | 99.85 |
| 90 | 4.587 | 1 | 0.02 | 99.87 |
| 91 | 4.693 | 1 | 0.01 | 99.88 |
| 92 | 4.715 | 1 | 0.01 | 99.90 |

| | | | | |
|---|---|---|---|---|
| 93 | 4.716 | 1 | 0.01 | 99.91 |
| 94 | 4.733 | 1 | 0.01 | 99.93 |
| 95 | 4.829 | 1 | 0.01 | 99.94 |
| 96 | 4.850 | 1 | 0.01 | 99.95 |
| 97 | 5.067 | 1 | 0.01 | 99.96 |
| 98 | 5.085 | 1 | 0.01 | 99.96 |
| 99 | 5.094 | 1 | 0.01 | 99.97 |
| 100 | 5.109 | 1 | 0.01 | 99.98 |
| 101 | 5.215 | 1 | 0.01 | 99.99 |
| 102 | 5.248 | 1 | 0.01 | 99.99 |
| 103 | 5.311 | 1 | 0.01 | 100.00 |
| 104 | 5.474 | 1 | 0.00 | 100.00 |

Conformers below 3 kcal/mol: 31
Lowest energy conformer : -36.160579 Eh

Ensemble summary for Figure **4h**

| Conformer | Energy (kcal/mol) | Degen. | % total | % cumul. |
|---|---|---|---|---|
| 0 | 0.000 | 1 | 22.71 | 22.71 |
| 1 | 0.121 | 1 | 18.52 | 41.23 |
| 2 | 0.218 | 1 | 15.71 | 56.94 |
| 3 | 0.245 | 1 | 15.01 | 71.96 |
| 4 | 0.656 | 1 | 7.50 | 79.46 |
| 5 | 0.662 | 1 | 7.43 | 86.89 |
| 6 | 0.880 | 1 | 5.14 | 92.04 |
| 7 | 1.271 | 1 | 2.66 | 94.69 |
| 8 | 1.289 | 1 | 2.58 | 97.27 |
| 9 | 2.022 | 1 | 0.75 | 98.02 |
| 10 | 2.852 | 1 | 0.18 | 98.21 |
| 11 | 2.928 | 1 | 0.16 | 98.37 |
| 12 | 2.967 | 1 | 0.15 | 98.52 |
| 13 | 2.971 | 1 | 0.15 | 98.67 |
| 14 | 2.984 | 1 | 0.15 | 98.82 |
| 15 | 3.058 | 1 | 0.13 | 98.95 |
| 16 | 3.182 | 1 | 0.11 | 99.05 |
| 17 | 3.228 | 1 | 0.10 | 99.15 |
| 18 | 3.304 | 1 | 0.09 | 99.24 |
| 19 | 3.309 | 1 | 0.09 | 99.32 |
| 20 | 3.339 | 1 | 0.08 | 99.40 |
| 21 | 3.344 | 1 | 0.08 | 99.48 |
| 22 | 3.359 | 1 | 0.08 | 99.56 |
| 23 | 3.398 | 1 | 0.07 | 99.64 |

| | | | | |
|---|---|---|---|---|
| 24 | 3.520 | 1 | 0.06 | 99.70 |
| 25 | 3.710 | 1 | 0.04 | 99.74 |
| 26 | 3.739 | 1 | 0.04 | 99.78 |
| 27 | 3.750 | 1 | 0.04 | 99.82 |
| 28 | 3.750 | 1 | 0.04 | 99.86 |
| 29 | 3.969 | 1 | 0.03 | 99.89 |
| 30 | 4.286 | 1 | 0.02 | 99.91 |
| 31 | 4.449 | 1 | 0.01 | 99.92 |
| 32 | 4.679 | 1 | 0.01 | 99.93 |
| 33 | 4.736 | 1 | 0.01 | 99.93 |
| 34 | 4.783 | 1 | 0.01 | 99.94 |
| 35 | 4.855 | 1 | 0.01 | 99.95 |
| 36 | 5.021 | 1 | 0.00 | 99.95 |
| 37 | 5.050 | 1 | 0.00 | 99.96 |
| 38 | 5.053 | 1 | 0.00 | 99.96 |
| 39 | 5.068 | 1 | 0.00 | 99.97 |
| 40 | 5.099 | 1 | 0.00 | 99.97 |
| 41 | 5.101 | 1 | 0.00 | 99.97 |
| 42 | 5.133 | 1 | 0.00 | 99.98 |
| 43 | 5.168 | 1 | 0.00 | 99.98 |
| 44 | 5.303 | 1 | 0.00 | 99.98 |
| 45 | 5.309 | 1 | 0.00 | 99.99 |
| 46 | 5.319 | 1 | 0.00 | 99.99 |
| 47 | 5.358 | 1 | 0.00 | 99.99 |
| 48 | 5.415 | 1 | 0.00 | 100.00 |
| 49 | 5.470 | 1 | 0.00 | 100.00 |
| 50 | 5.593 | 1 | 0.00 | 100.00 |
| 51 | 5.971 | 1 | 0.00 | 100.00 |

Conformers below 3 kcal/mol: 15
Lowest energy conformer : -36.160328 Eh

#### 5. Copies of NMR spectra

$^1$H and $^{13}$C spectra of **1.4a**

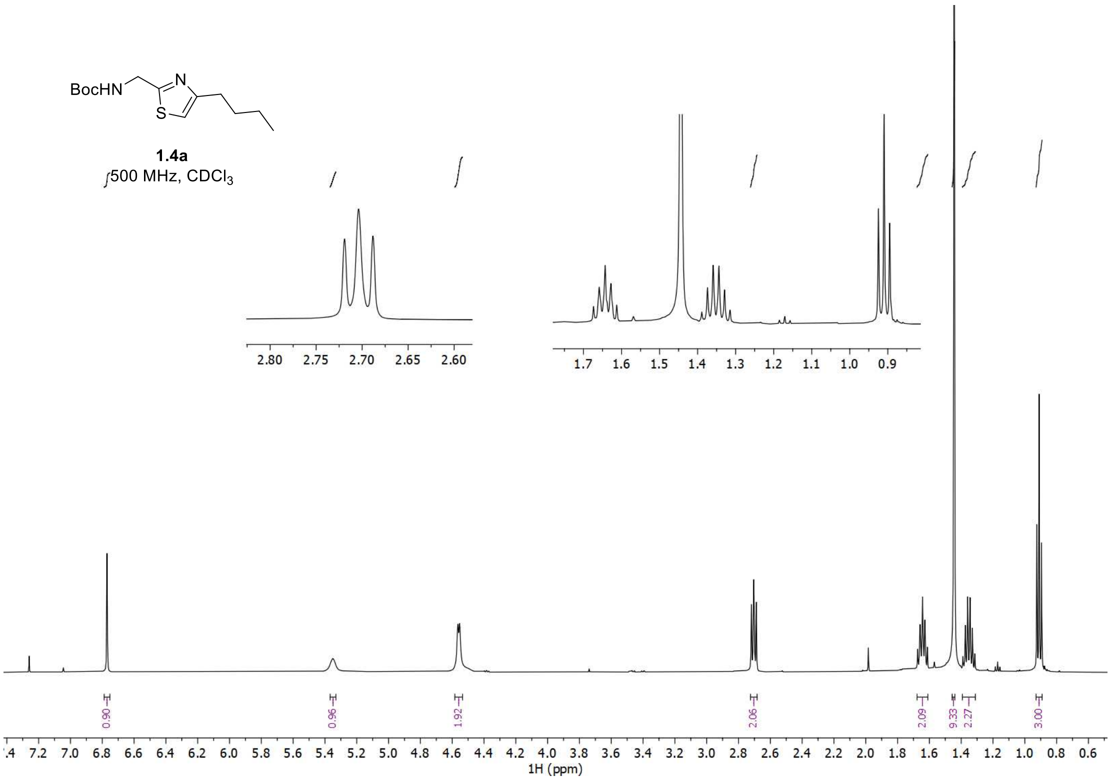


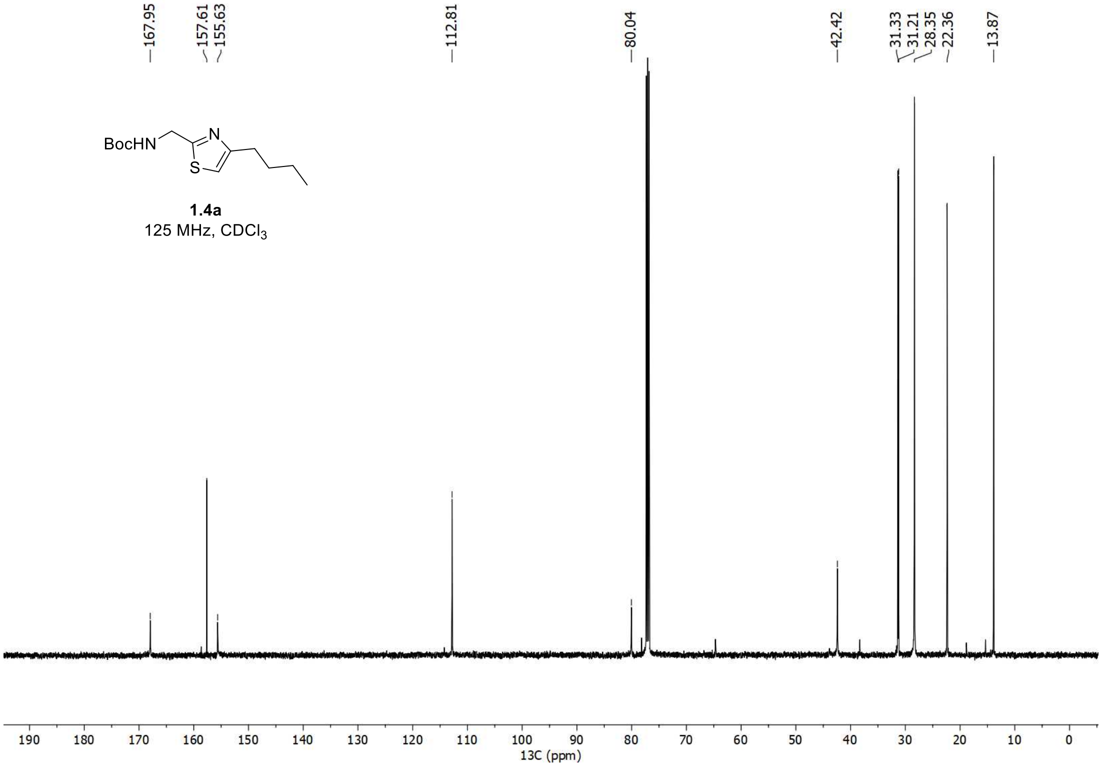

#### $^{1}H$ and $^{13}C$ spectra of **1a**

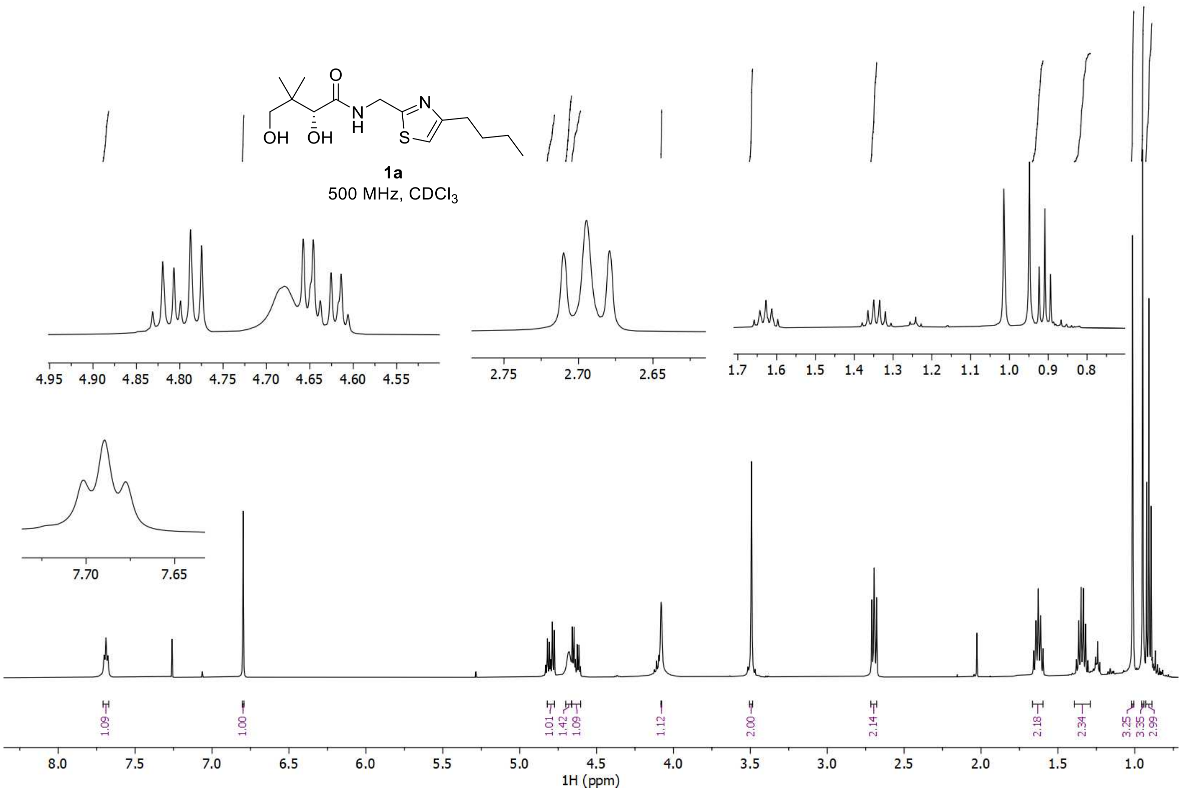


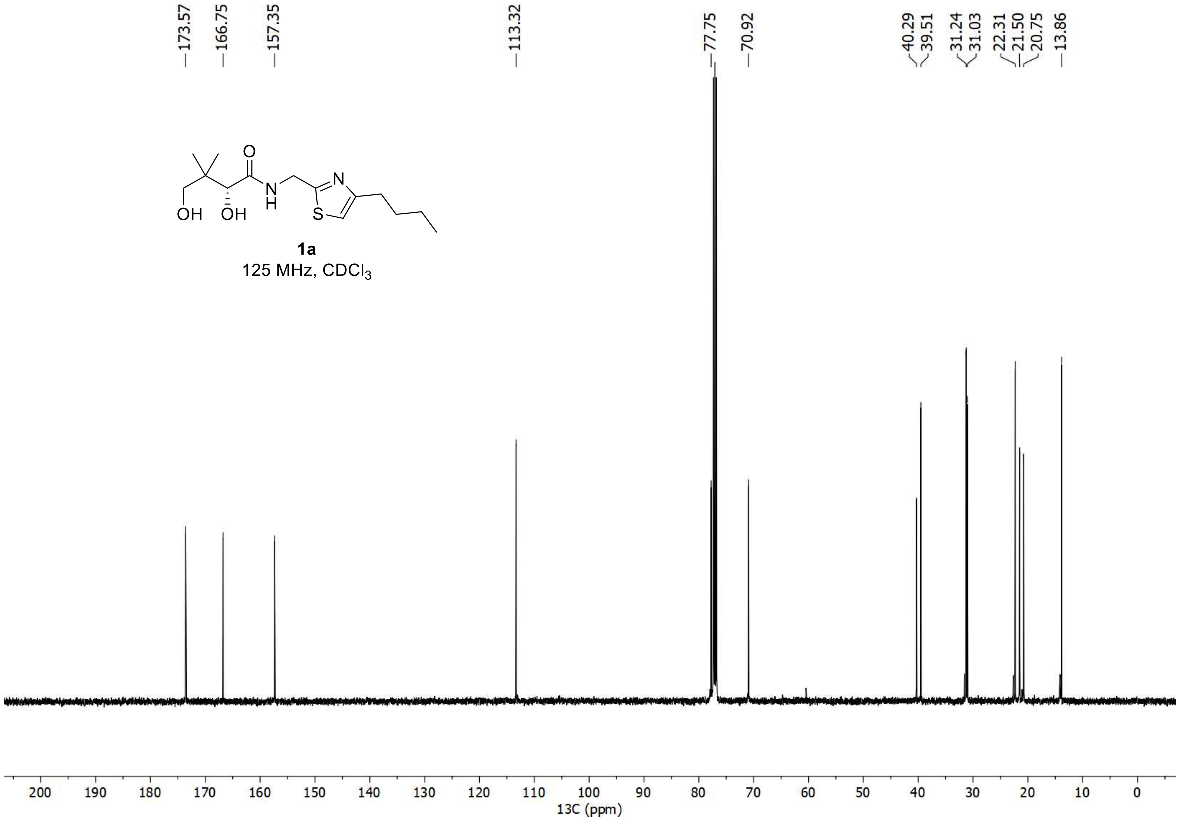

<u><sup></sup></u>$^{1}$H and $^{13}$C spectra of **1.4b**

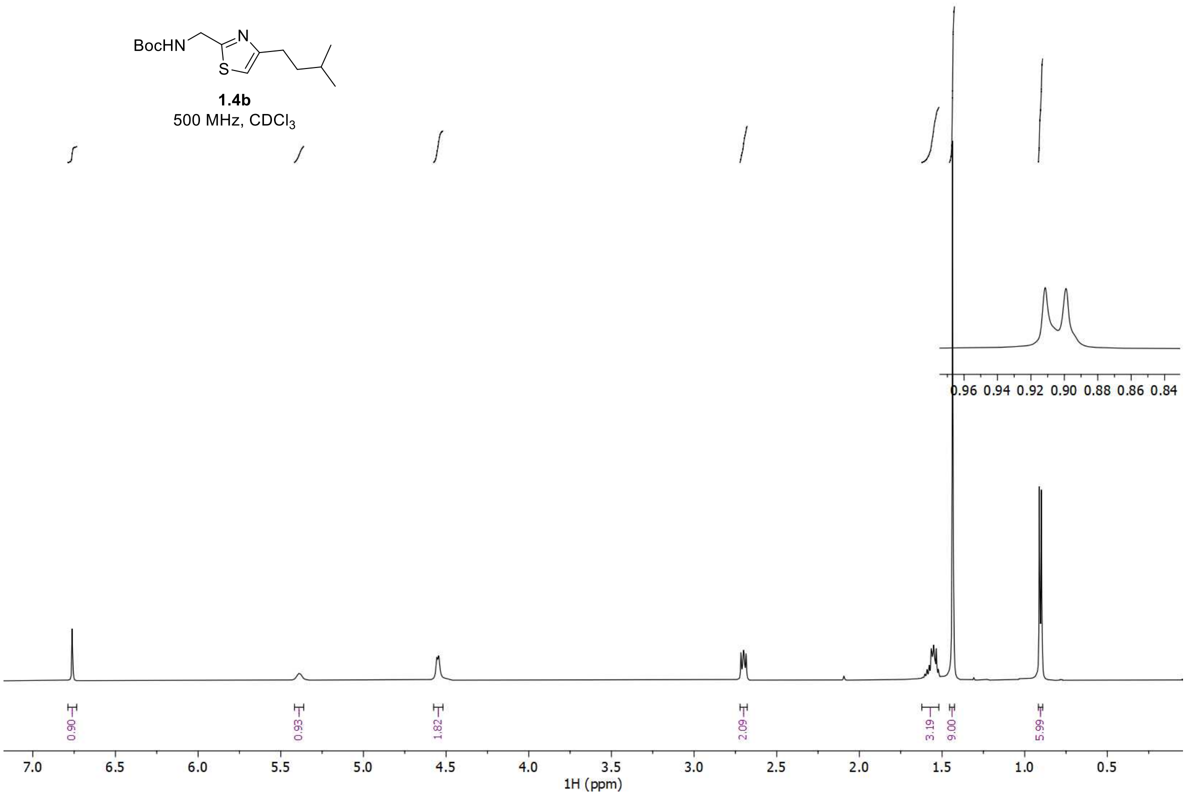


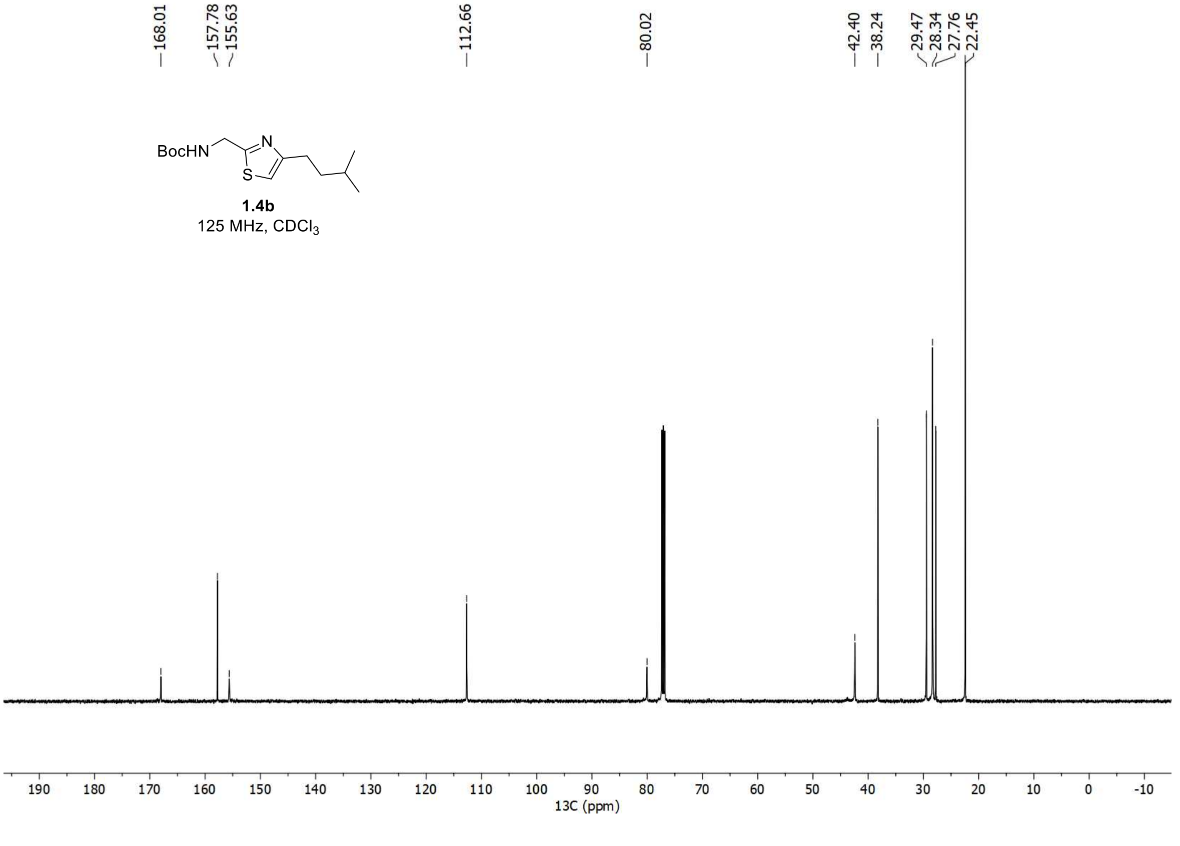

<u>$^{1}H$ and $^{13}C$ spectra of **1b**</u>

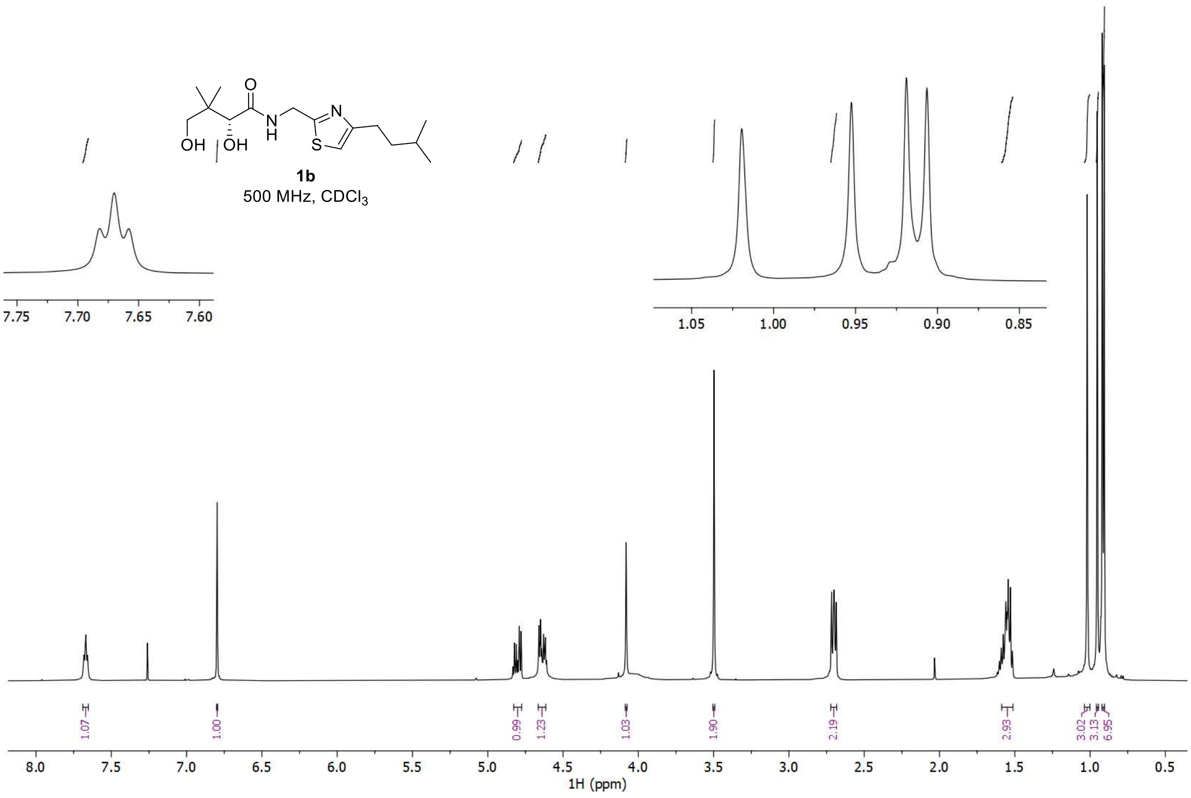


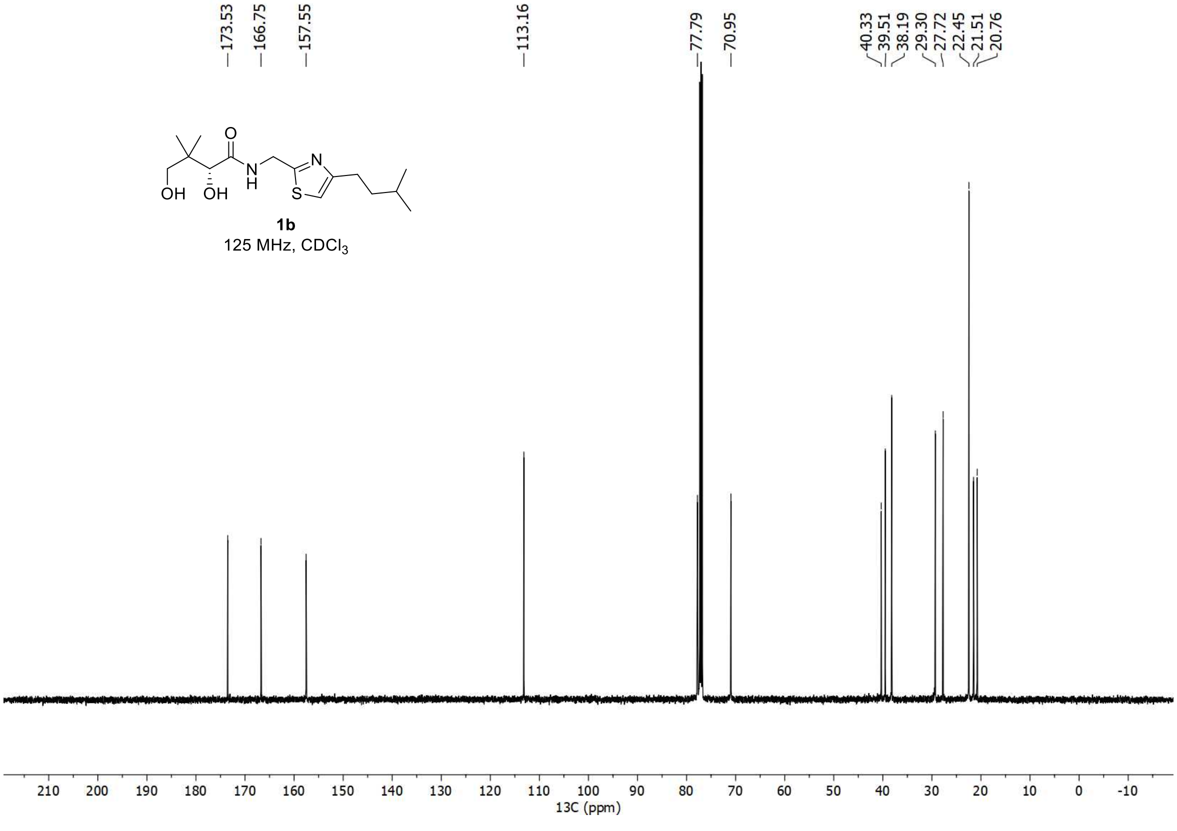

[1]H and [13]C spectra of **1c**

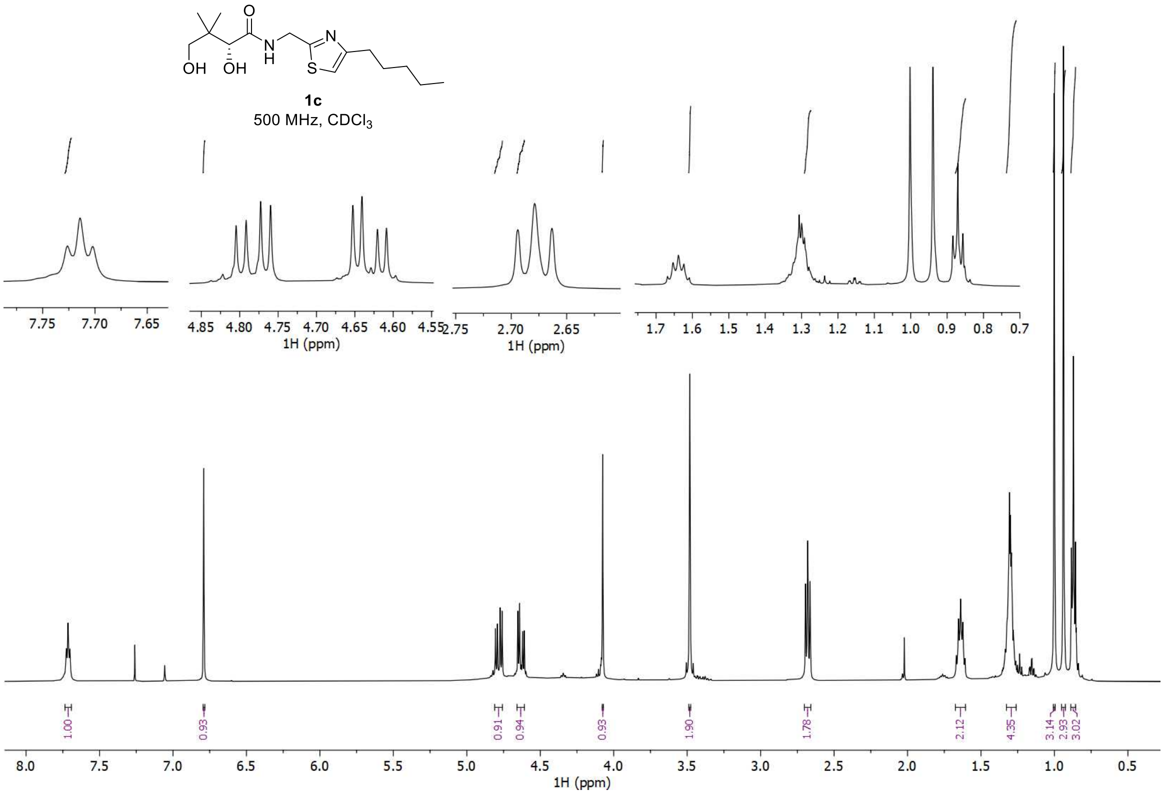


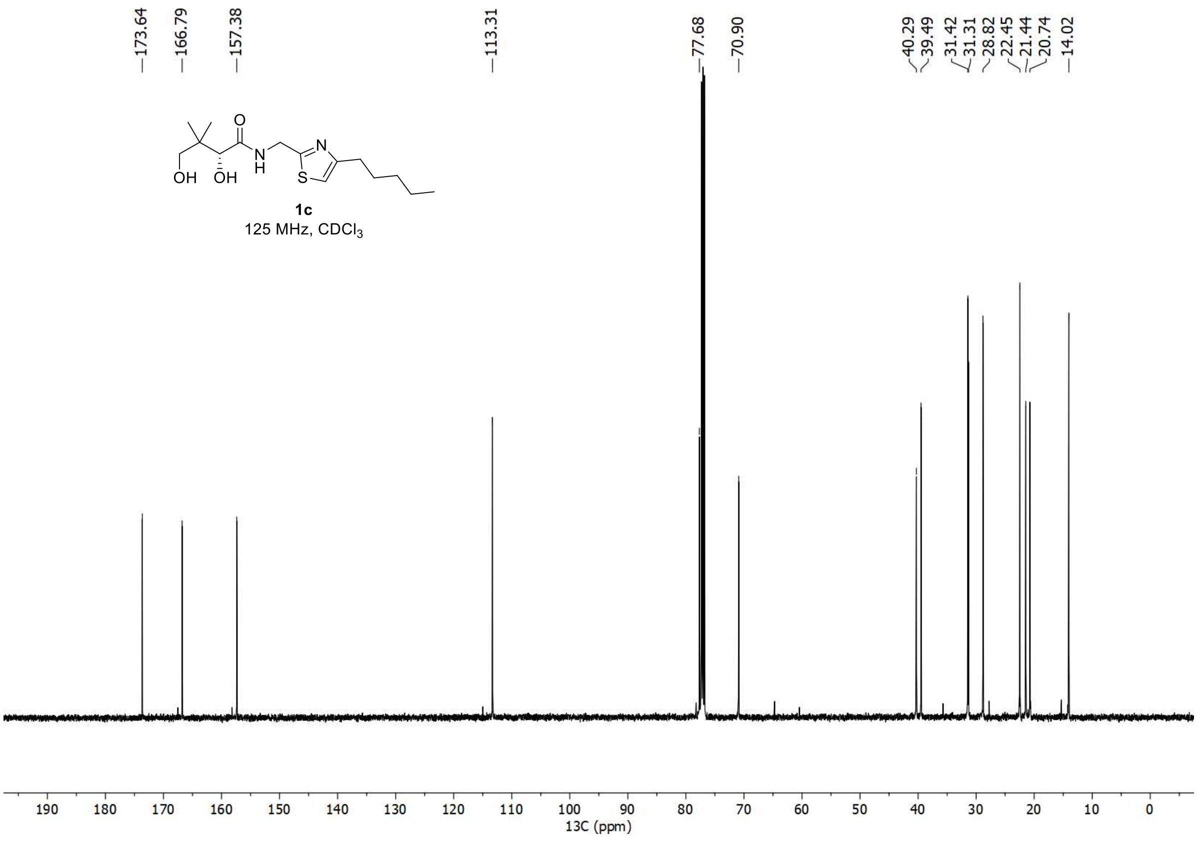

<u>$^{1}$H and $^{13}$C spectra of **1.4d**</u>

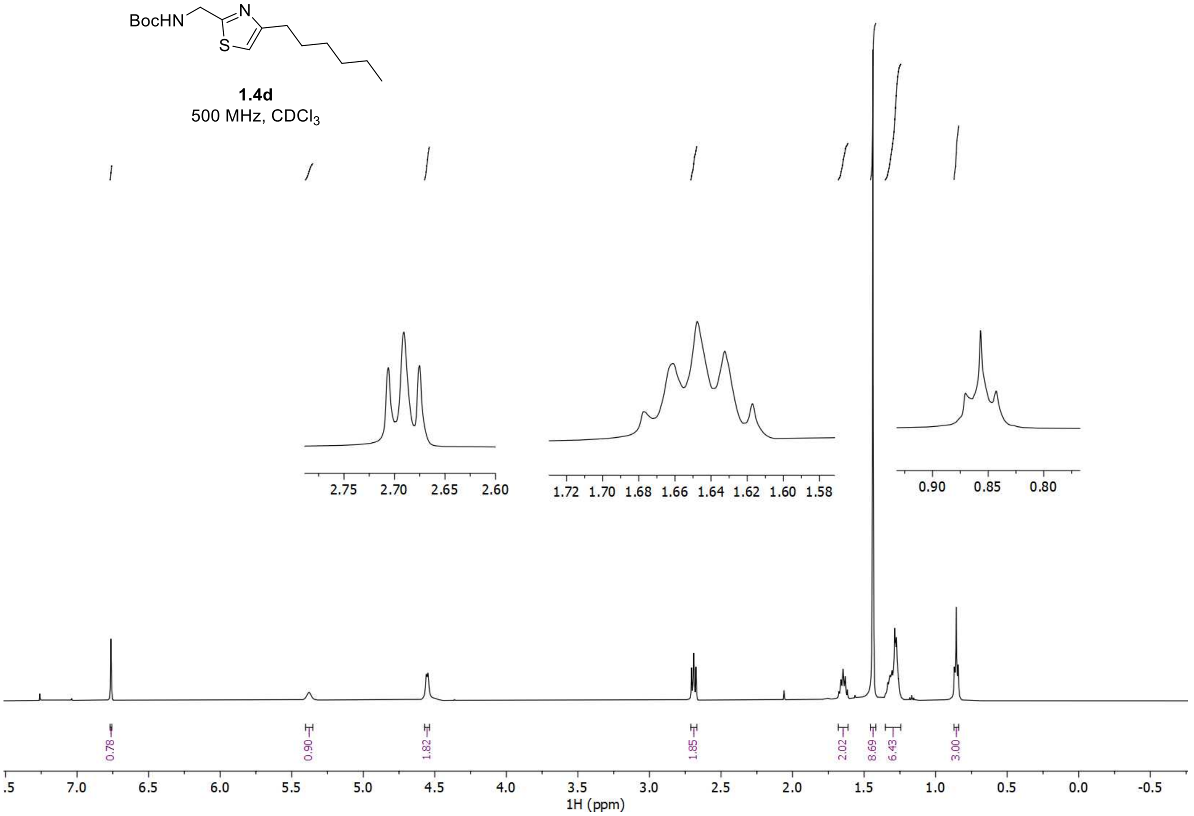


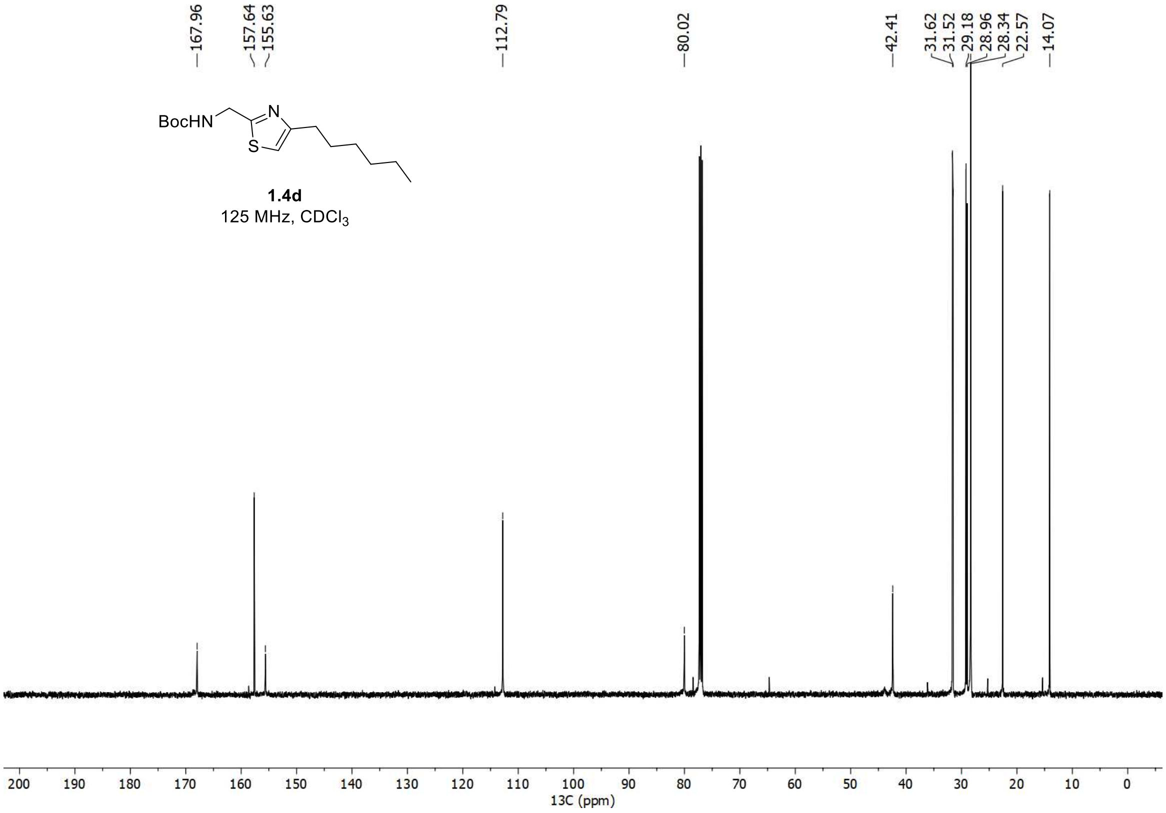

#### $^{1}$H and $^{13}$C spectra of **1d**

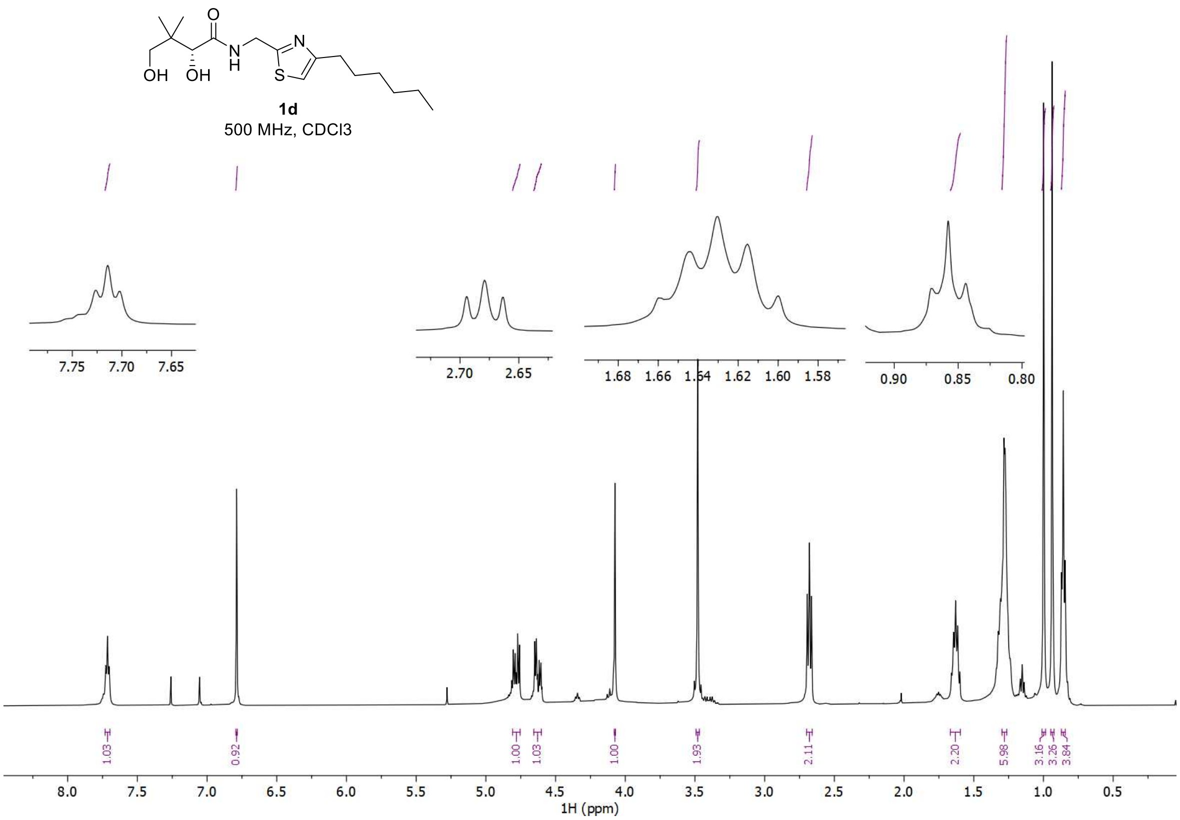


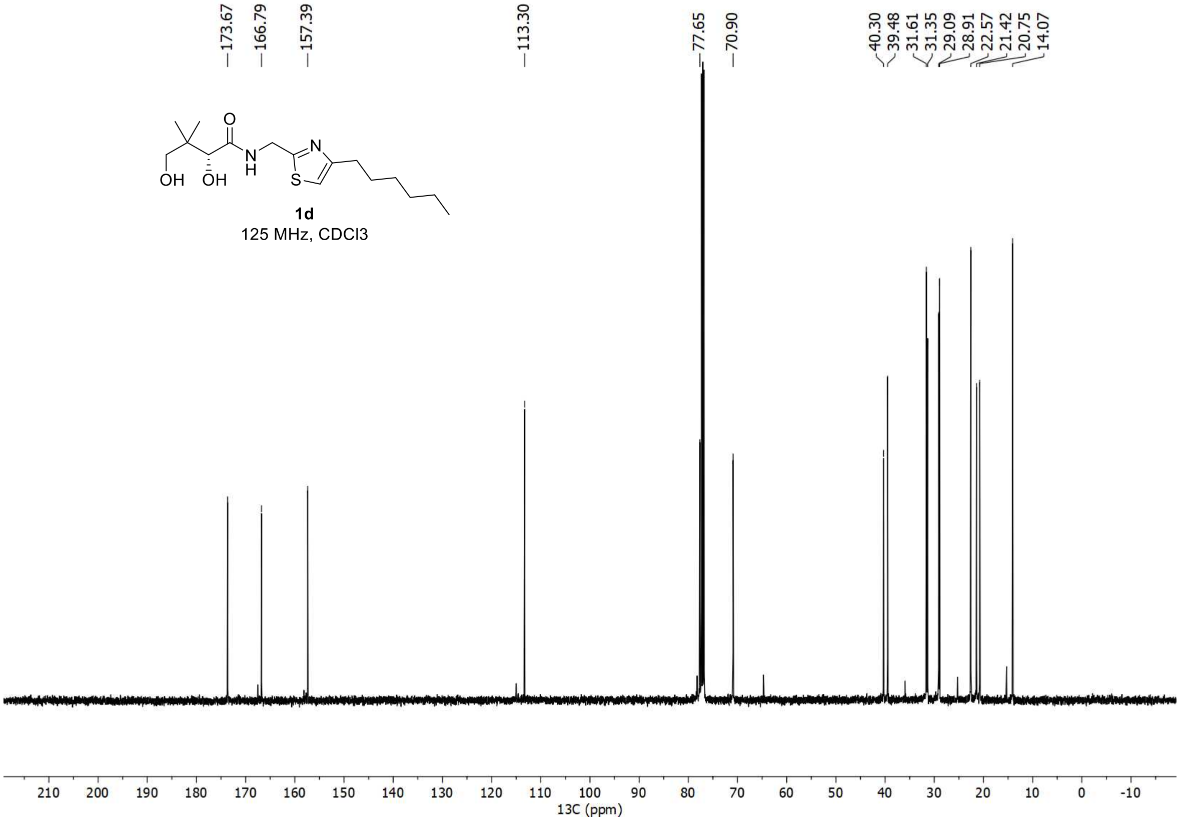

$^{1}H$ and $^{13}C$ spectra of **1.4e**

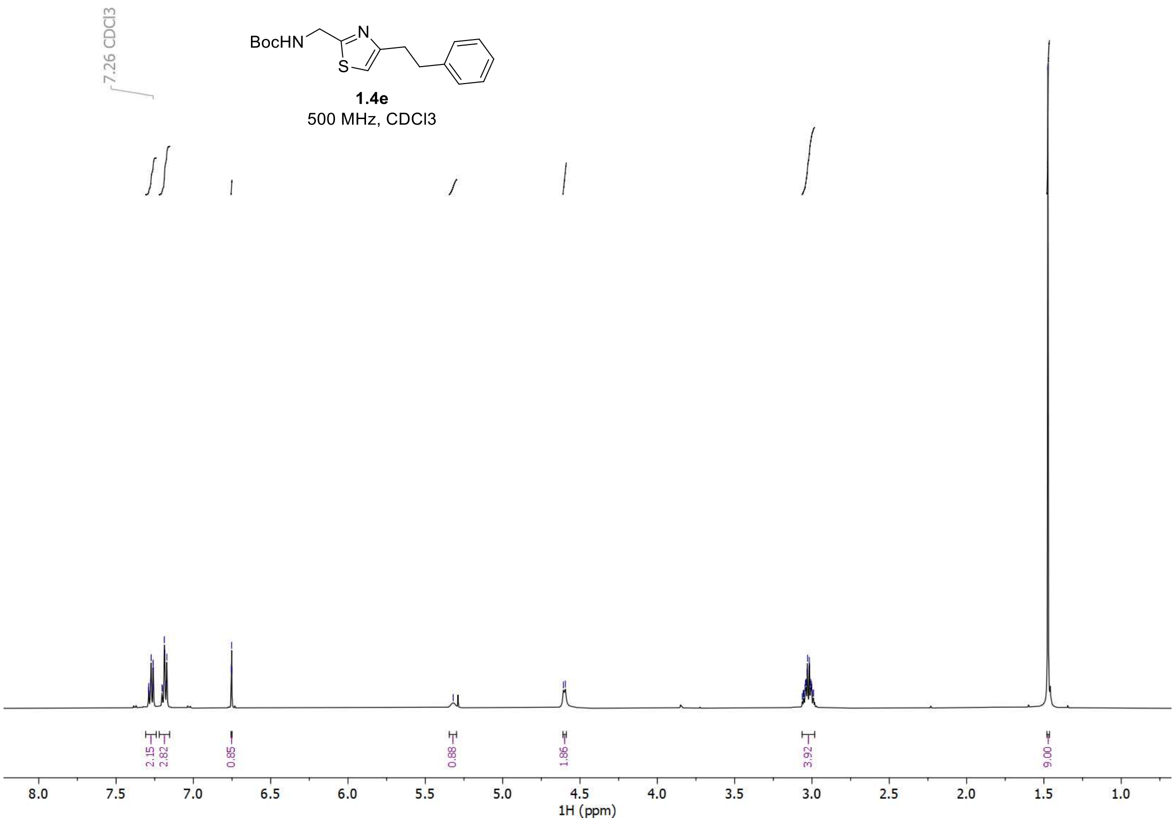


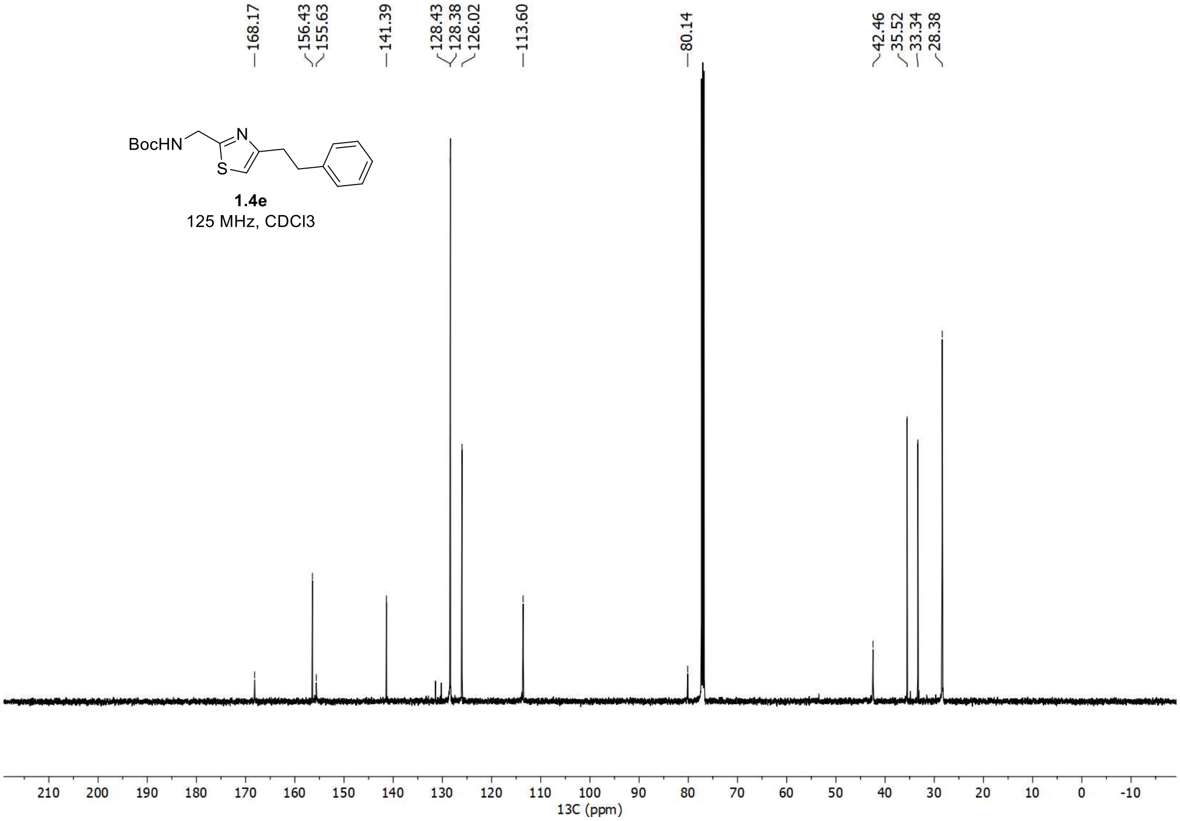

<u>$^{1}$H and $^{13}$C spectra of **1e**</u>

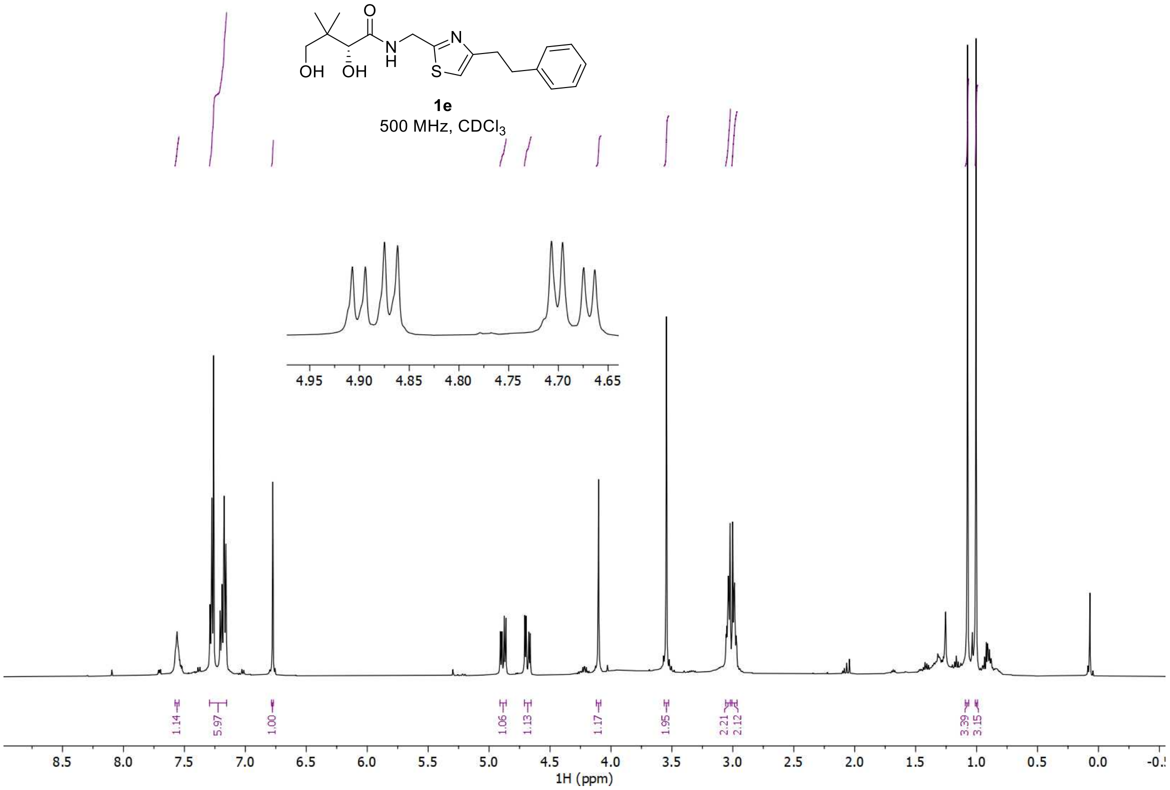


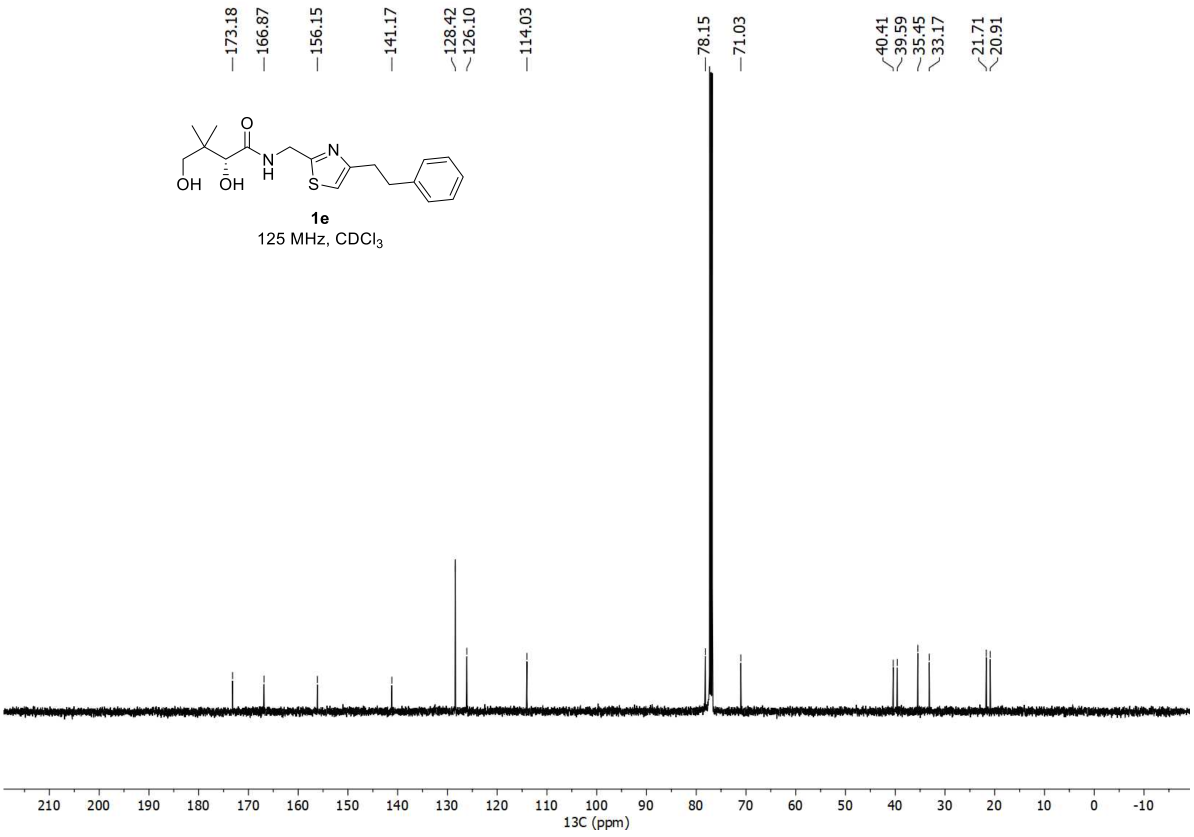

$^{1}H$ and $^{13}C$ spectra of **2.5a**

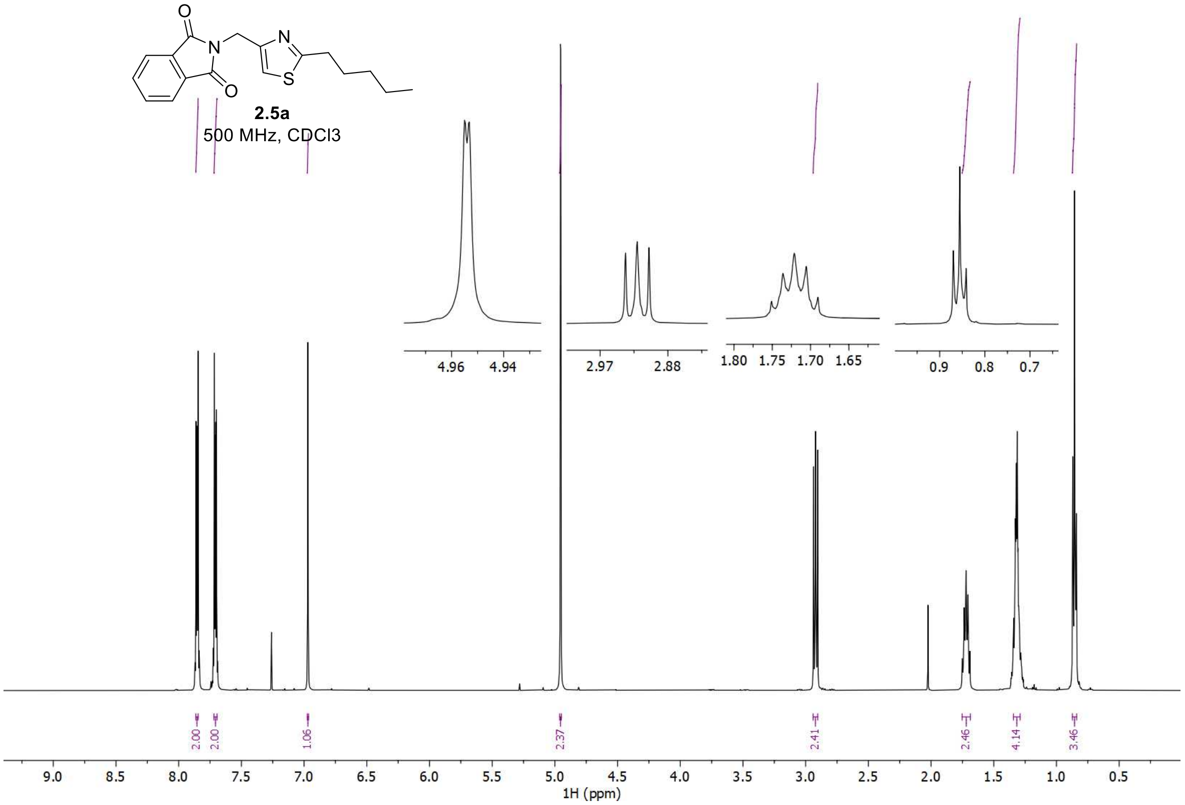


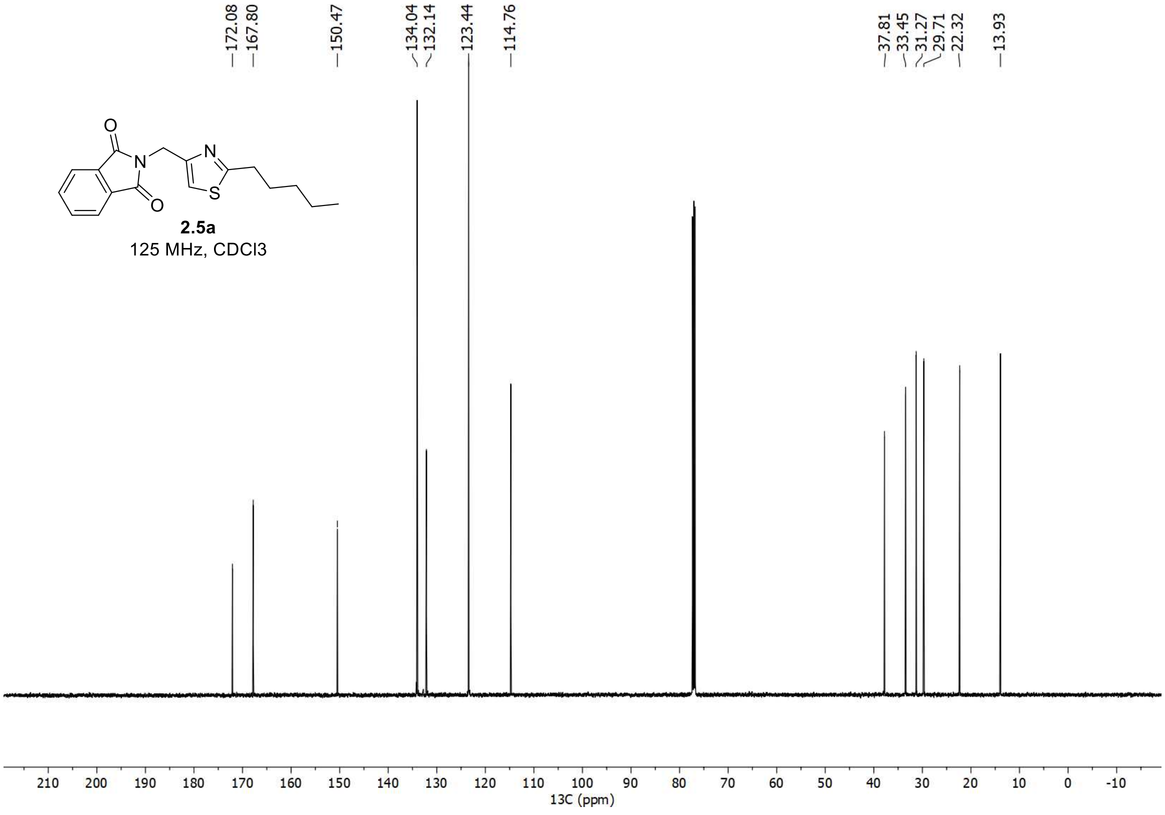

<u>[1]H and [13]C spectra of **2.6a**</u>

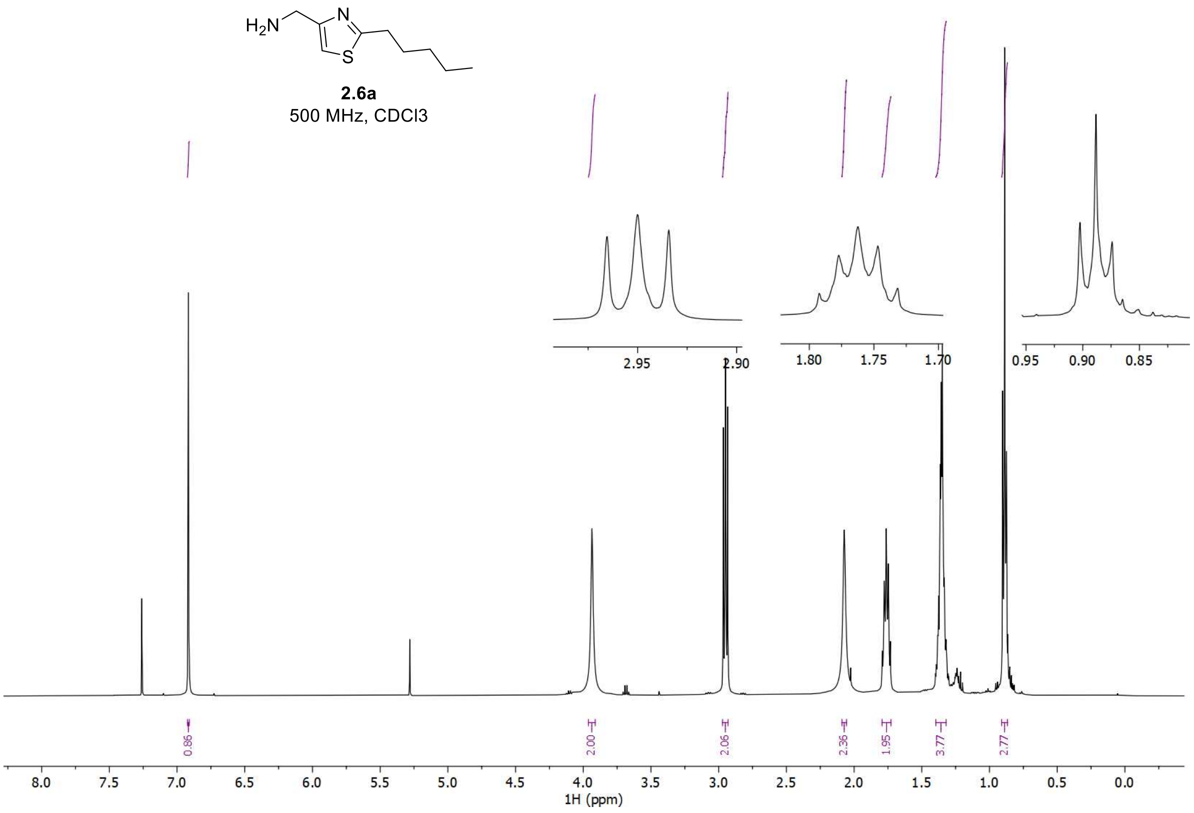


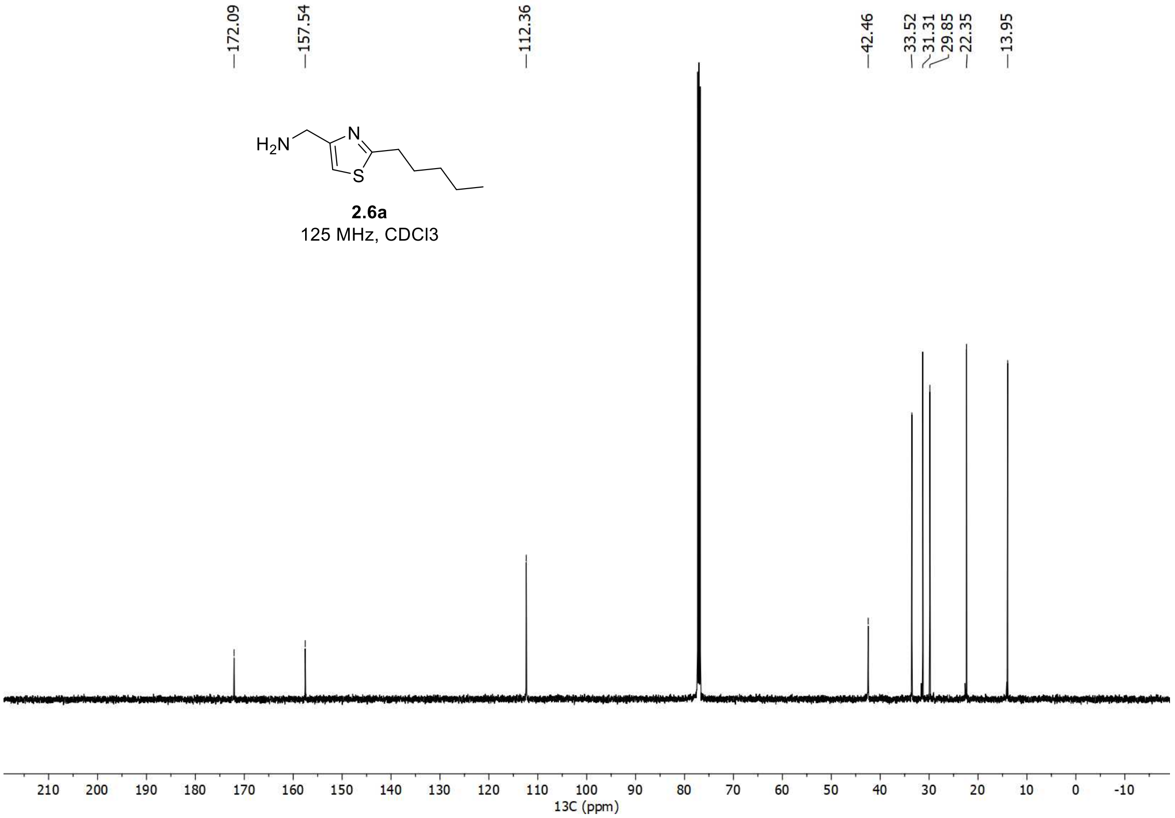

<u>$^{1}$H and $^{13}$C spectra of **2a**</u>

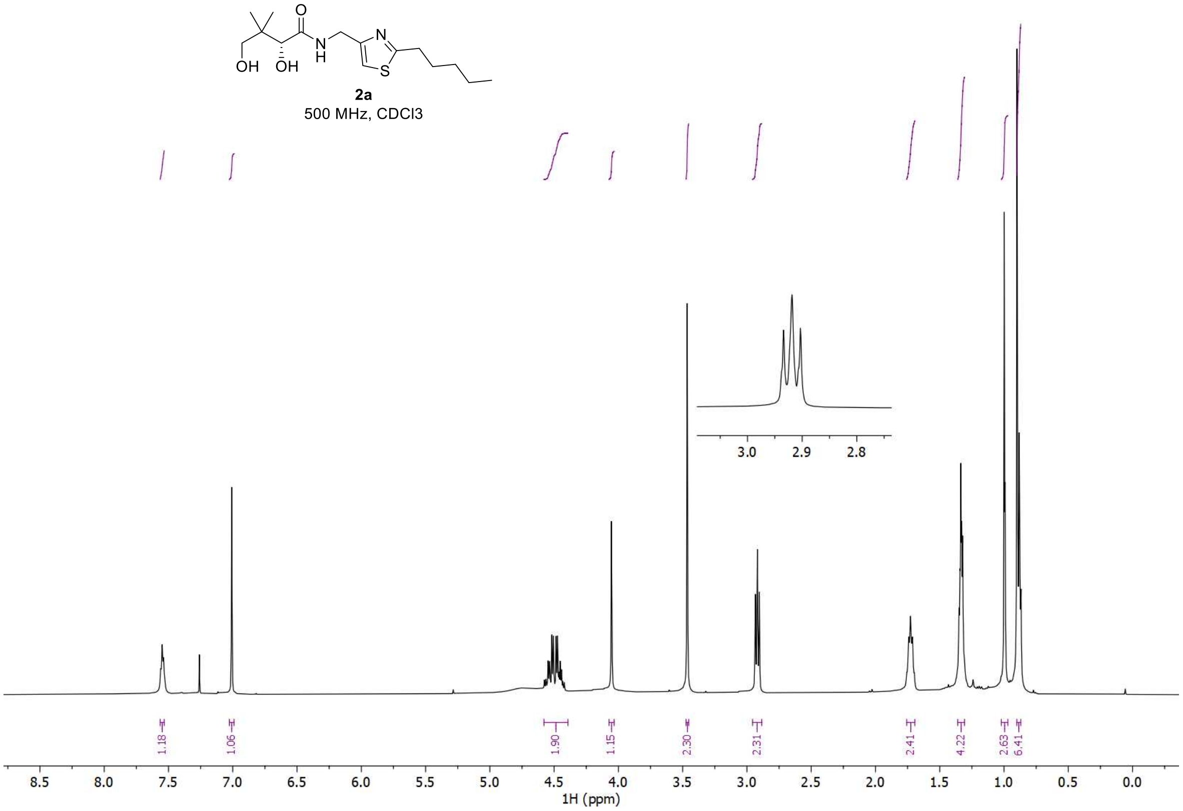


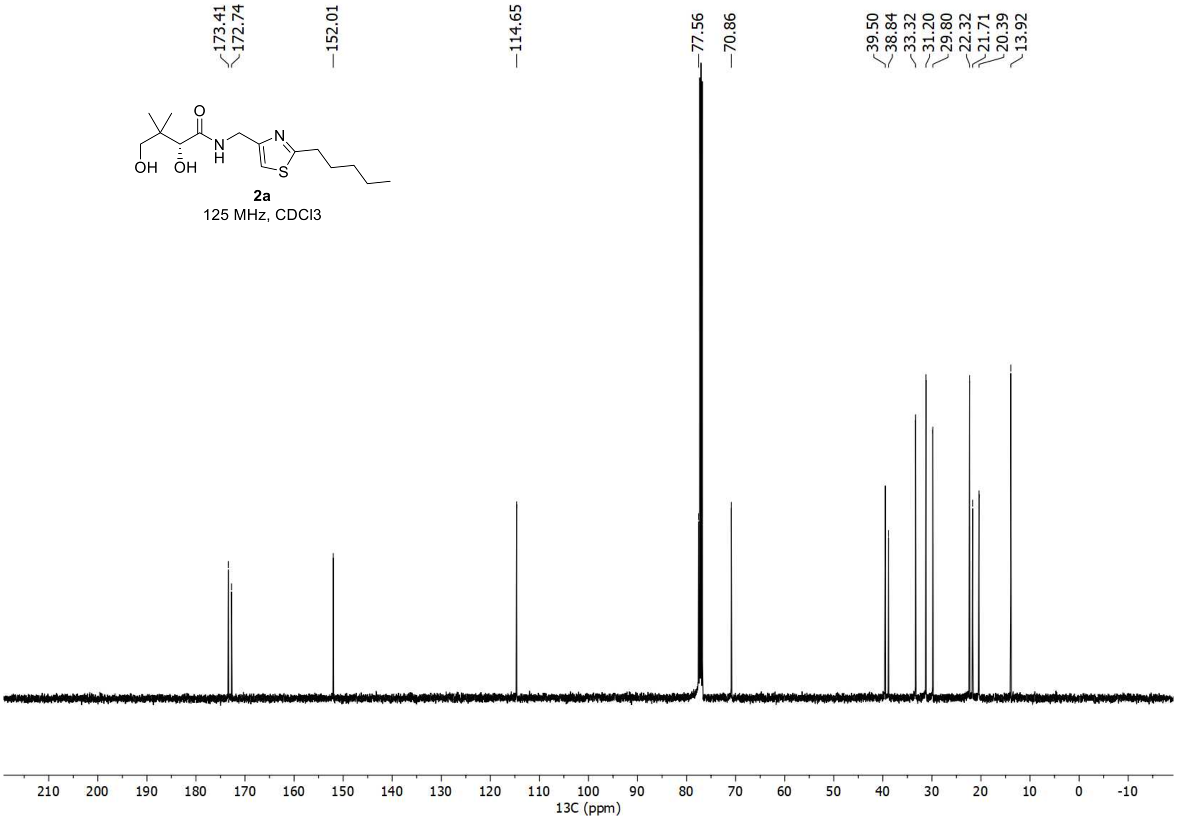

### $^1$H and $^{13}$C spectra of **2.5b**

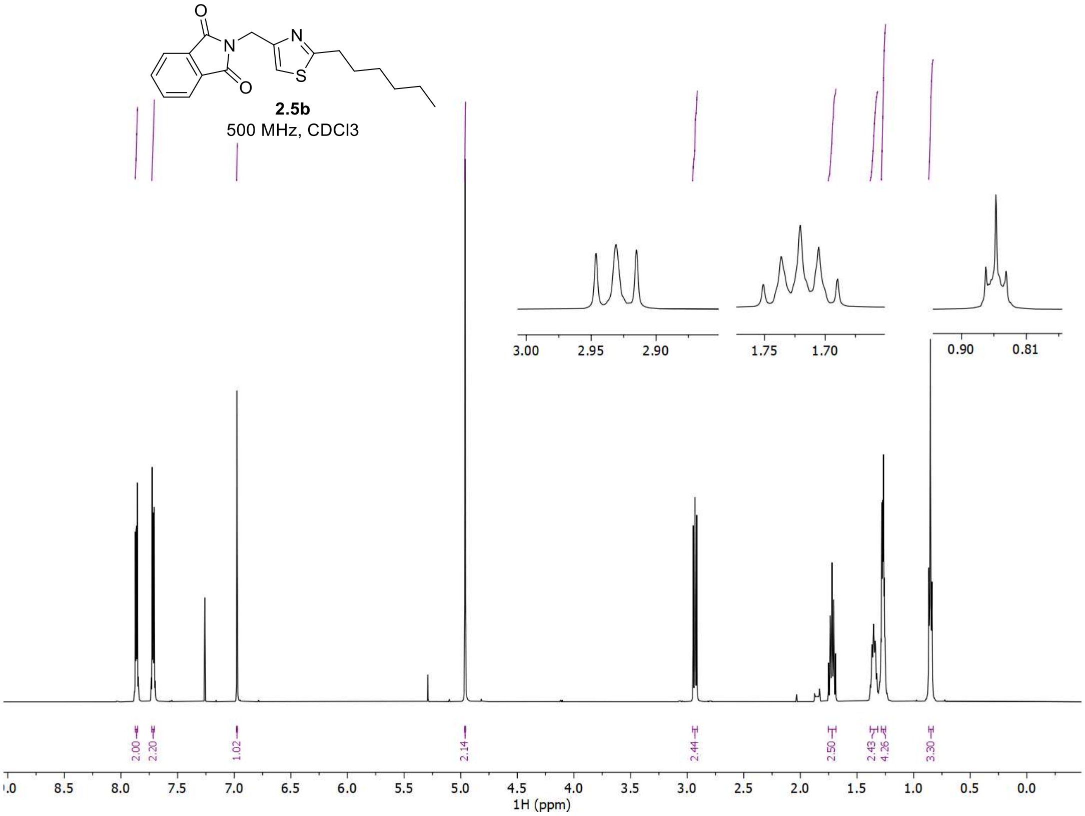


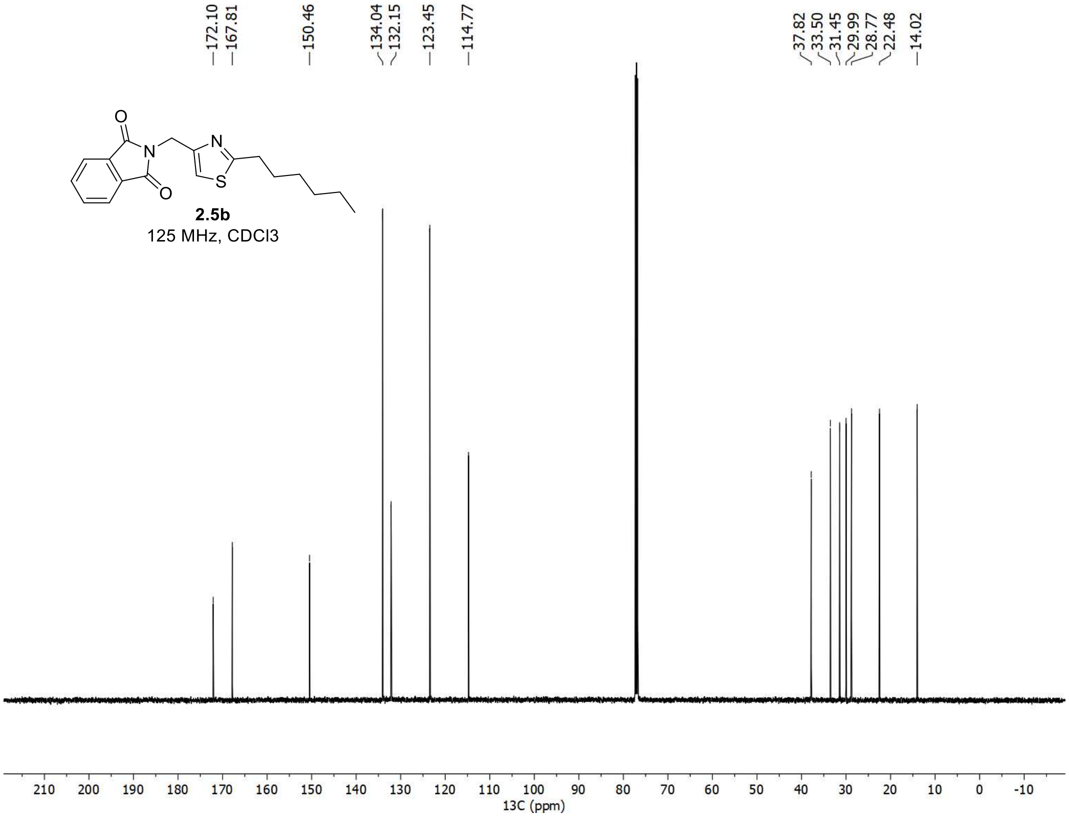

$^{1}H$ and $^{13}C$ spectra of **2.6b**

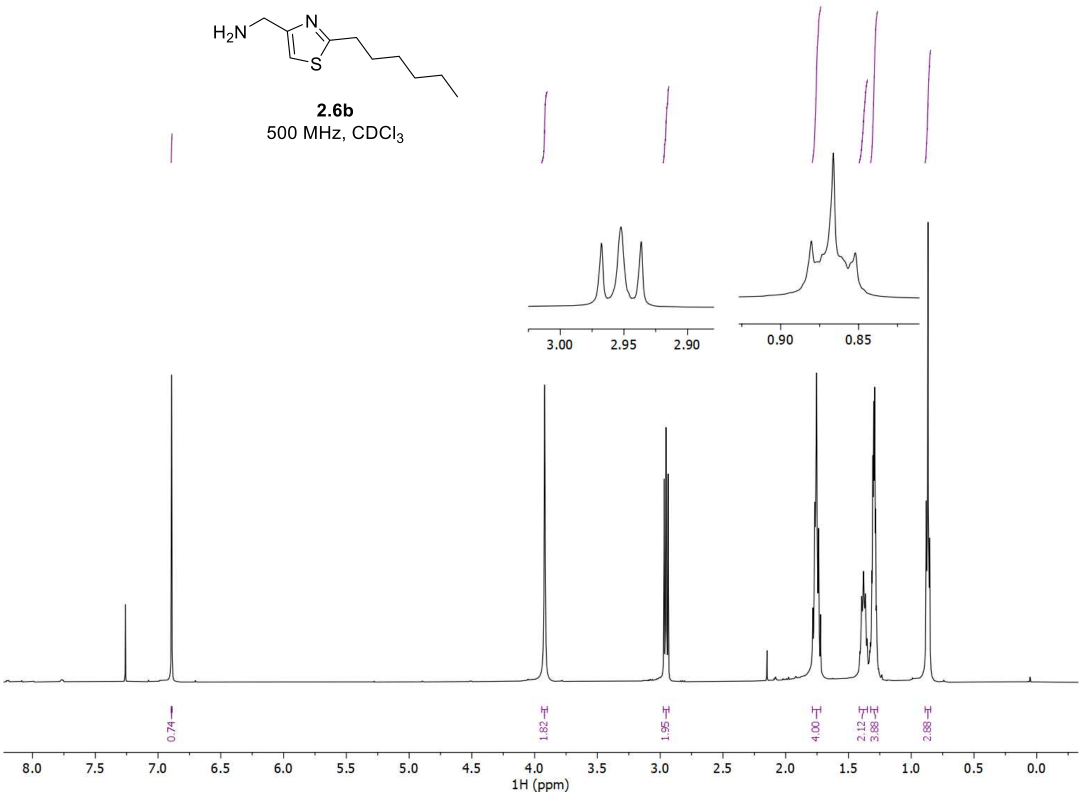


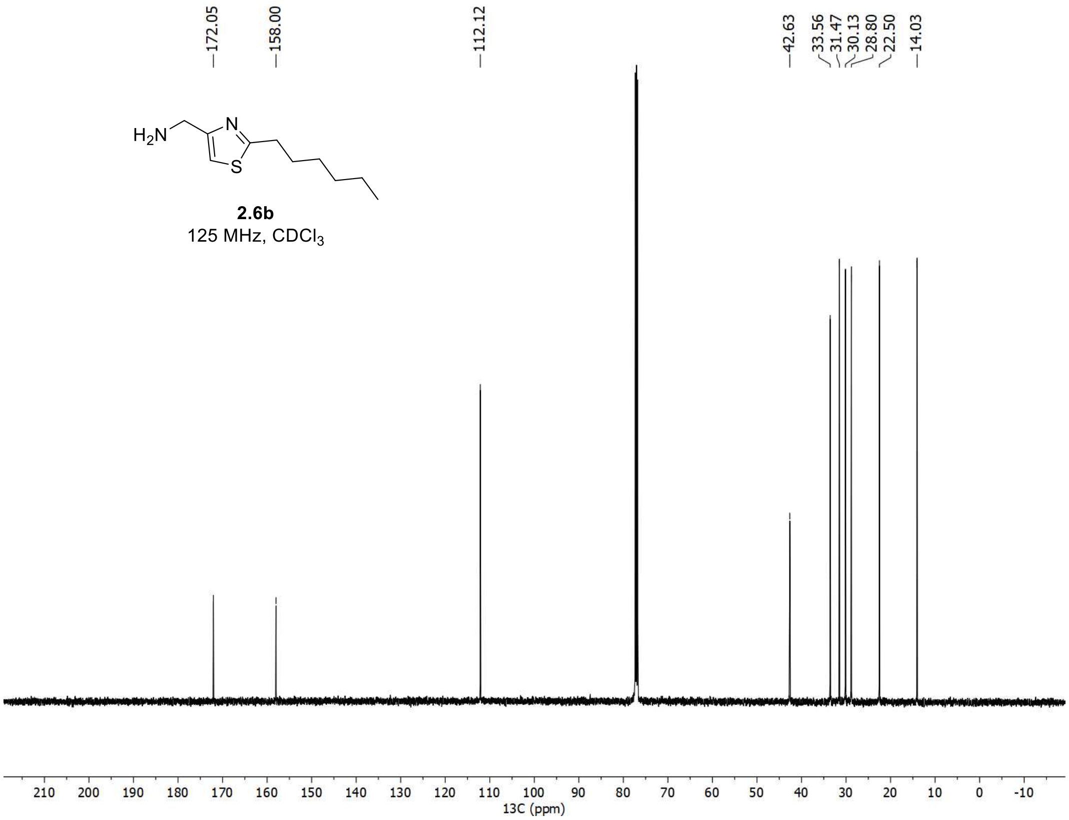

[1]H and [13]C spectra of **2b**

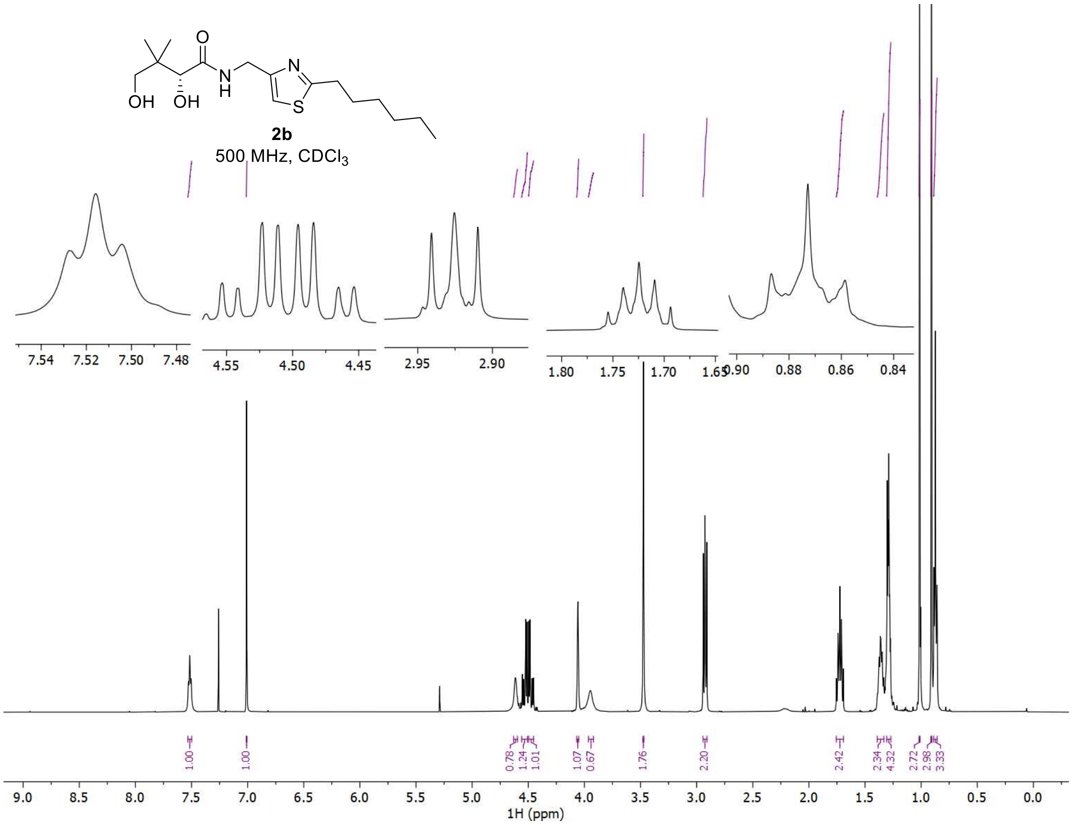


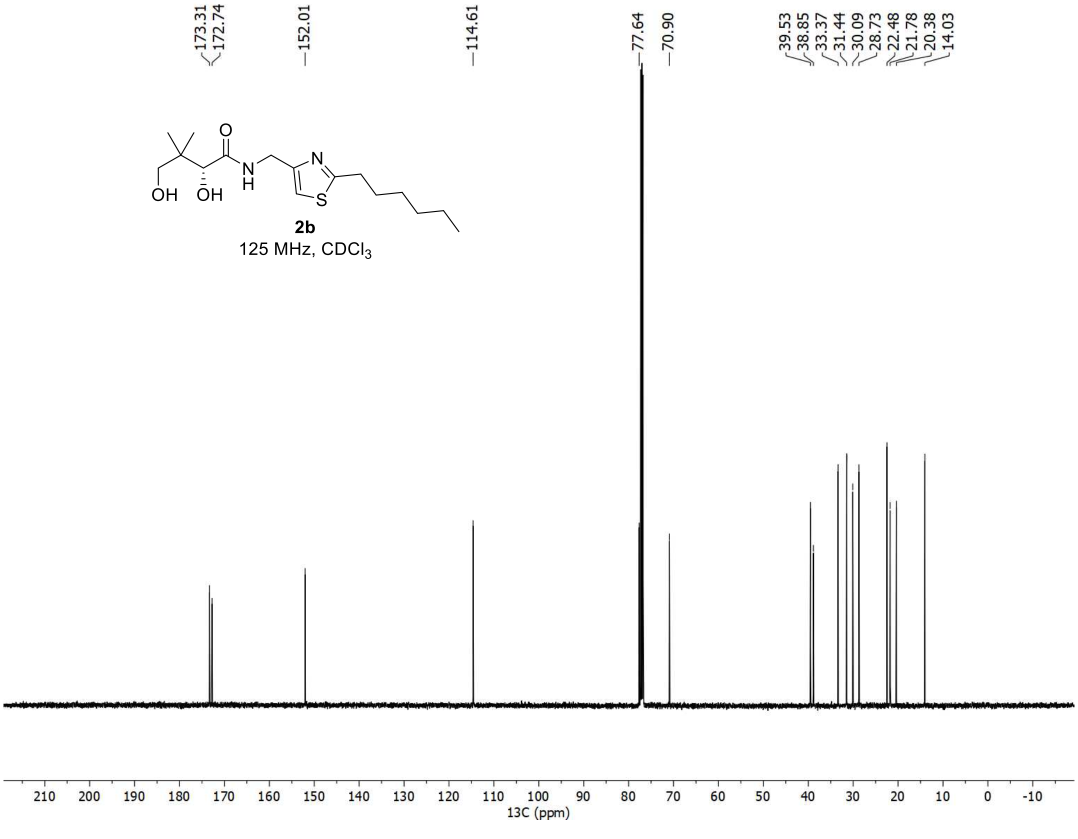

<u>$^{1}$H and $^{13}$C spectra of **2.5c**</u>

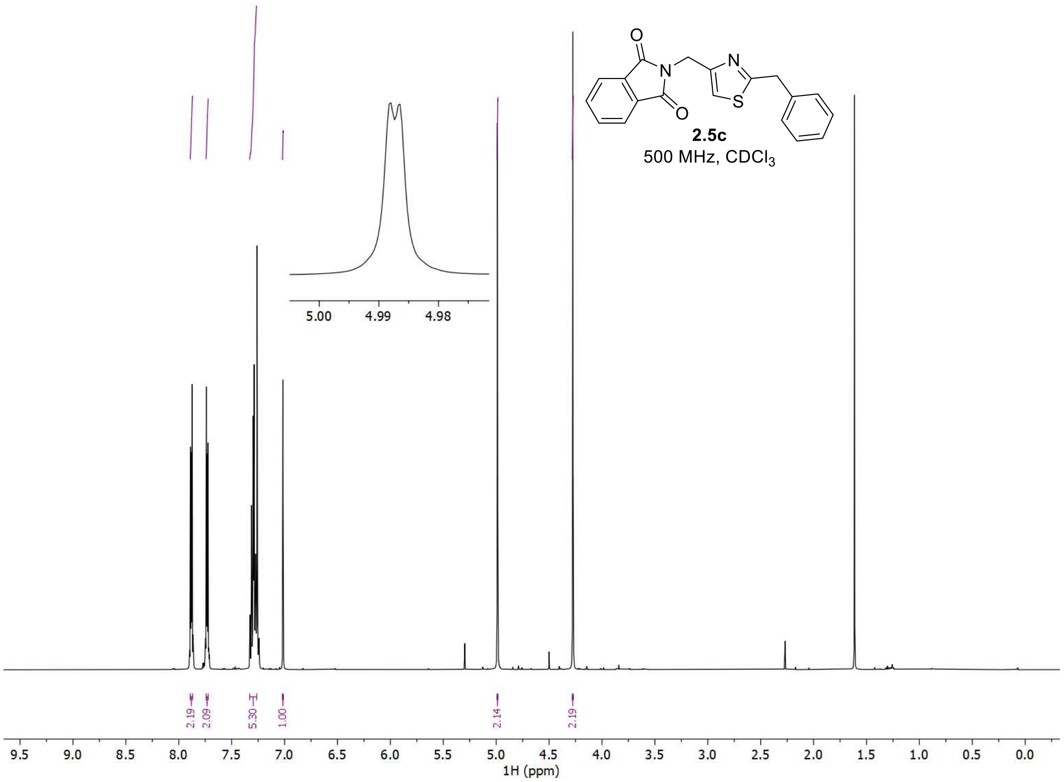


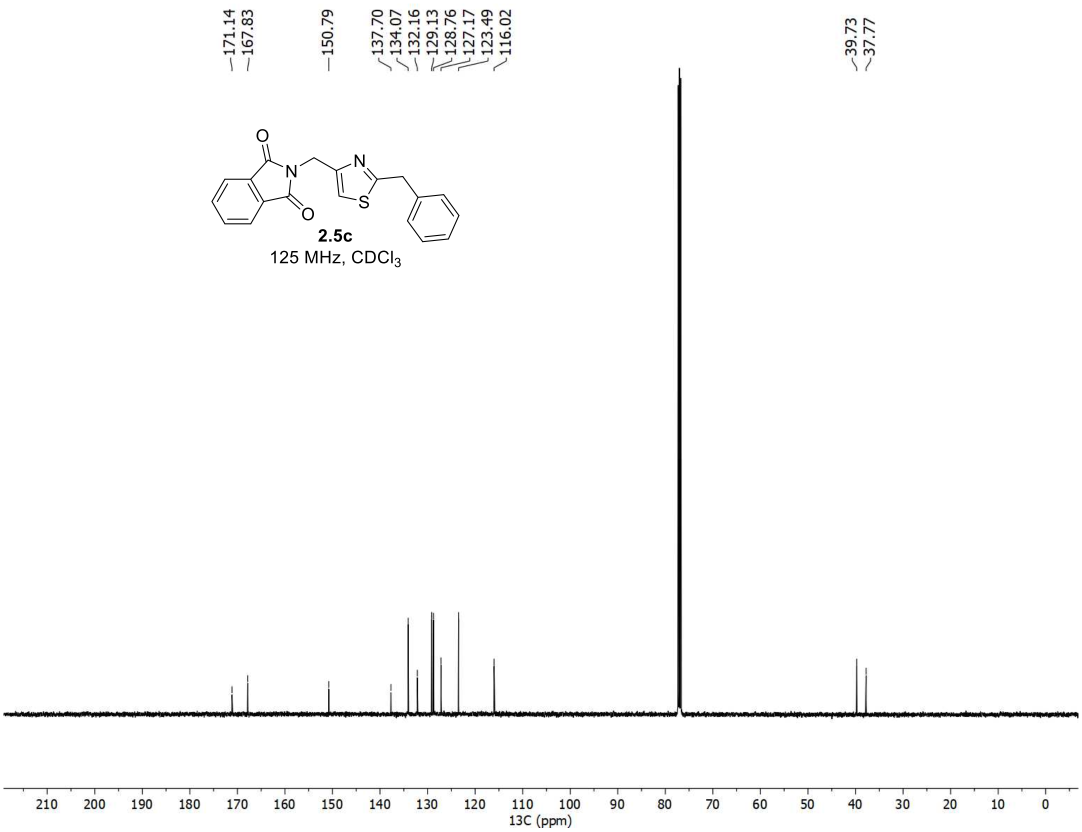

$^{1}H$ and $^{13}C$ spectra of **2.6c**

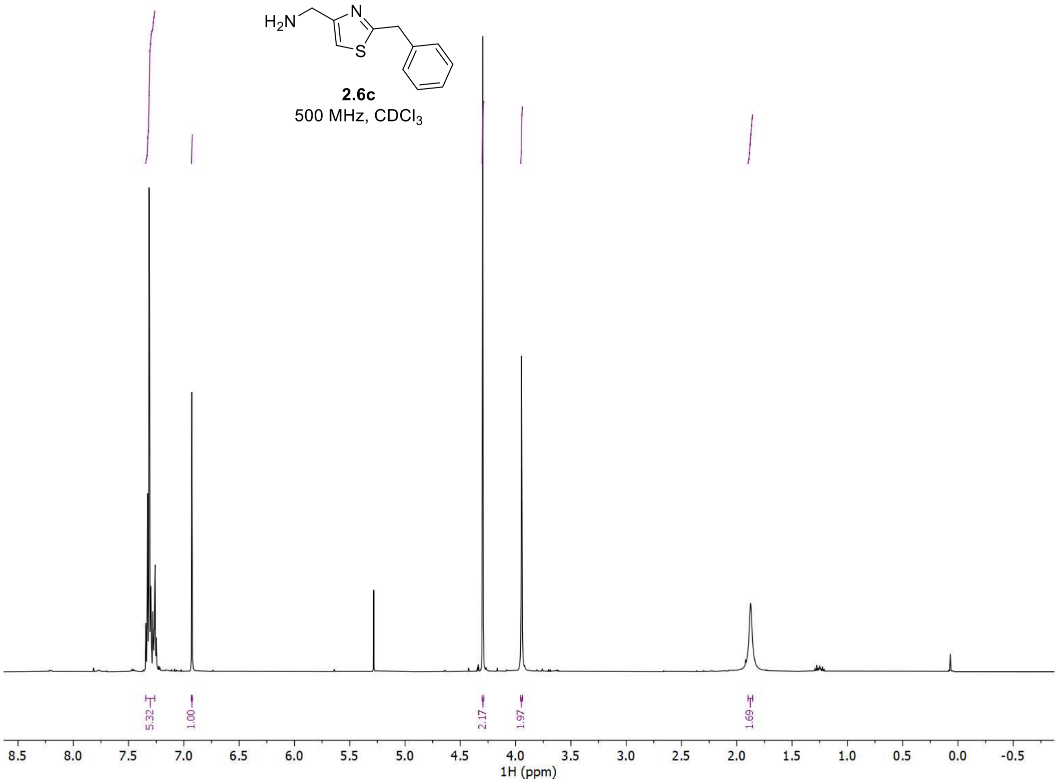


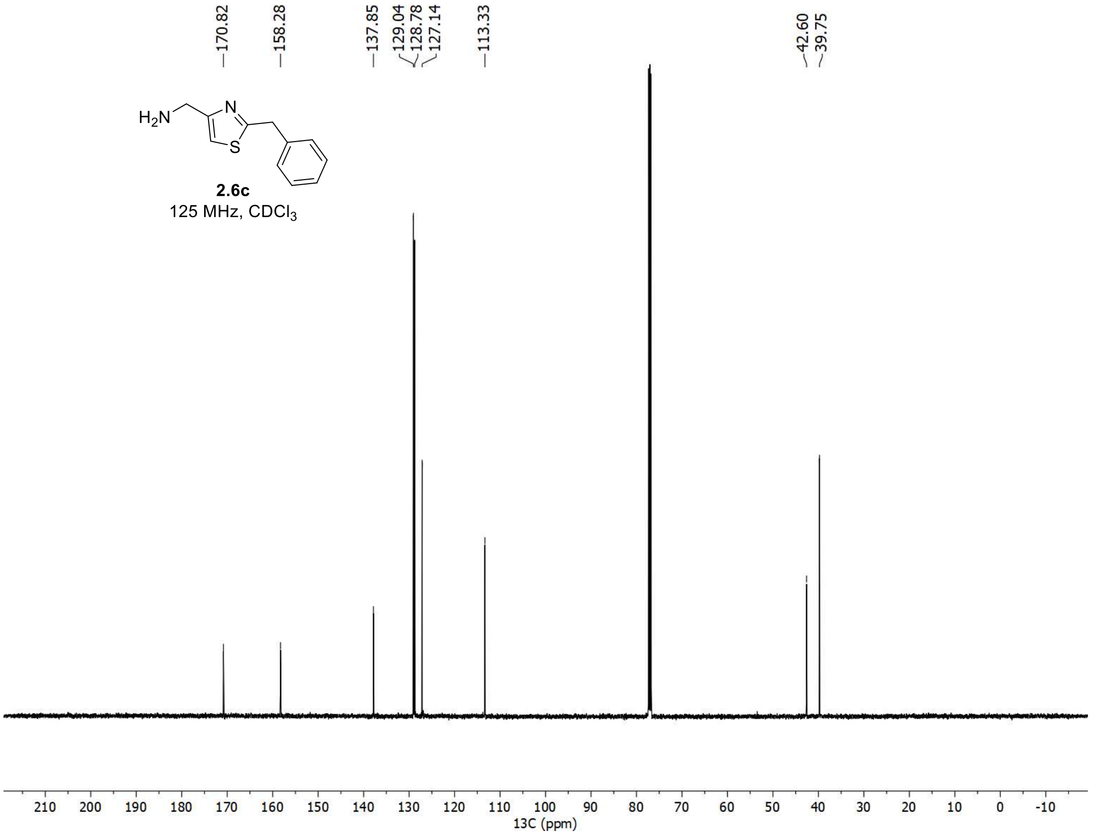

<u>$^{1}$H and $^{13}$C spectra of **2c**</u>

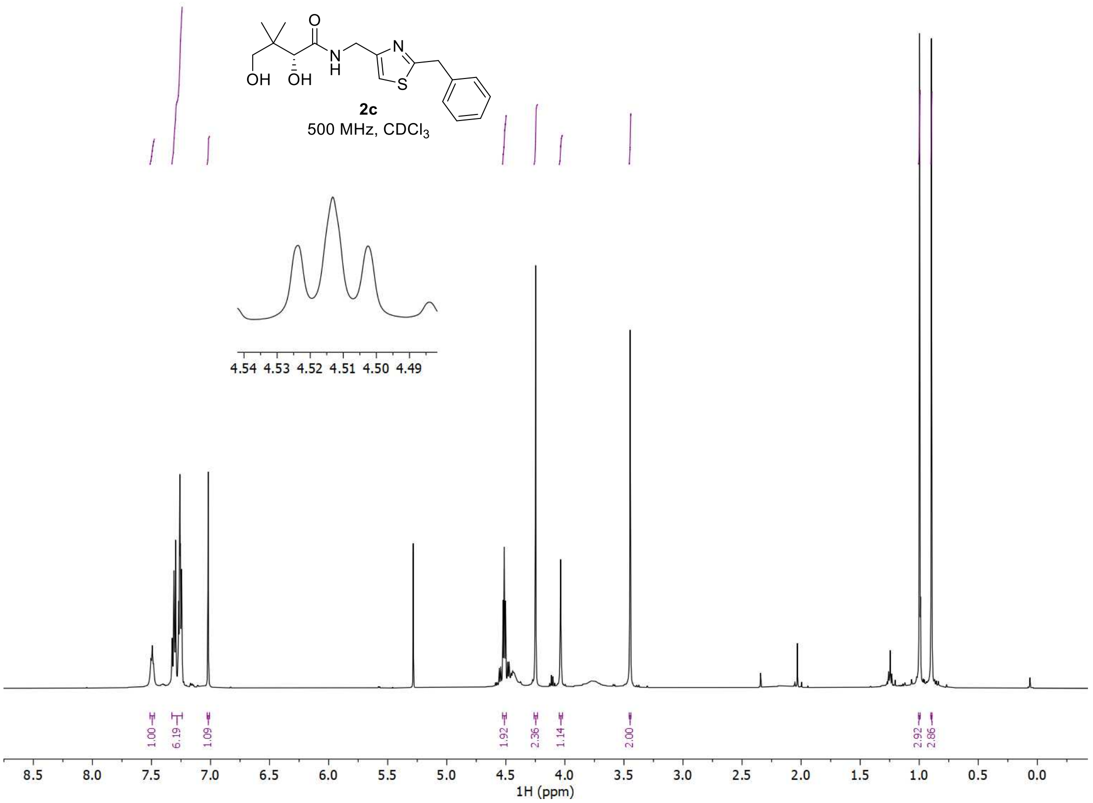


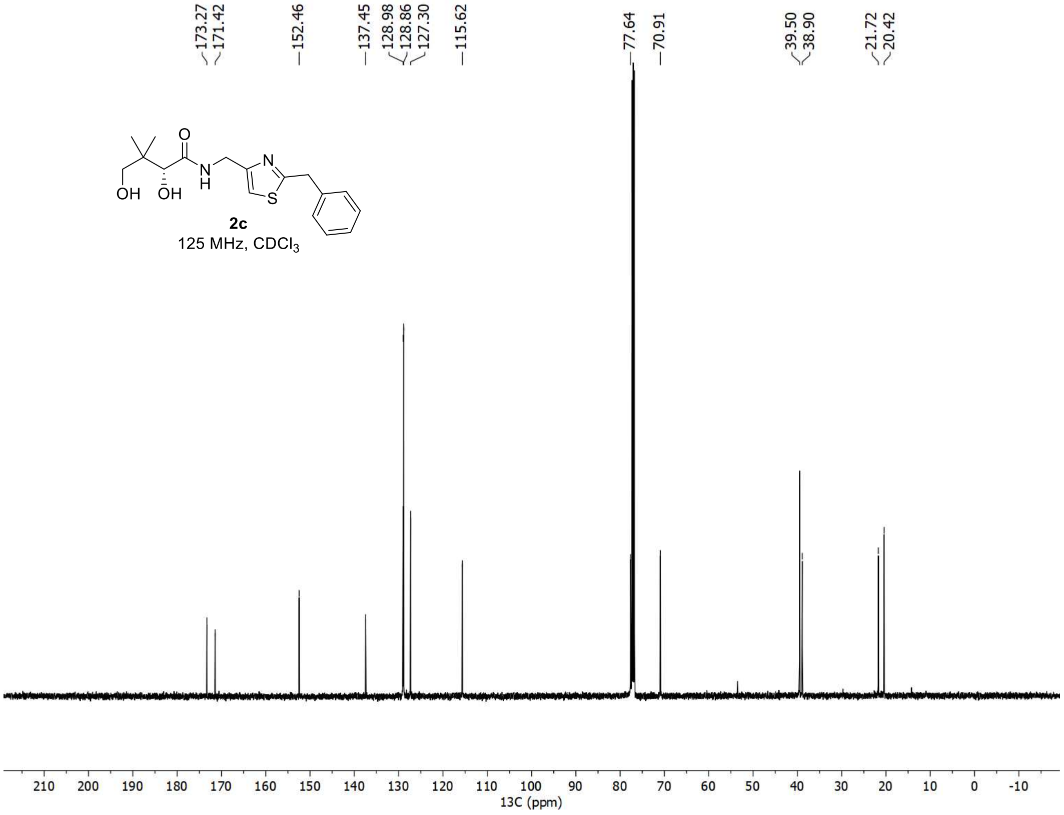

<sup></sup>

[1]H and [13]C spectra of **2.5d**

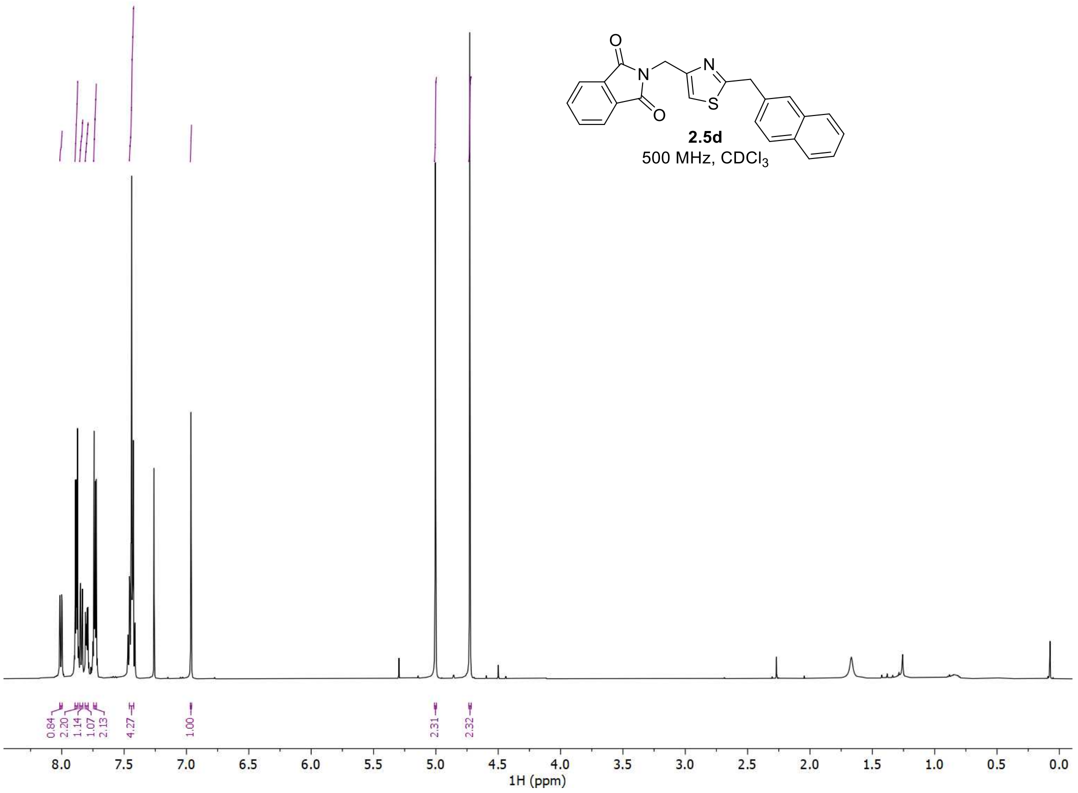


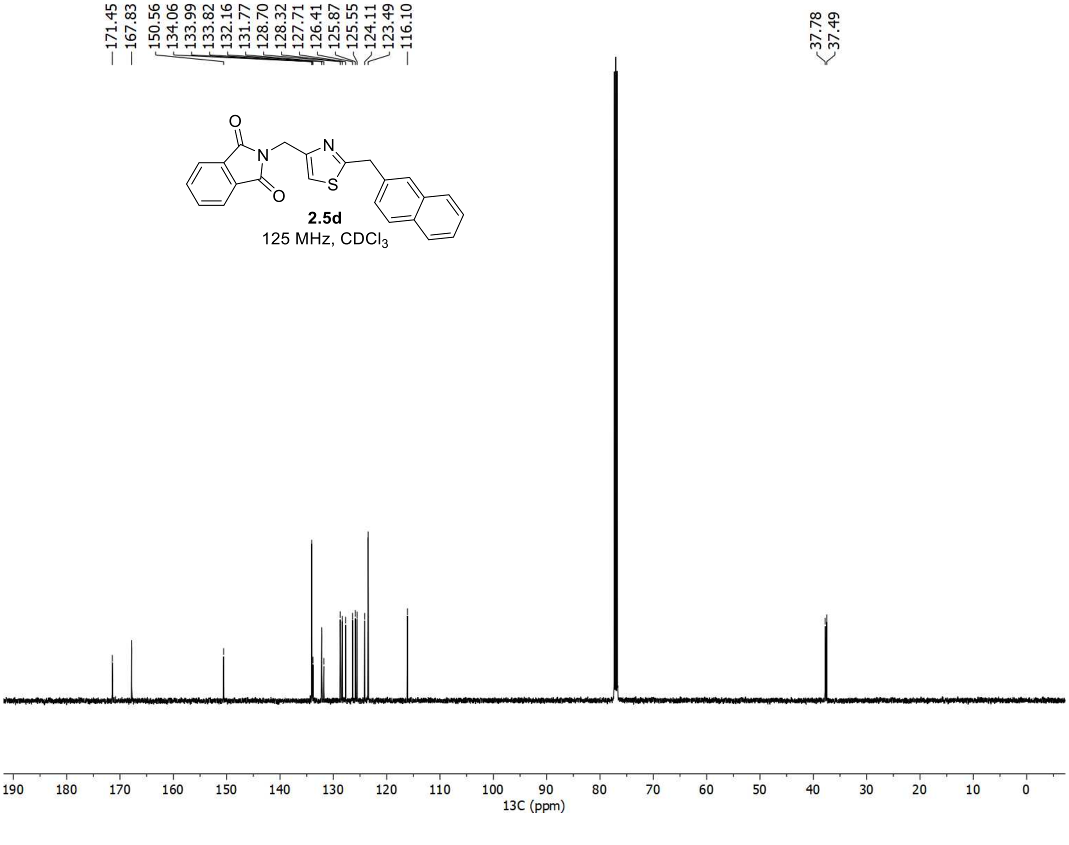

[1]H and [13]C spectra of **2.6d**

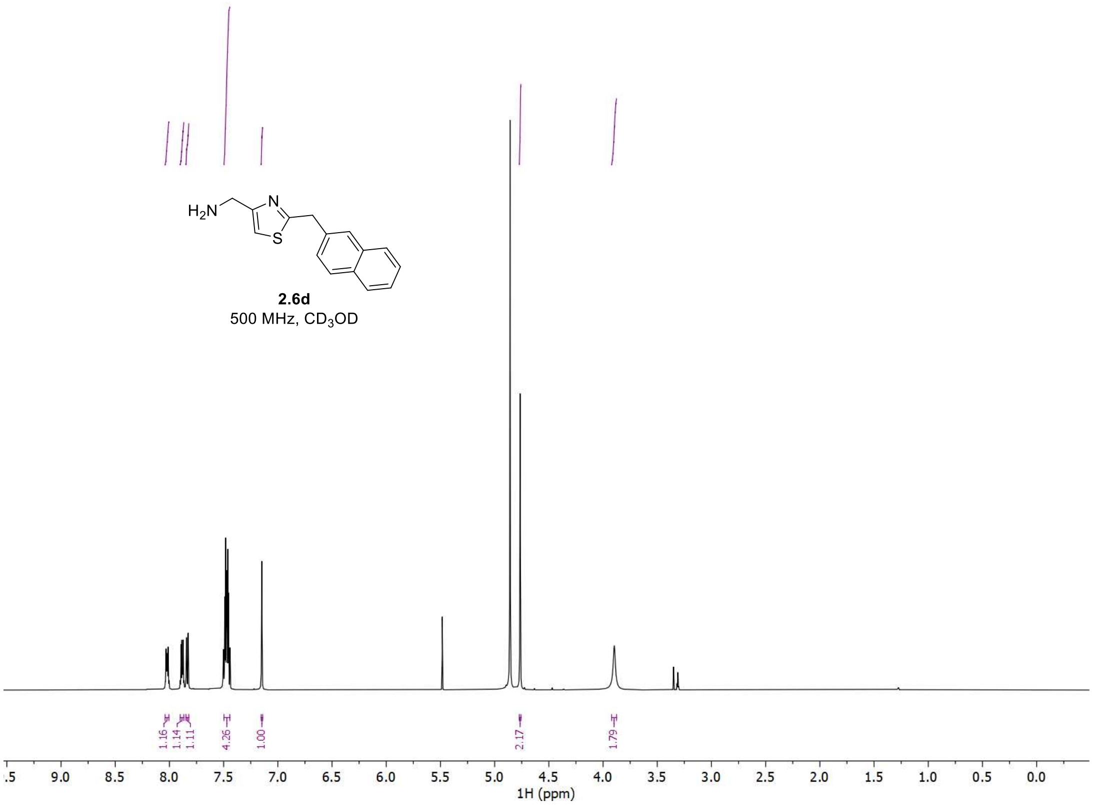


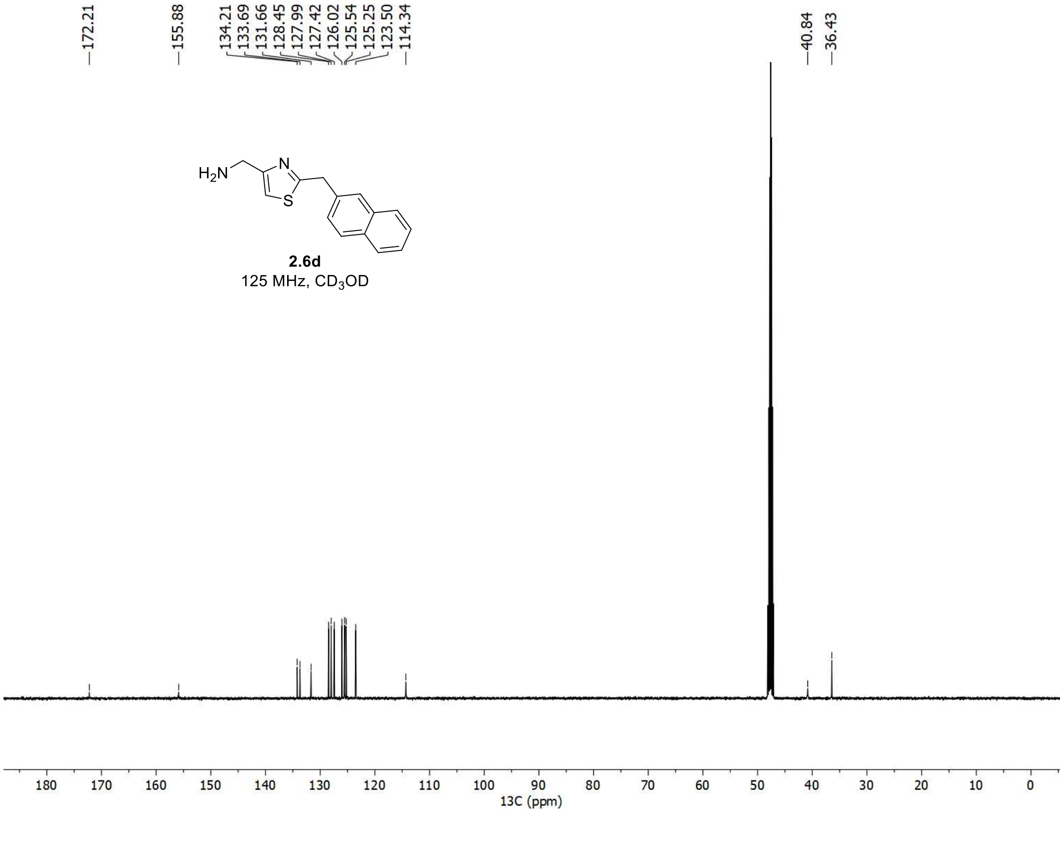

[1]H and [13]C spectra of **2d**

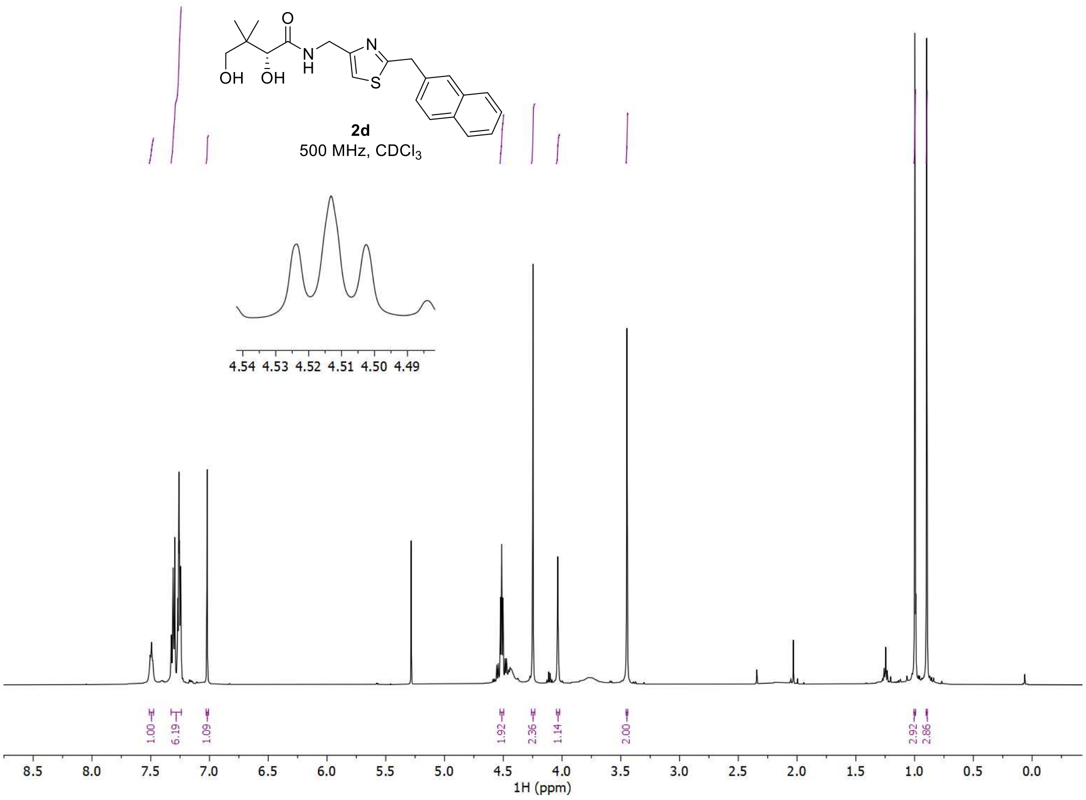


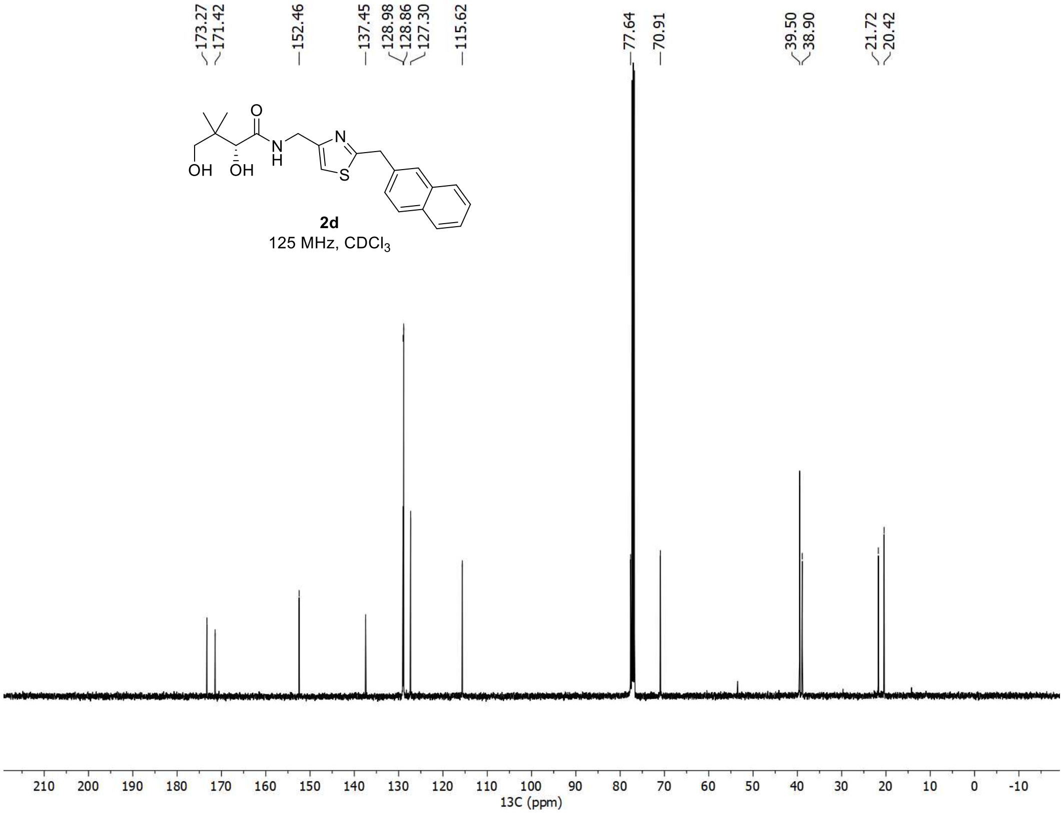

<u>$^{1}$H and $^{13}$C spectra of **2.5e**</u>

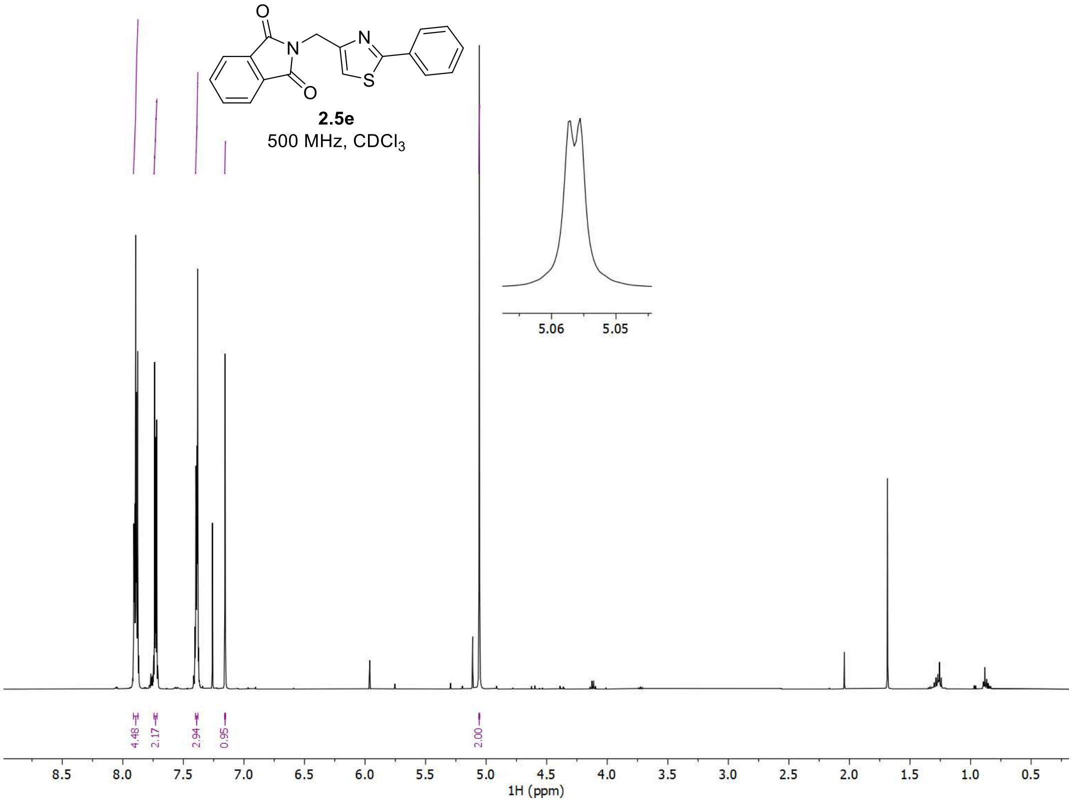


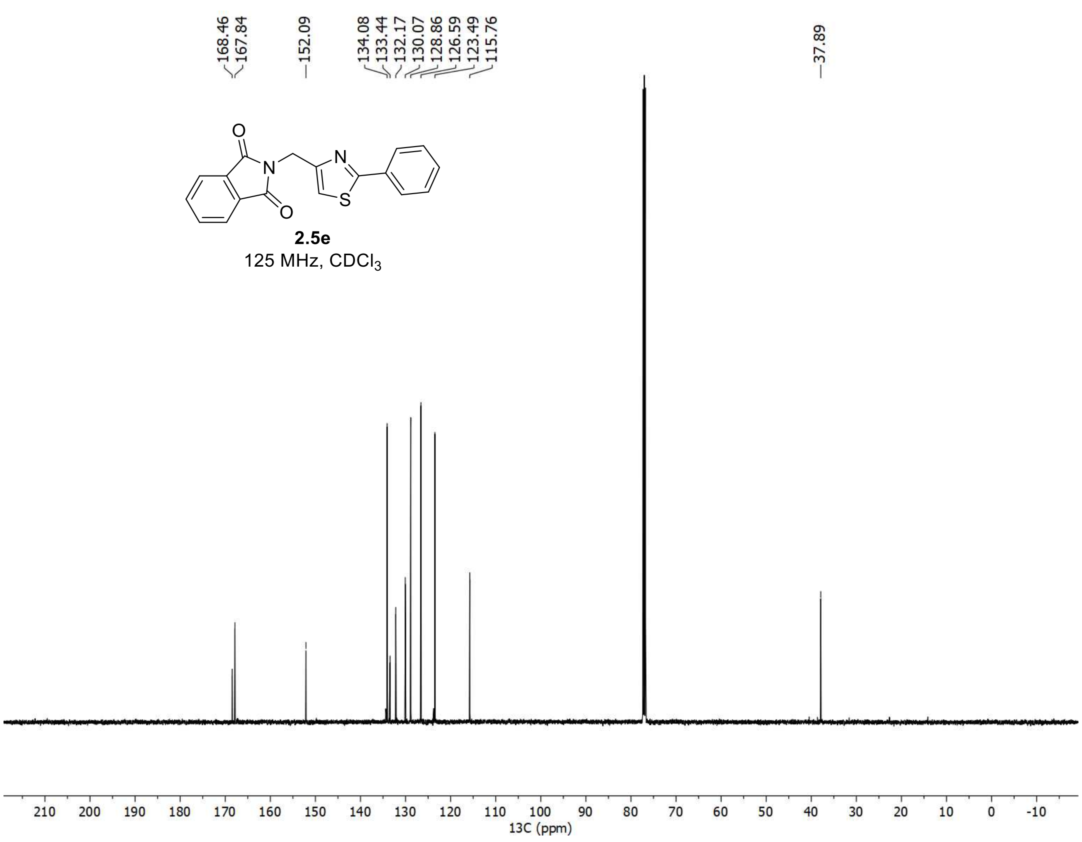

[1]H and [13]C spectra of **2.6e**

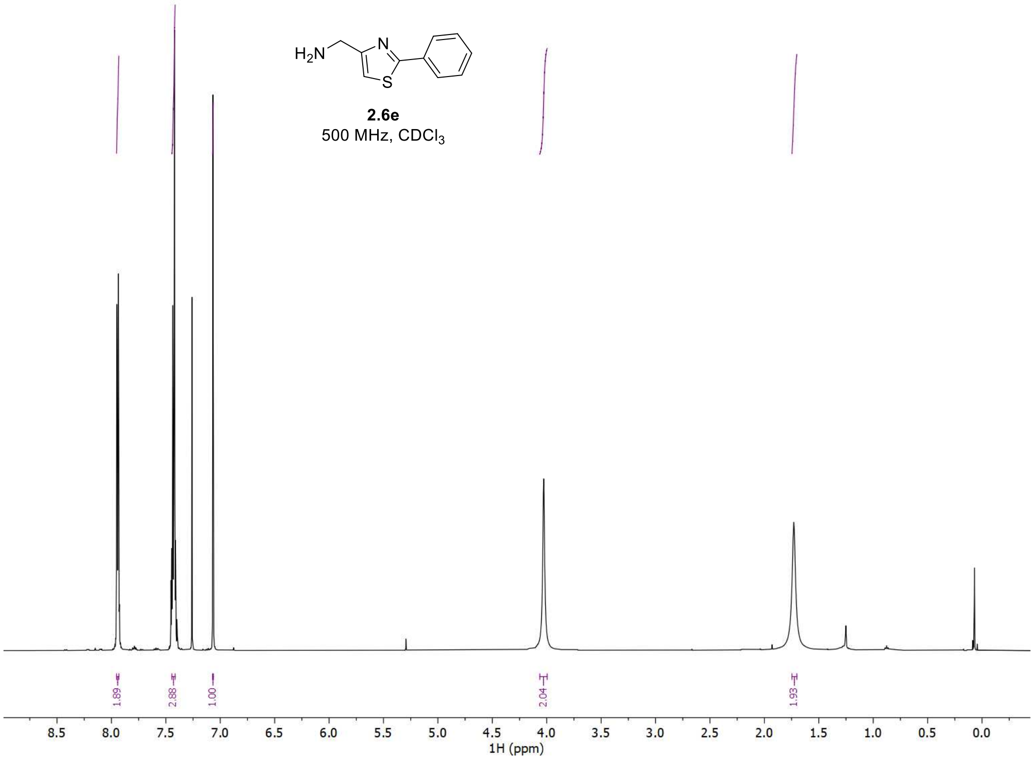


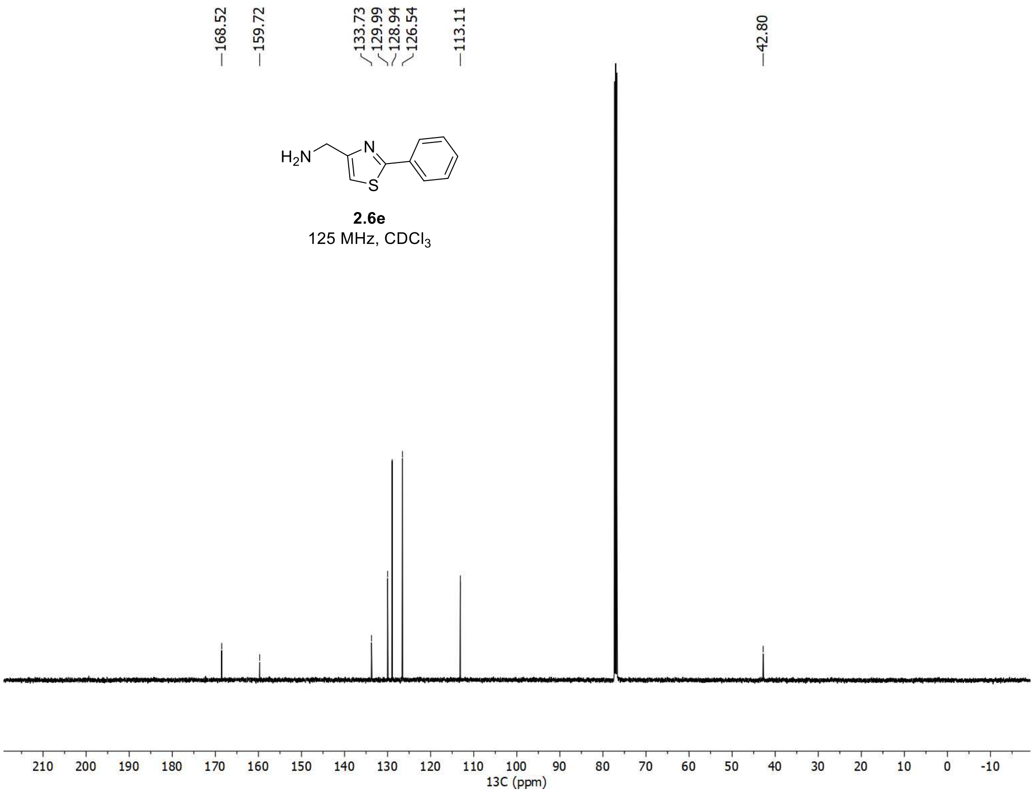

[1]H and [13]C spectra of **2e**

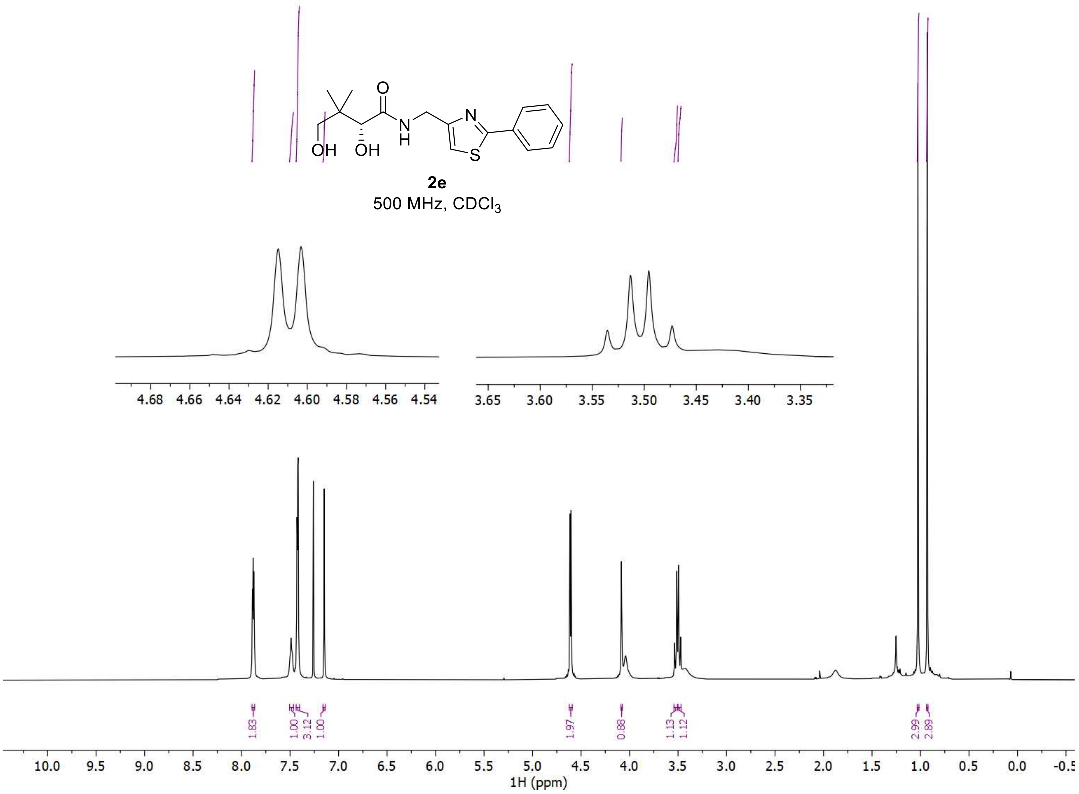


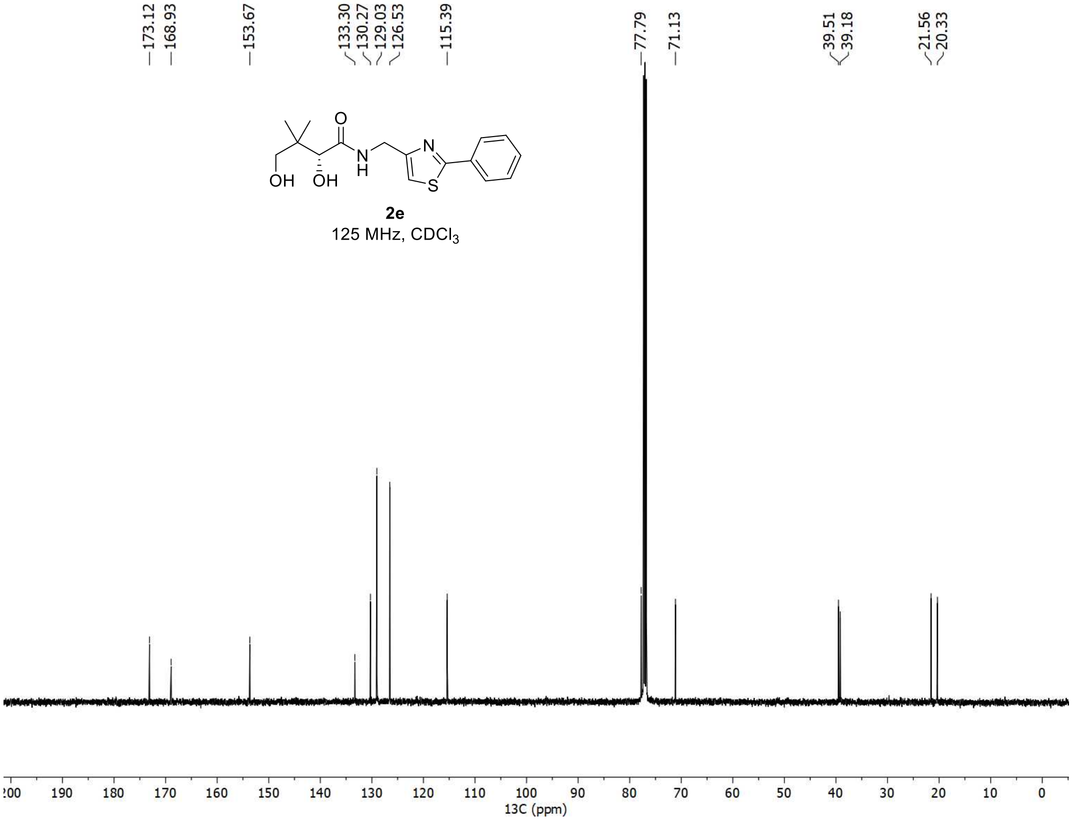

[1]H and [13]C spectra of **2.5f**

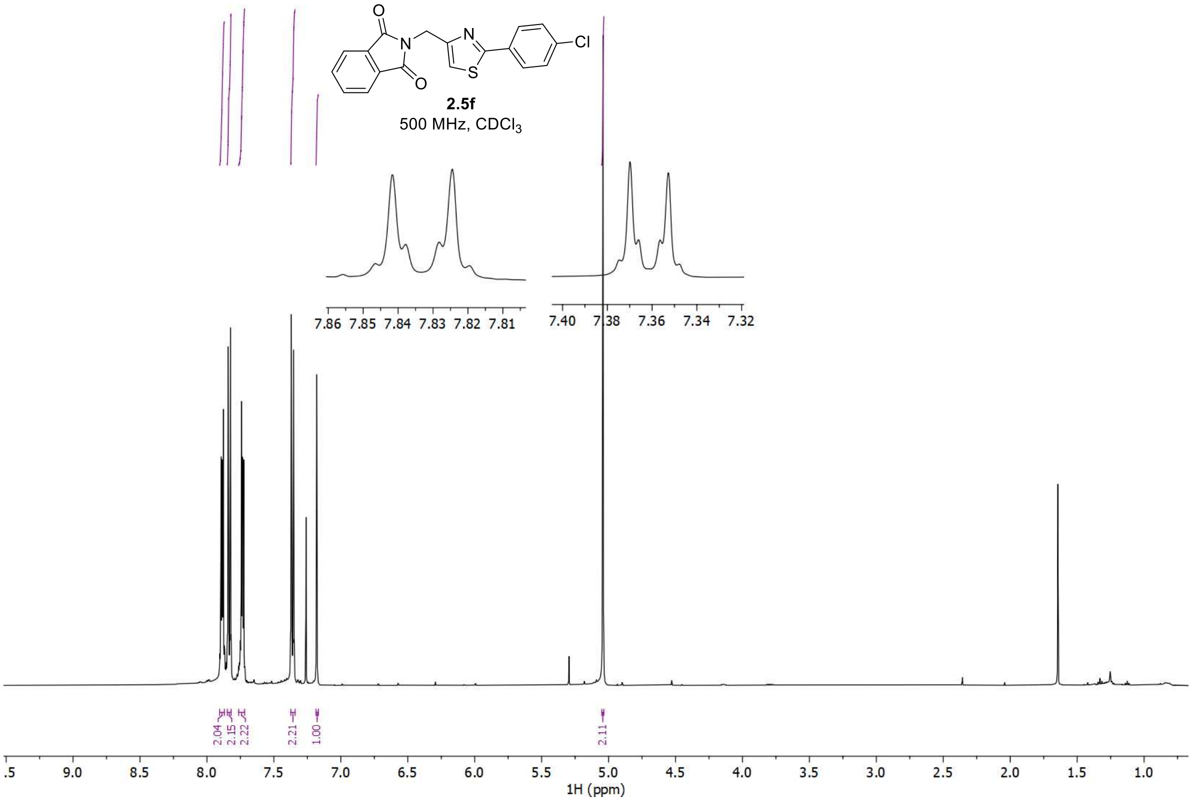


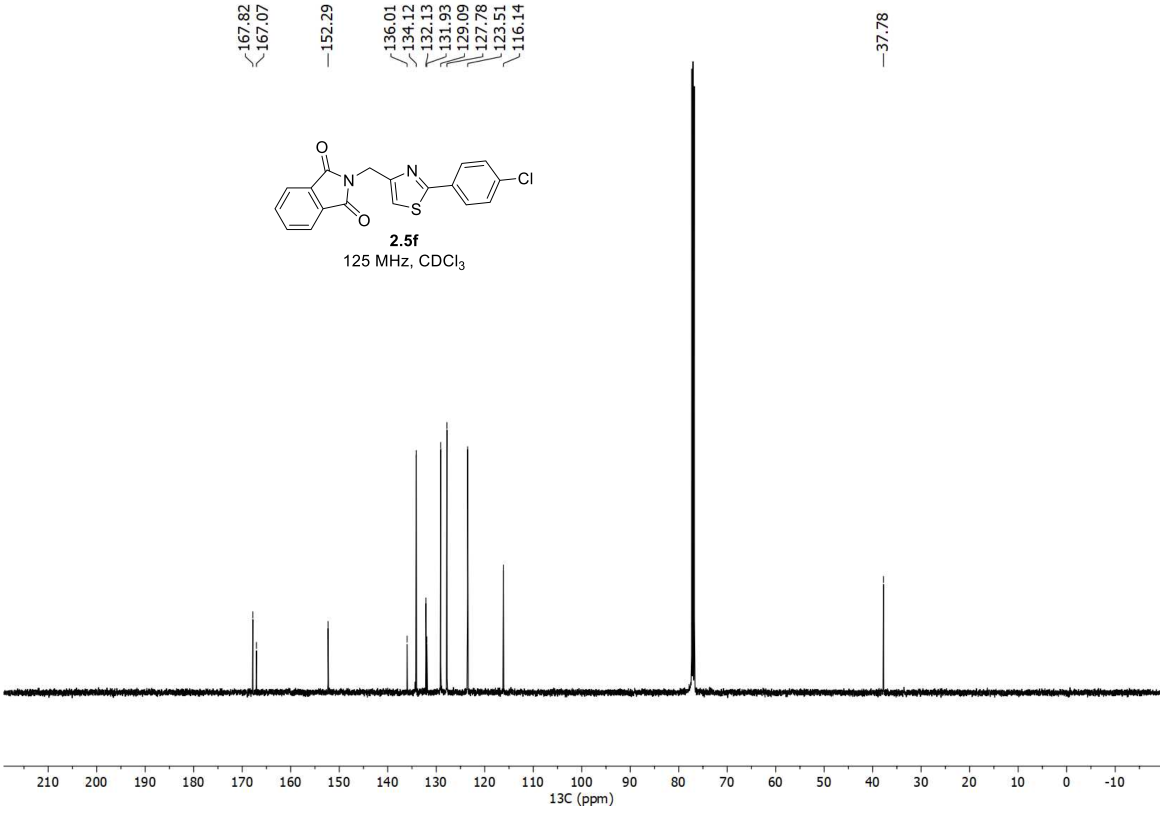

$^{1}H$ and $^{13}C$ spectra of **2.6f**

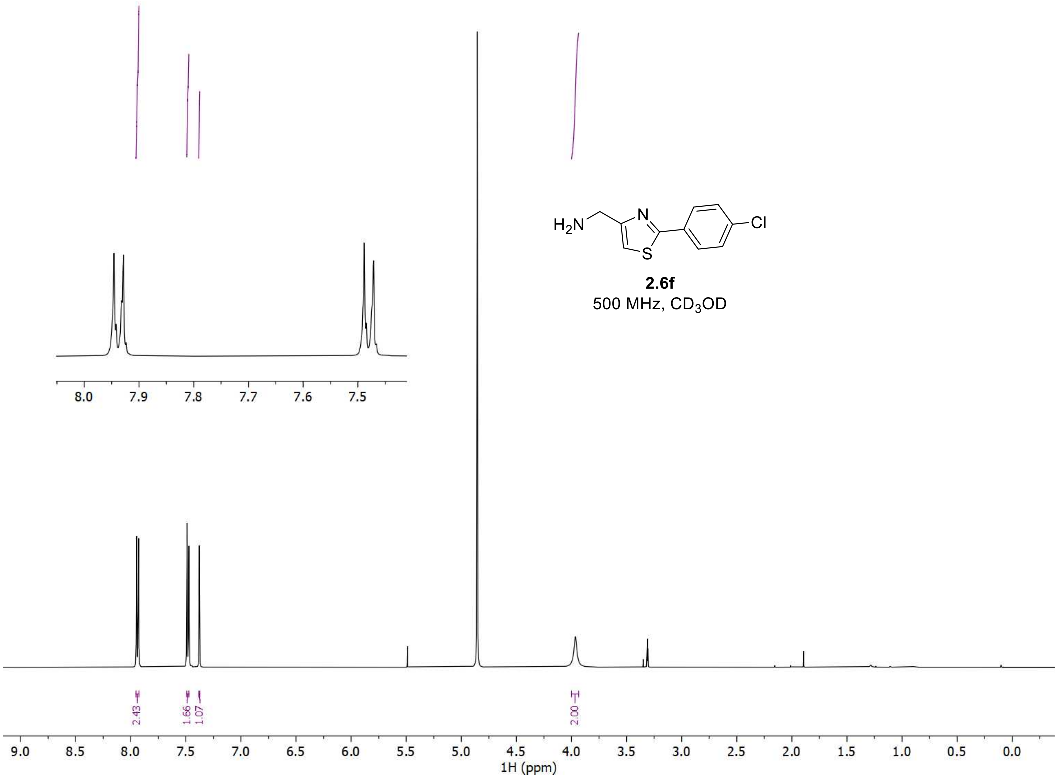


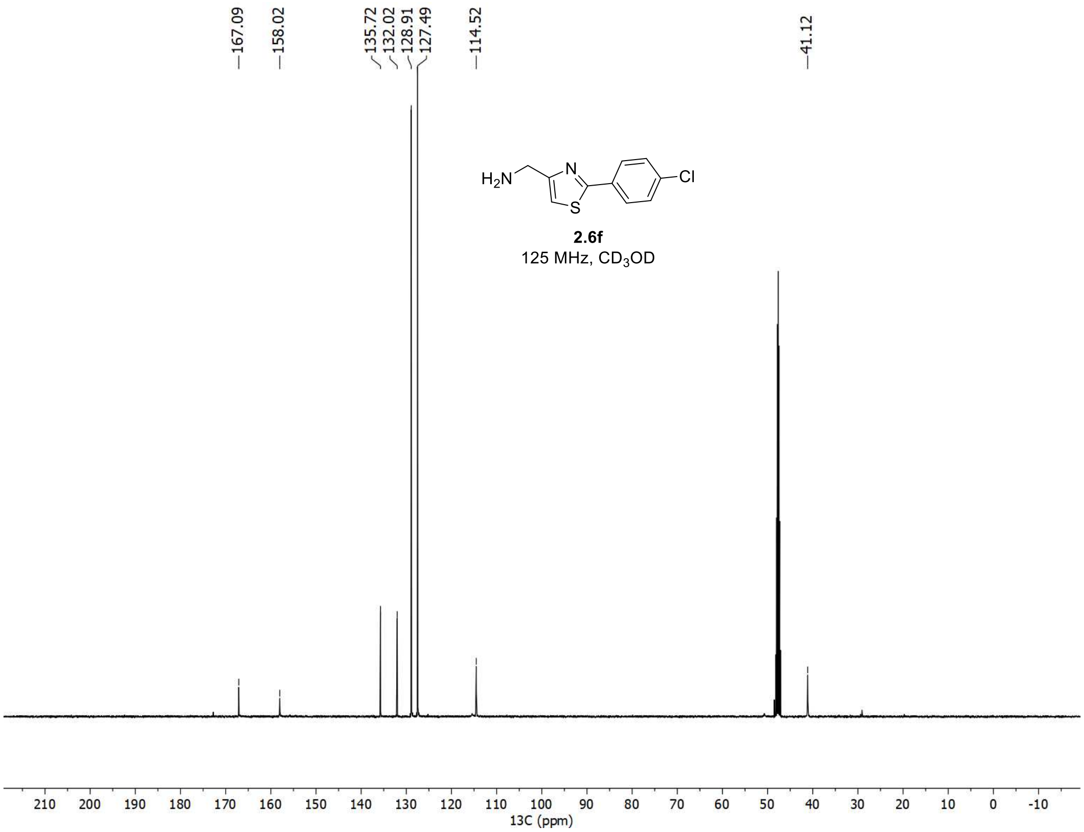

[1]H and [13]C spectra of **2f**

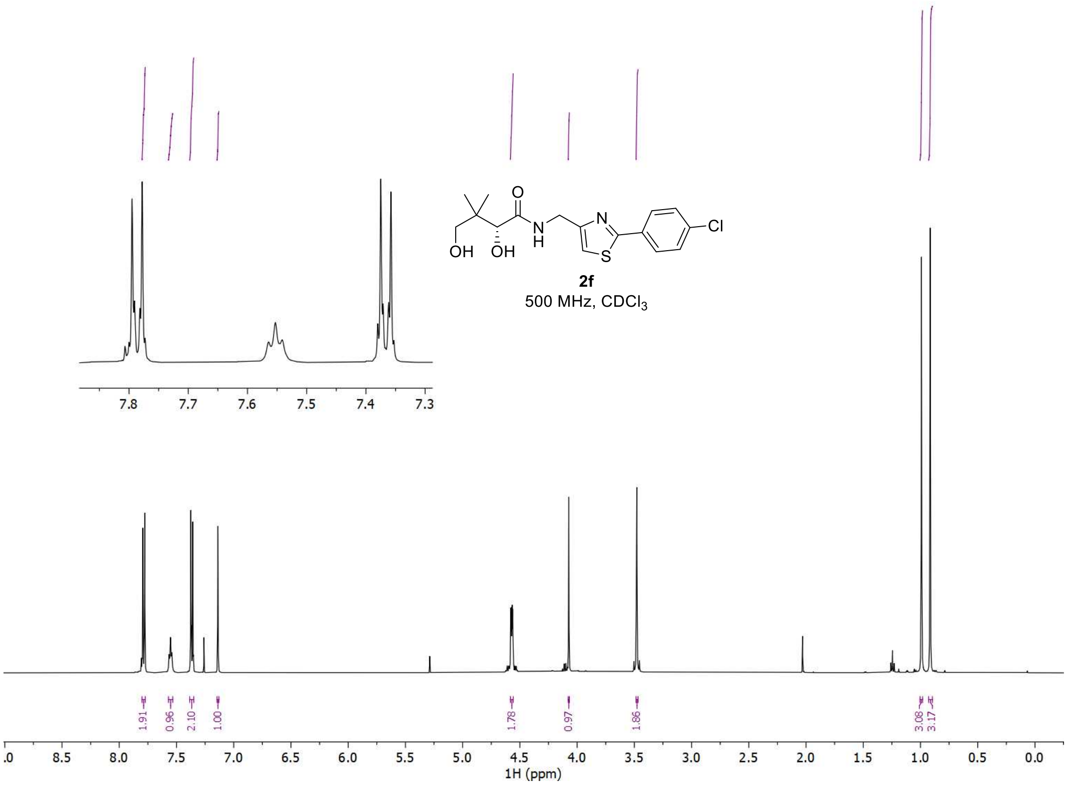


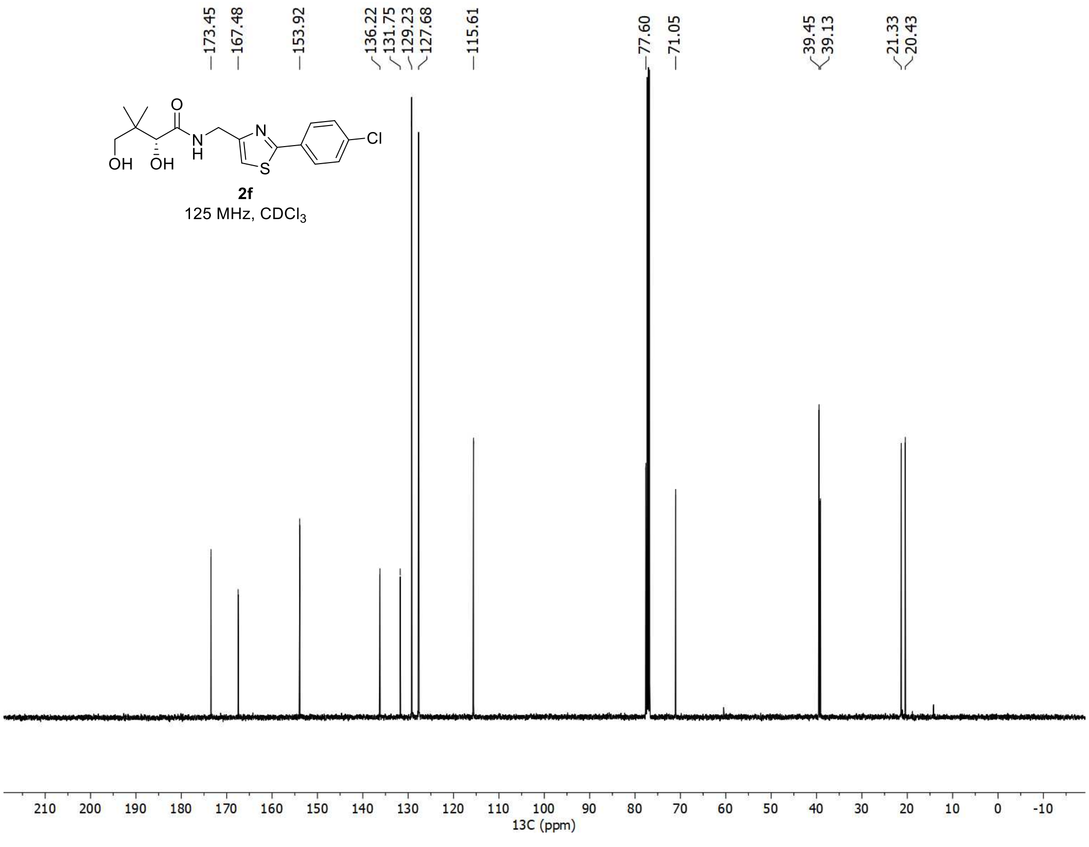

$^{1}H$, $^{19}F$, and $^{13}C$ spectra of **2.5g**

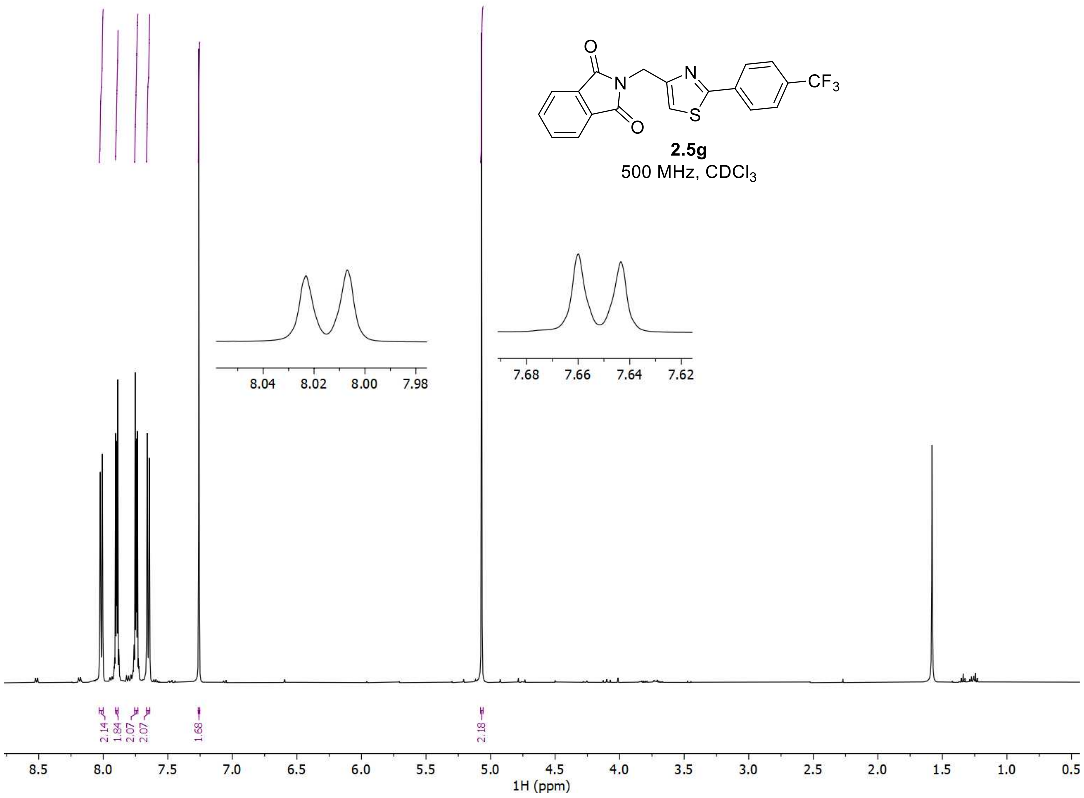


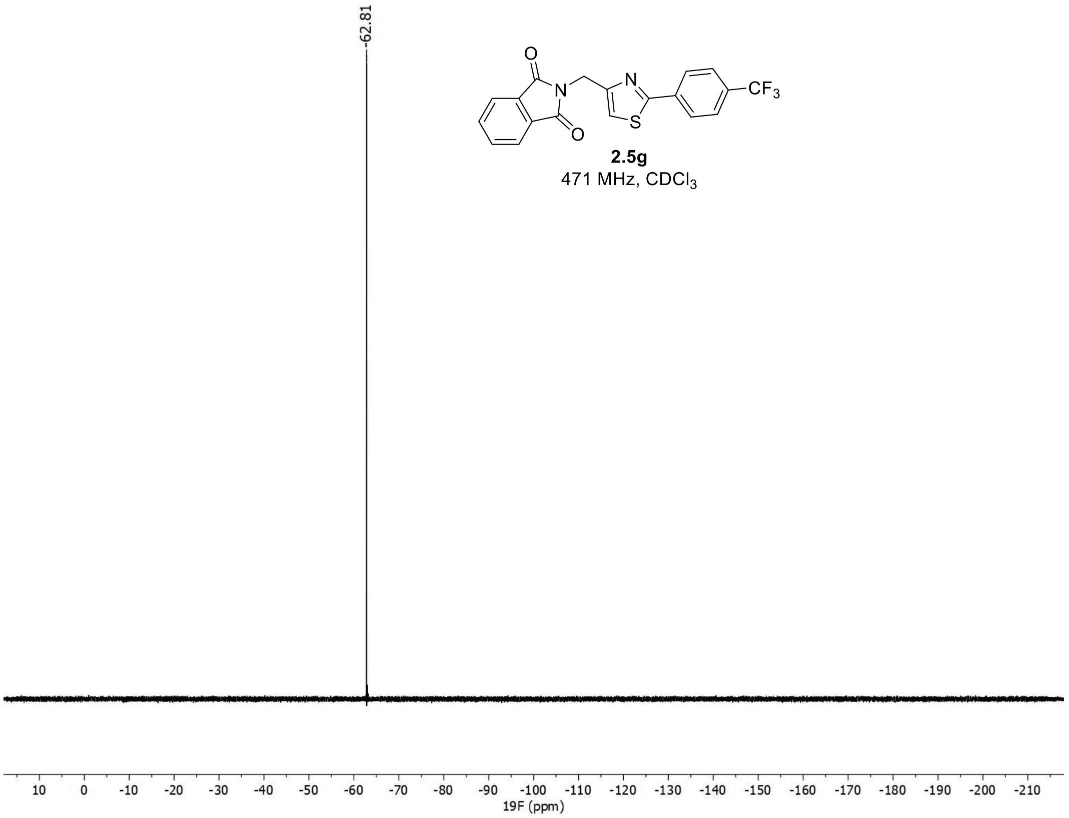

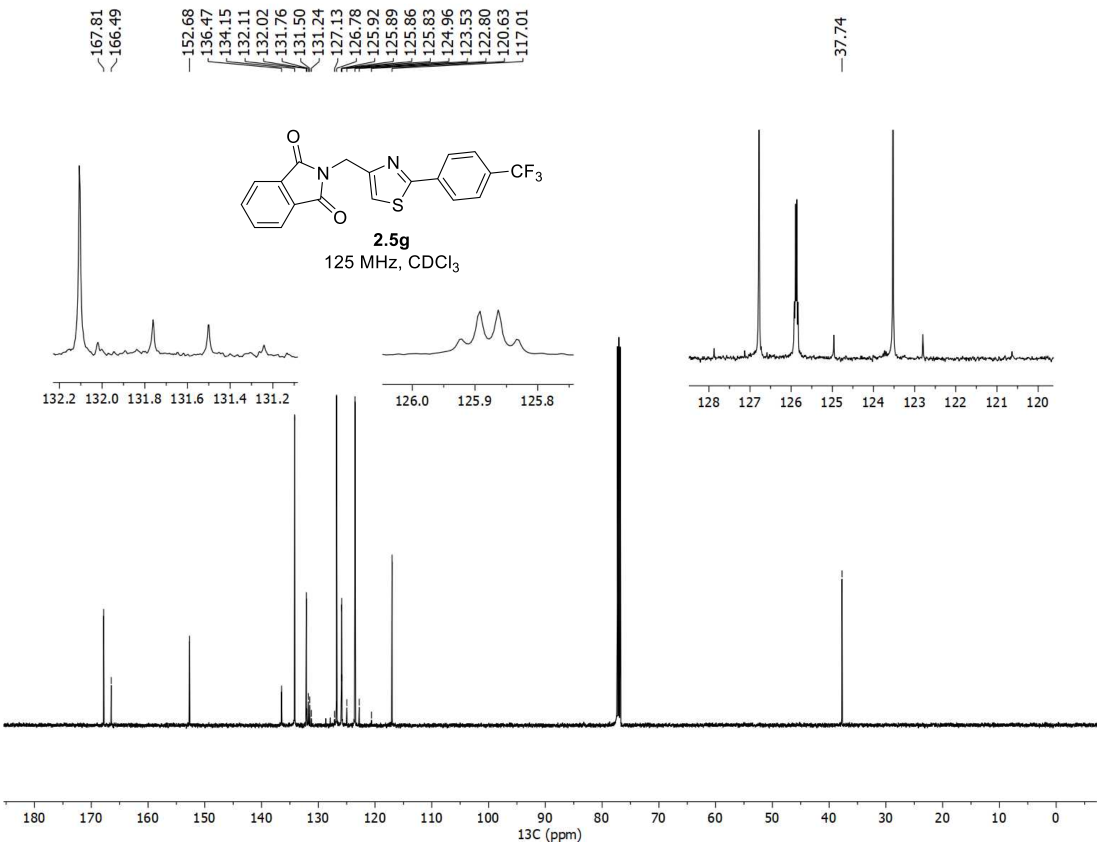

167.81
166.49
152.68
136.47
134.15
132.11
132.02
131.76
131.50
131.24
127.13
126.78
125.92
125.89
125.86
125.83
124.96
123.53
122.80
120.63
117.01
37.74
O
N
O
N
S
CF3
2.5g
125 MHz, CDCl3
132.2 132.0 131.8 131.6 131.4 131.2
126.0 125.9 125.8
128 127 126 125 124 123 122 121 120
180 170 160 150 140 130 120 110 100 90 80 70 60 50 40 30 20 10 0
13C (ppm)

$^{1}H$, $^{19}F$, and $^{13}C$ spectra of **2.6g**

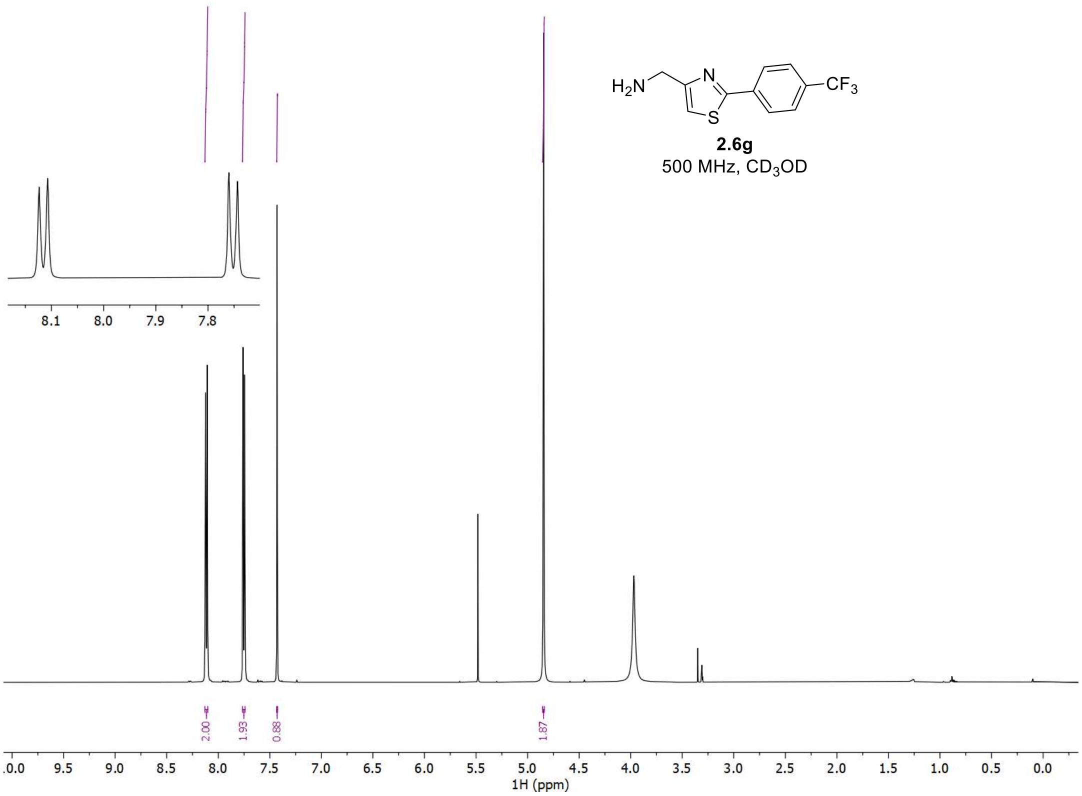


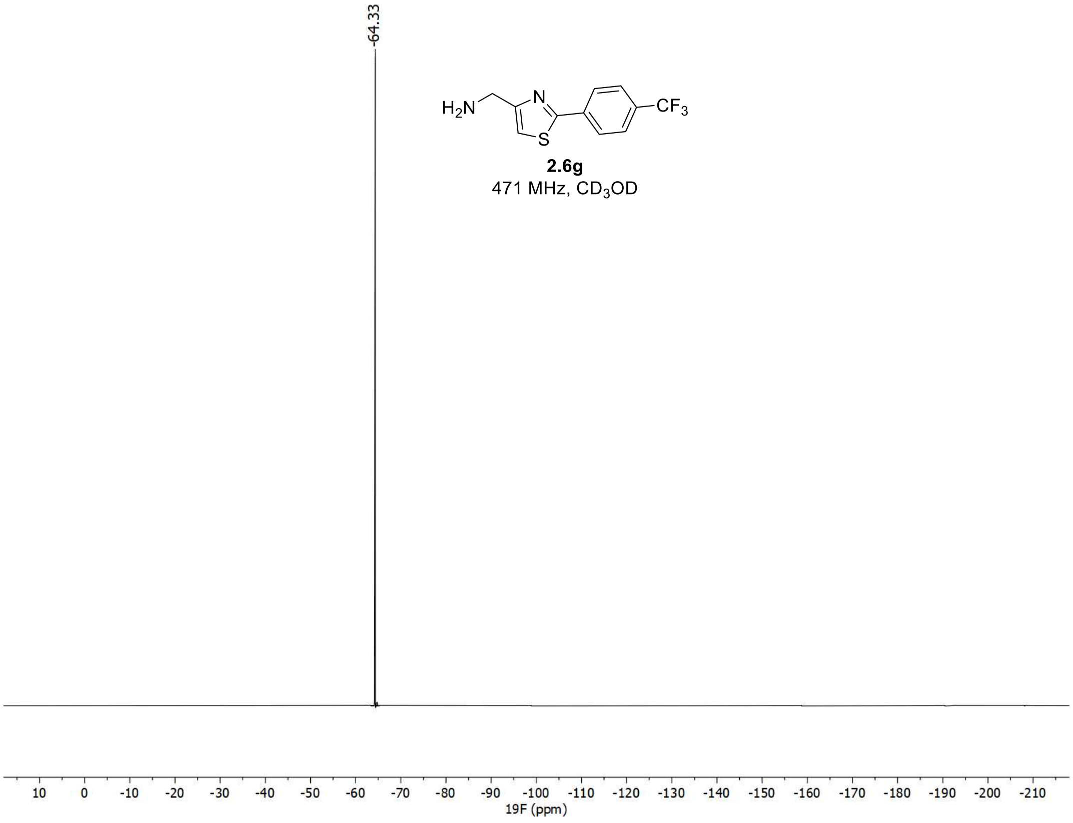

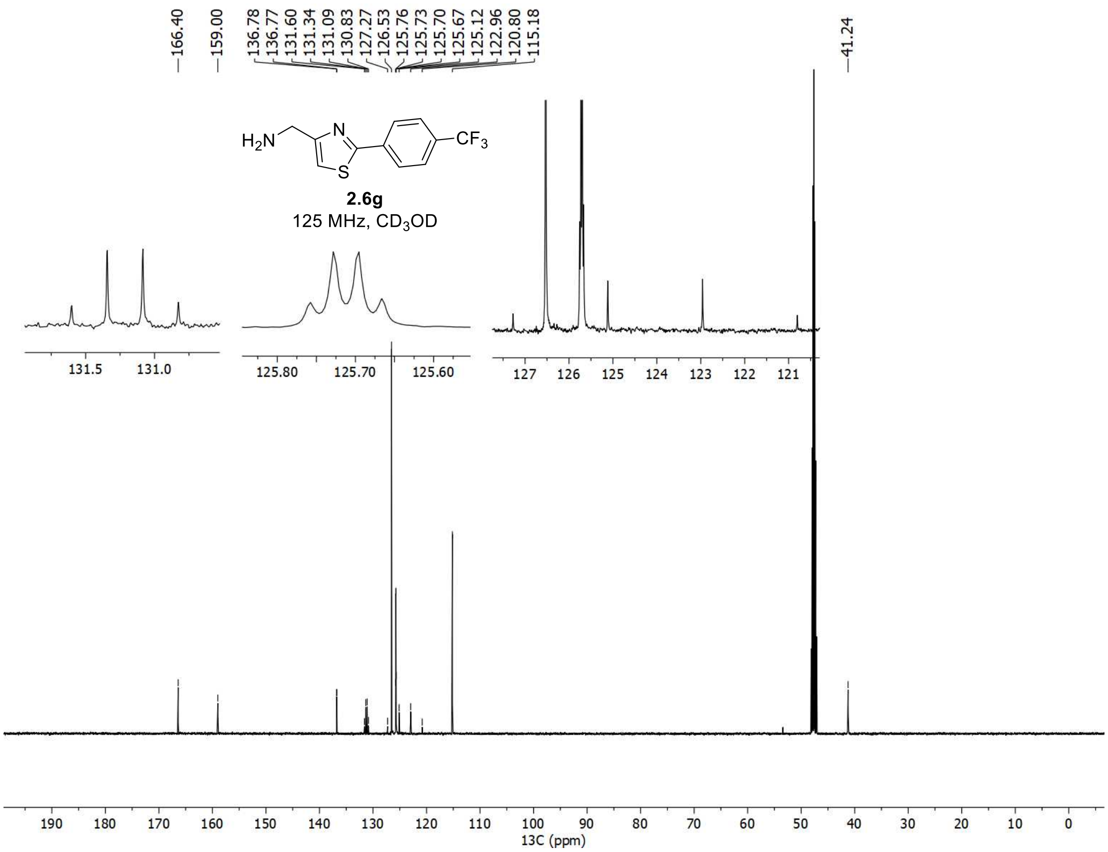

166.40
159.00
136.78
136.77
131.60
131.34
131.09
130.83
127.27
126.53
125.76
125.73
125.70
125.67
125.12
122.96
120.80
115.18
41.24
$H_2N$
N
S
$CF_3$
2.6g
125 MHz, $CD_3OD$
13C (ppm)

$^{1}H$, $^{19}F$, and $^{13}C$ spectra of **2g**

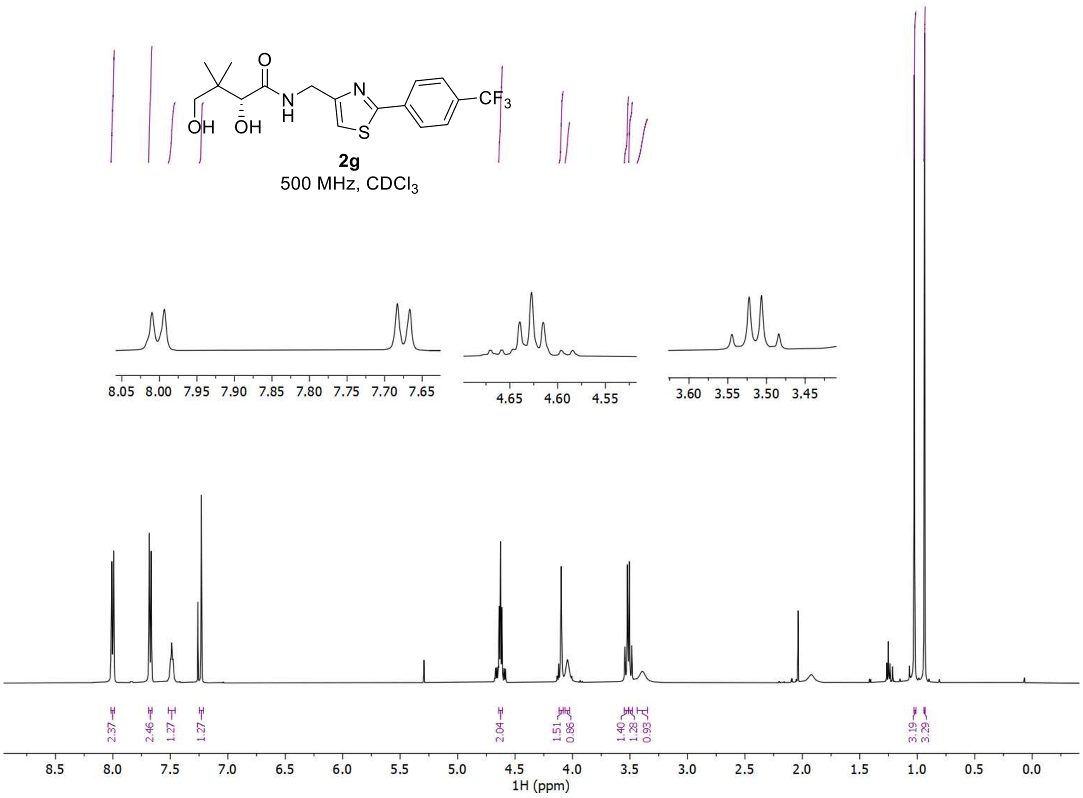


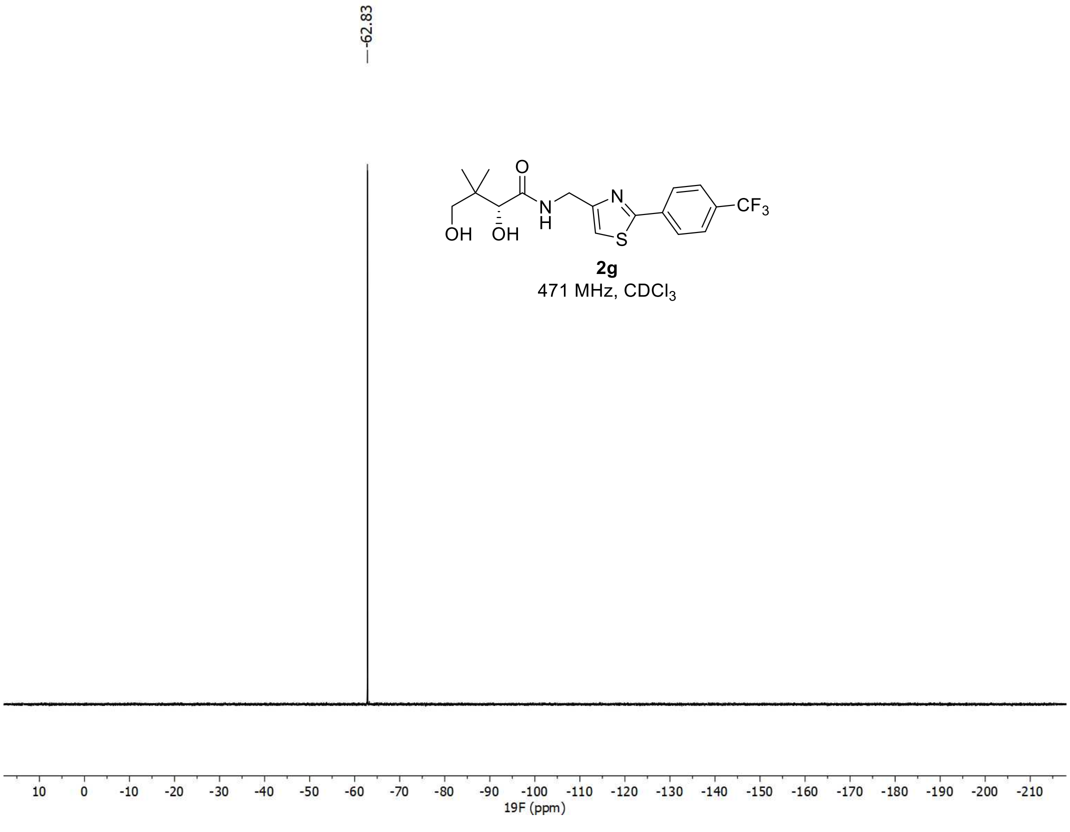

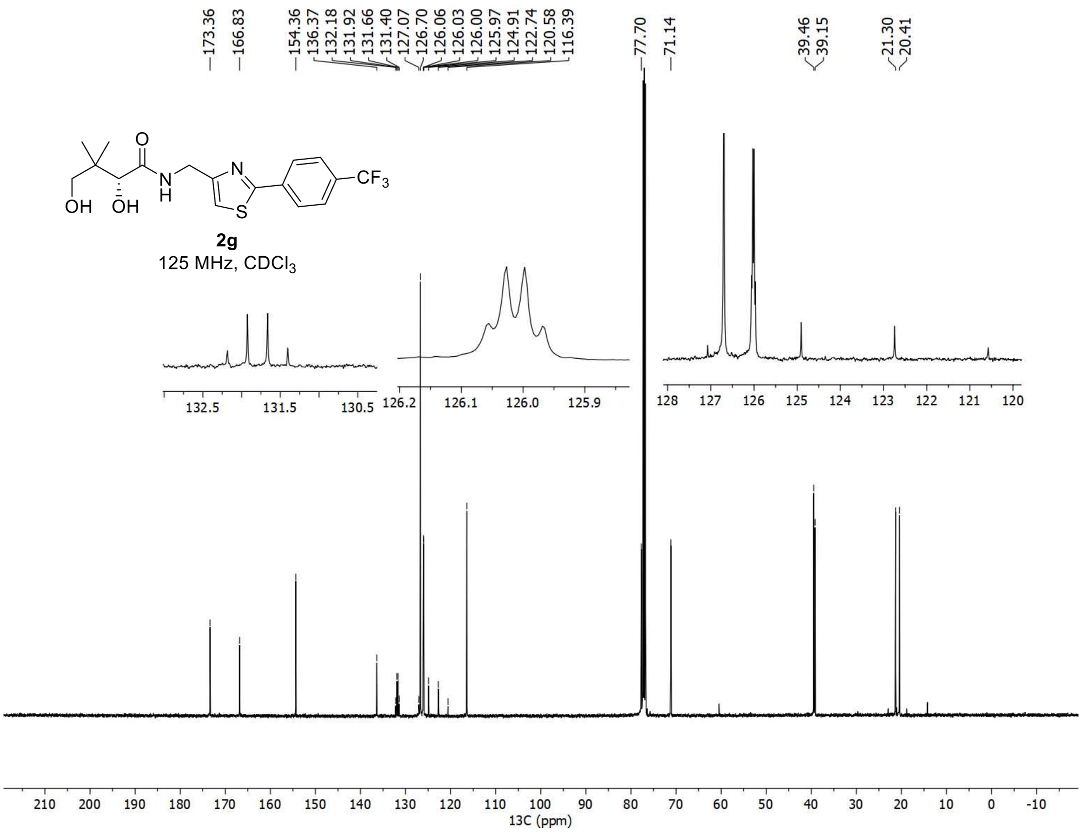

173.36
166.83
154.36
136.37
132.18
131.92
131.66
131.40
127.07
126.70
126.06
126.03
126.00
125.97
124.91
122.74
120.58
116.39
77.70
71.14
39.46
39.15
21.30
20.41
OH
OH
O
N
H
N
S
$CF_3$
2g
125 MHz, $CDCl_3$
13C (ppm)

<sup></sup>

[1]H, [19]F, and [13]C spectra of **2.5h**

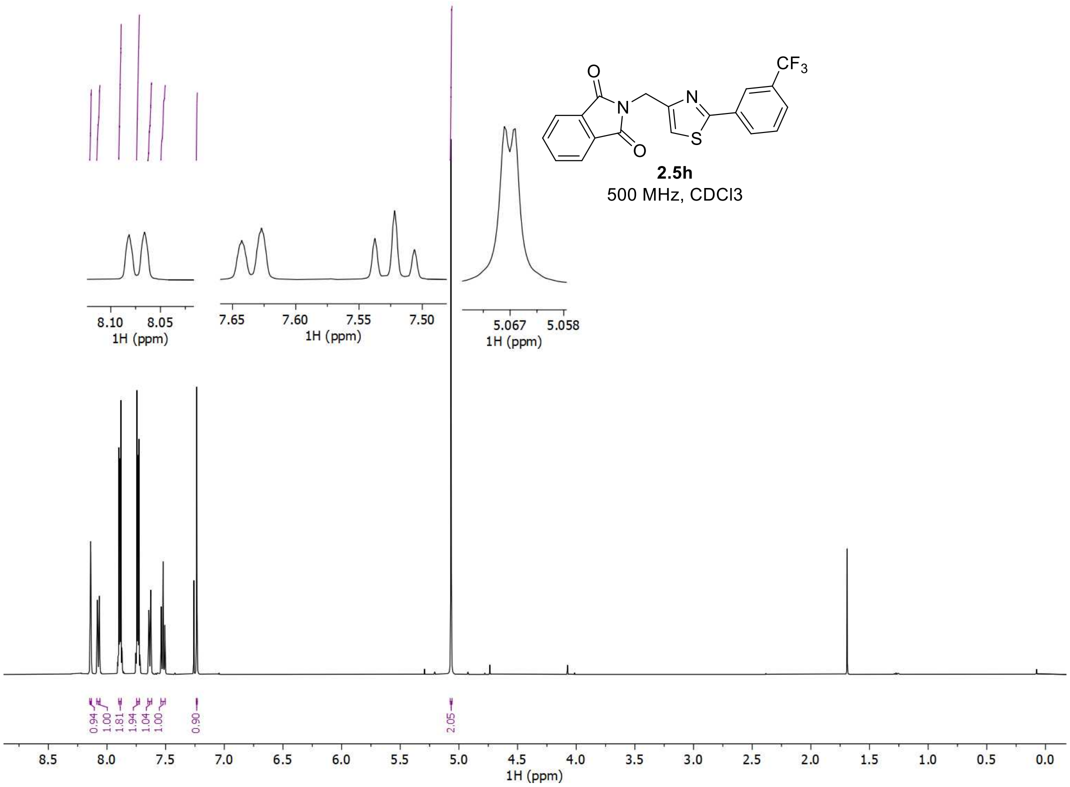


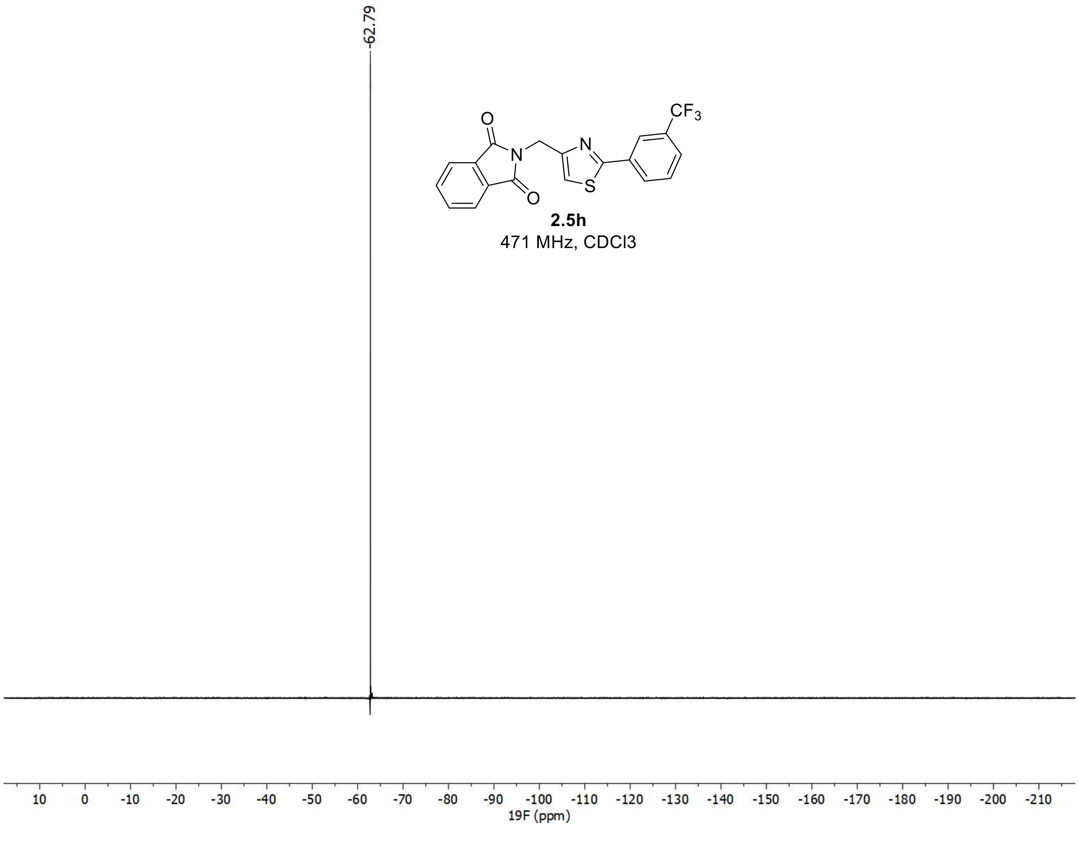

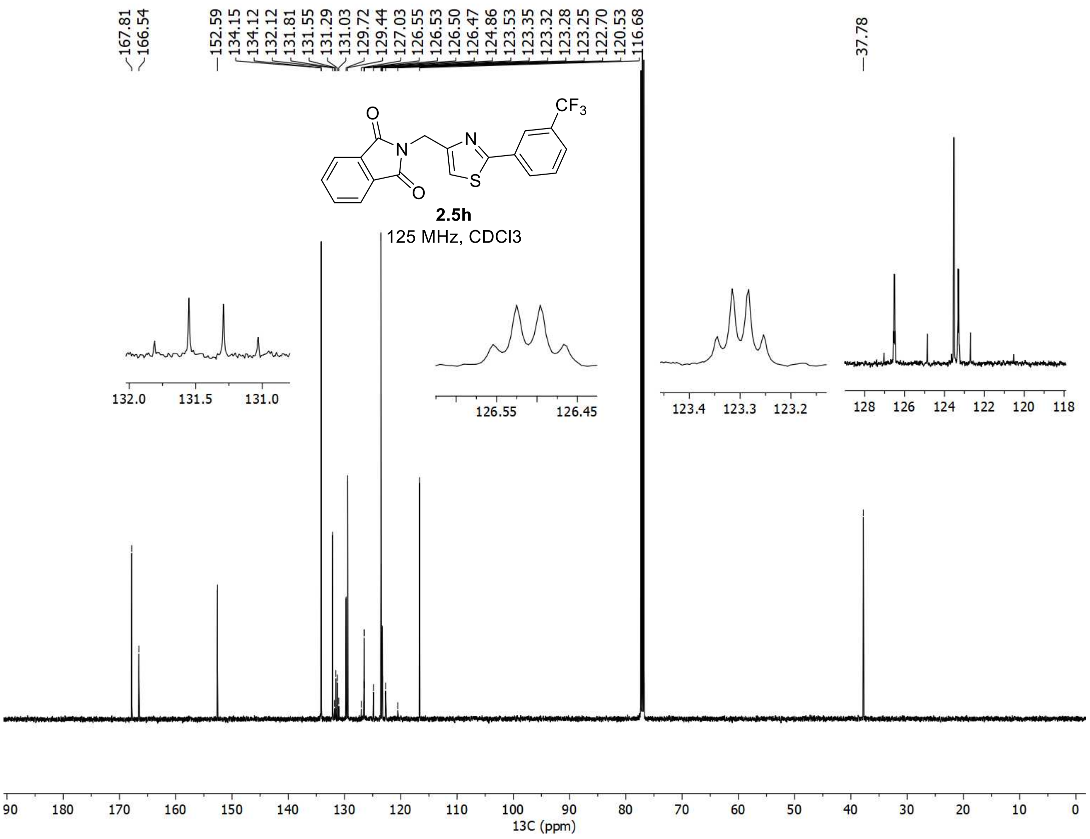

167.81
166.54
152.59
134.15
134.12
132.12
131.81
131.55
131.29
131.03
129.72
129.44
127.03
126.55
126.53
126.50
126.47
124.86
123.53
123.35
123.32
123.28
123.25
122.70
120.53
116.68
37.78
CF3
N
S
O
2.5h
125 MHz, CDCl3
13C (ppm)

<u>[1]H, [19]F, and [13]C spectra of **2.6h**</u>

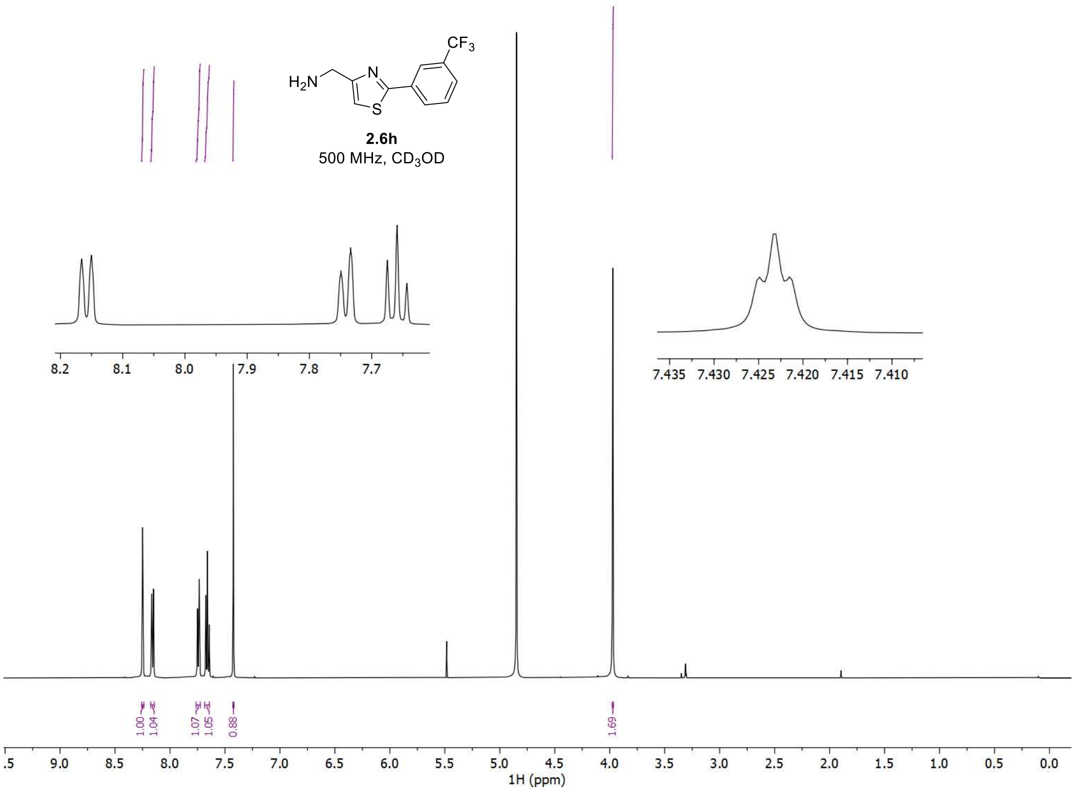


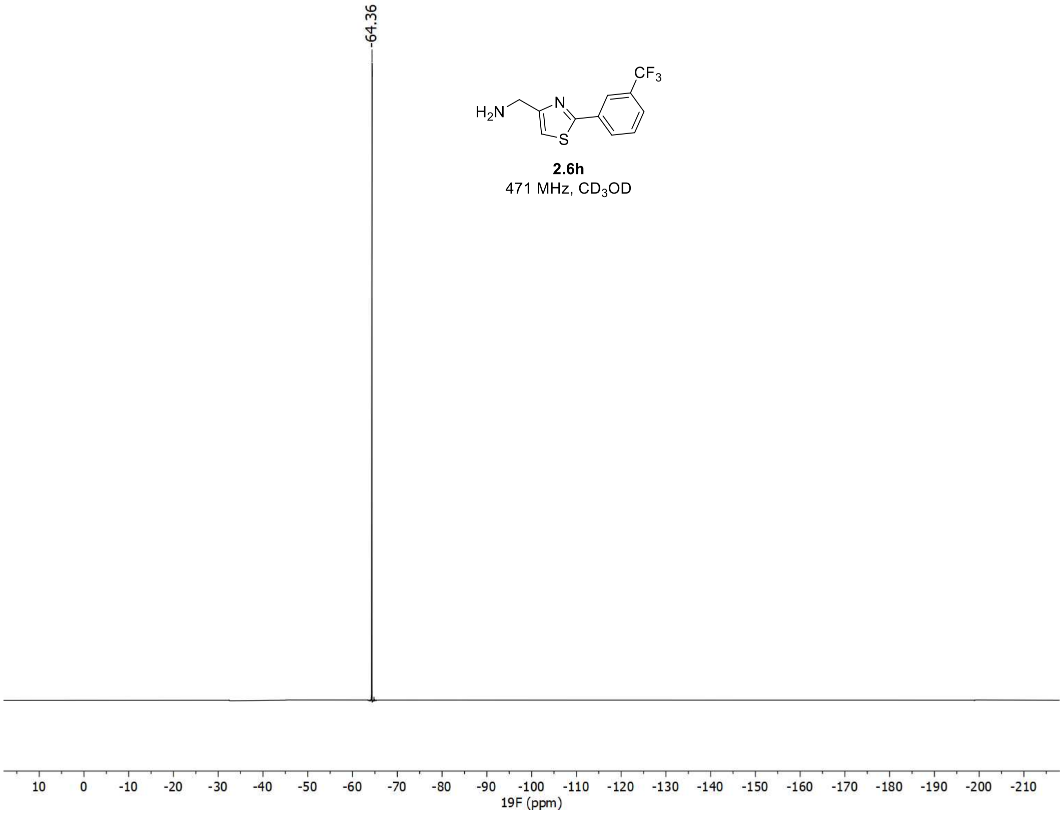

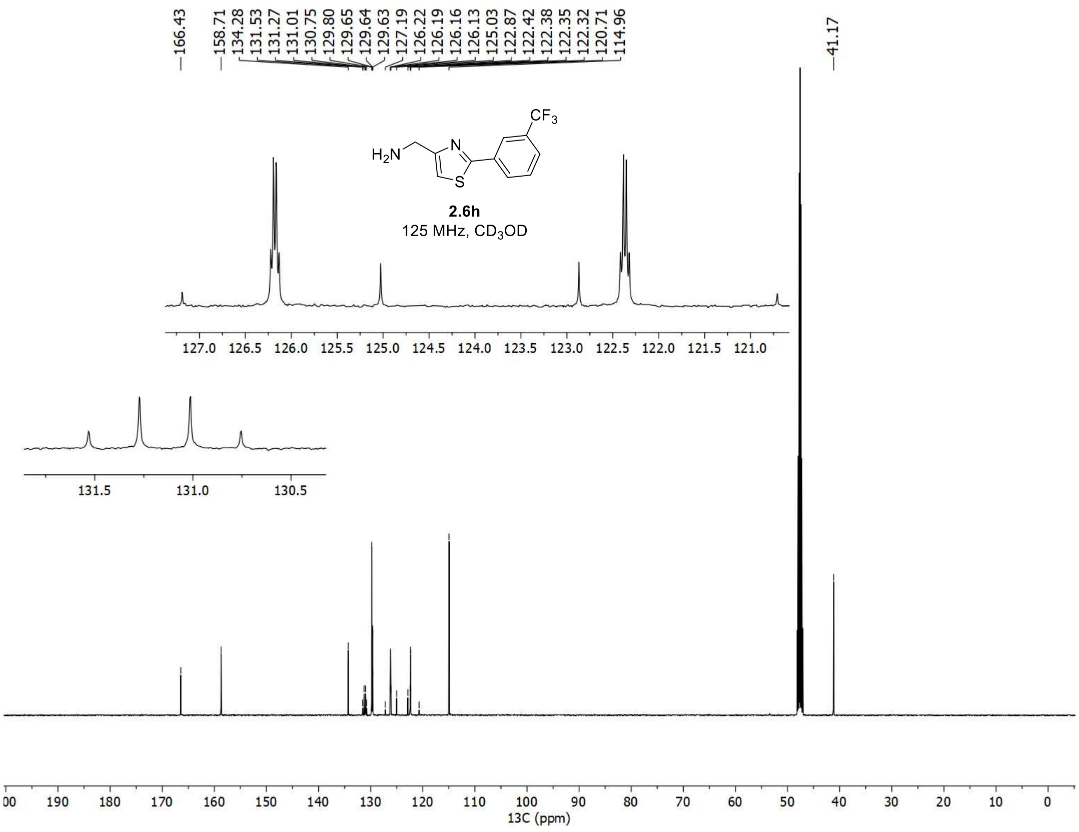

166.43
158.71
134.28
131.53
131.27
131.01
130.75
129.80
129.65
129.64
129.63
127.19
126.22
126.19
126.16
126.13
125.03
122.87
122.42
122.38
122.35
122.32
120.71
114.96
41.17
CF3
H2N
N
S
2.6h
125 MHz, CD3OD
13C (ppm)

<sup></sup>

$^{1}$H, $^{19}$F, and $^{13}$C spectra of **2h**

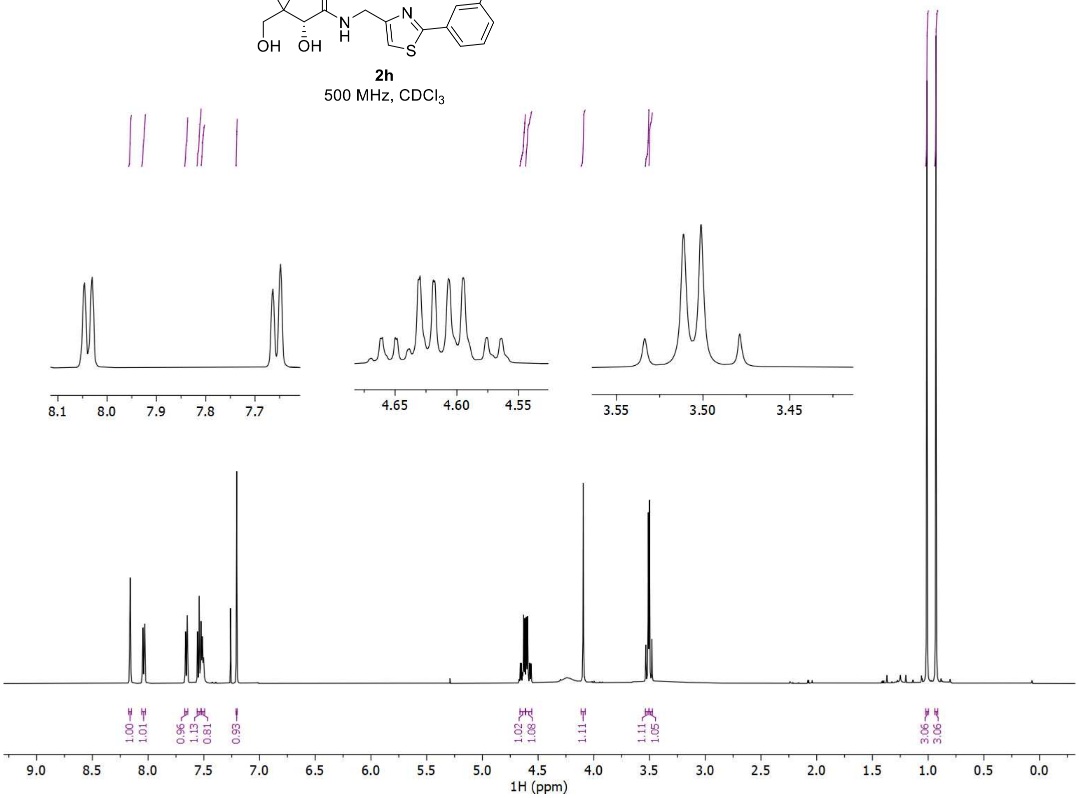


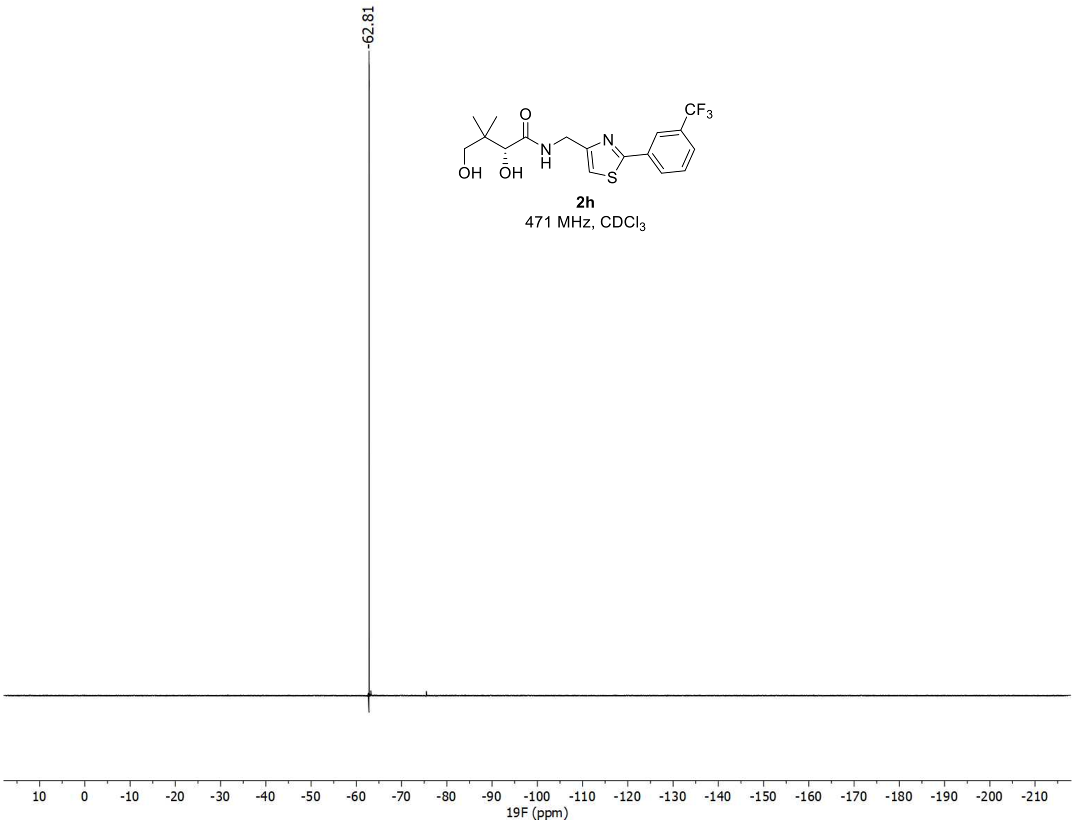

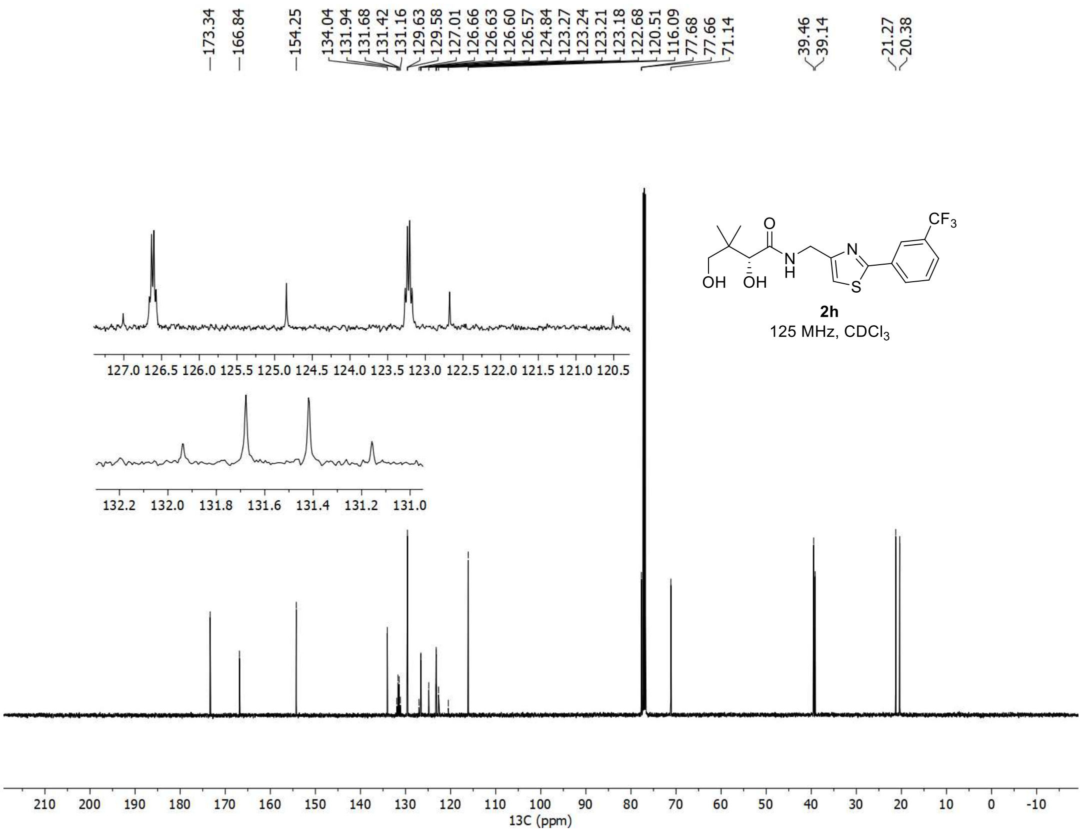
173.34
166.84
154.25
134.04
131.94
131.68
131.42
131.16
129.63
129.58
127.01
126.66
126.63
126.60
126.57
124.84
123.27
123.24
123.21
123.18
122.68
120.51
116.09
77.68
77.66
71.14
39.46
39.14
21.27
20.38
CF3
O
N
OH
OH
H
S
2h
125 MHz, CDCl3
13C (ppm)

[1]H and [13]C spectra of **3.1a**

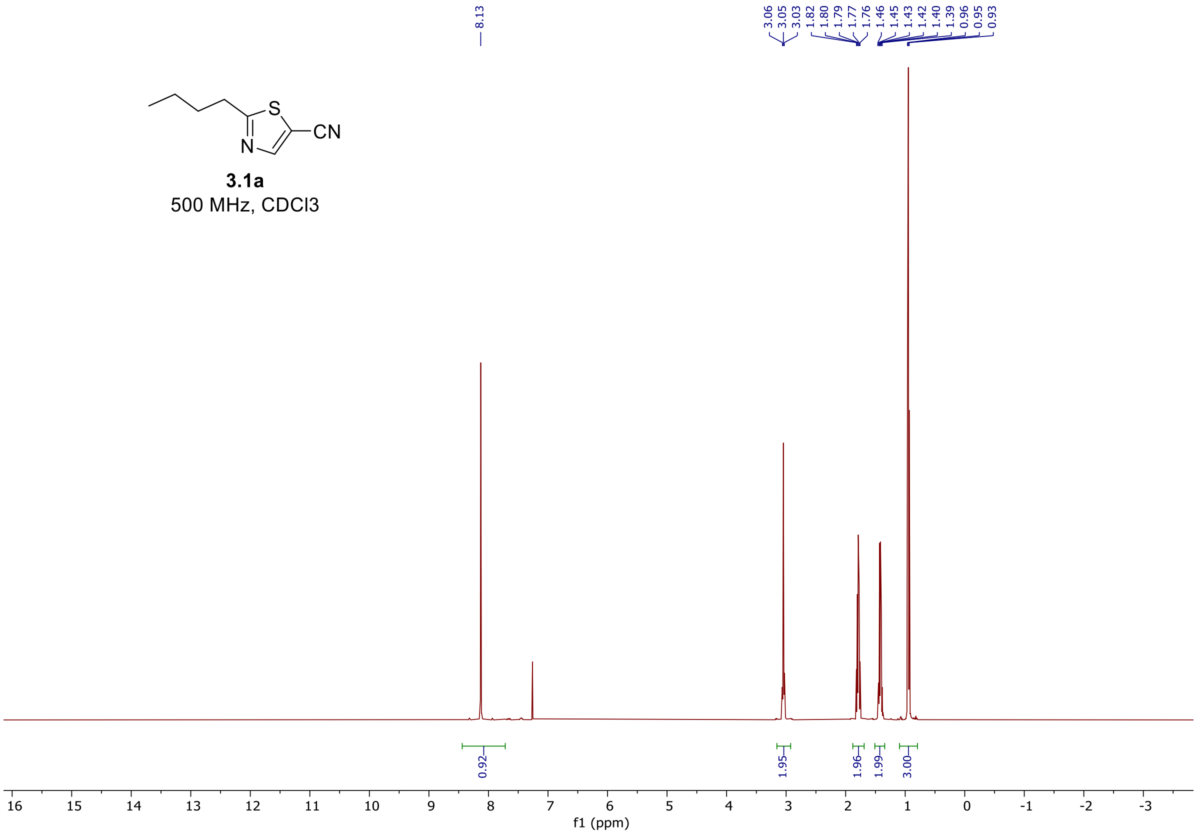


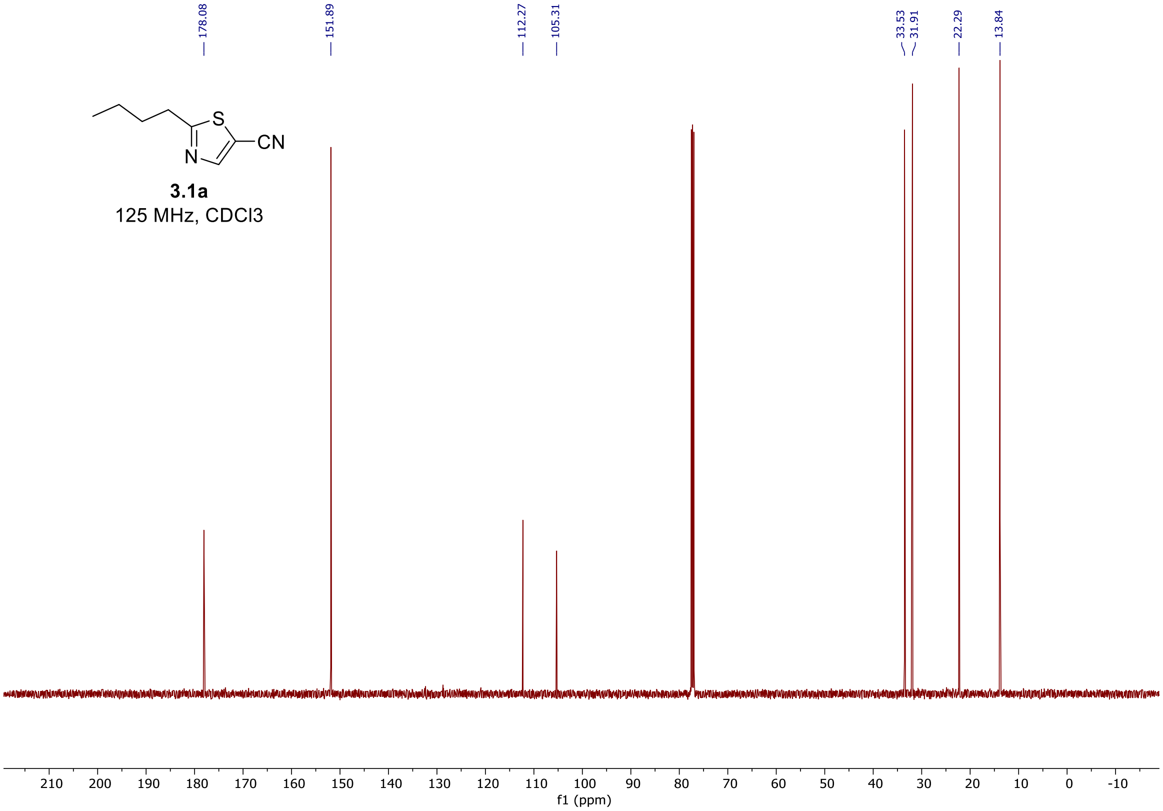

<u>$^{1}$H and $^{13}$C spectra of **3.2a**</u>

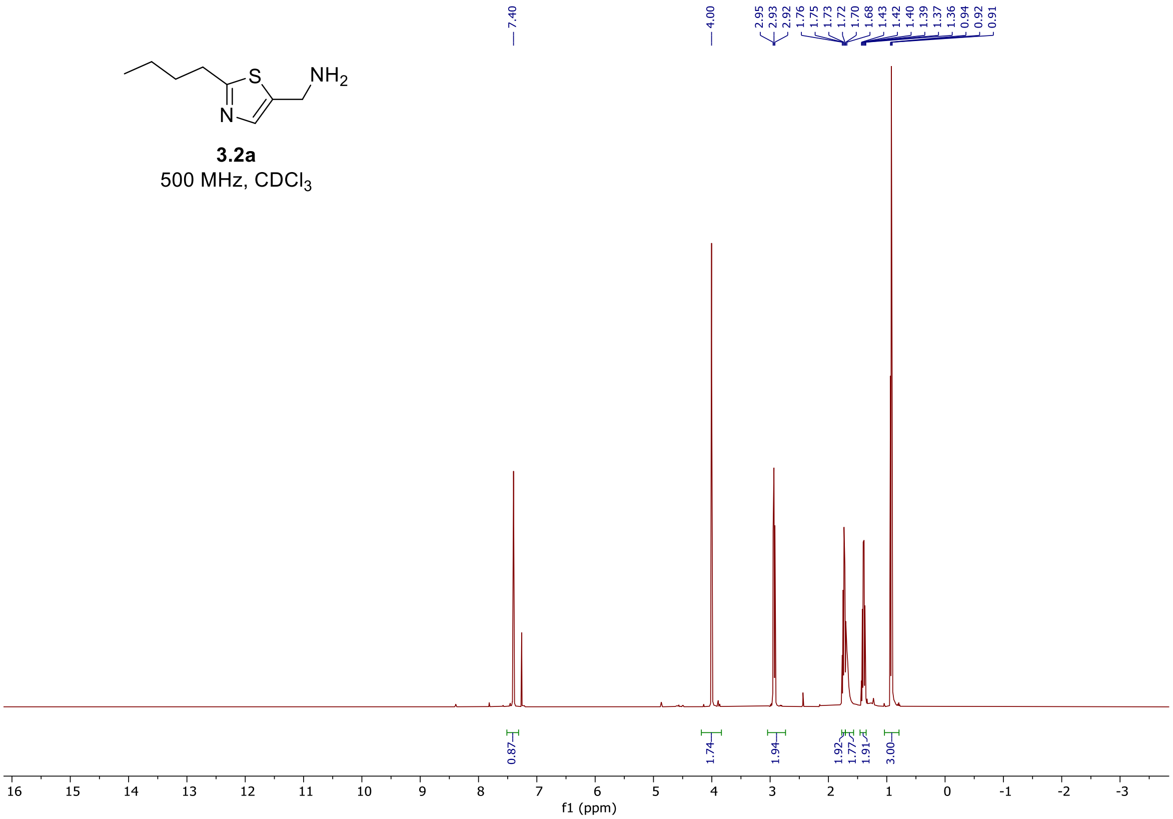


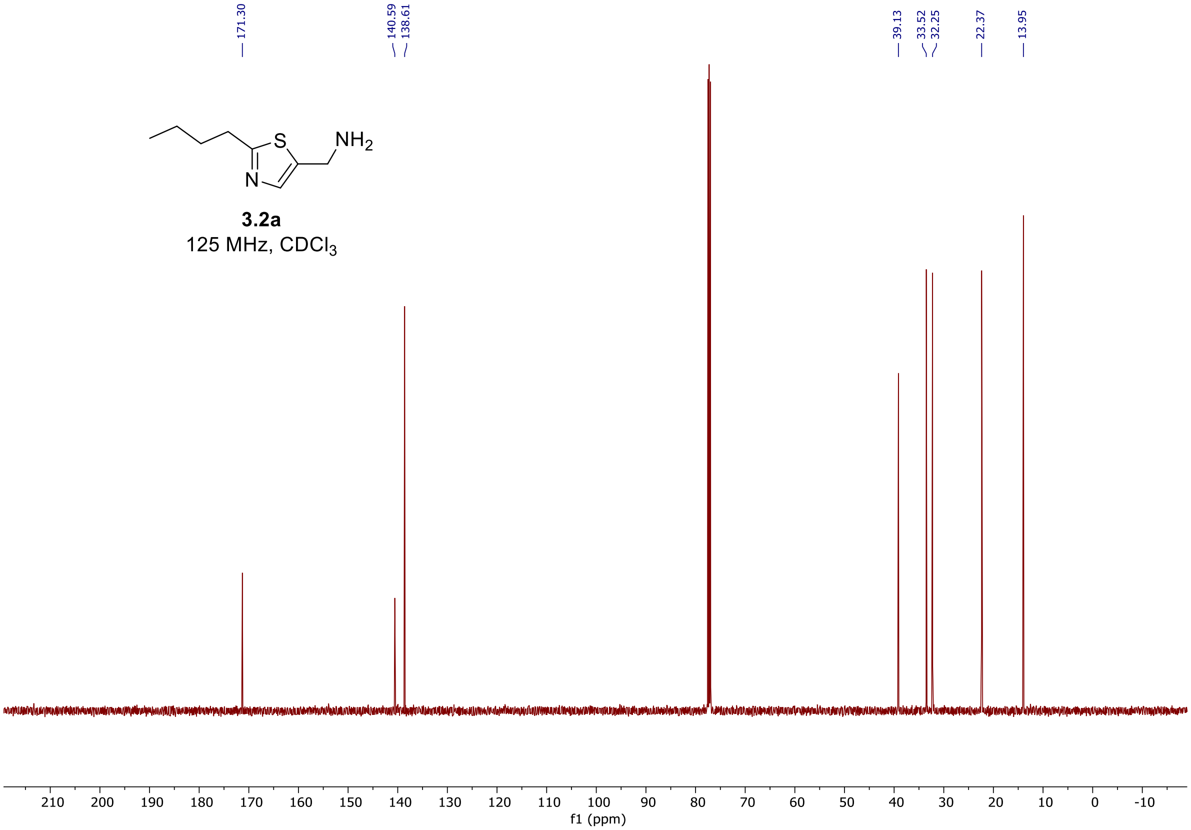

$^{1}$H and $^{13}$C spectra of **3a**

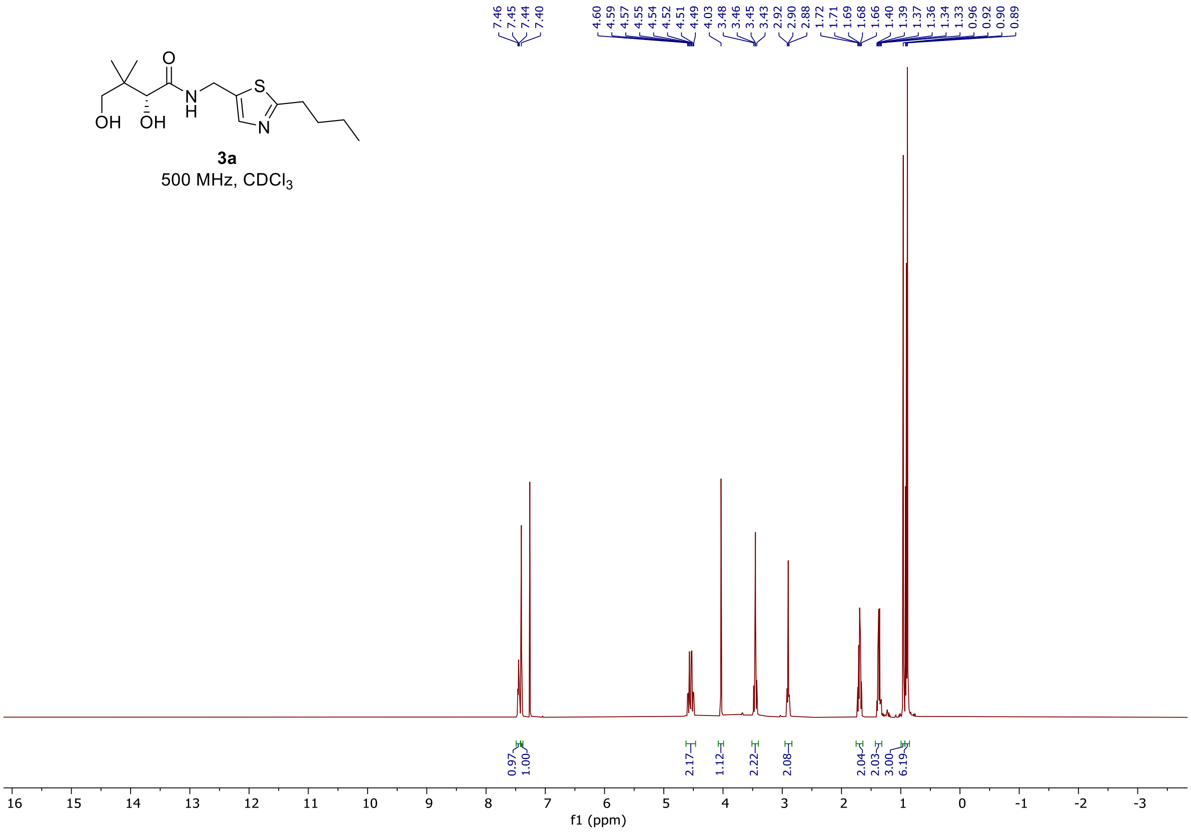


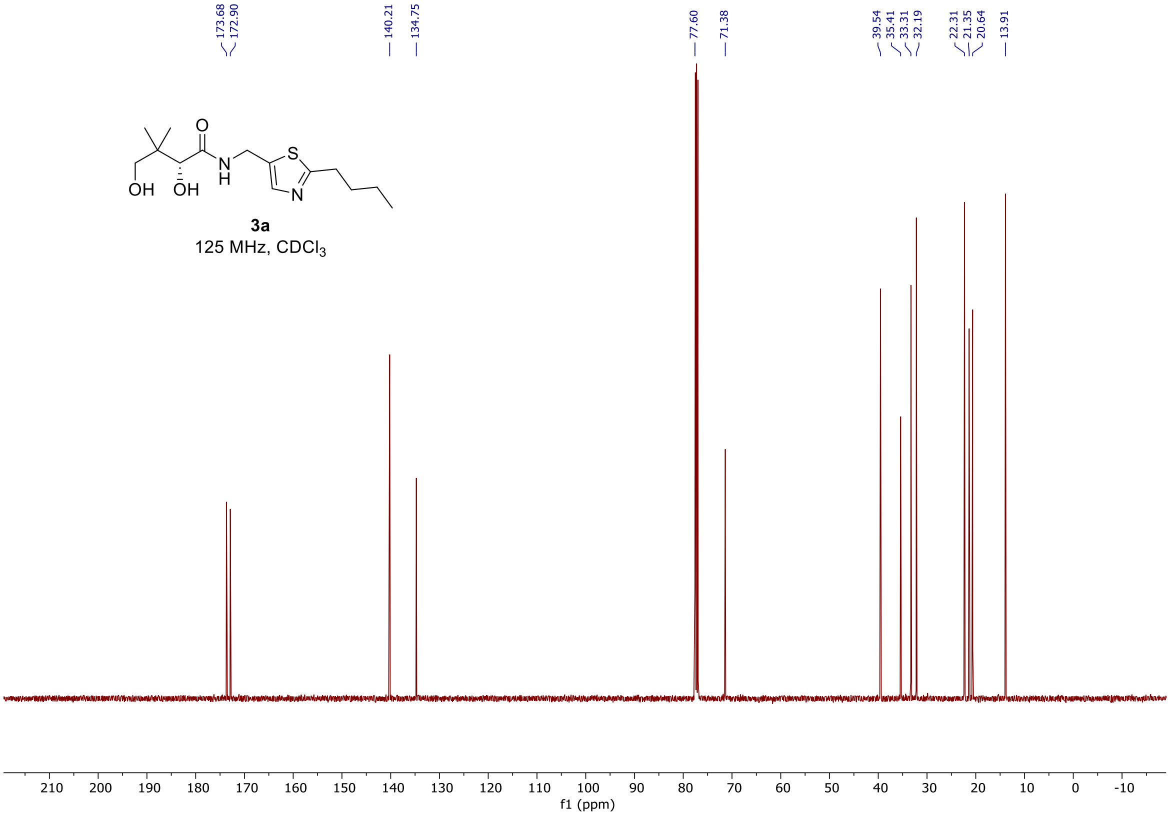

### $^{1}H$ and $^{13}C$ spectra of **3.1b**

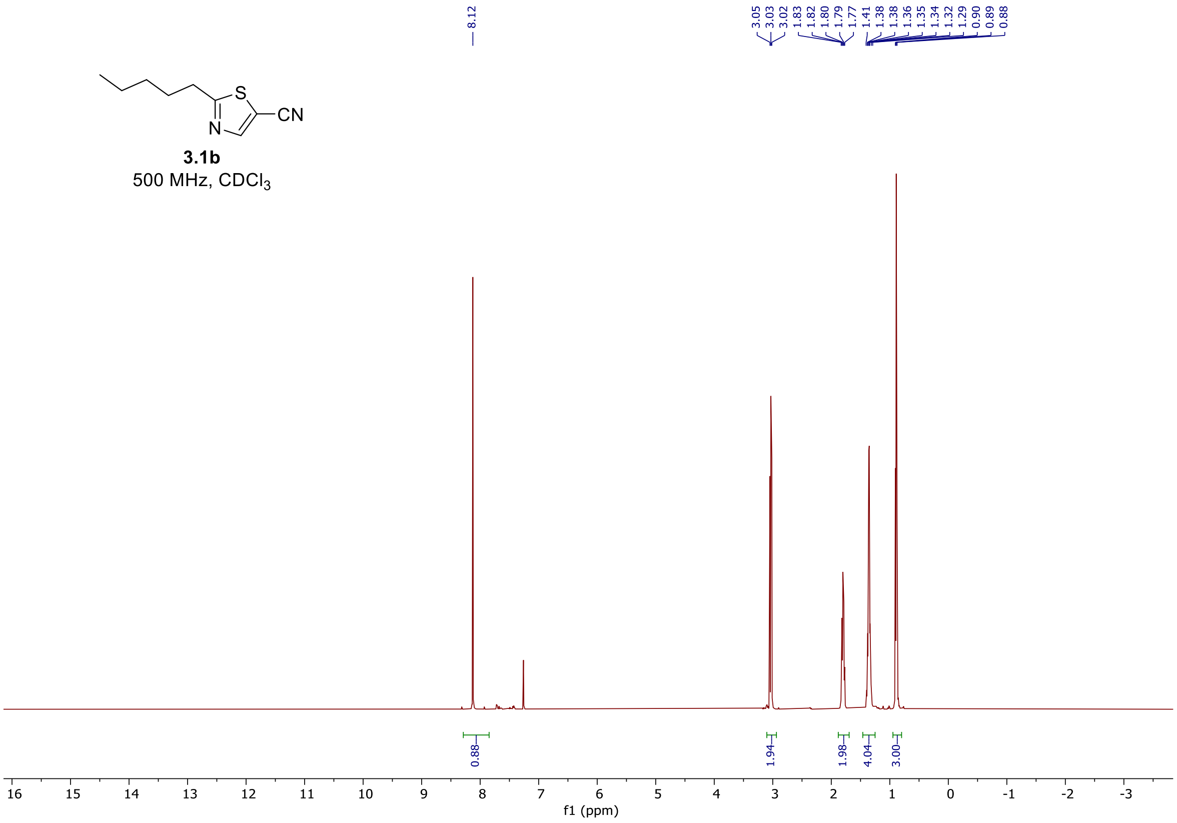


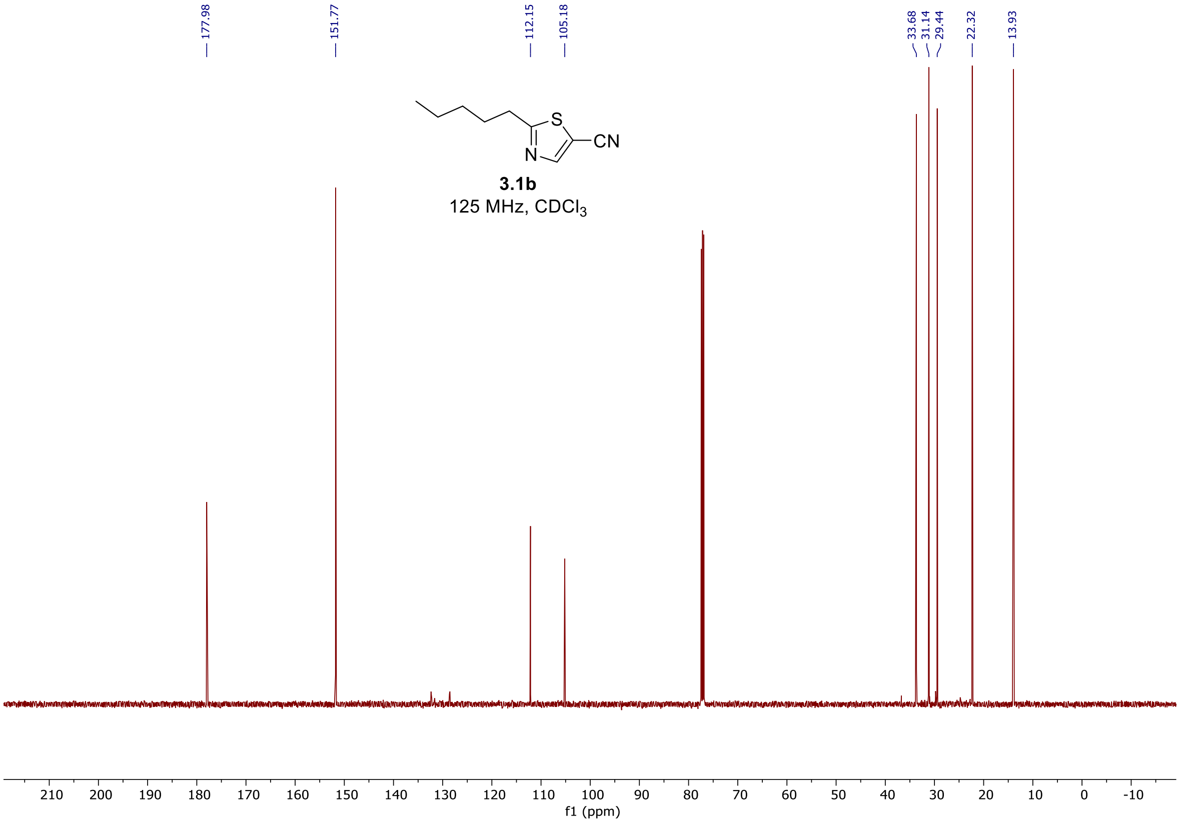

$^{1}H$ and $^{13}C$ spectra of **3.2b**

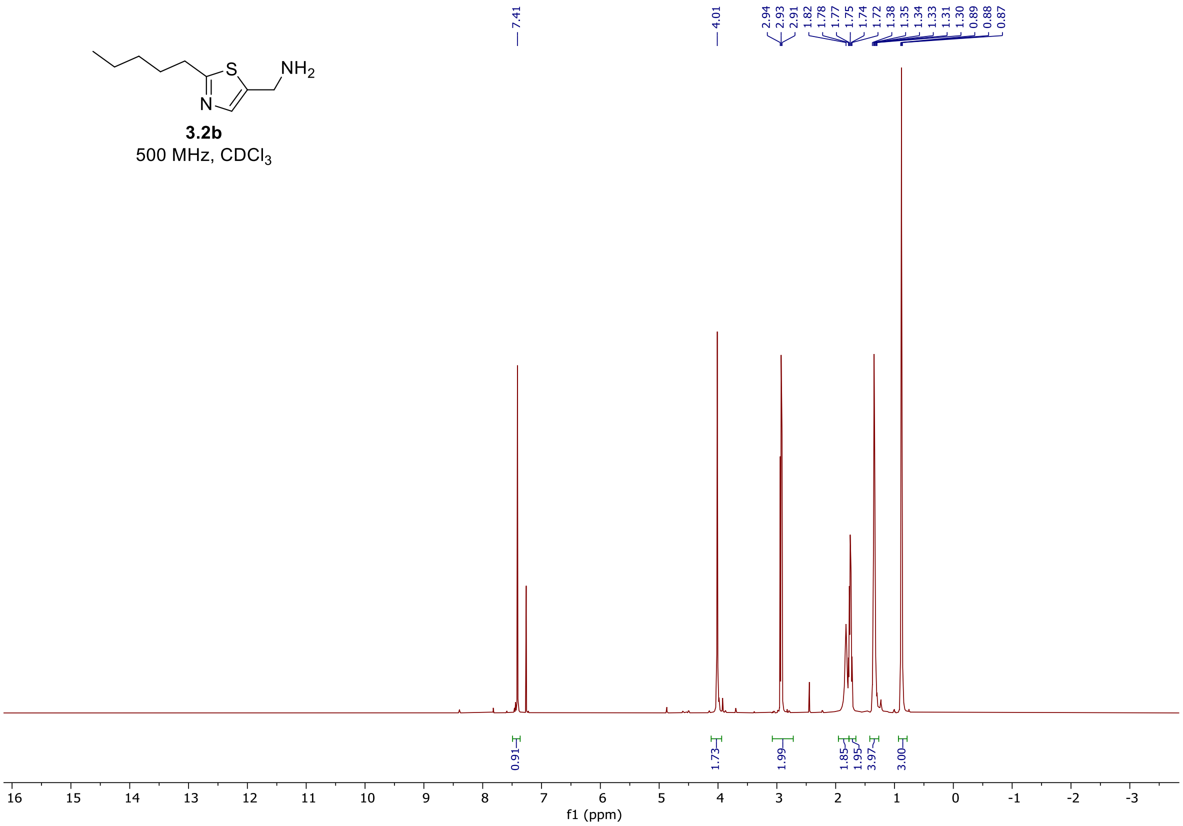


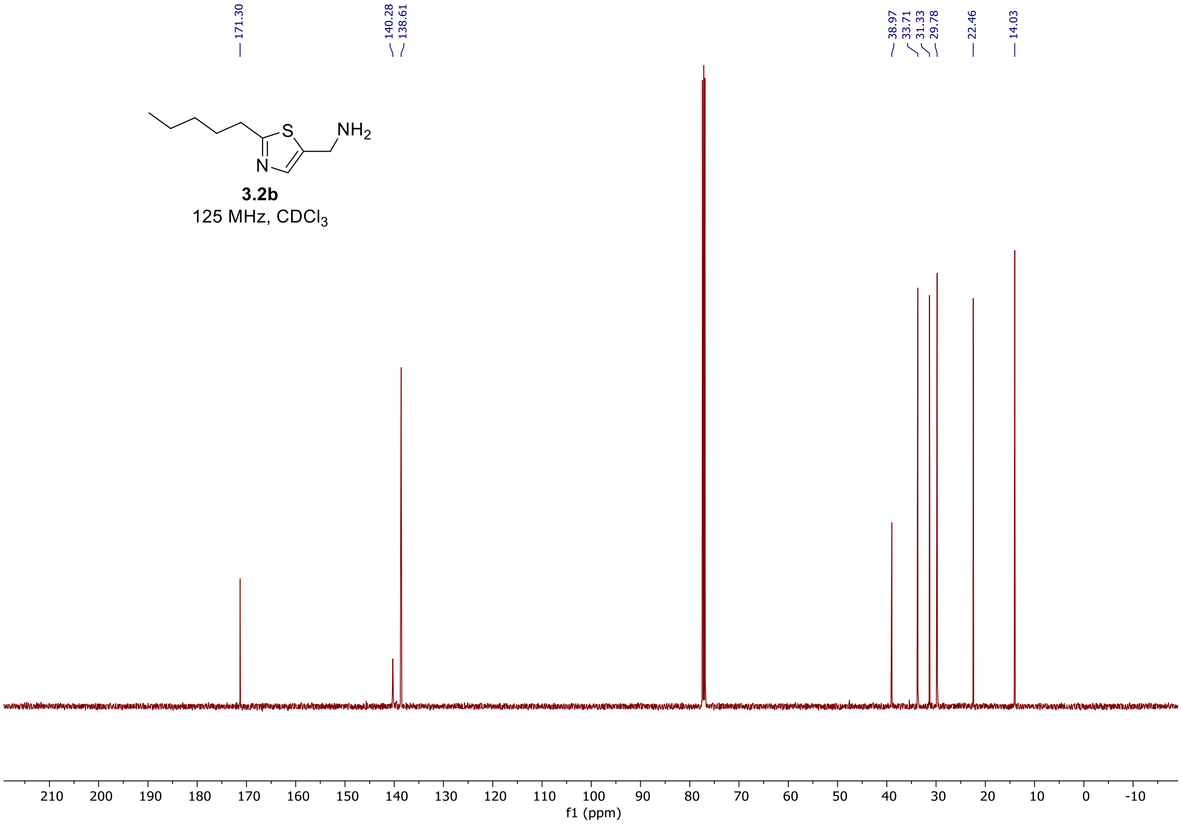

<sup></sup>

#### $^{1}$H and $^{13}$C spectra of **3b**

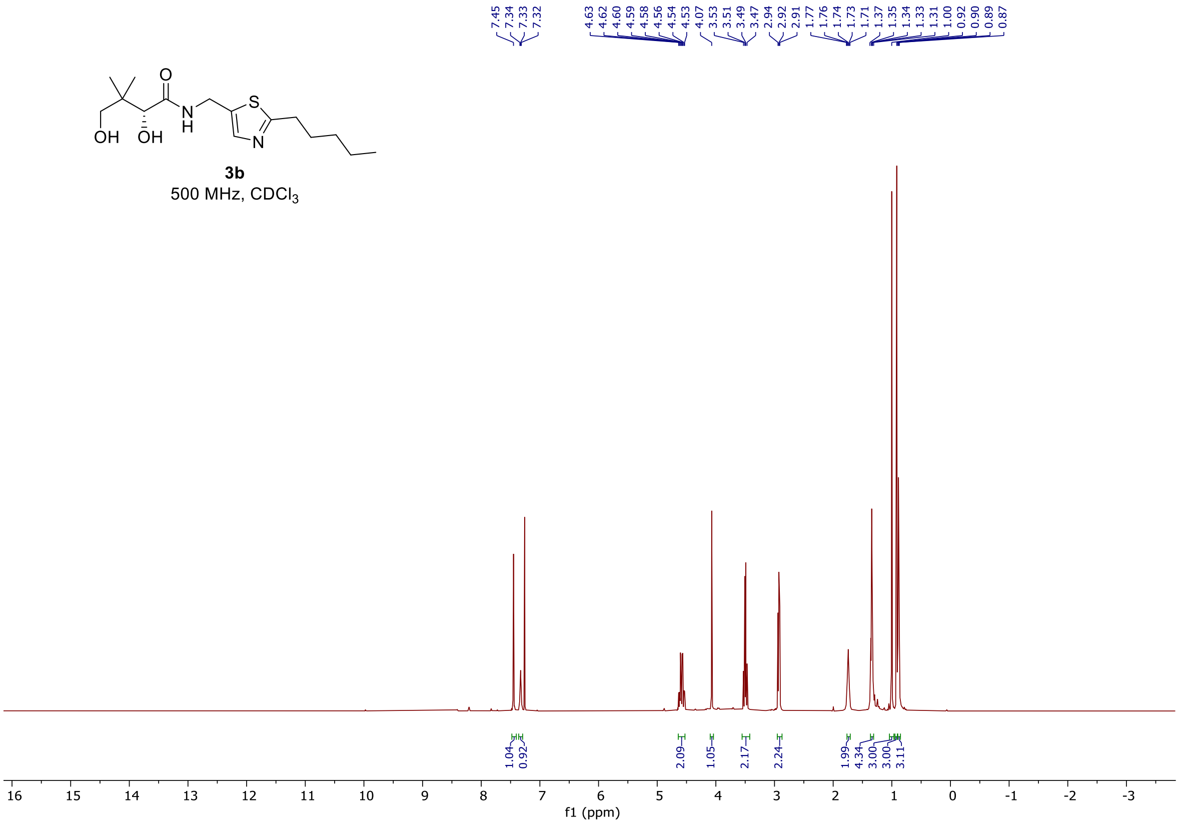


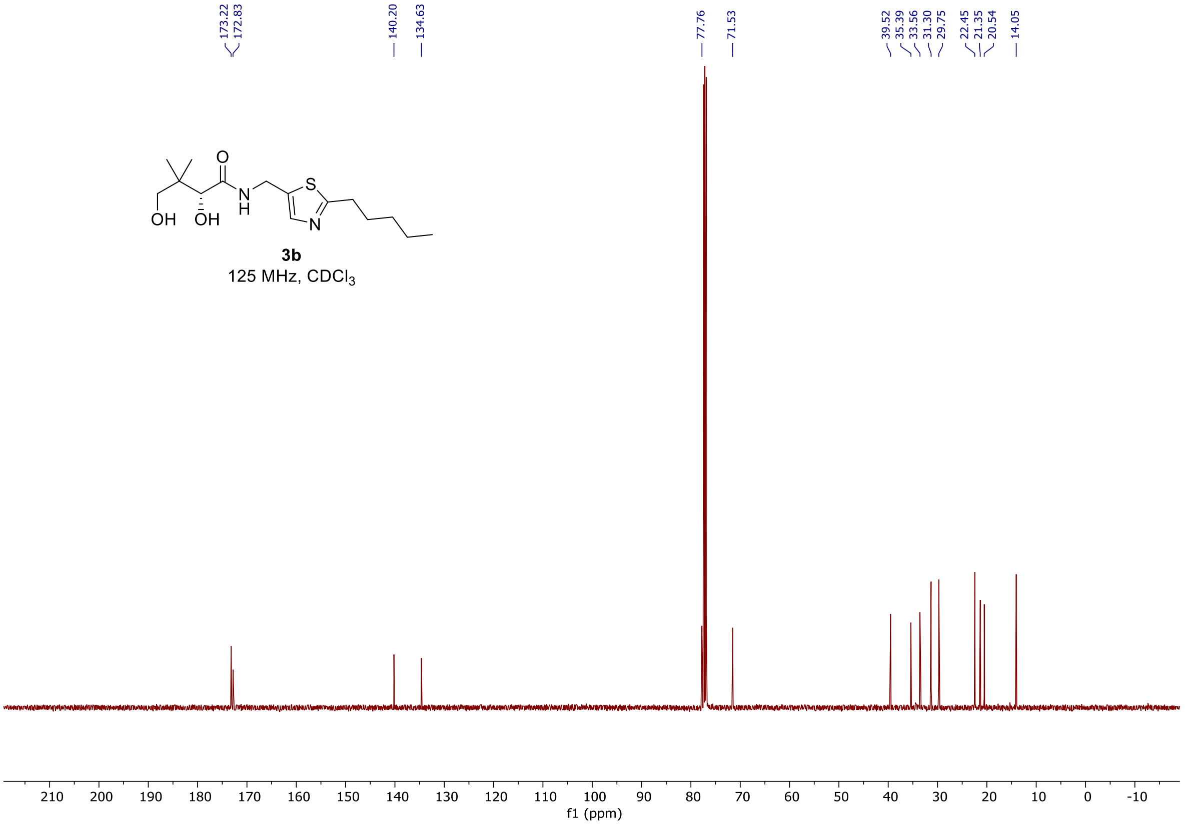

$^{1}H$ and $^{13}C$ spectra of **3.1c**

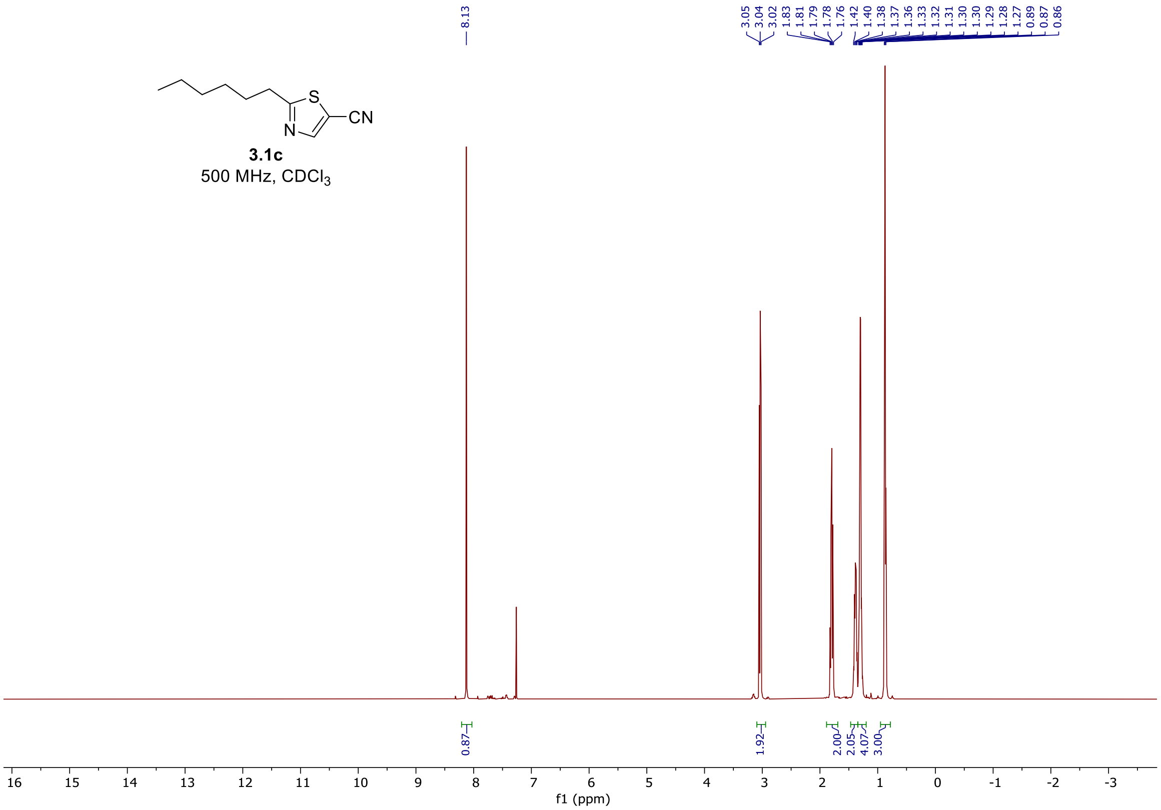


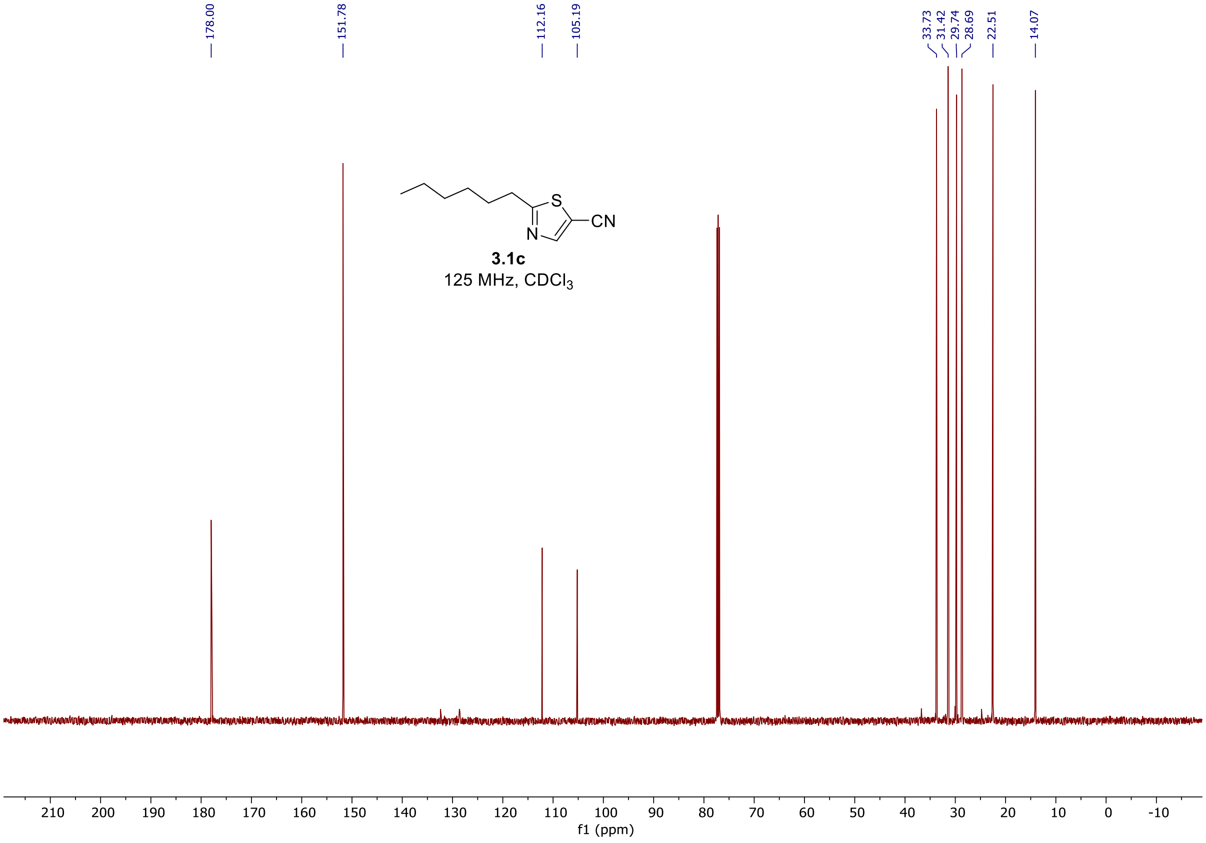

<sup></sup>

$^{1}$H and $^{13}$C spectra of **3.2c**

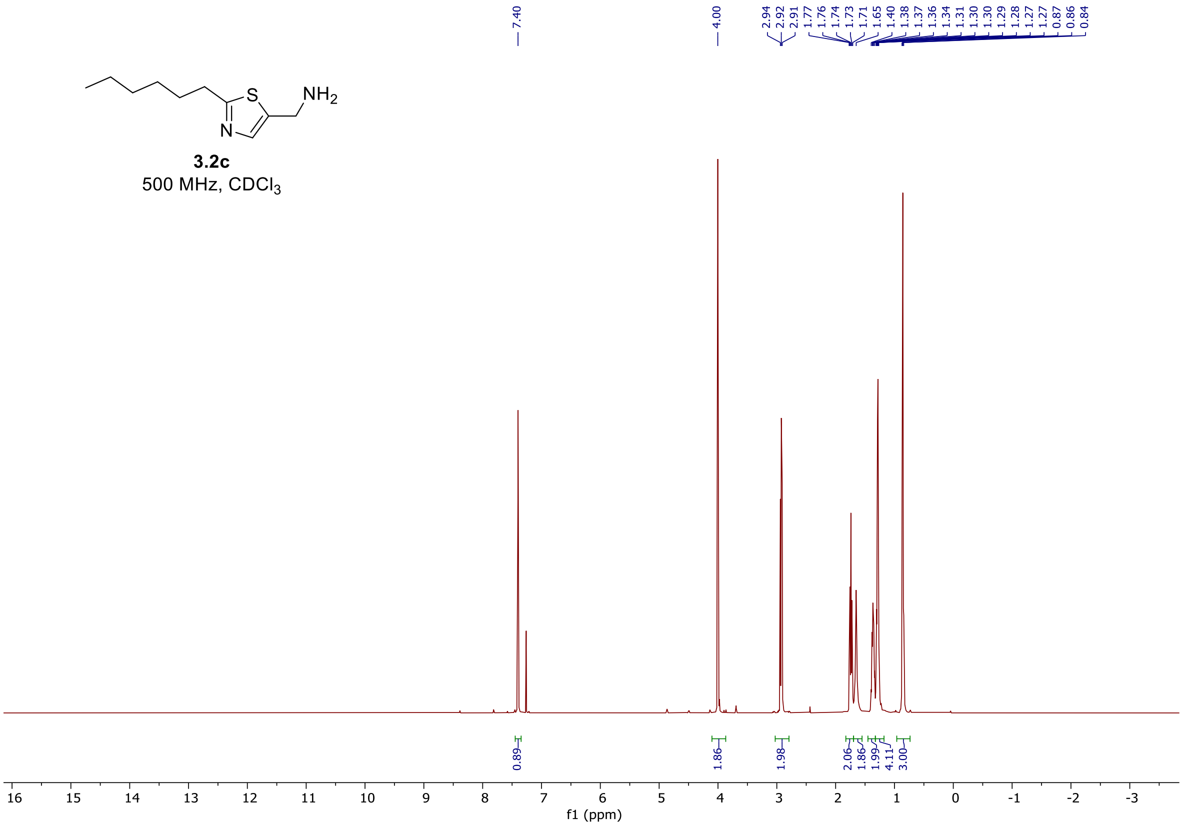


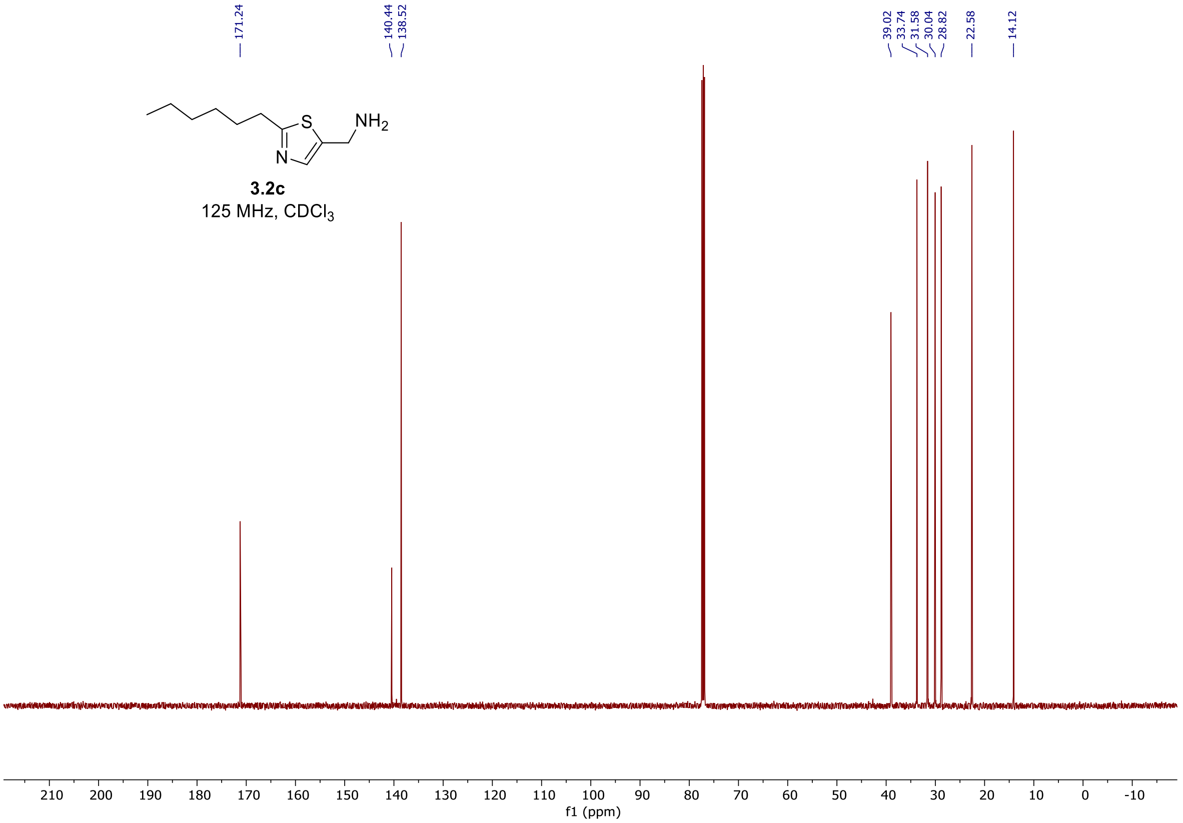

[1]H and [13]C spectra of **3c**

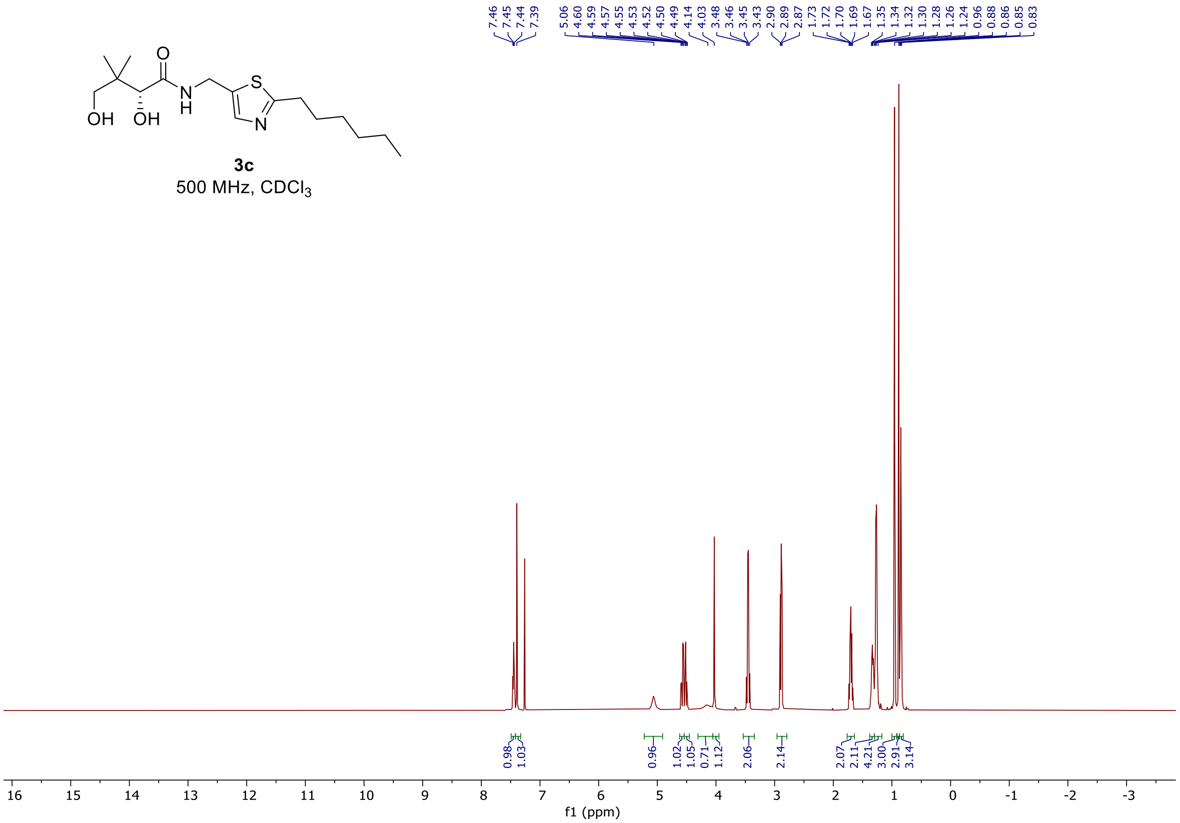


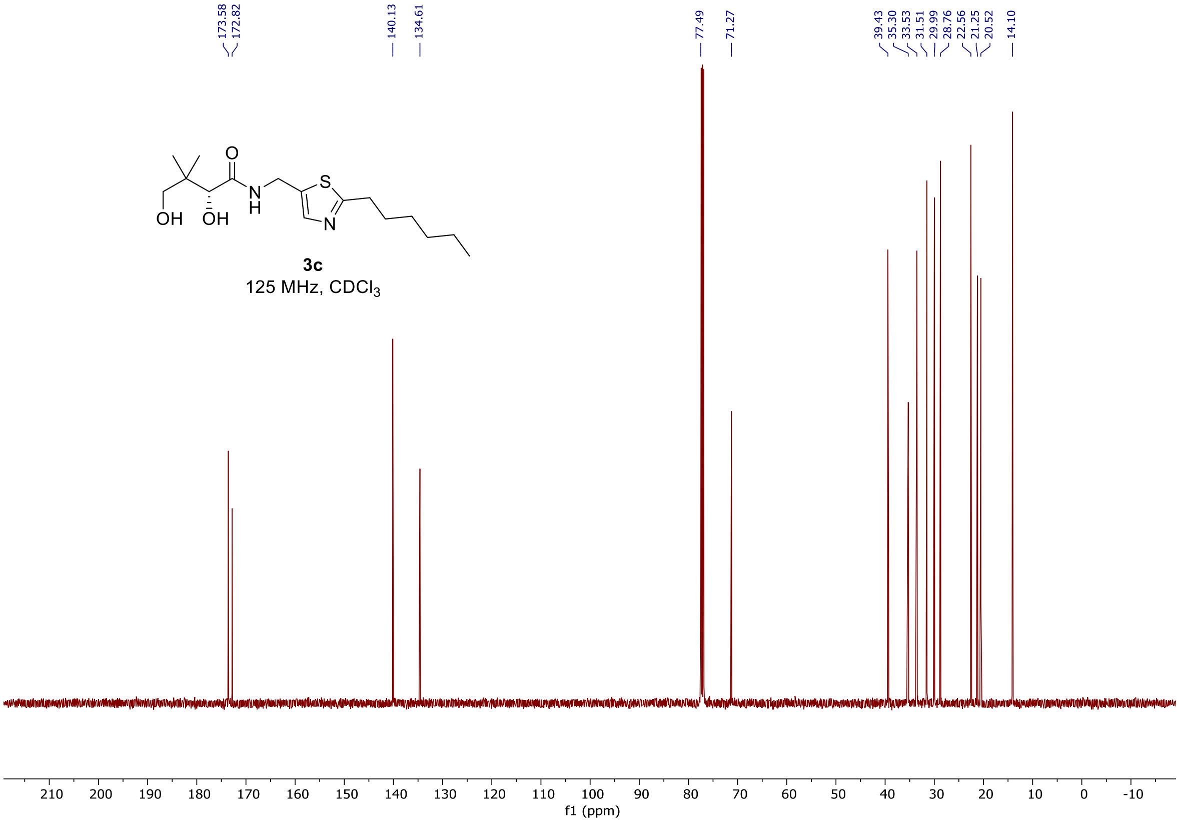

[1]H and [13]C spectra of **3.1d**

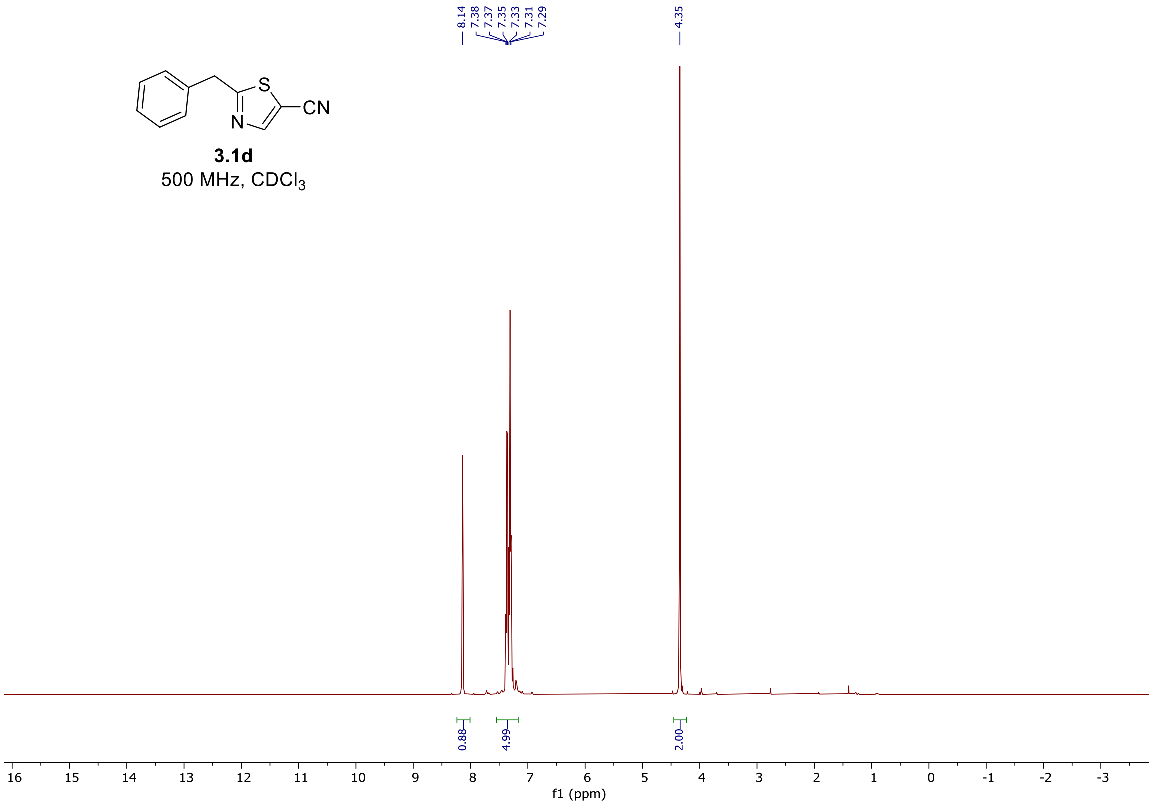


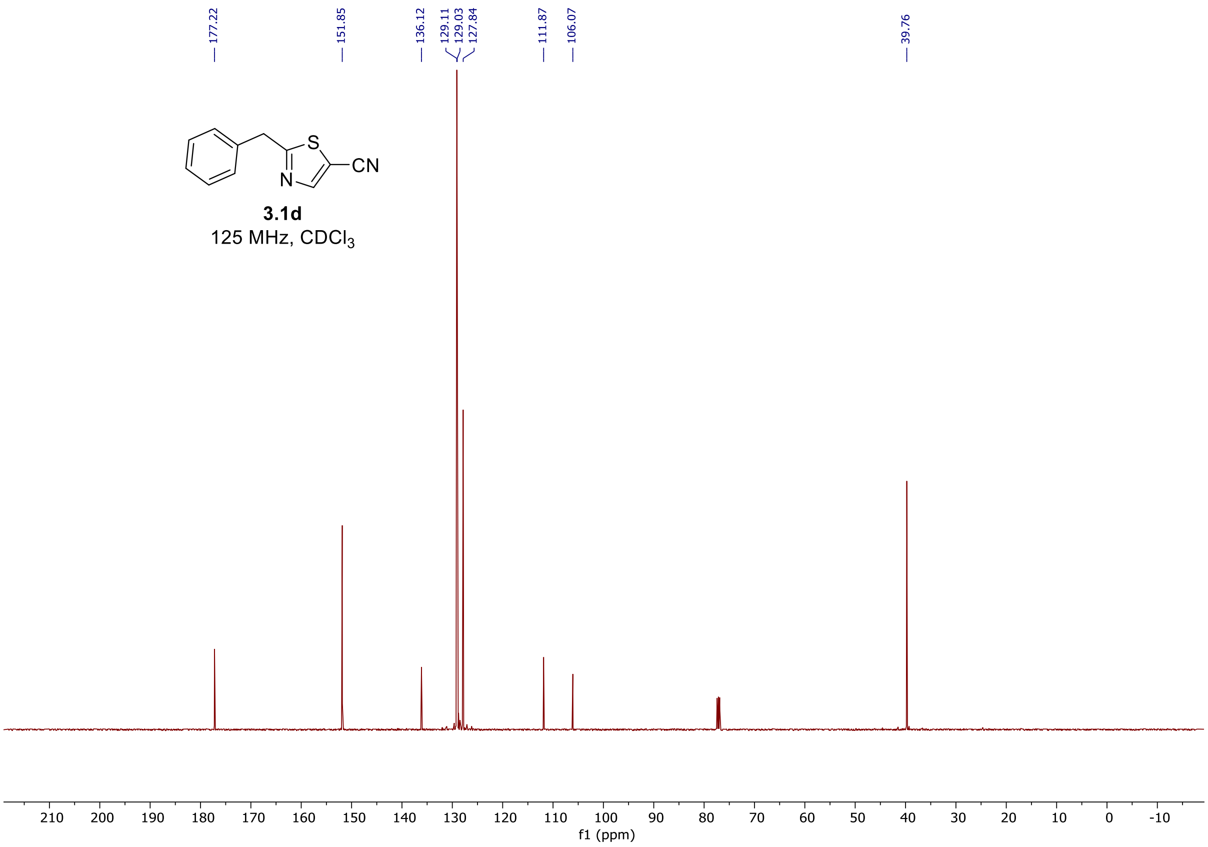

$^{1}H$ and $^{13}C$ spectra of **3.2d**

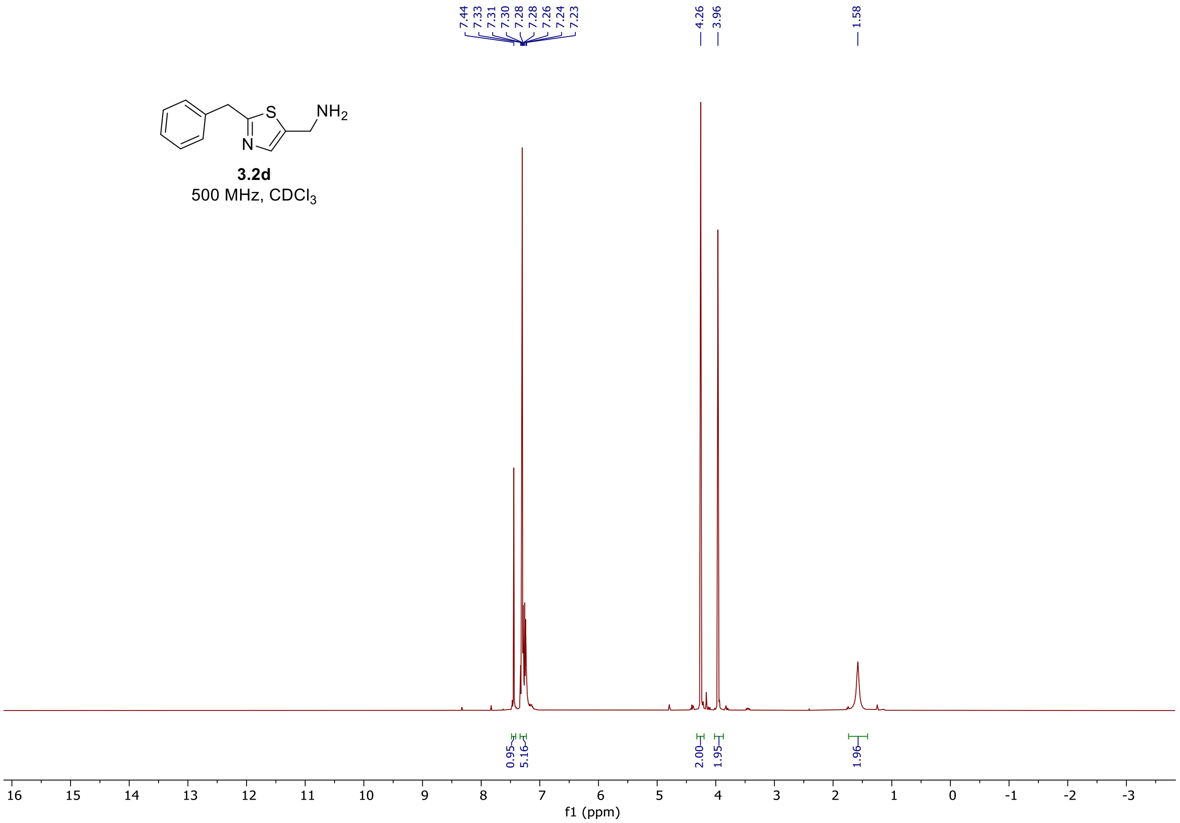


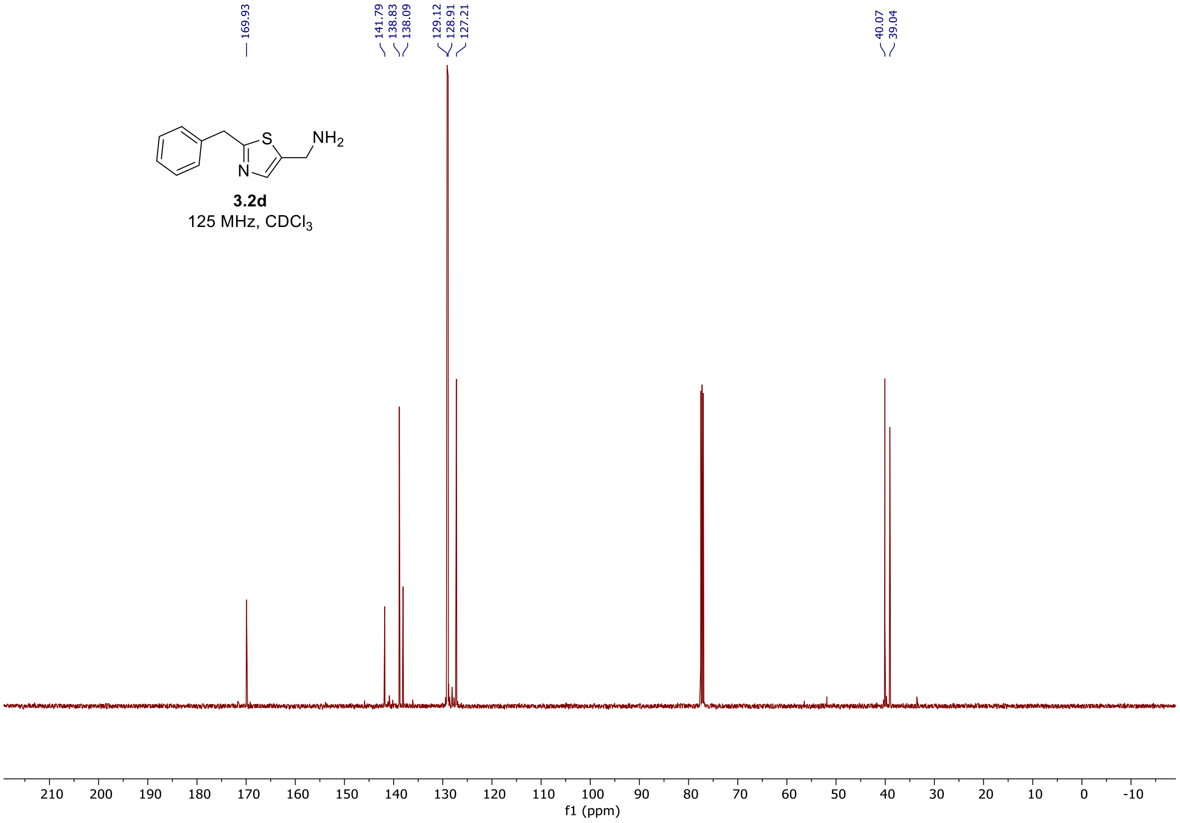

#### $^1$H and $^{13}$C spectra of **3d**

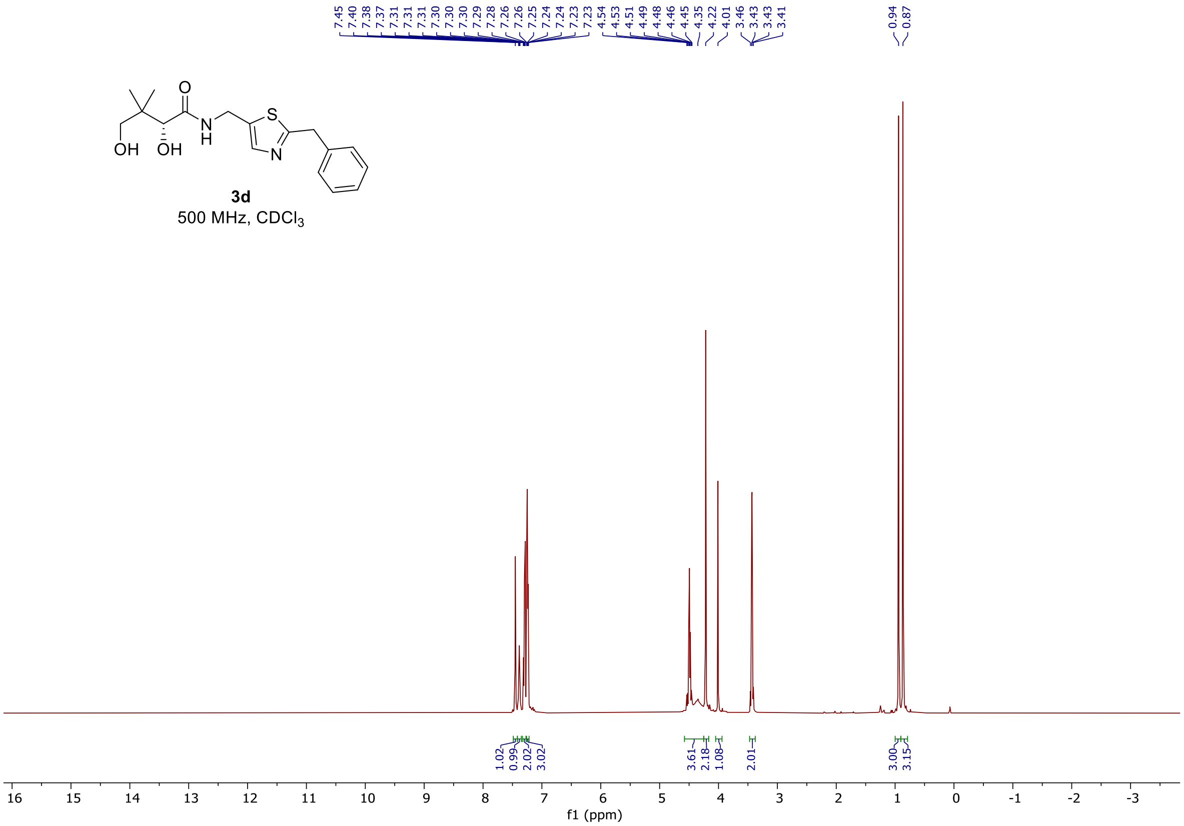


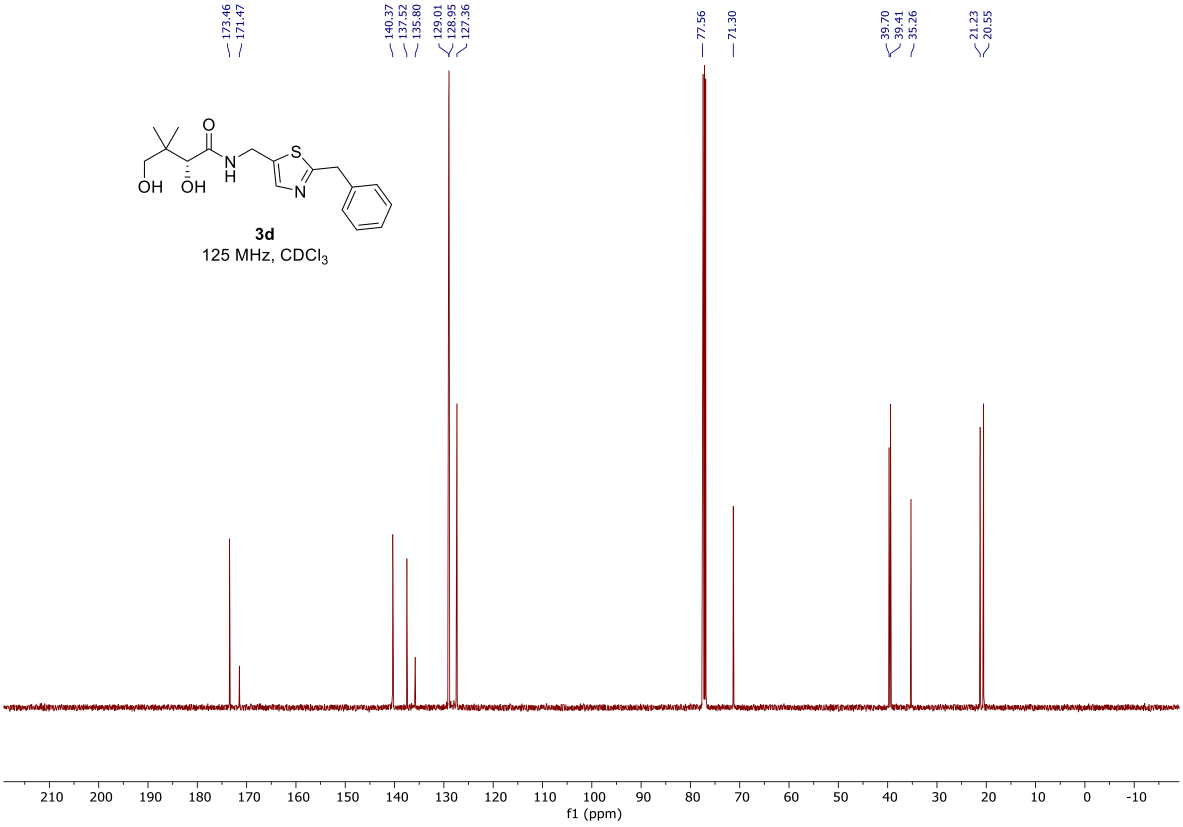

$^{1}H$ and $^{13}C$ spectra of **3.1e**

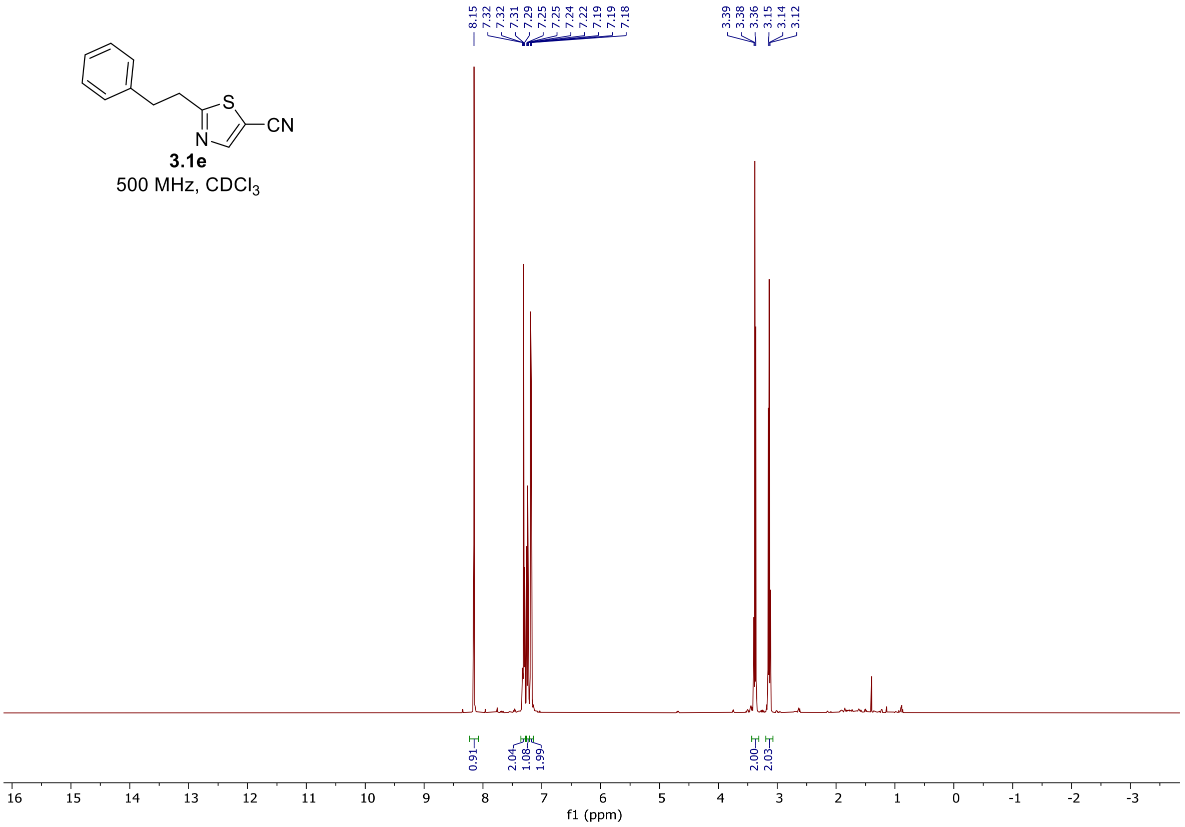


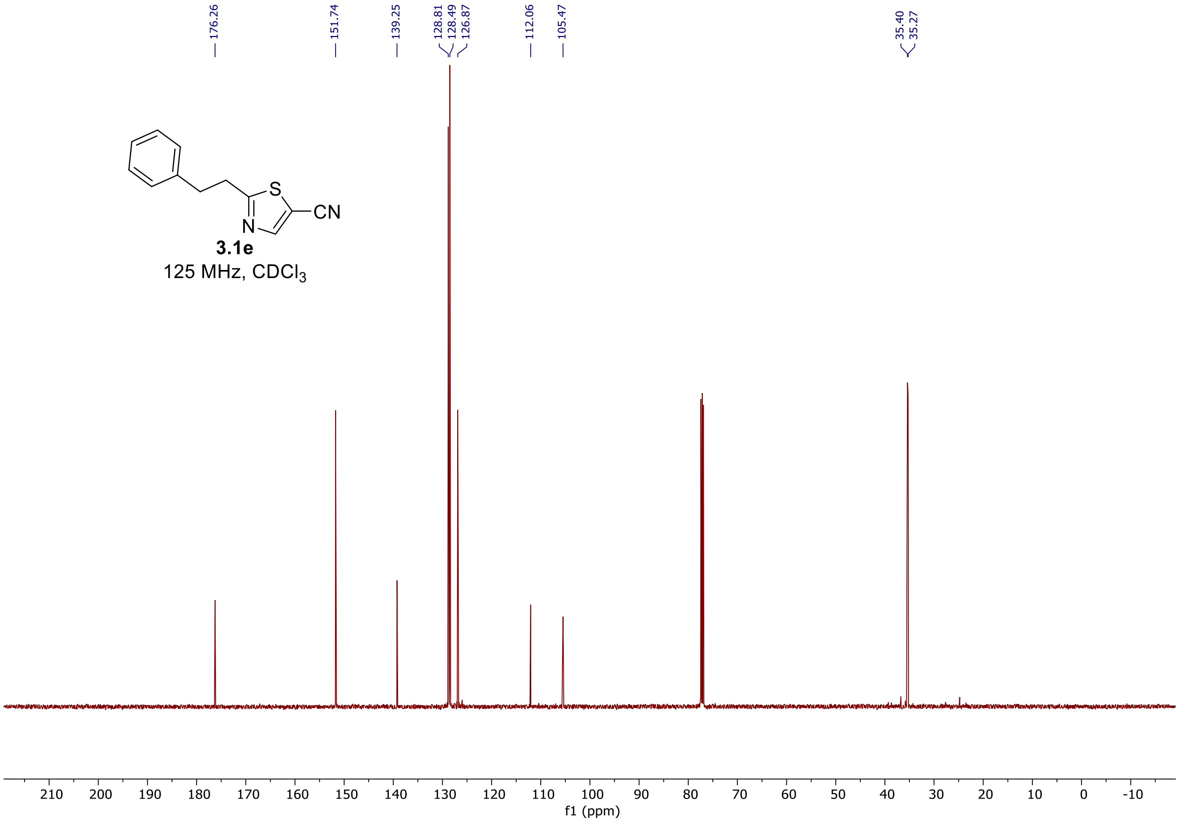

$^{1}H$ and $^{13}C$ spectra of **3.2e**

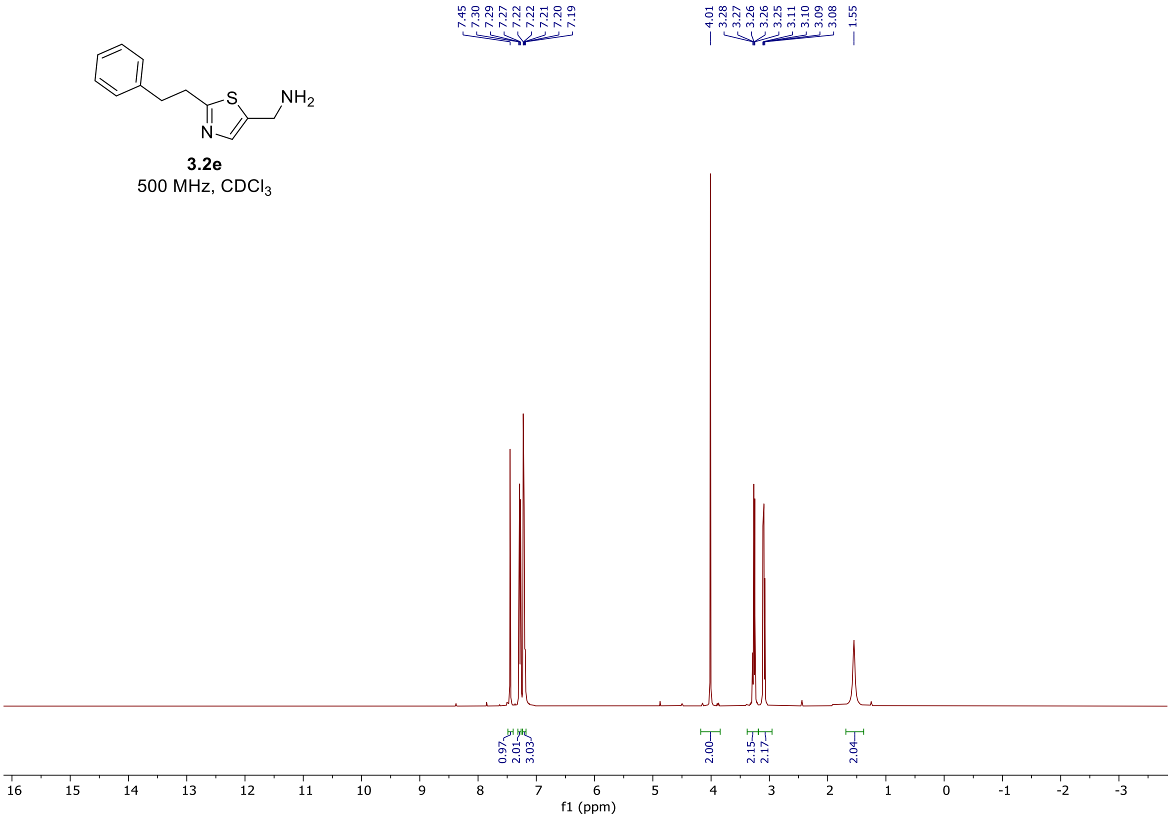


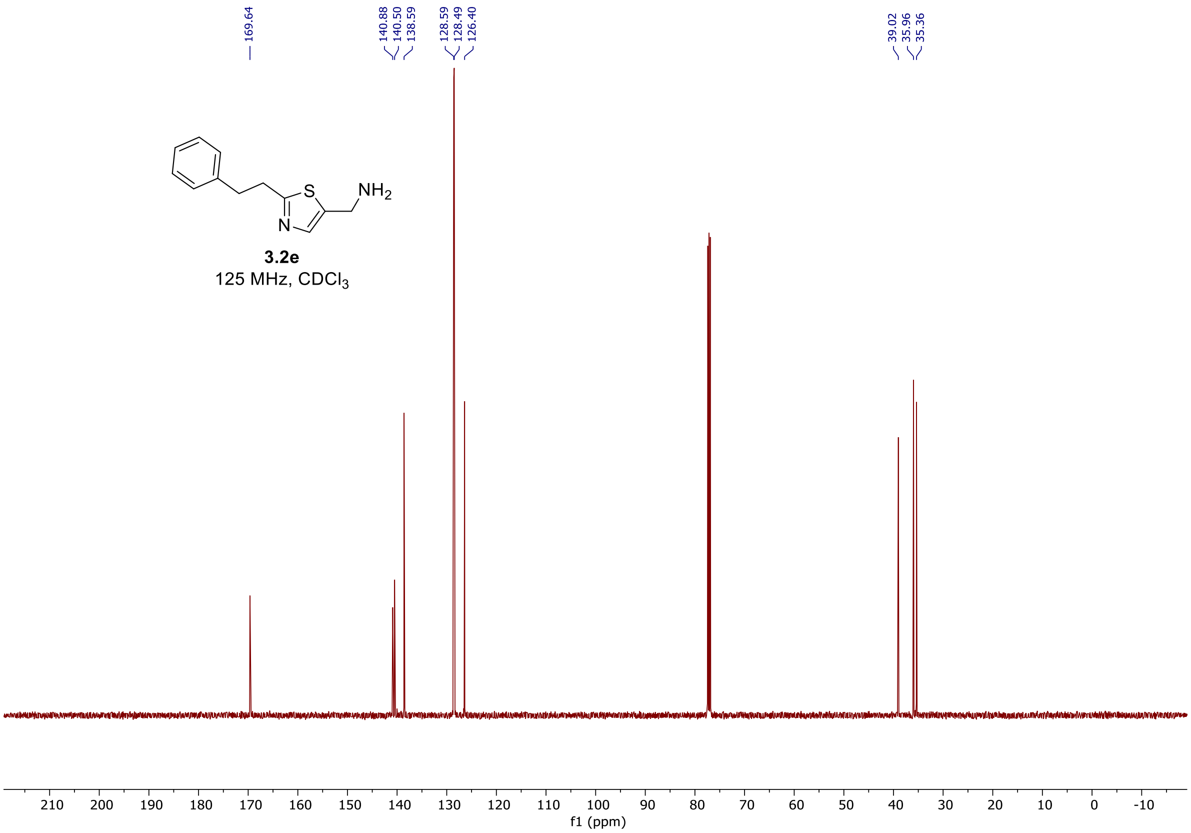

### $^{1}H$ and $^{13}C$ spectra of **3e**

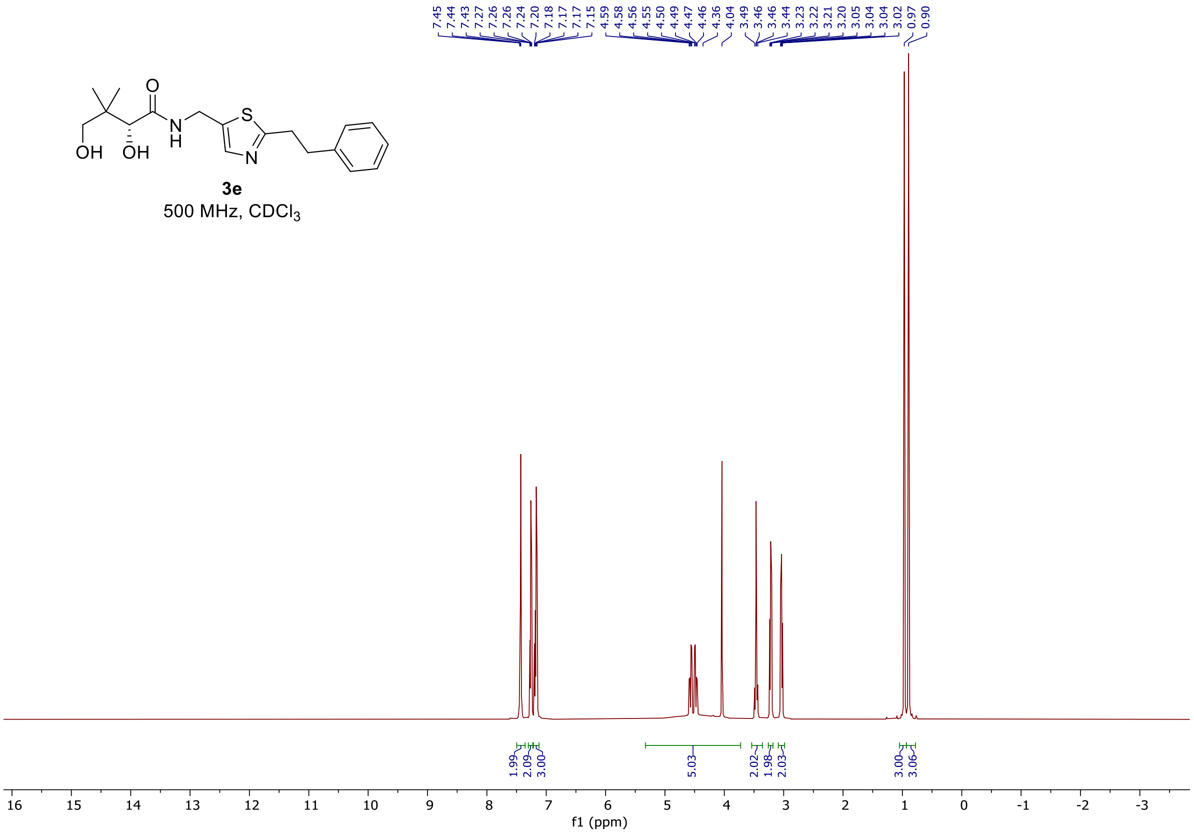


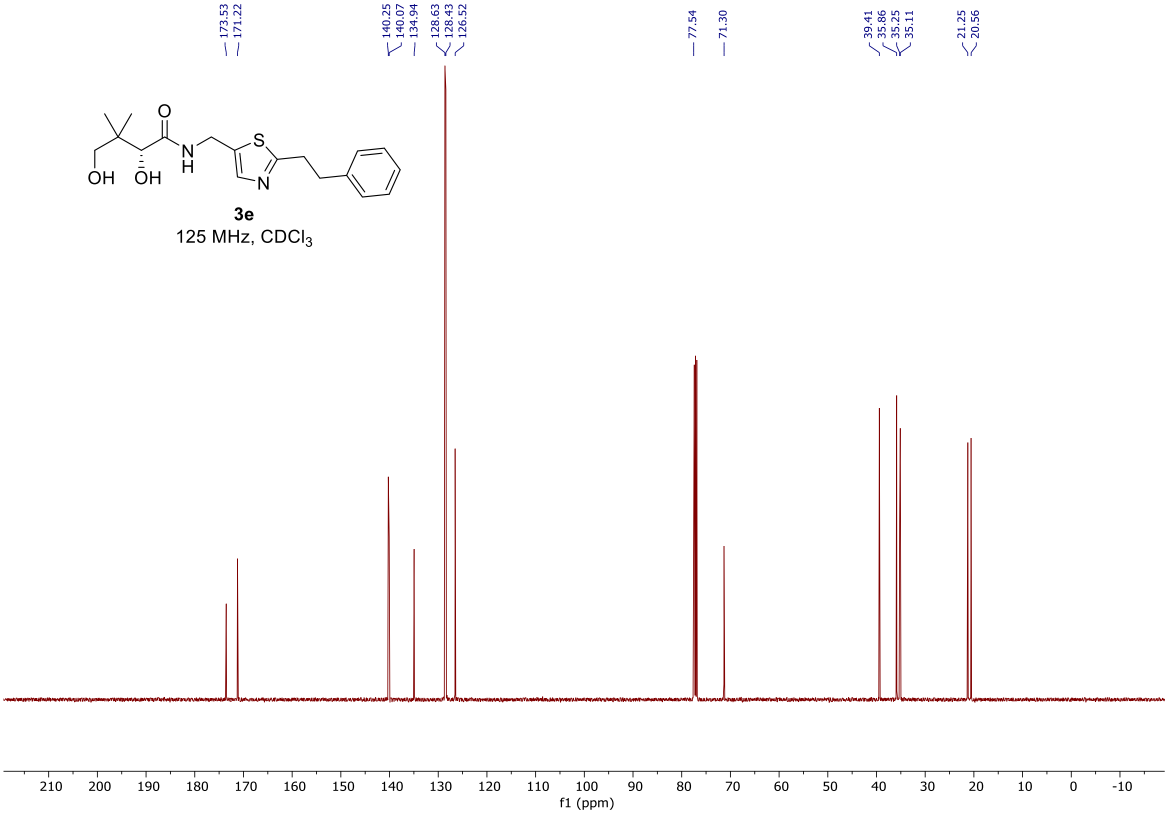

$^{1}H$ and $^{13}C$ spectra of **3.6f**

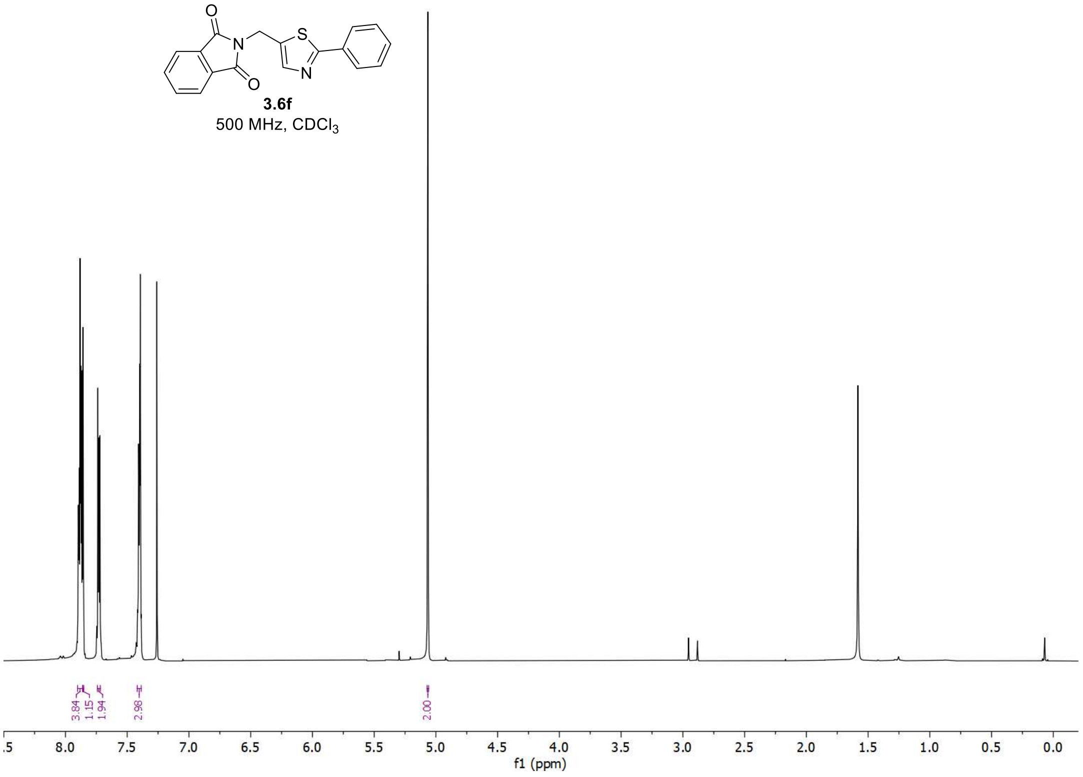


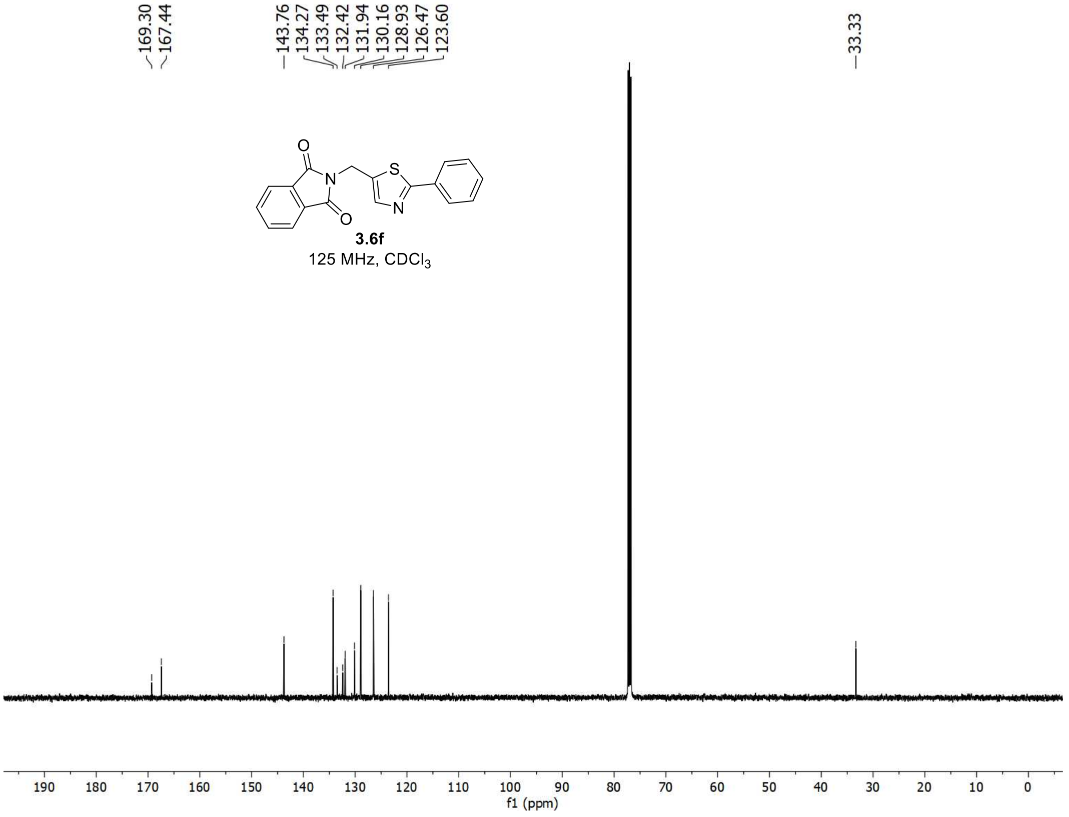

$^{1}H$ and $^{13}C$ spectra of **3.7f**

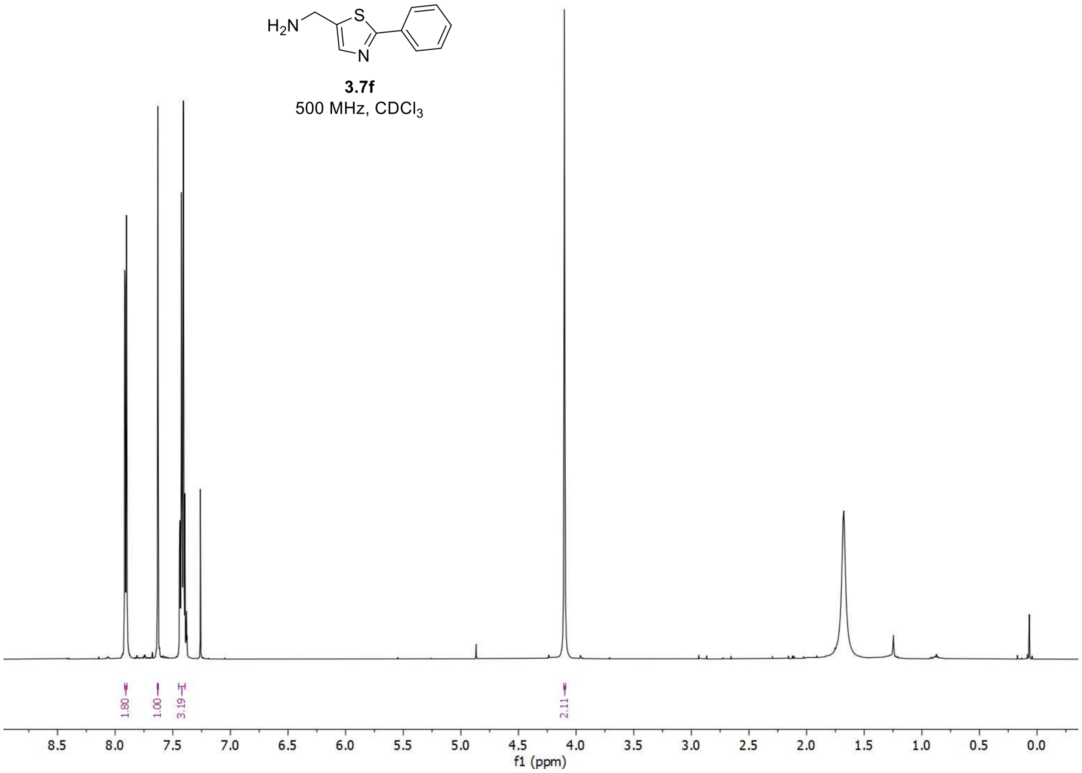


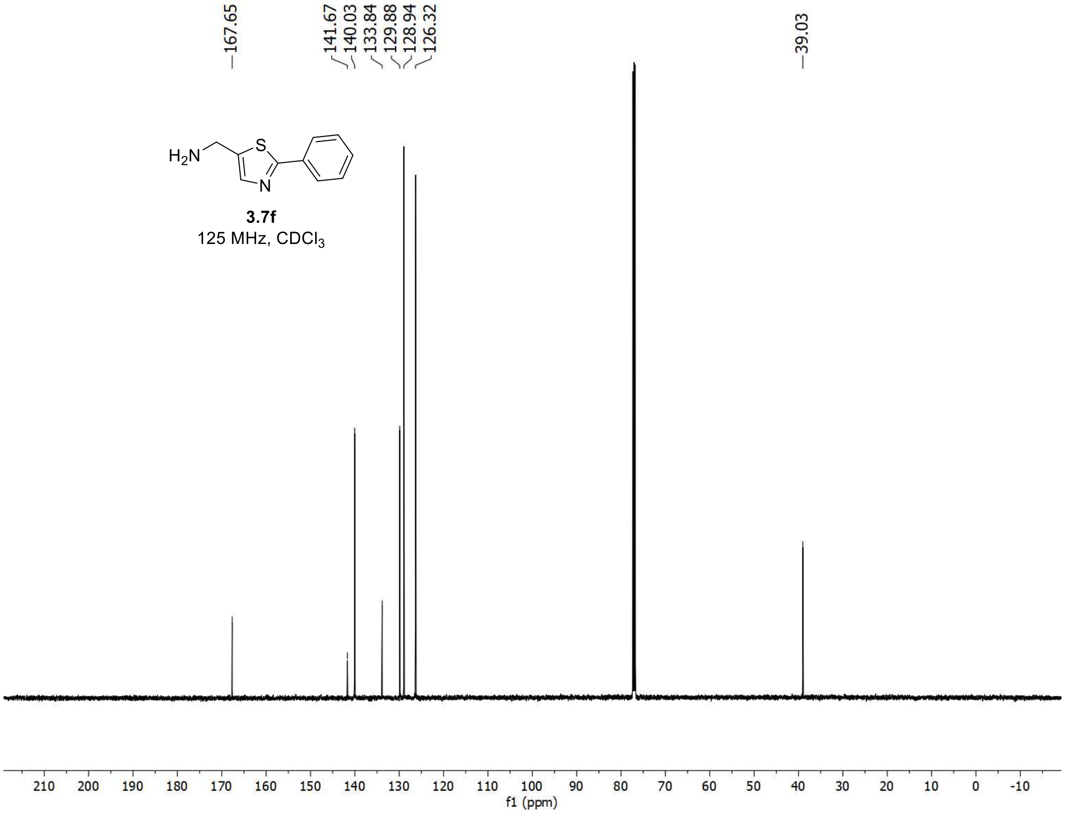

$^{1}H$ and $^{13}C$ spectra of **3f**

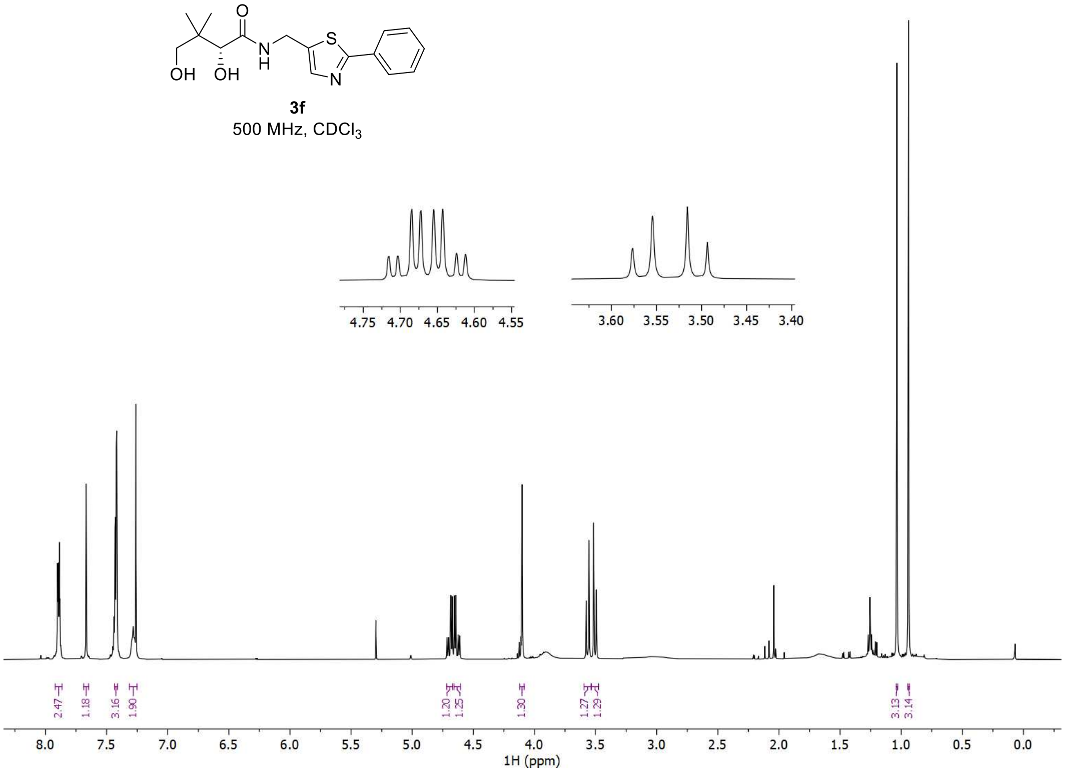


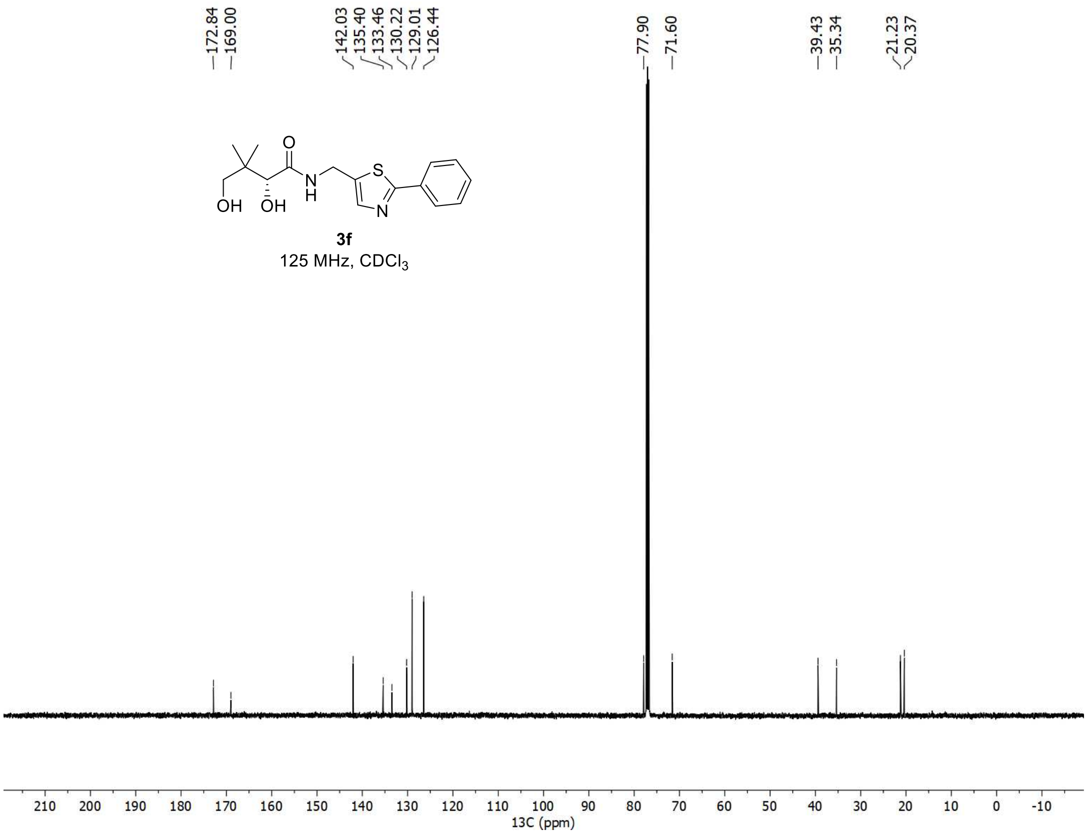

<sup></sup>

$^{1}H$, $^{19}F$, and $^{13}C$ spectra of **3.4g**

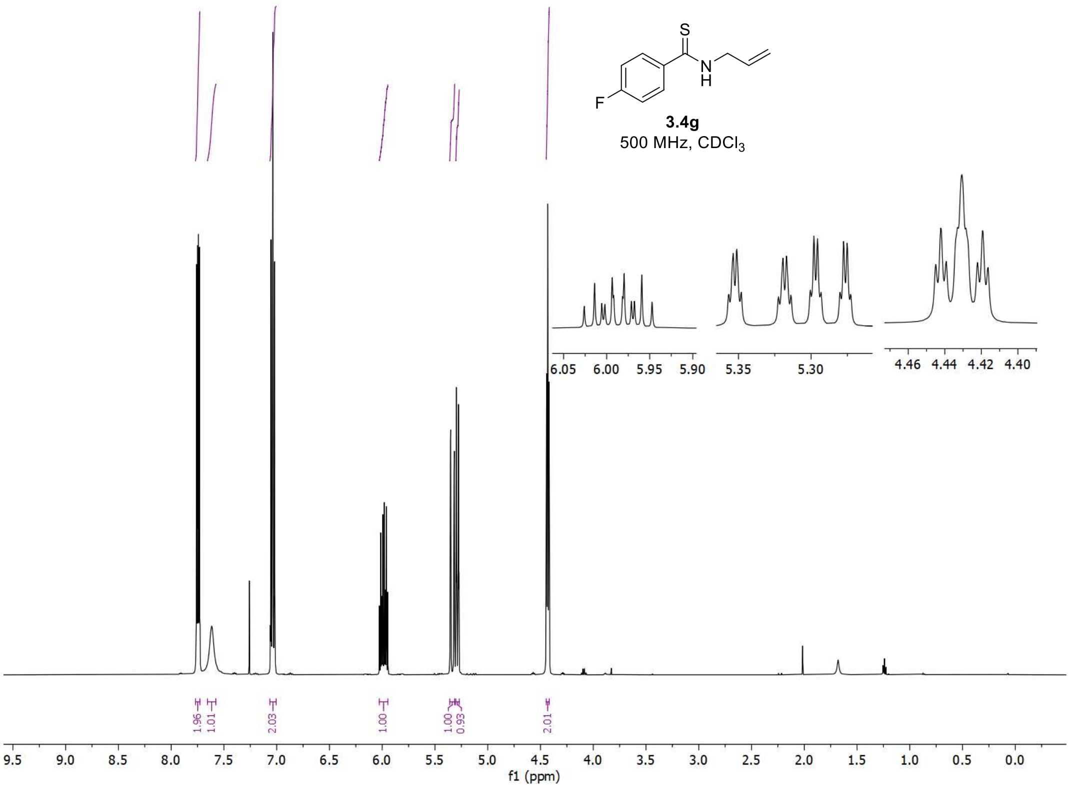


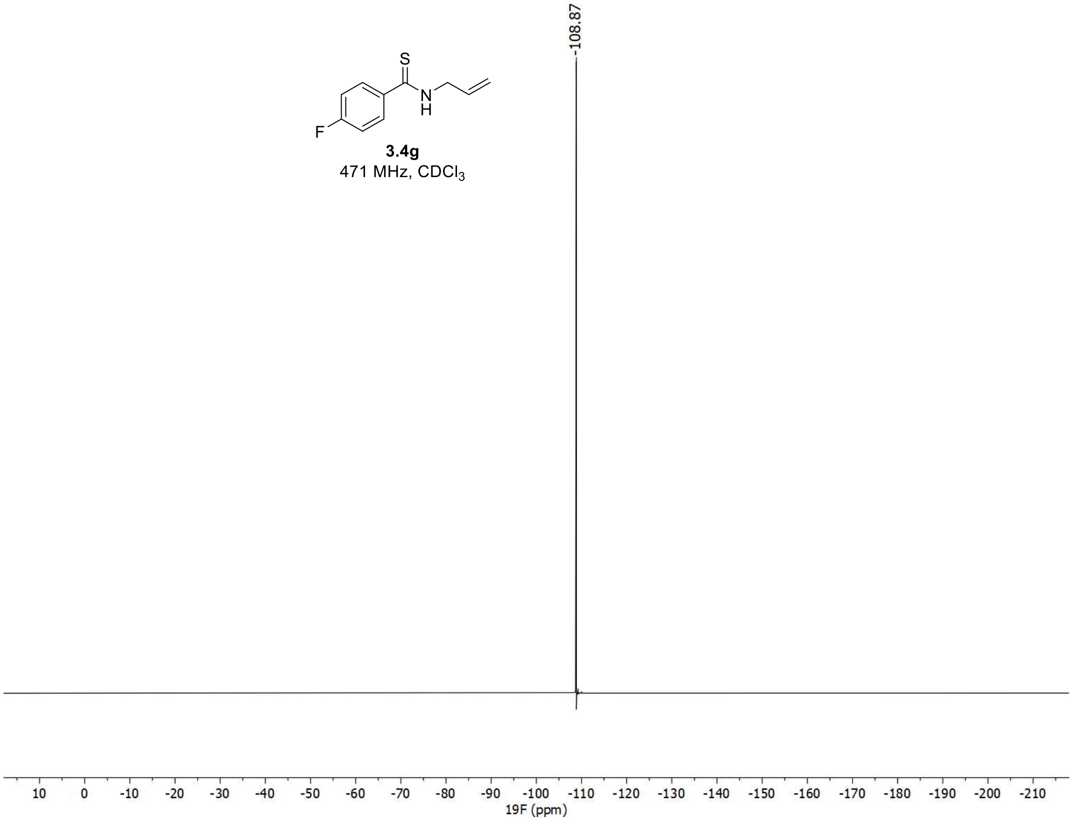

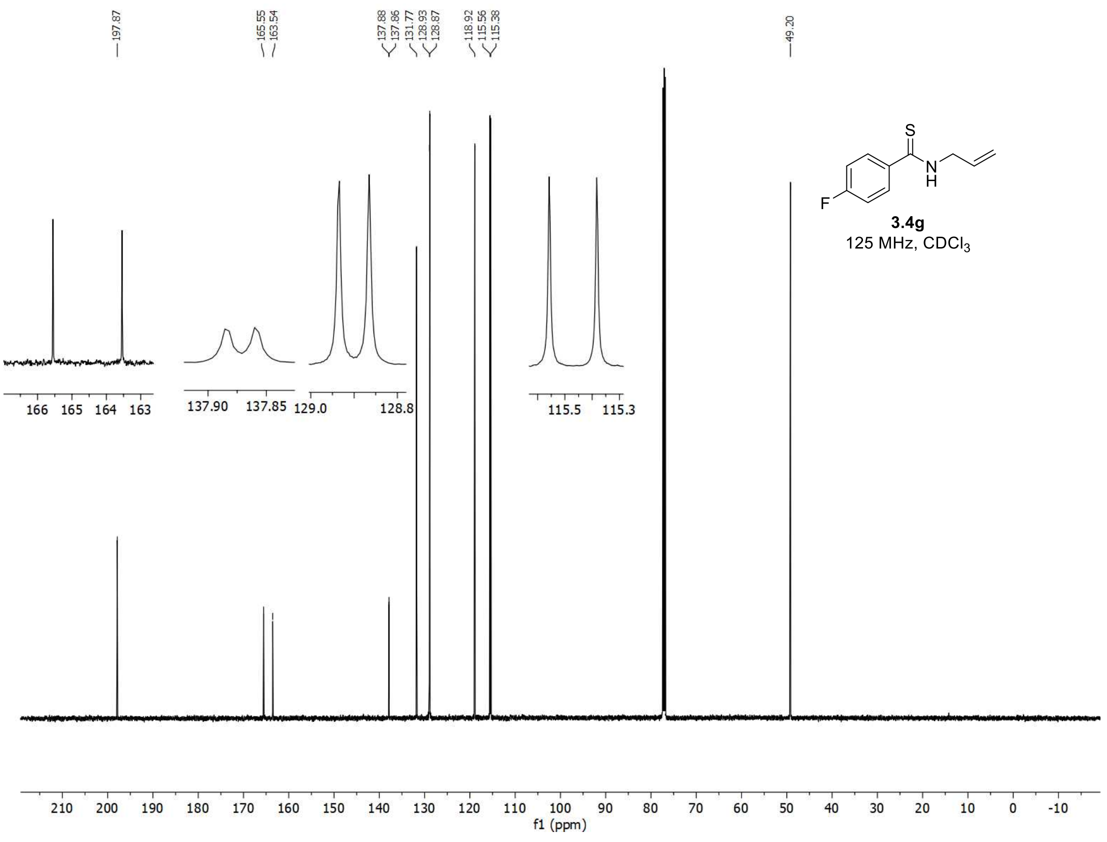
197.87
165.55
163.54
137.88
137.86
131.77
128.93
128.87
118.92
115.56
115.38
49.20
3.4g
125 MHz, CDCl3
f1 (ppm)

$^{1}H$, $^{19}F$, and $^{13}C$ spectra of **3.6g**

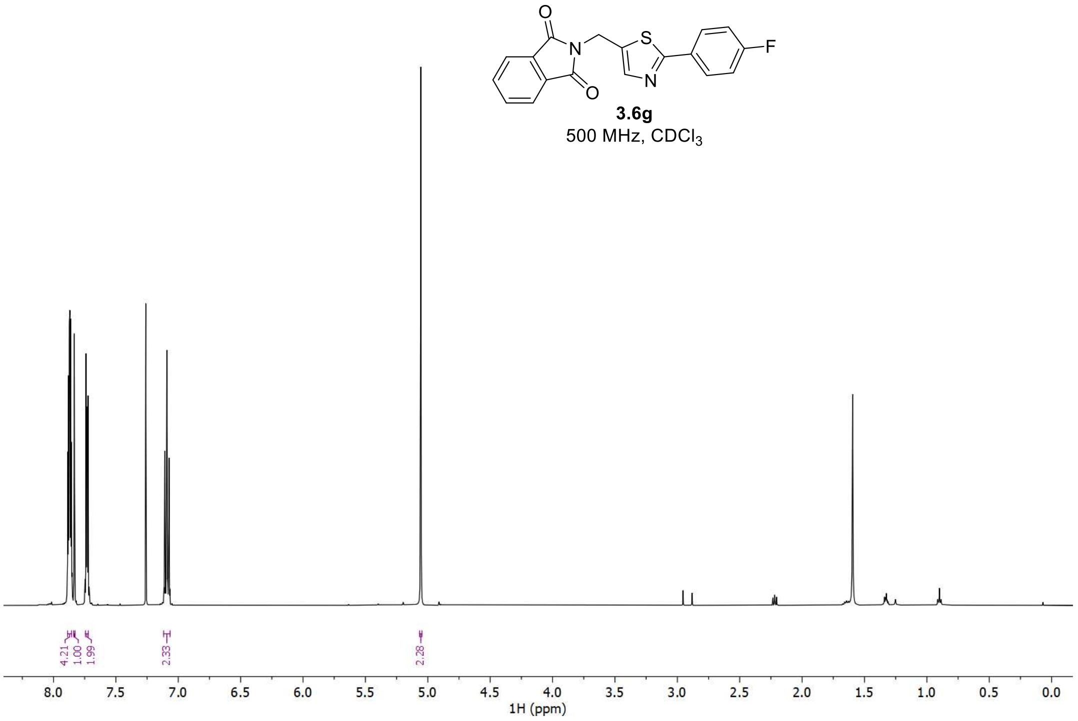


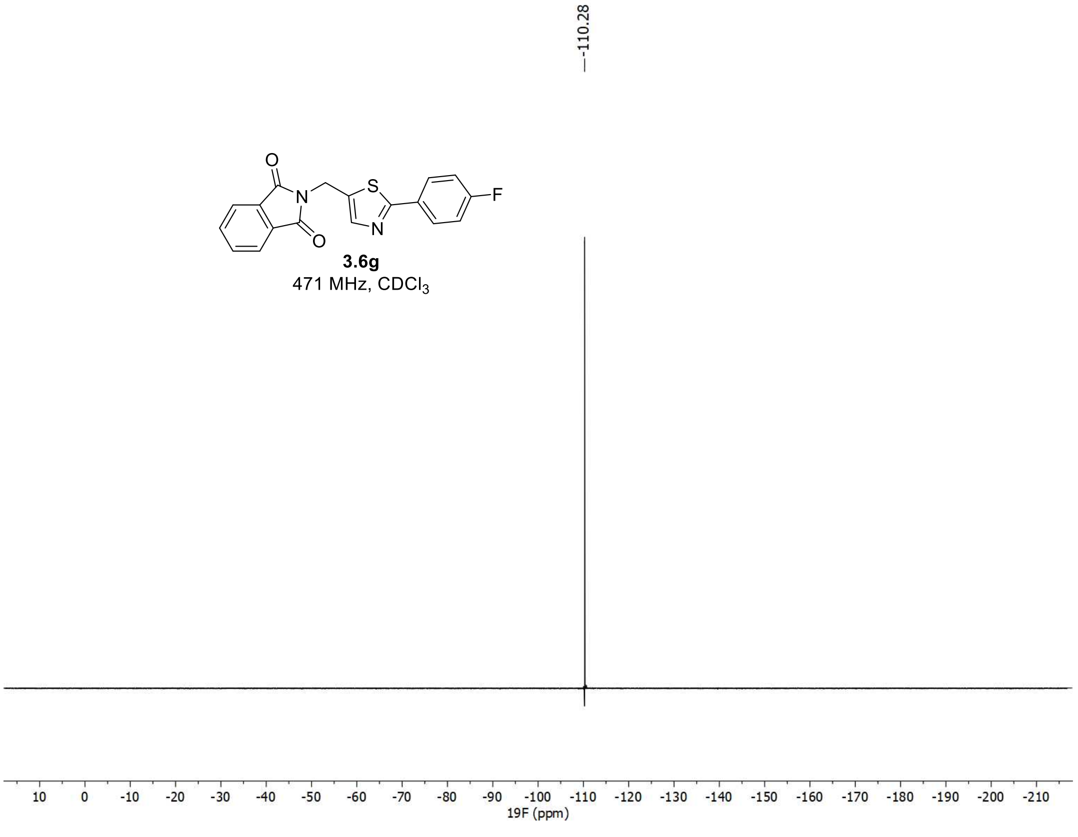

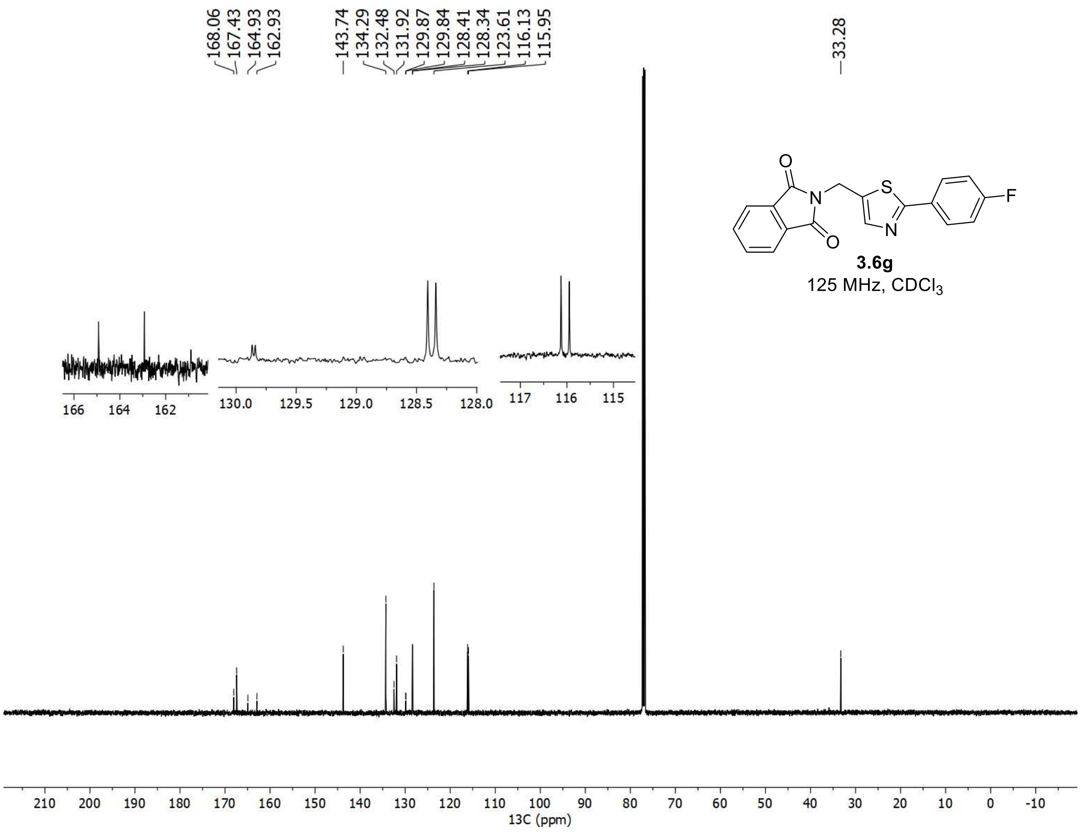
168.06
167.43
164.93
162.93
143.74
134.29
132.48
131.92
129.87
129.84
128.41
128.34
123.61
116.13
115.95
33.28
3.6g
125 MHz, $CDCl_3$
13C (ppm)

$^{1}H$, $^{19}F$, and $^{13}C$ spectra of **3.7g**

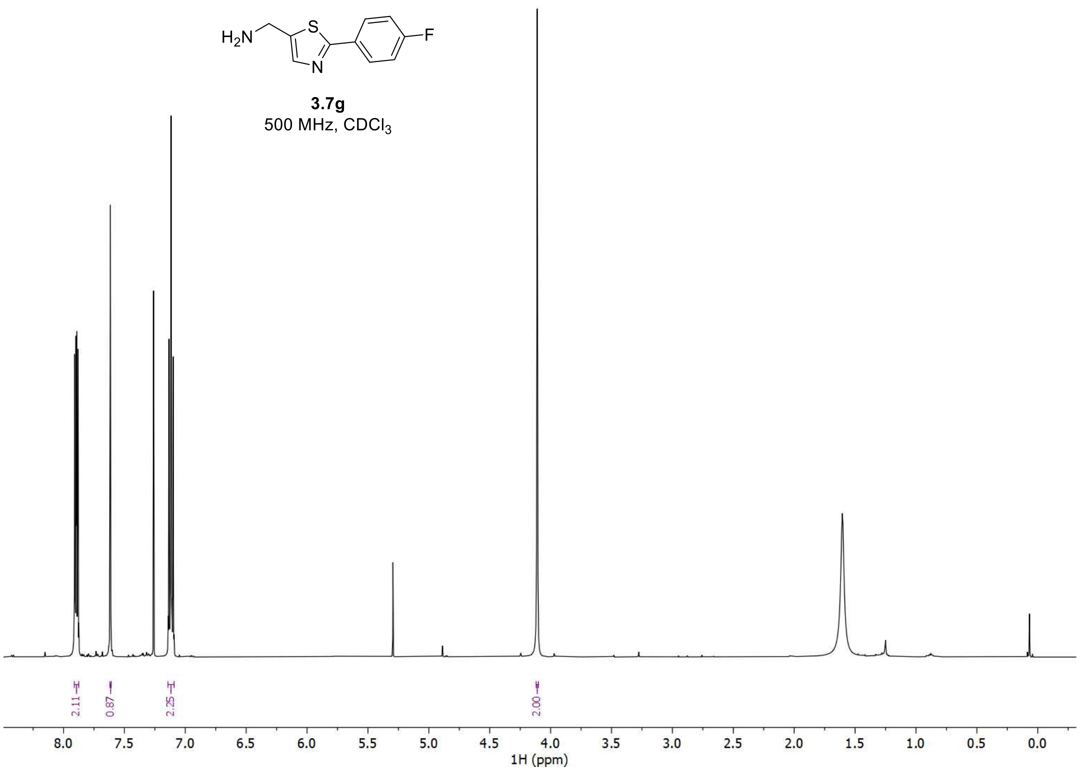


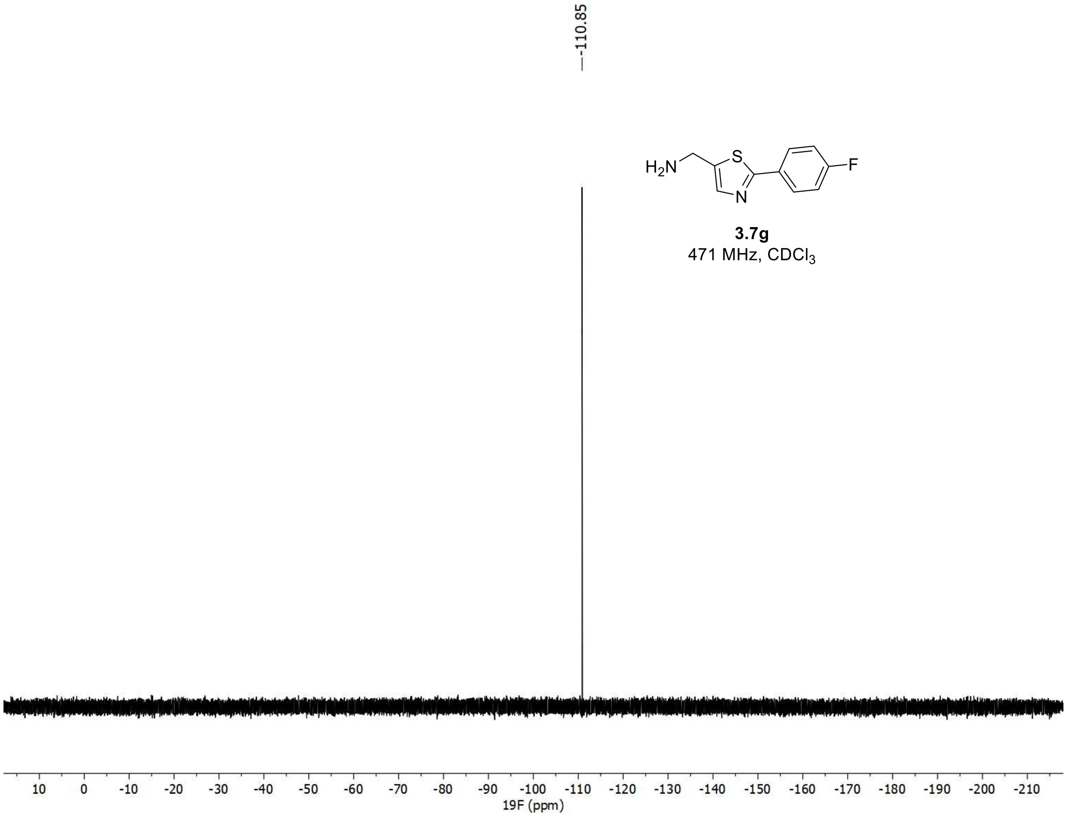

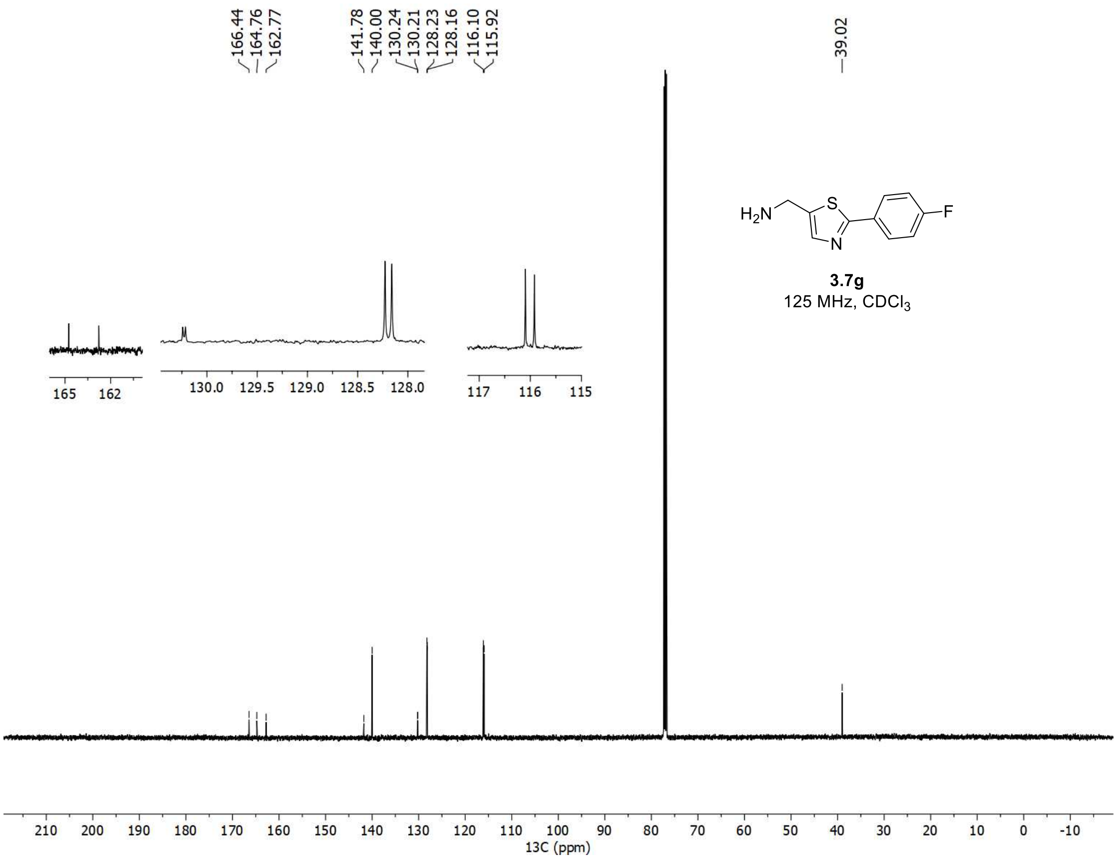

166.44
164.76
162.77
141.78
140.00
130.24
130.21
128.23
128.16
116.10
115.92
39.02
H2N
S
N
F
3.7g
125 MHz, CDCl3
165 162
130.0 129.5 129.0 128.5 128.0
117 116 115
210 200 190 180 170 160 150 140 130 120 110 100 90 80 70 60 50 40 30 20 10 0 -10
13C (ppm)

<u>[1]H, [19]F, and [13]C spectra of **3g**</u>

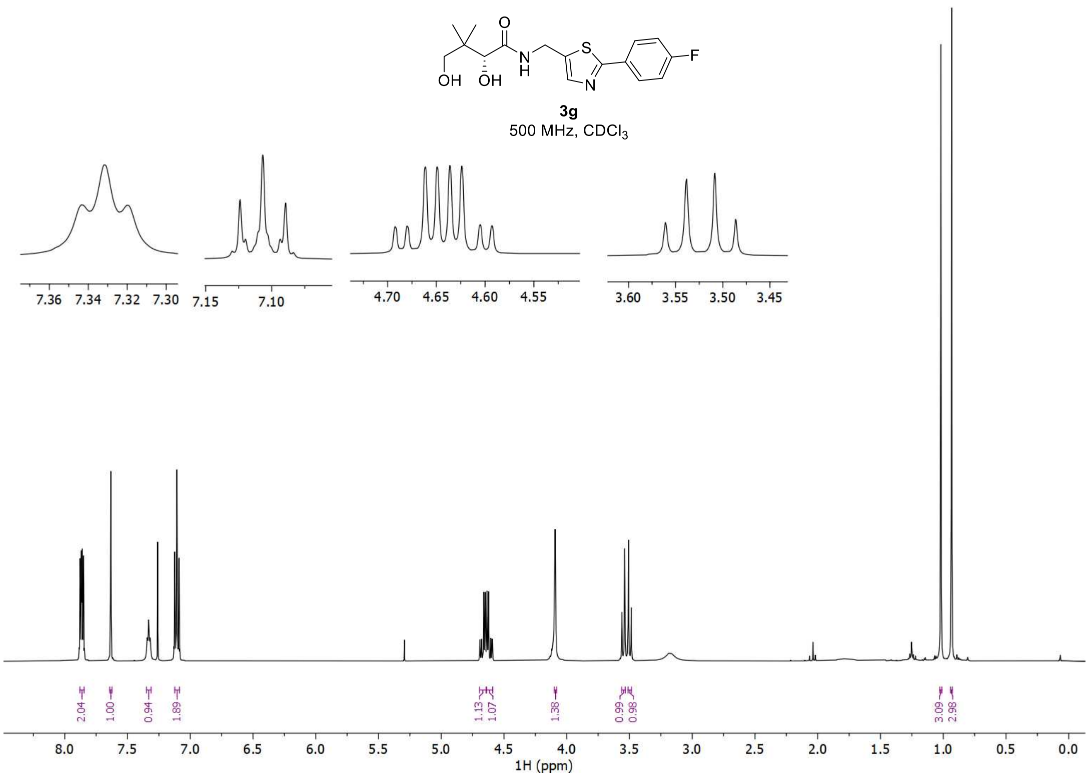


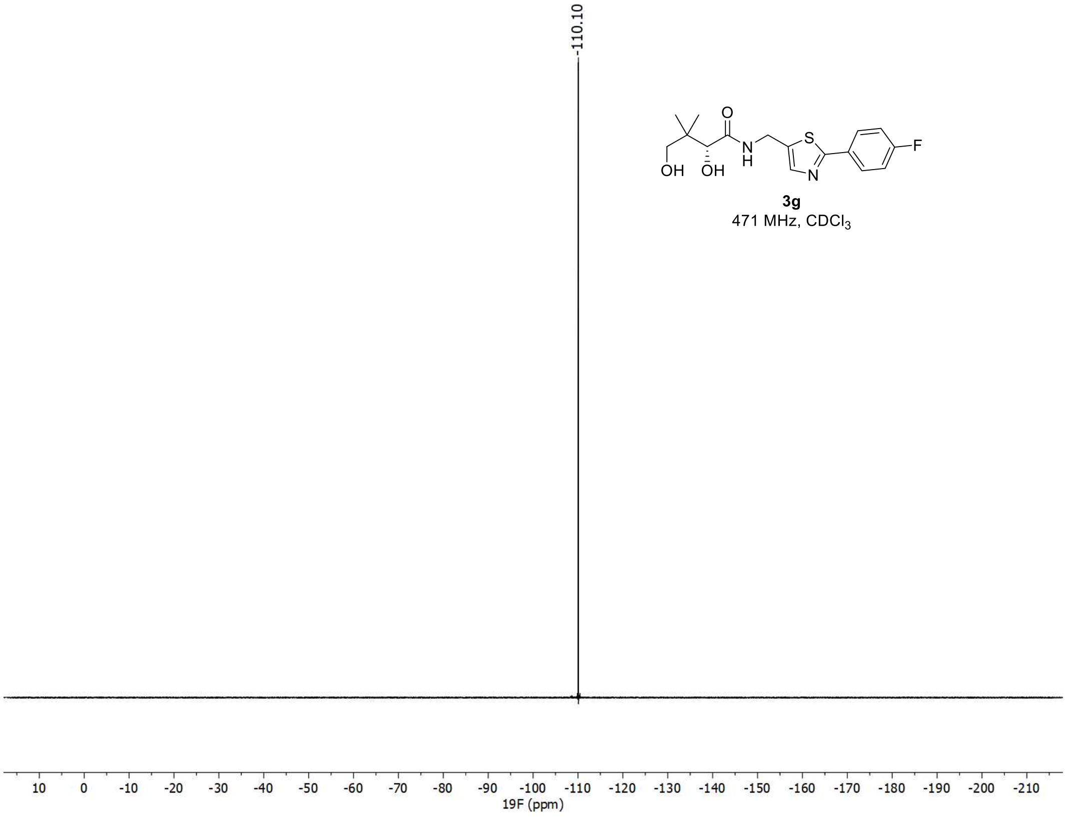

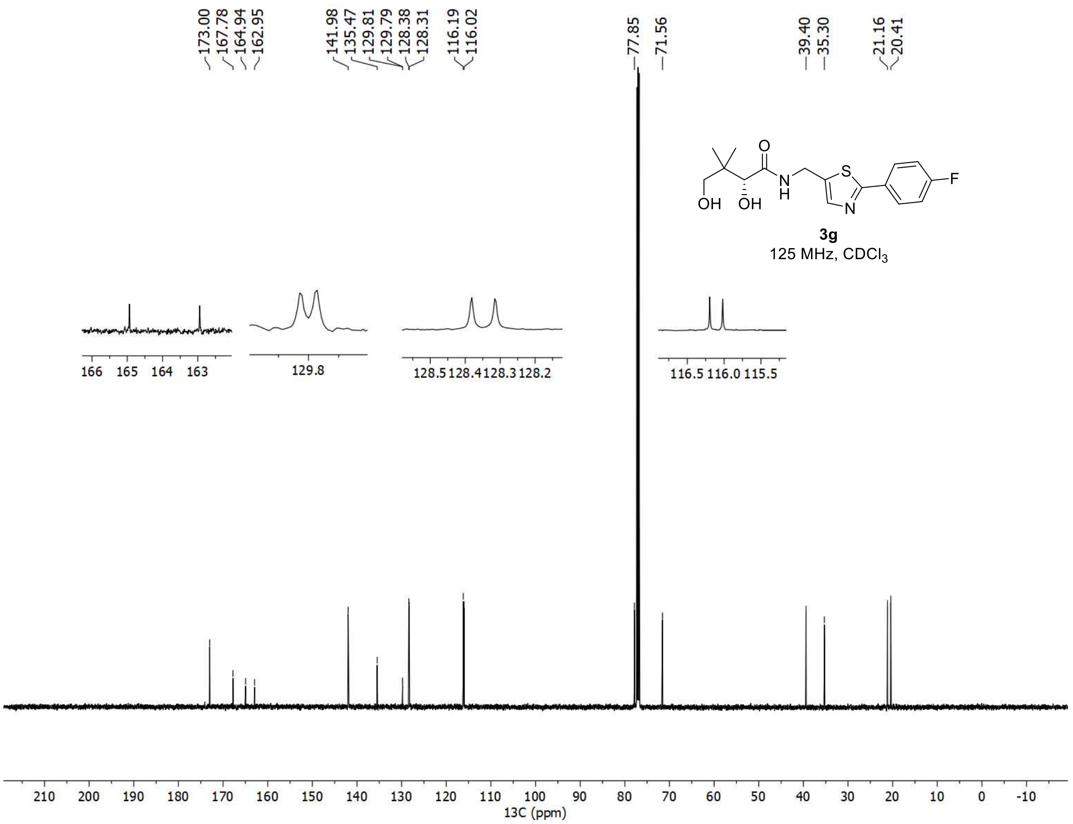
173.00
167.78
164.94
162.95
141.98
135.47
129.81
129.79
128.38
128.31
116.19
116.02
77.85
71.56
39.40
35.30
21.16
20.41
3g
125 MHz, CDCl3
166 165 164 163
129.8
128.5 128.4 128.3 128.2
116.5 116.0 115.5
210 200 190 180 170 160 150 140 130 120 110 100 90 80 70 60 50 40 30 20 10 0 -10
13C (ppm)

[1]H and [13]C spectra of **3.4h**

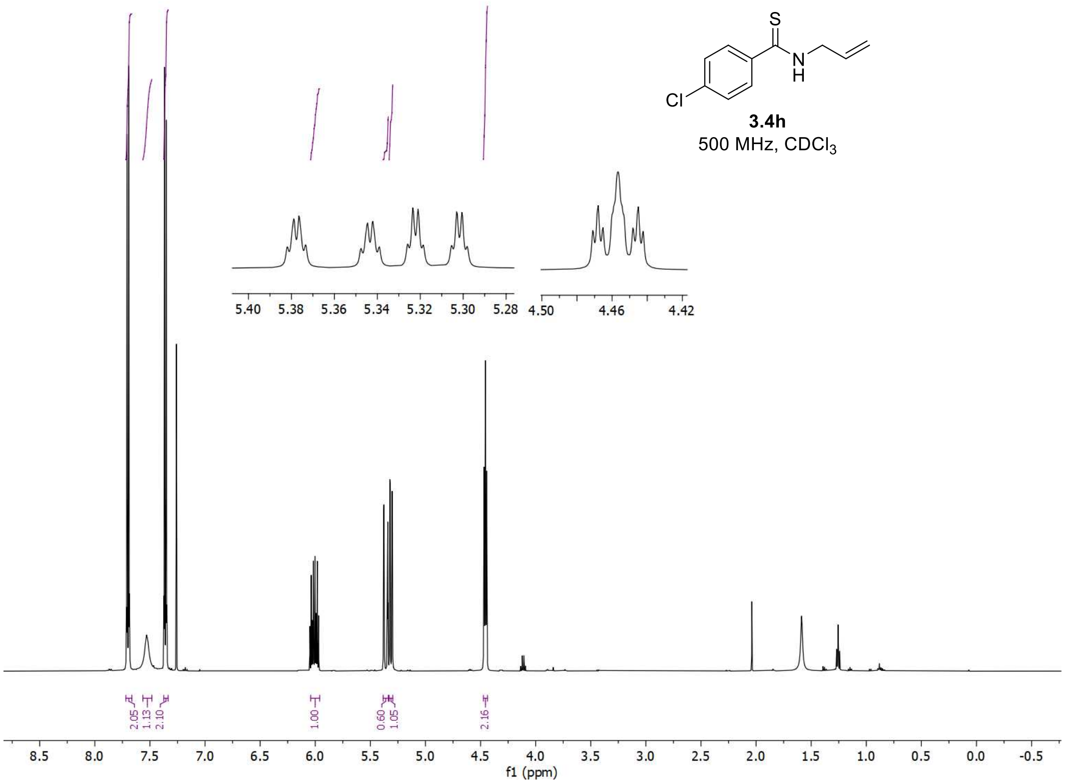


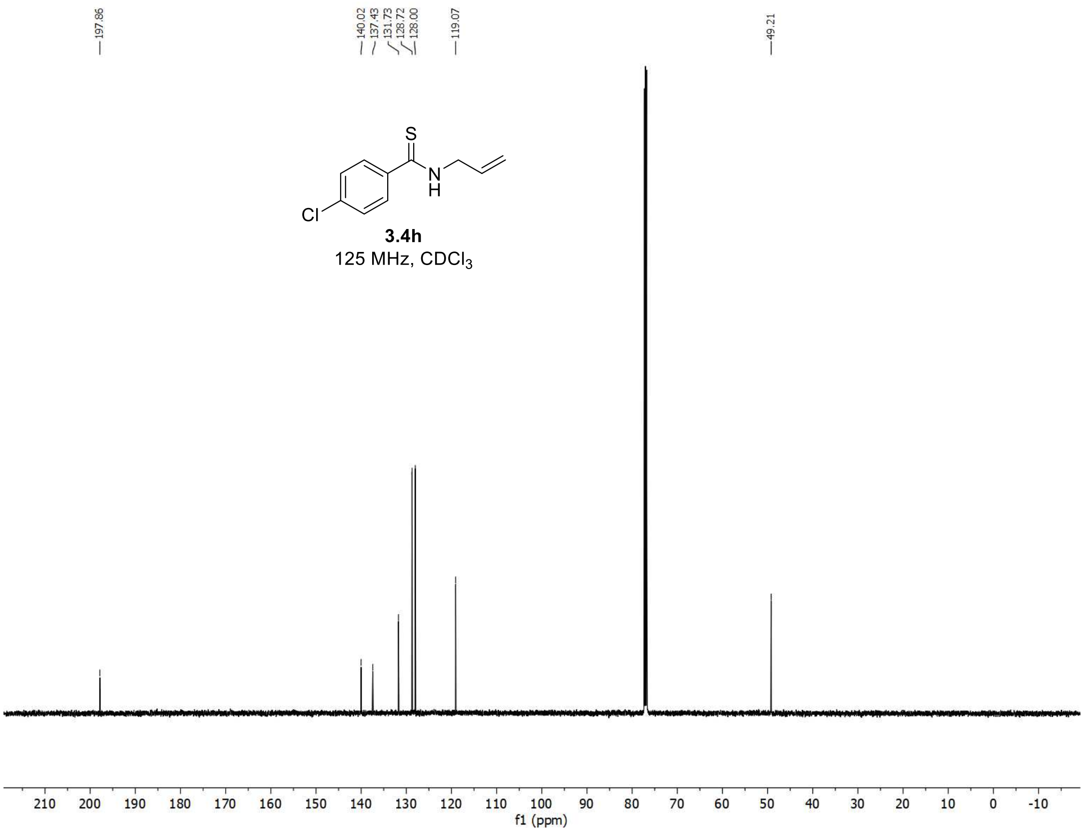

[1]H and [13]C spectra of **3.6h**

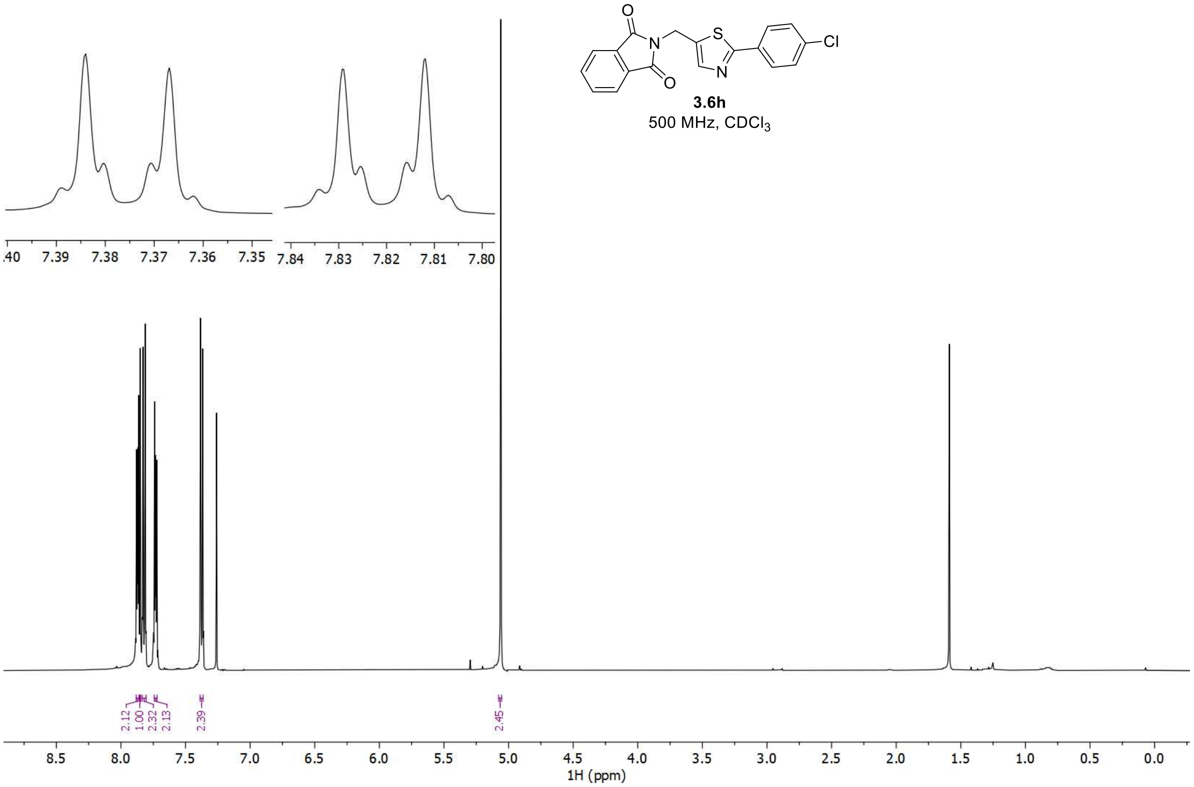


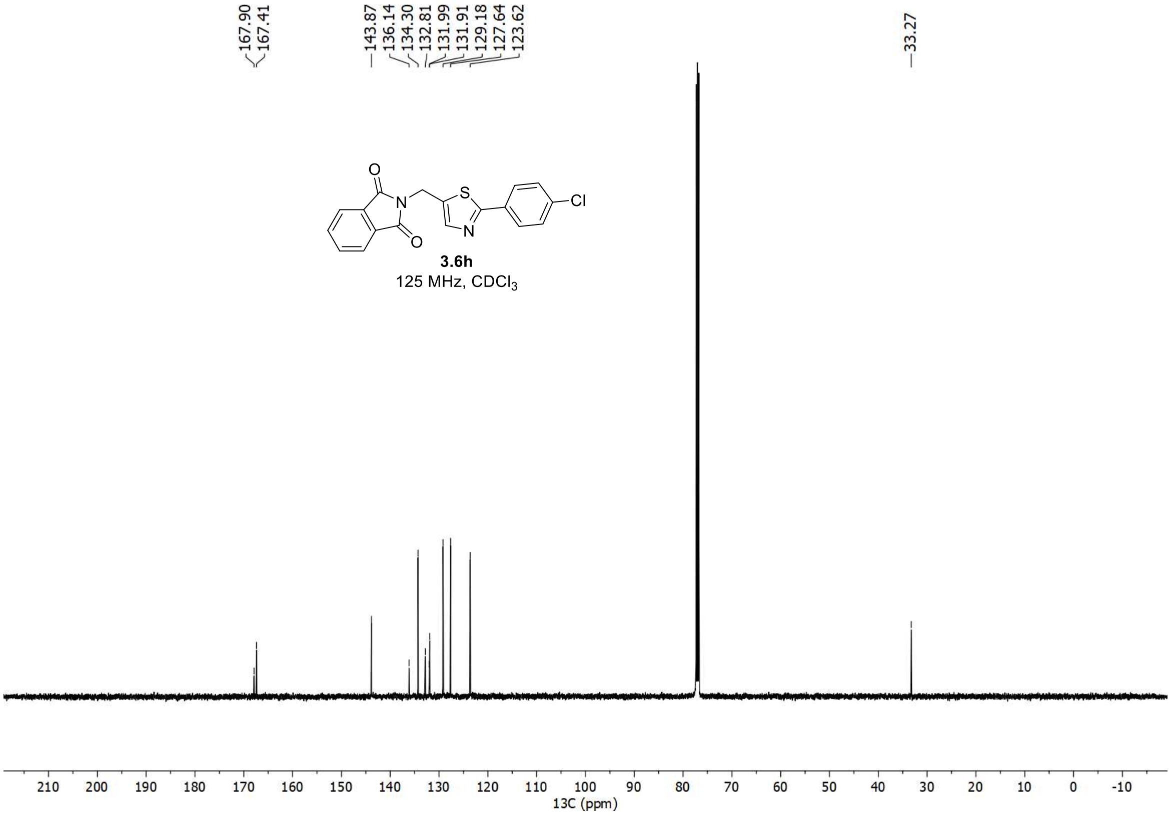

[1]H and [13]C spectra of **3.7h**

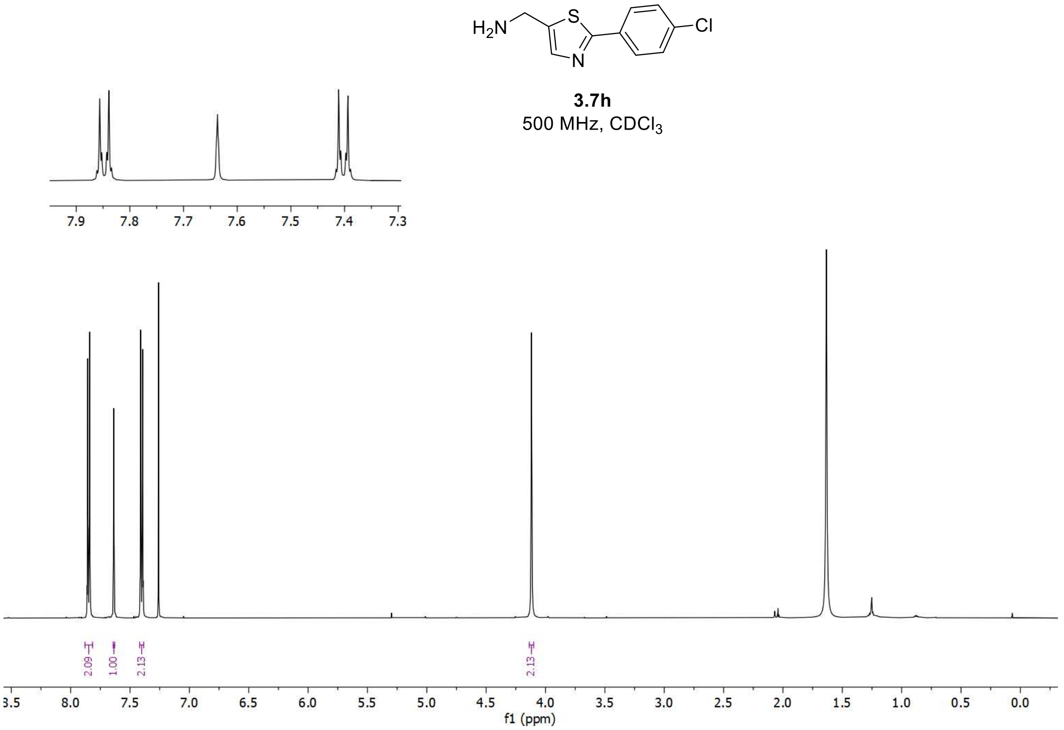


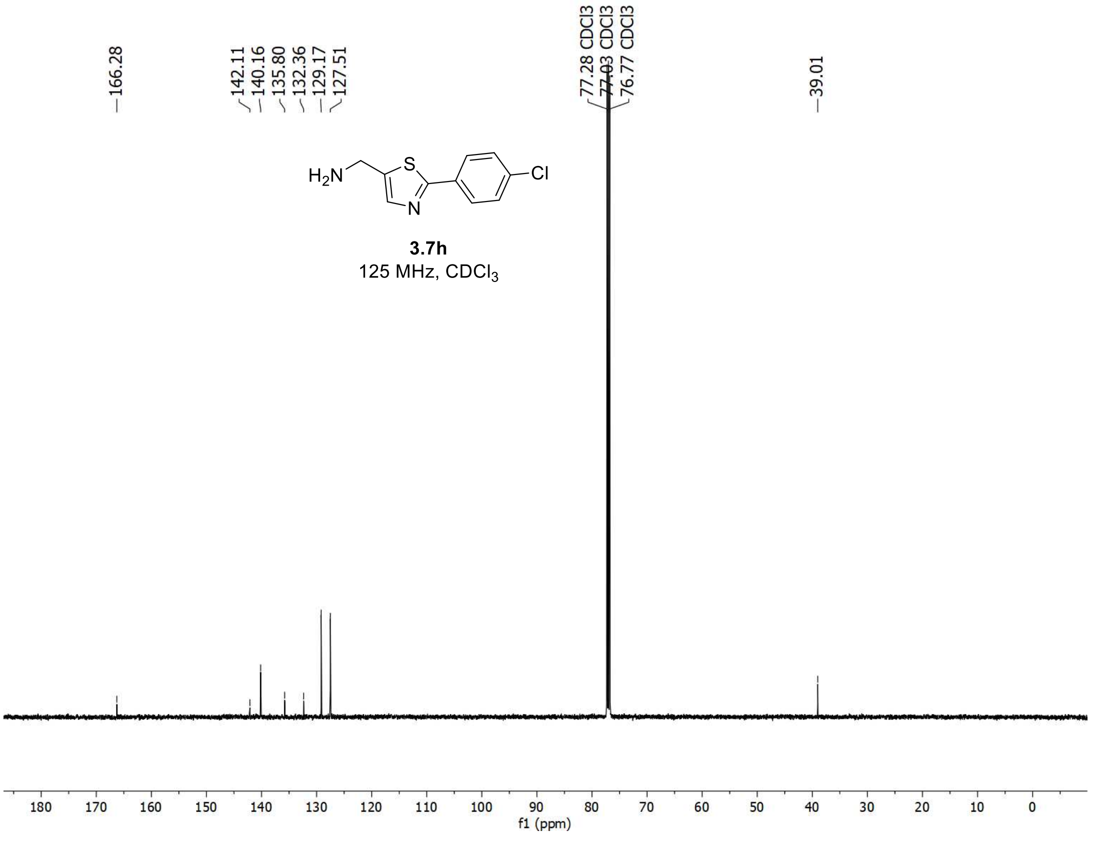

$^{1}H$ and $^{13}C$ spectra of **3h**

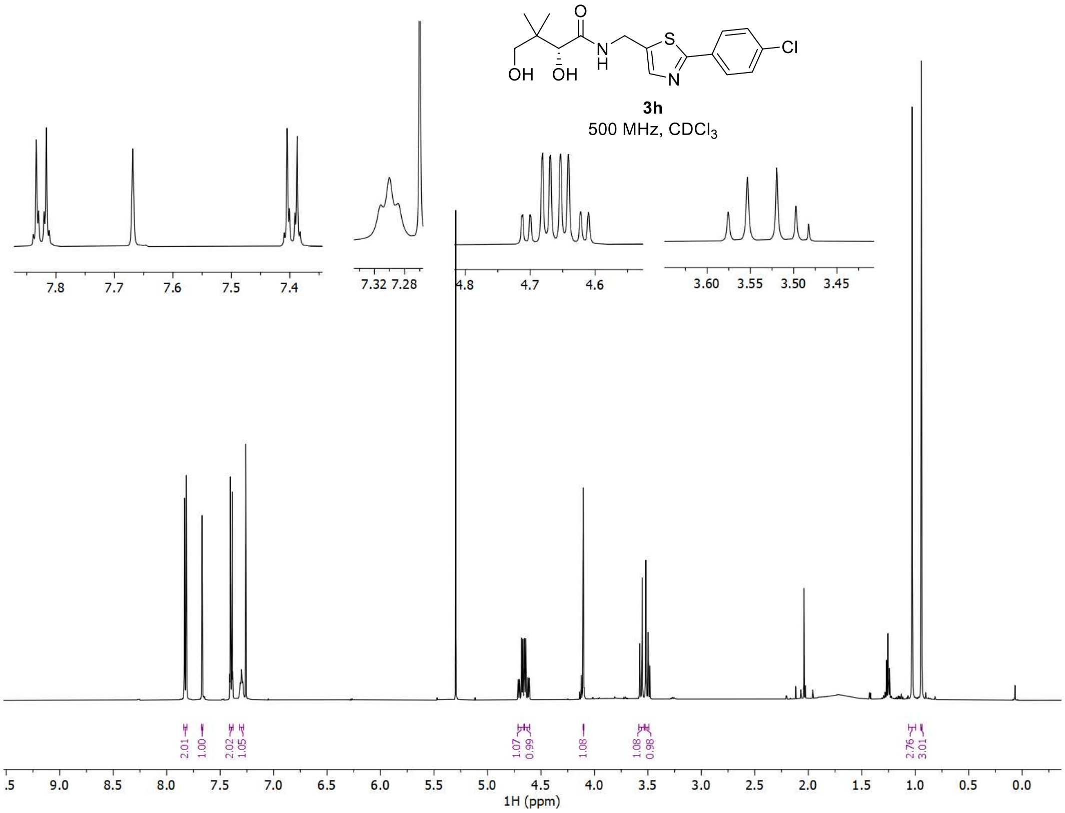


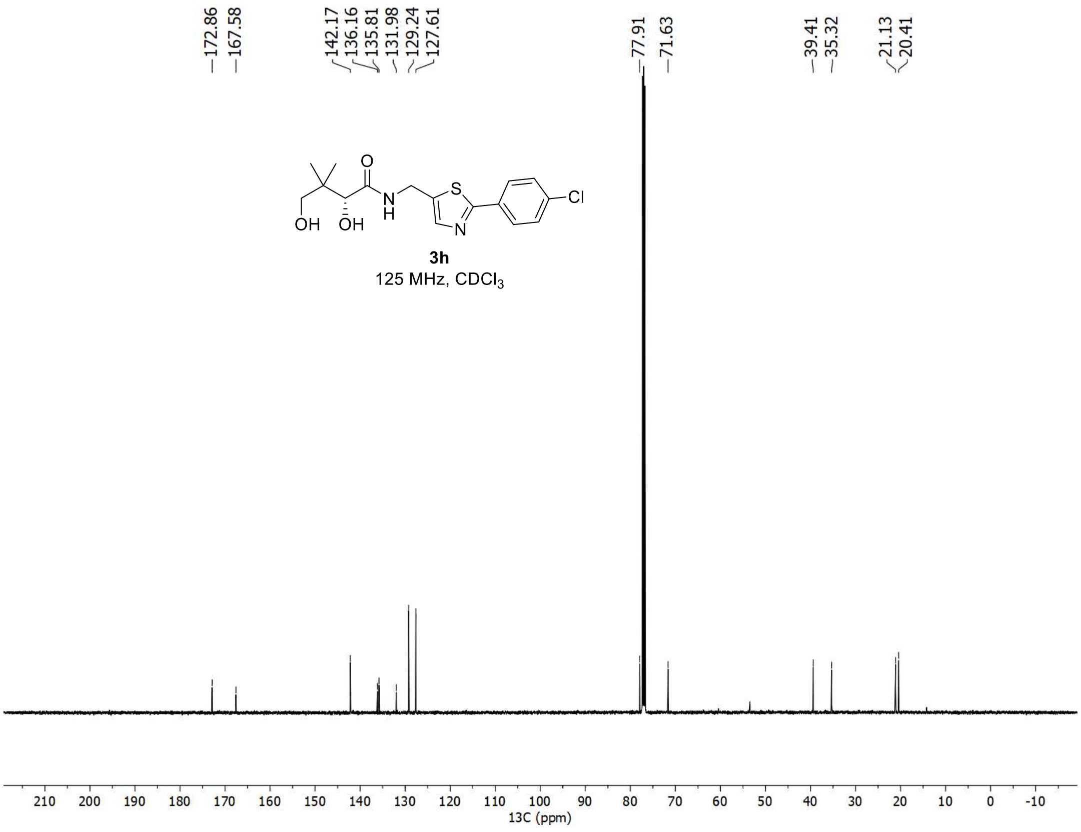

$^{1}H$, $^{19}F$, and $^{13}C$ spectra of **3.4i**

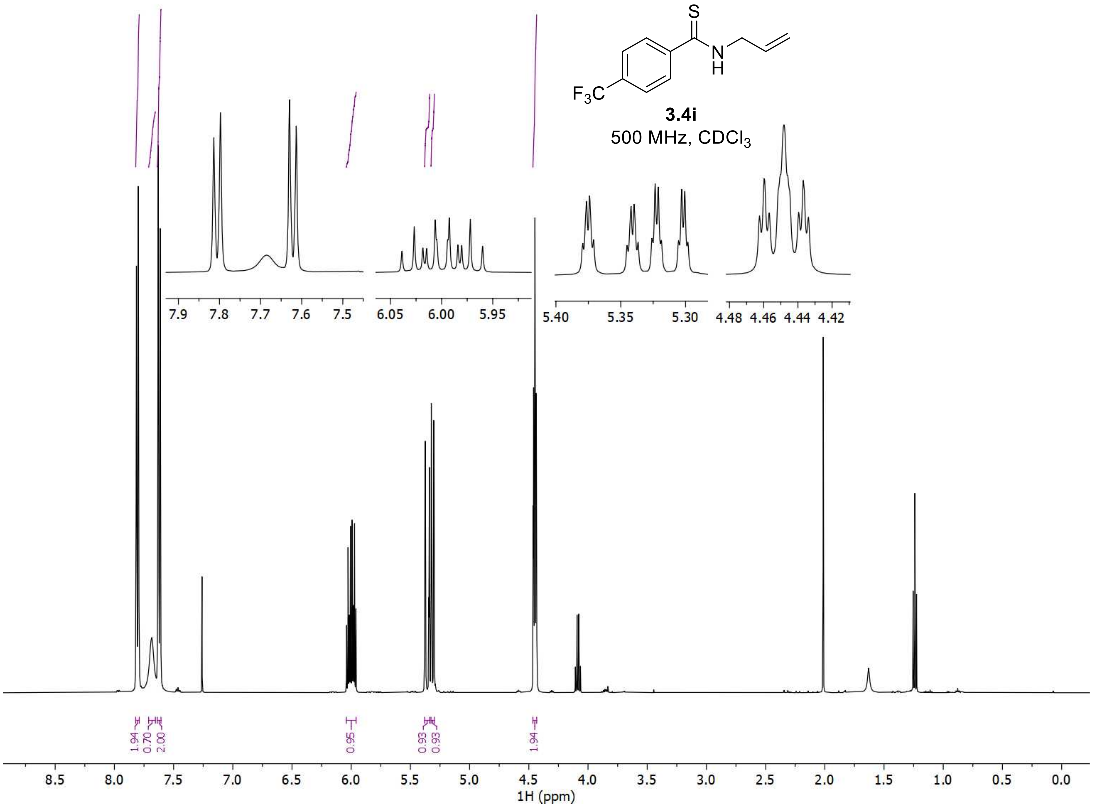


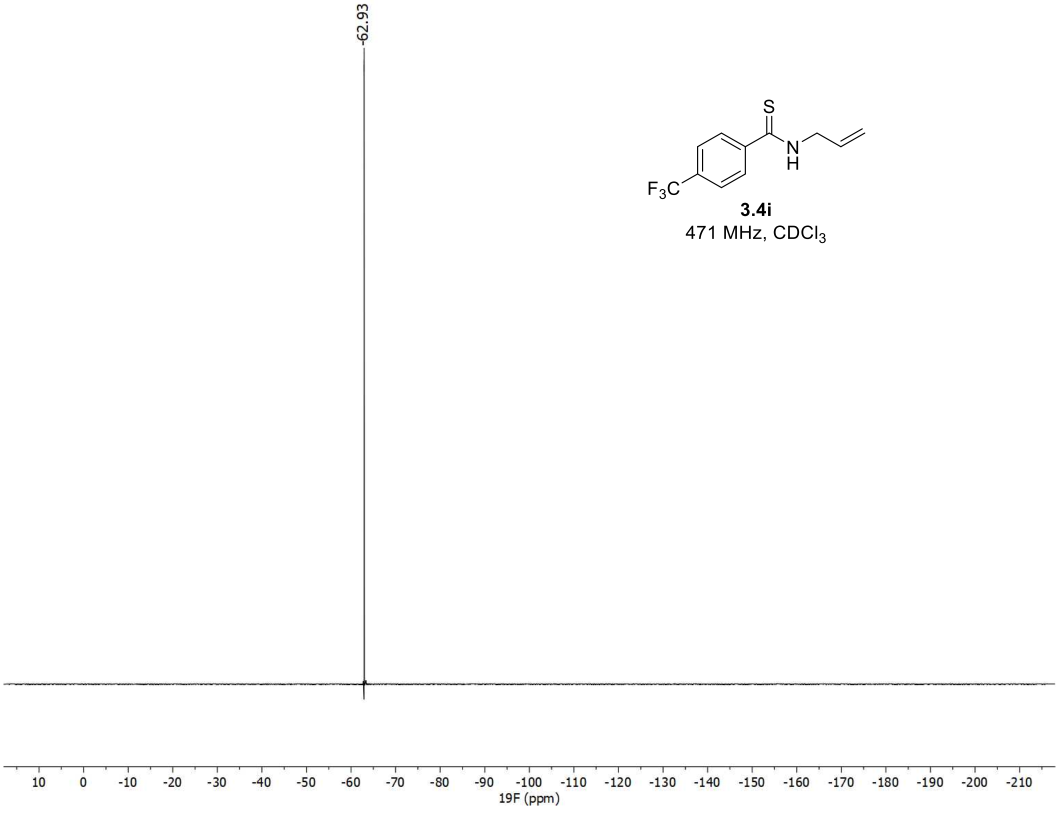

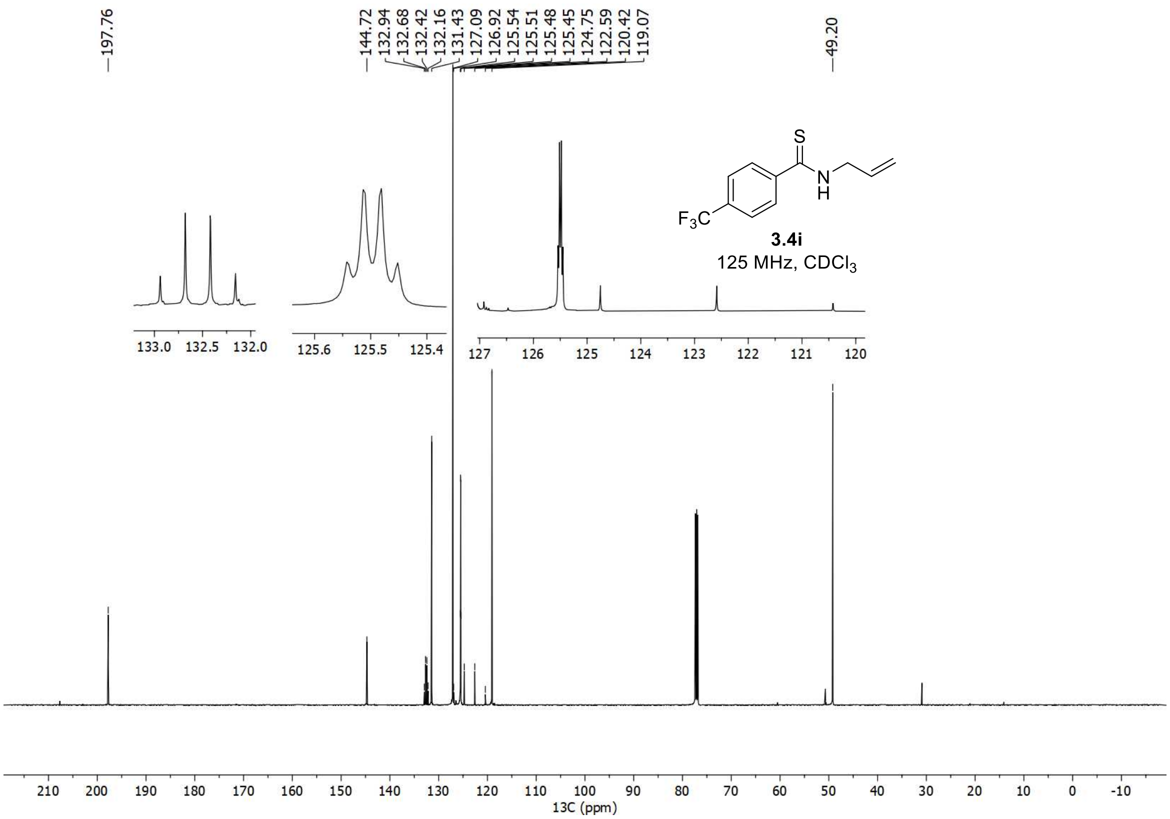
197.76
144.72
132.94
132.68
132.42
132.16
131.43
127.09
126.92
125.54
125.51
125.48
125.45
124.75
122.59
120.42
119.07
49.20
F3C
S
N
H
3.4i
125 MHz, CDCl3
13C (ppm)

<u>$^{1}$H, $^{19}$F, and $^{13}$C spectra of **3.6i**</u>

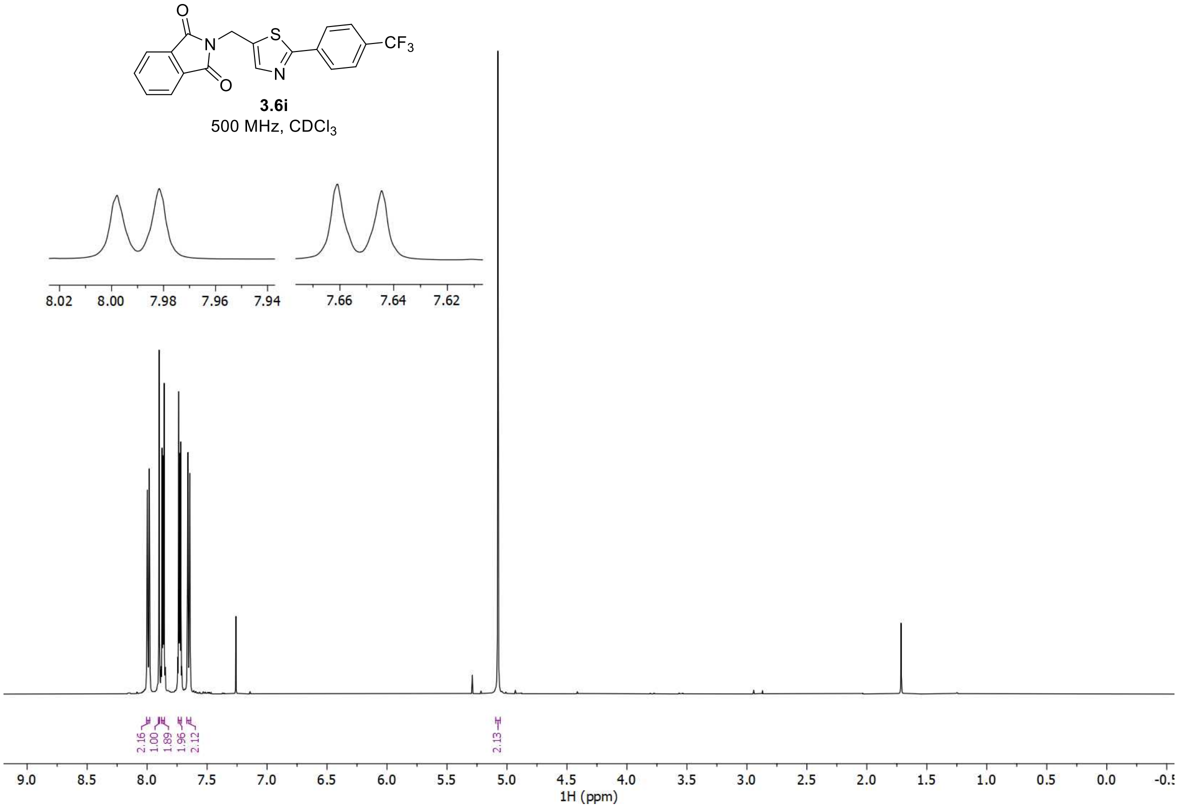


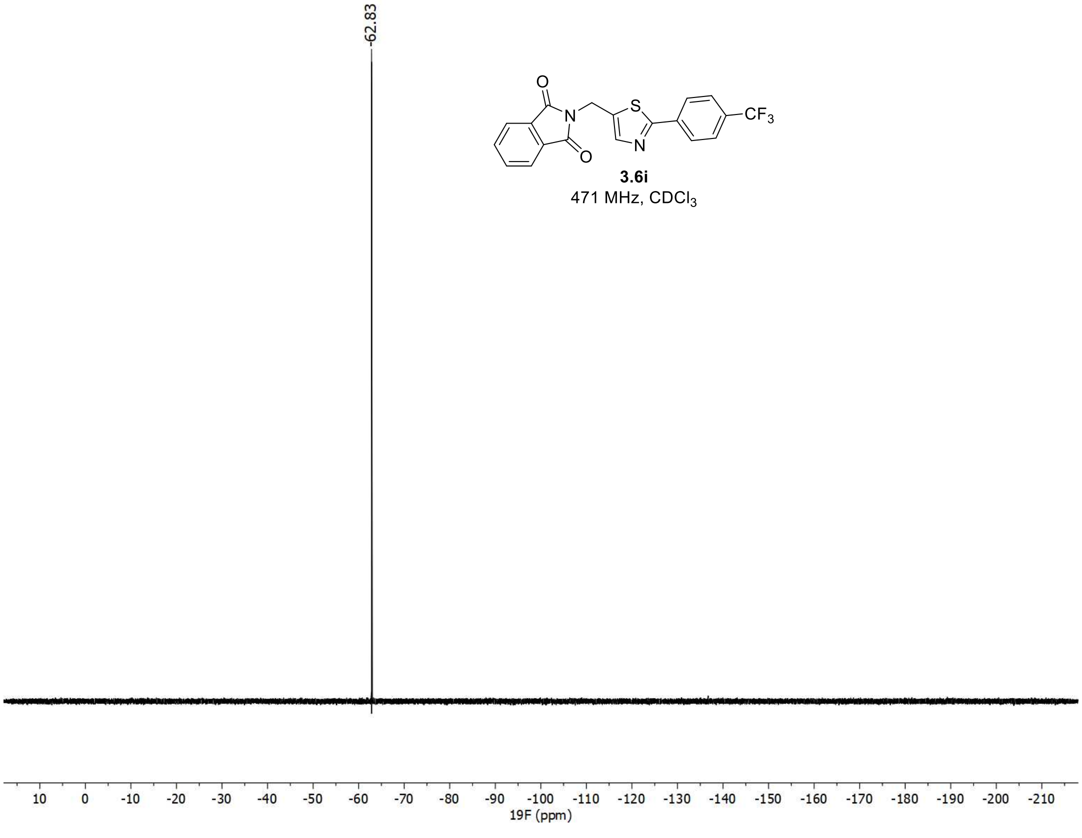

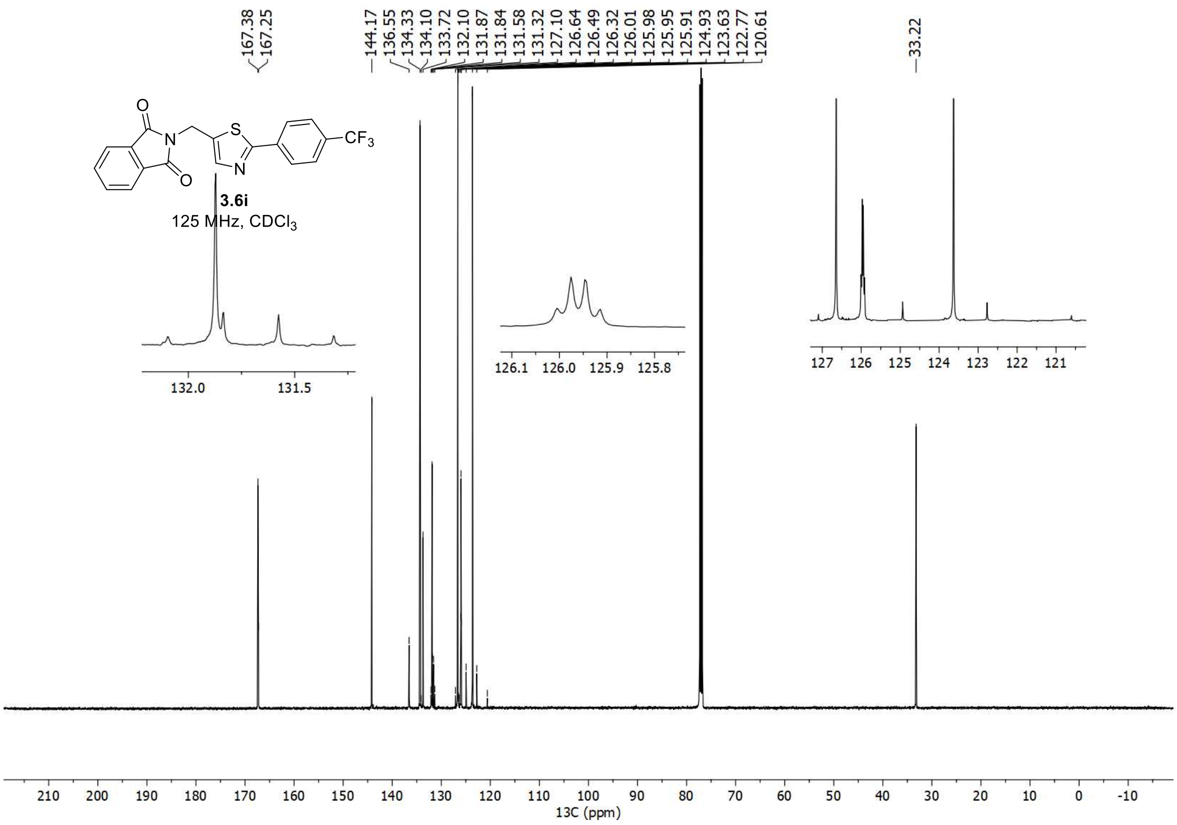
167.38
167.25
144.17
136.55
134.33
134.10
133.72
132.10
131.87
131.84
131.58
131.32
127.10
126.64
126.49
126.32
126.01
125.98
125.95
125.91
124.93
123.63
122.77
120.61
33.22
O
N
S
N
O
CF3
3.6i
125 MHz, CDCl3
132.0
131.5
126.1
126.0
125.9
125.8
127
126
125
124
123
122
121
210
200
190
180
170
160
150
140
130
120
110
100
90
80
70
60
50
40
30
20
10
0
-10
13C (ppm)

$^1$H, $^{19}$F, and $^{13}$C spectra of **3.7i**

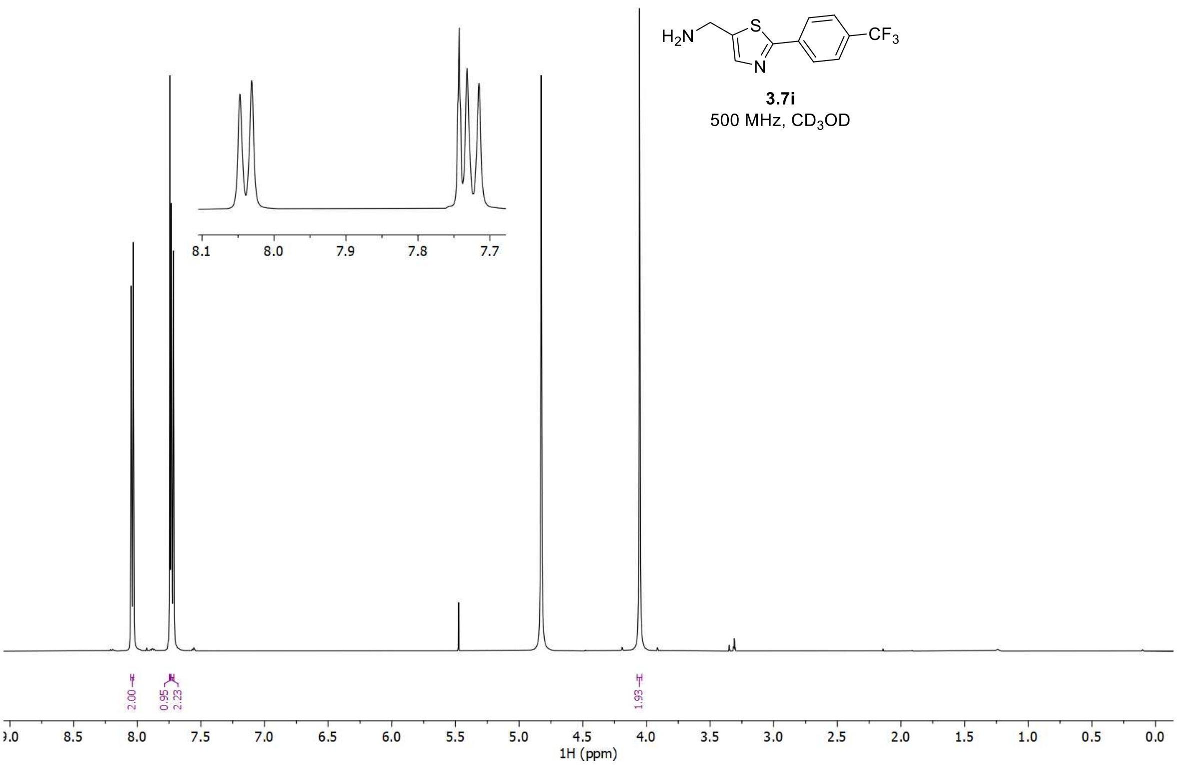


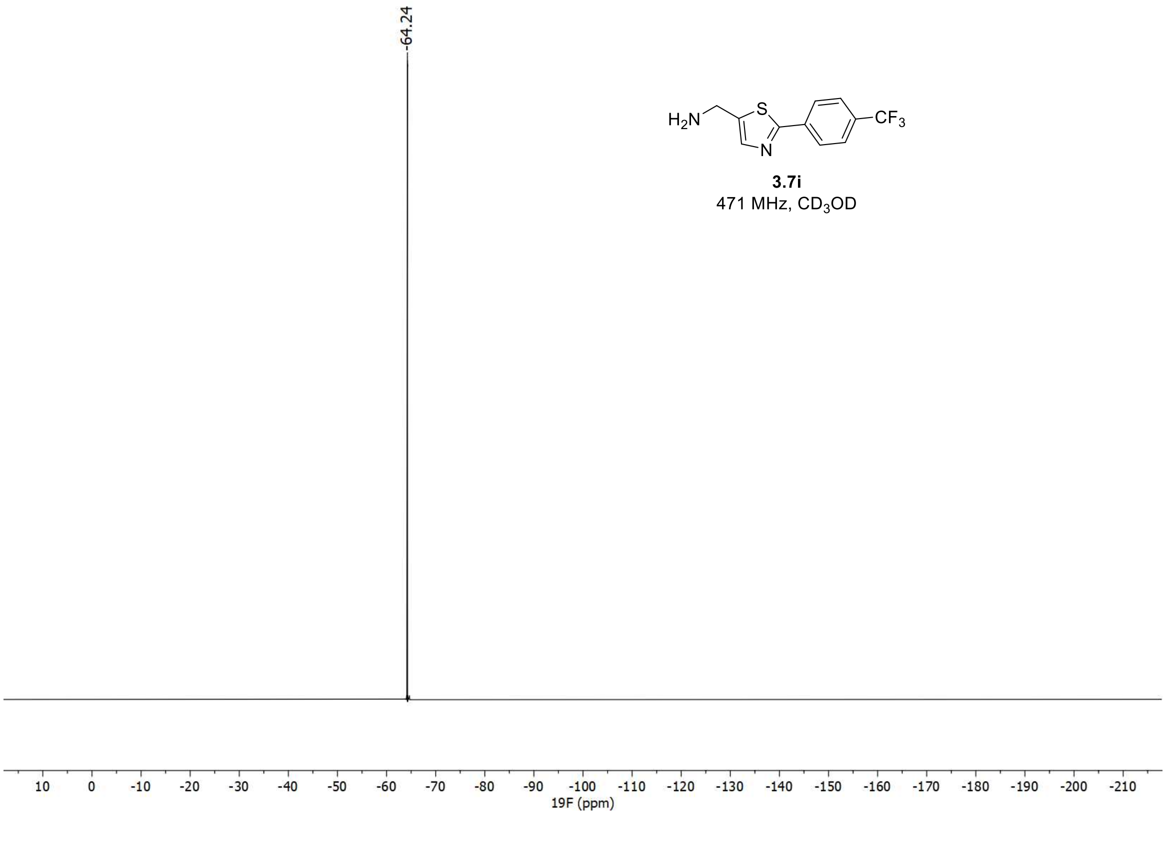

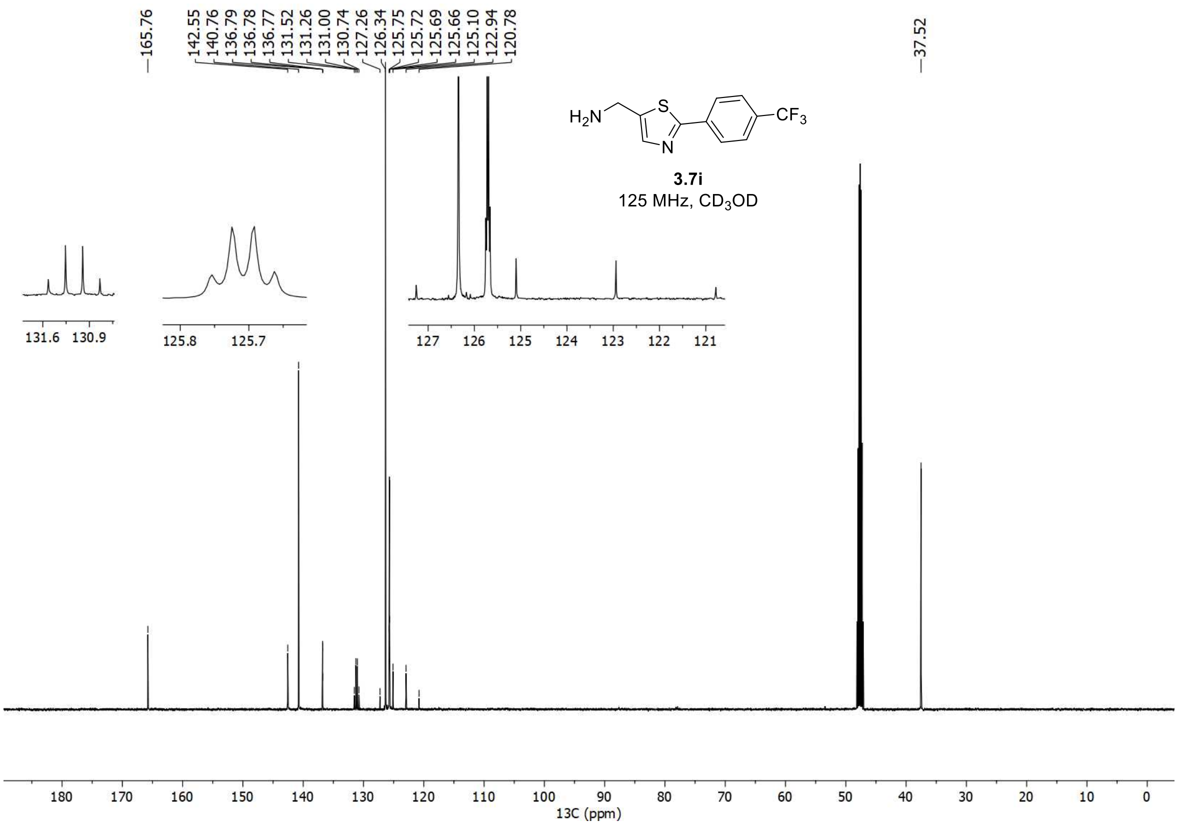

165.76
142.55
140.76
136.79
136.78
136.77
131.52
131.26
131.00
130.74
127.26
126.34
125.75
125.72
125.69
125.66
125.10
122.94
120.78
37.52
$H_2N$
S
N
$CF_3$
3.7i
125 MHz, $CD_3OD$
13C (ppm)

$^{1}H$, $^{19}F$, and $^{13}C$ spectra of **3i**

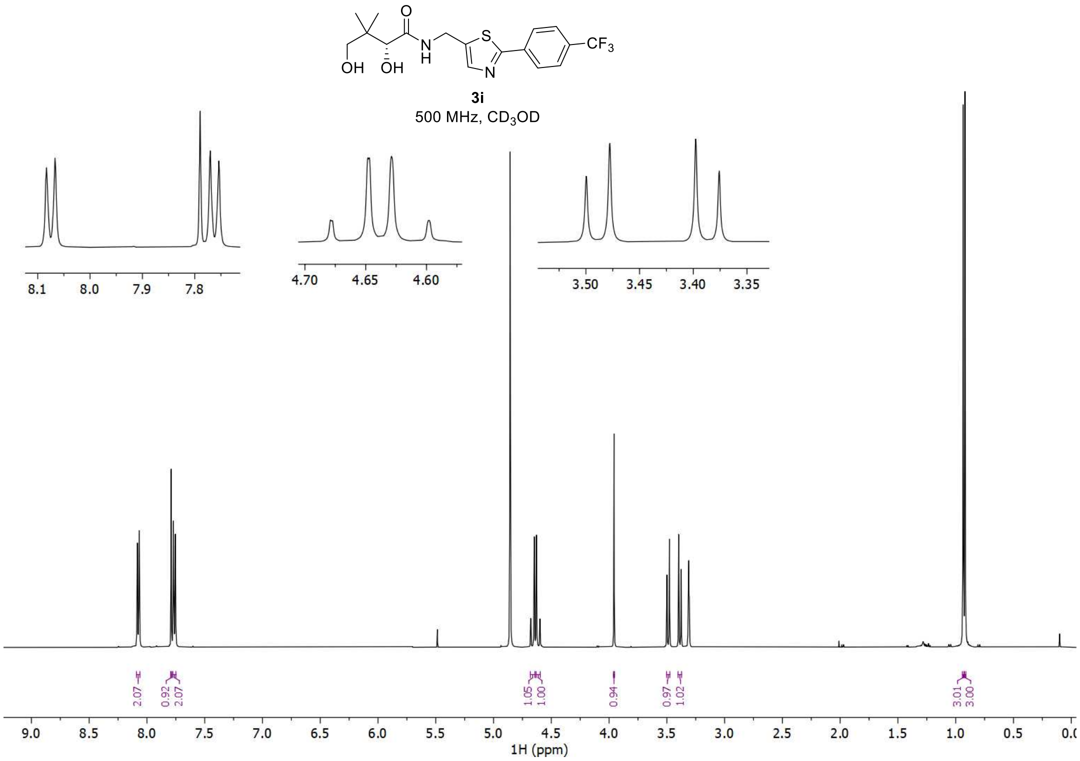


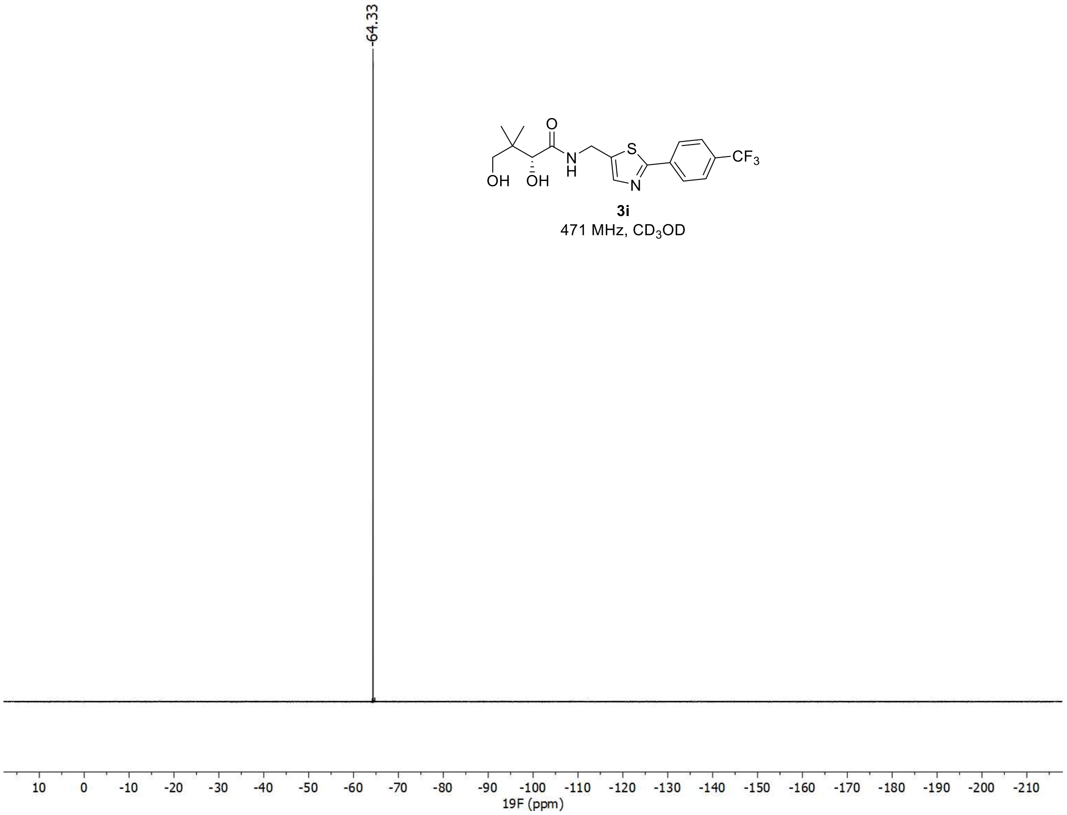

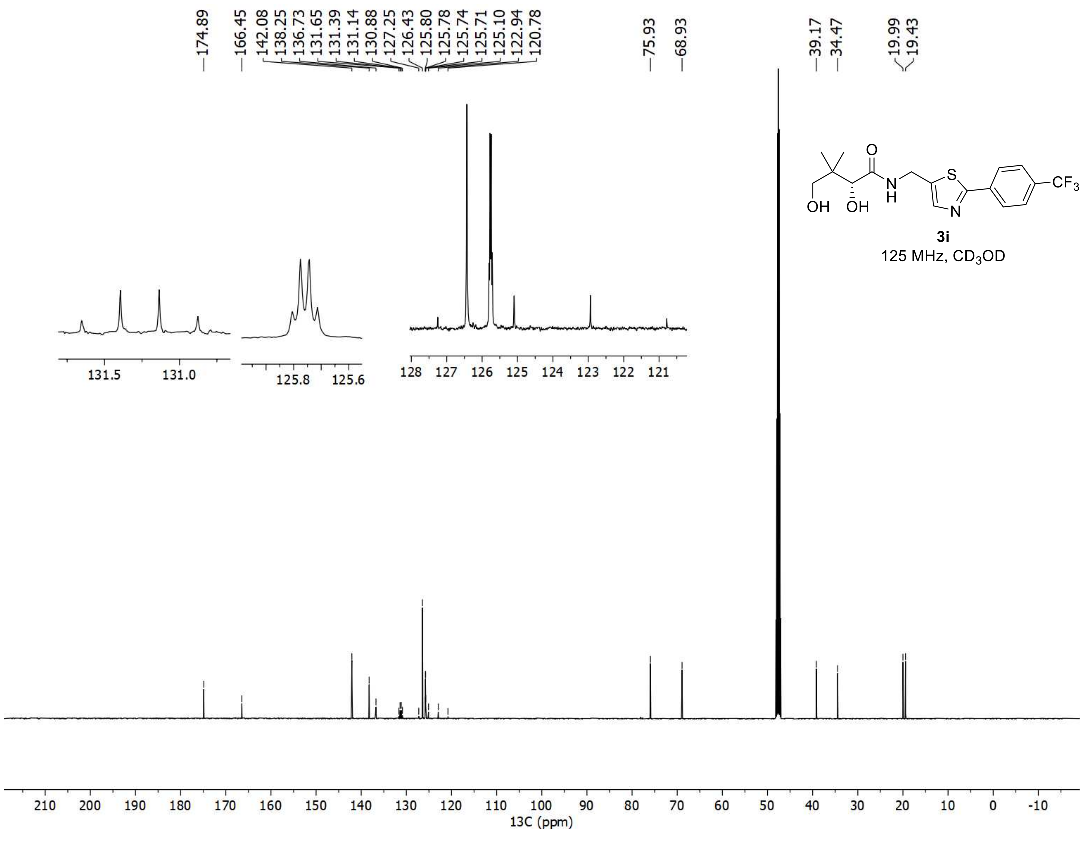
174.89
166.45
142.08
138.25
136.73
131.65
131.39
131.14
130.88
127.25
126.43
125.80
125.78
125.74
125.71
125.10
122.94
120.78
75.93
68.93
39.17
34.47
19.99
19.43
3i
125 MHz, CD3OD
13C (ppm)

$^{1}H$, $^{19}F$, and $^{13}C$ spectra of **3.4j**

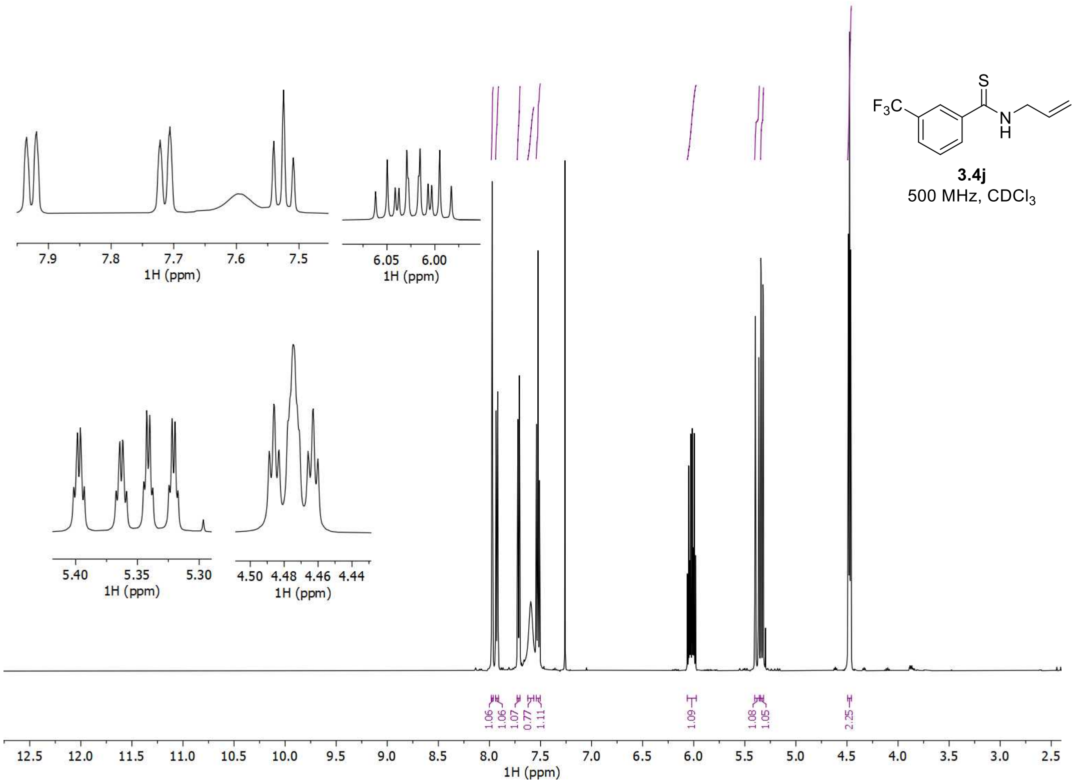


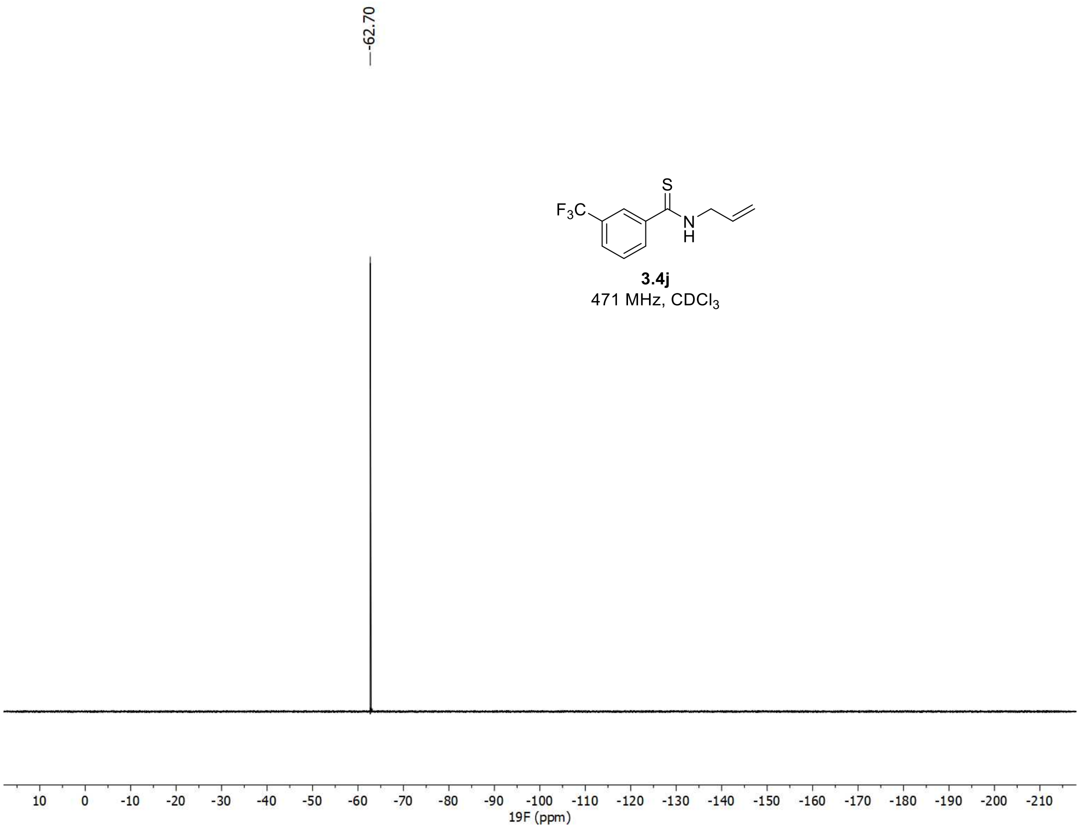

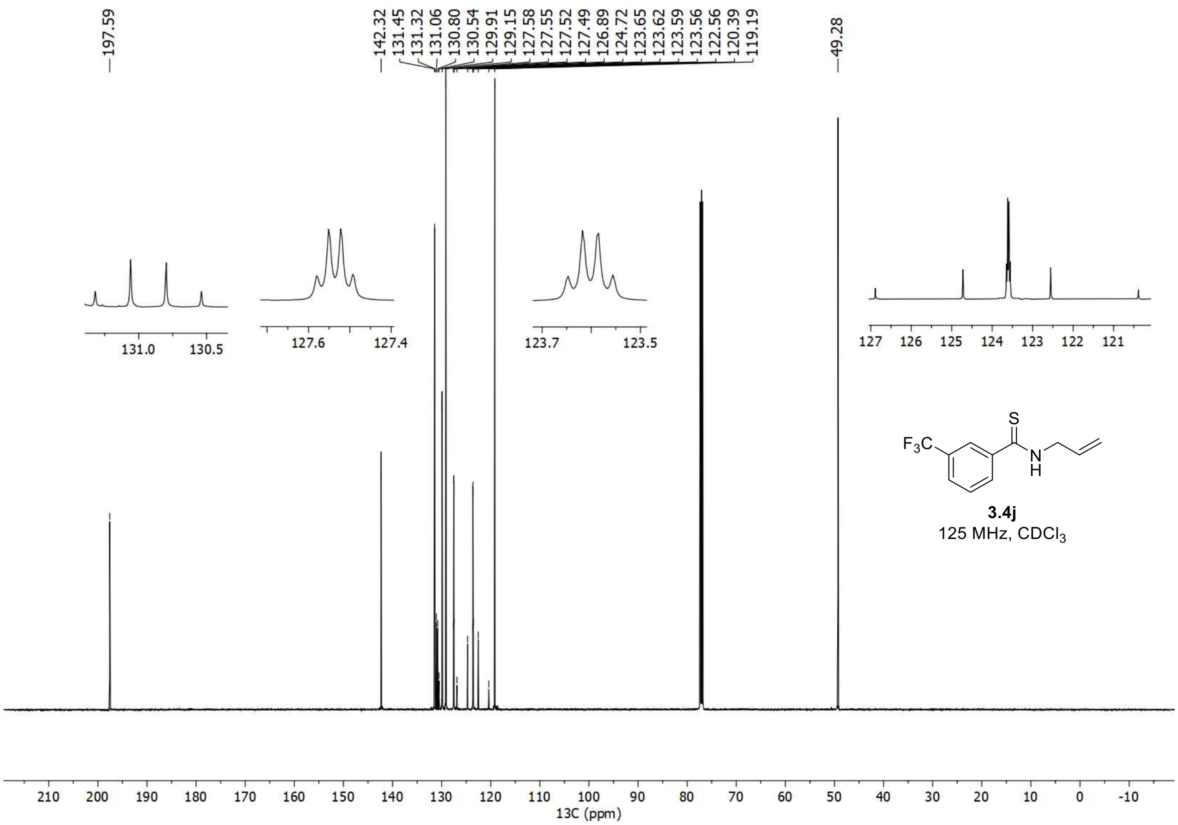


**3.4j**

125 MHz, $CDCl_3$

[1]H, [19]F, and [13]C spectra of **3.6j**

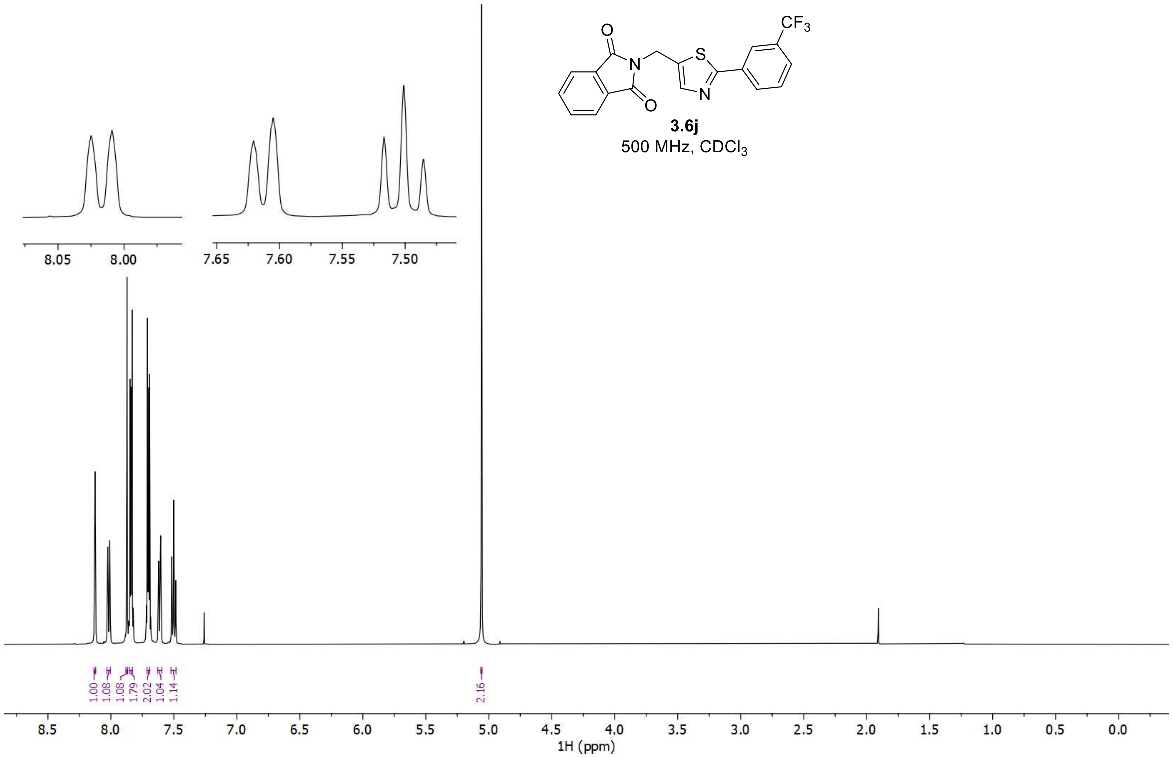


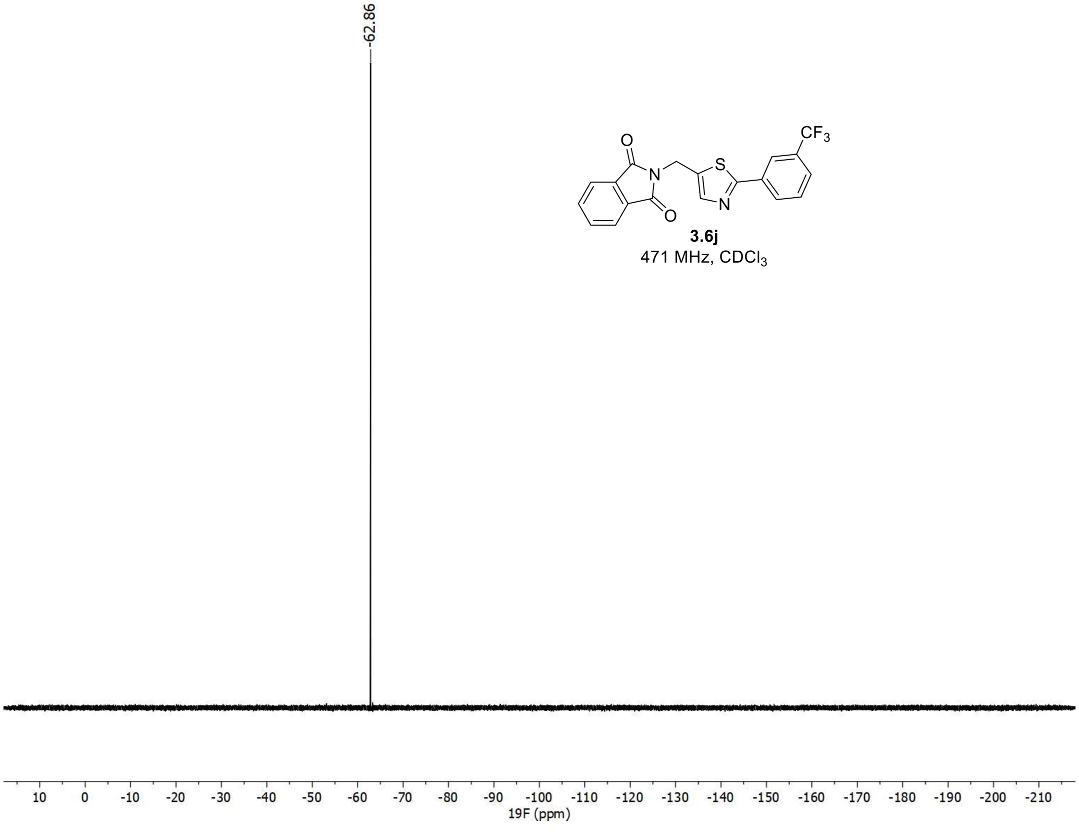

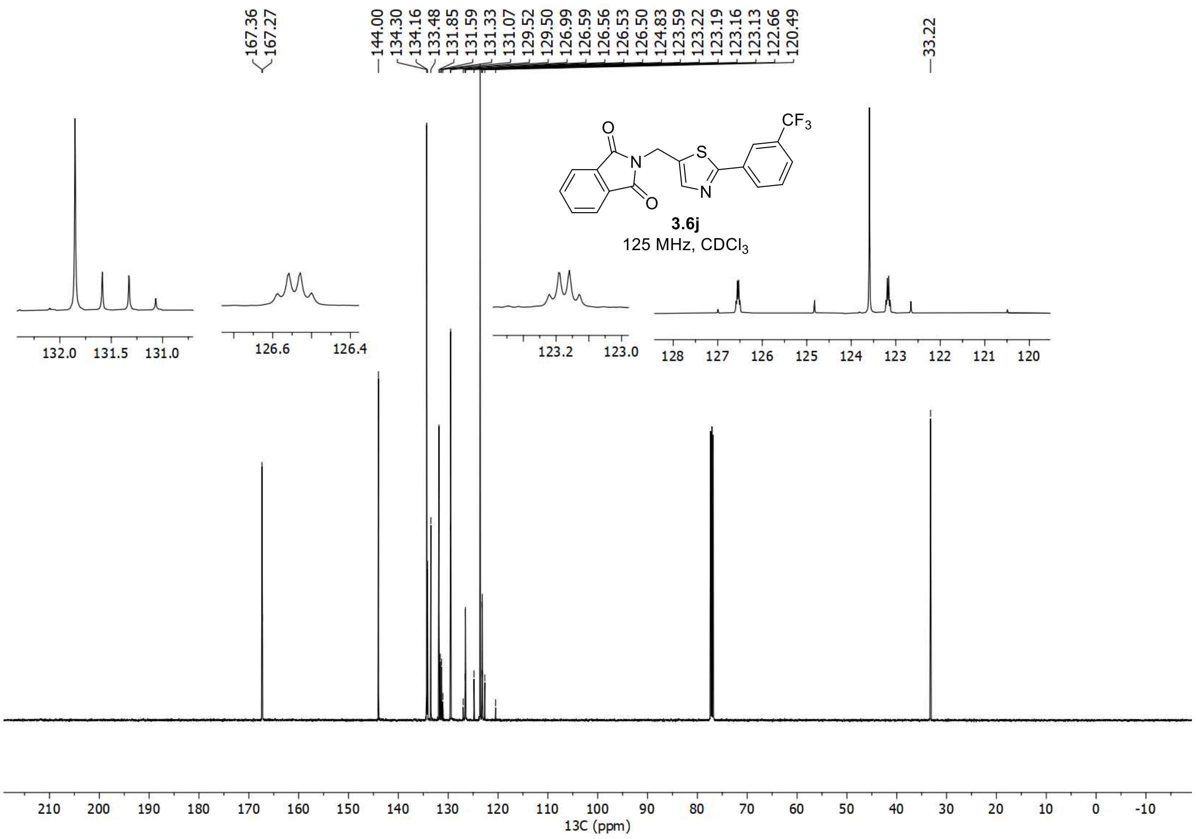

167.36
167.27
144.00
134.30
134.16
133.48
131.85
131.59
131.33
131.07
129.52
129.50
126.99
126.59
126.56
126.53
126.50
124.83
123.59
123.22
123.19
123.16
123.13
122.66
120.49
33.22
CF3
O
N
S
N
O
3.6j
125 MHz, CDCl3
13C (ppm)

$^{1}H$, $^{19}F$, and $^{13}C$ spectra of **3.7j**

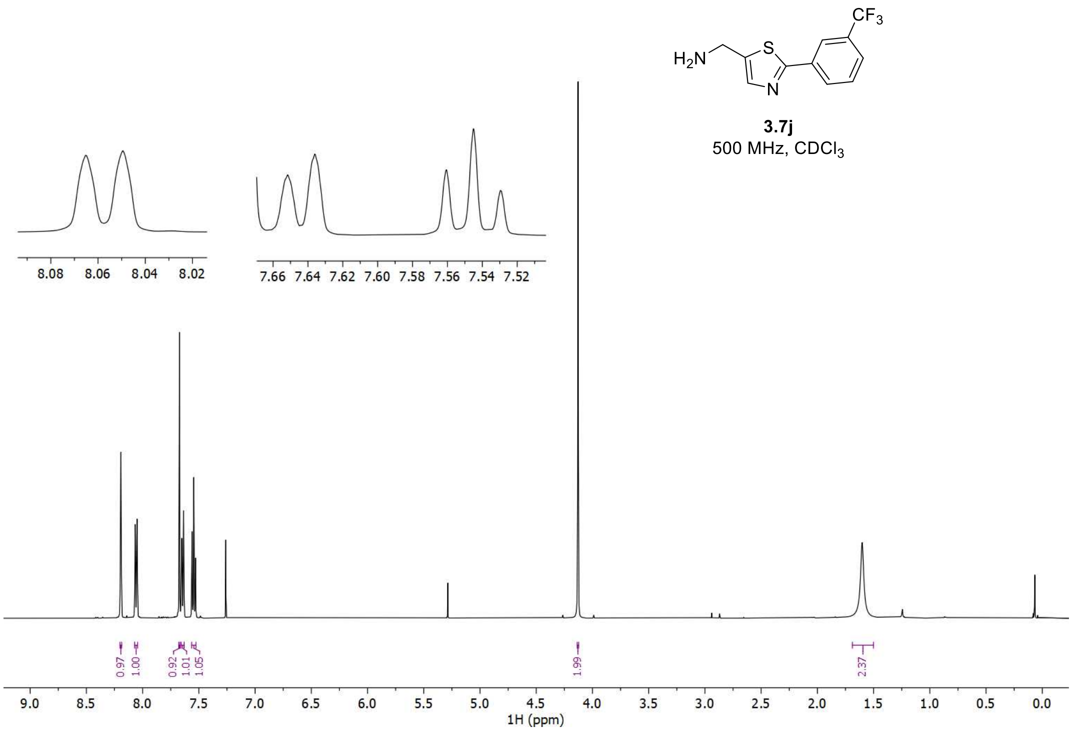


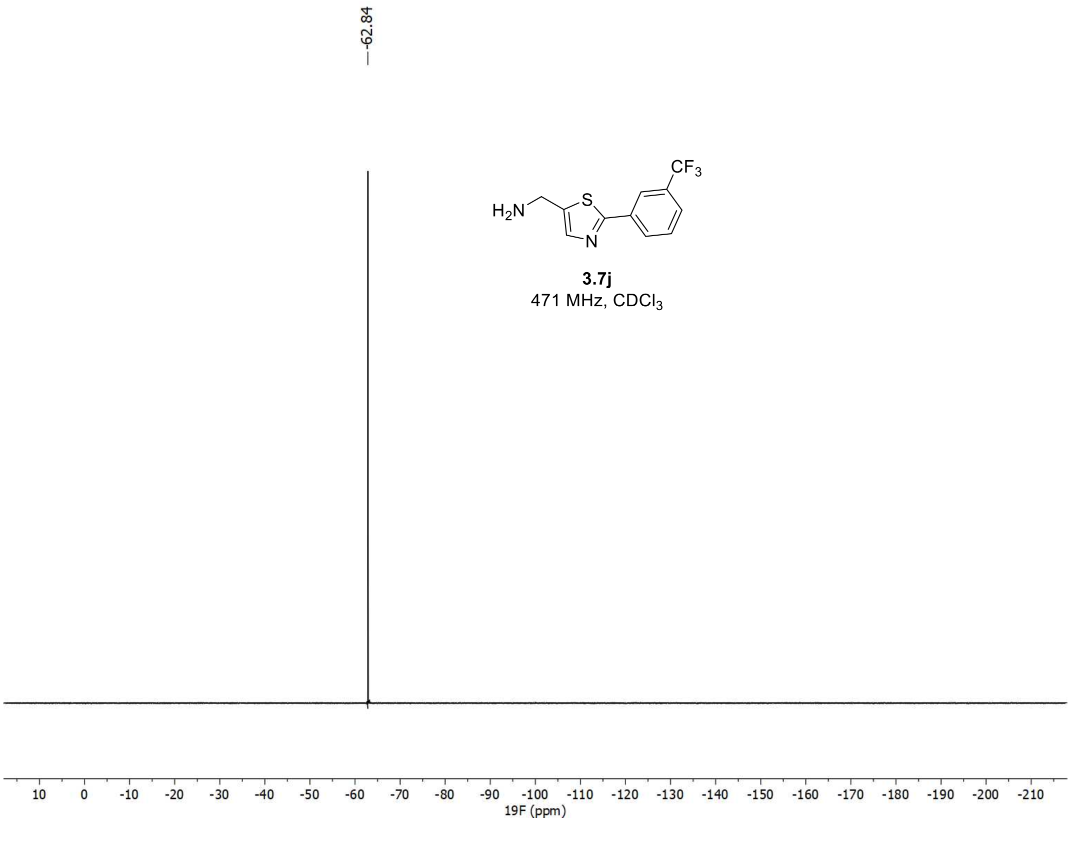

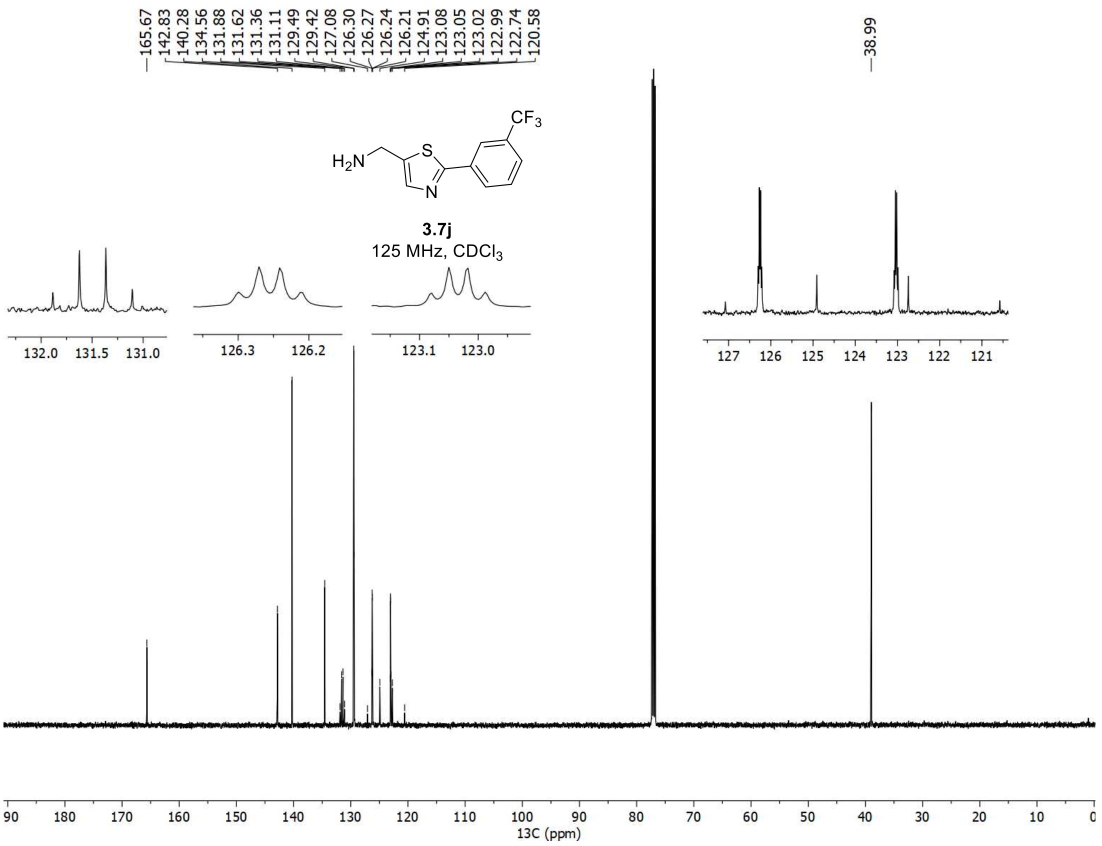

165.67
142.83
140.28
134.56
131.88
131.62
131.36
131.11
129.49
129.42
127.08
126.30
126.27
126.24
126.21
124.91
123.08
123.05
123.02
122.99
122.74
120.58
38.99
$CF_3$
$H_2N$
S
N
3.7j
125 MHz, $CDCl_3$
13C (ppm)

$^{1}H$, $^{19}F$, and $^{13}C$ spectra of **3j**

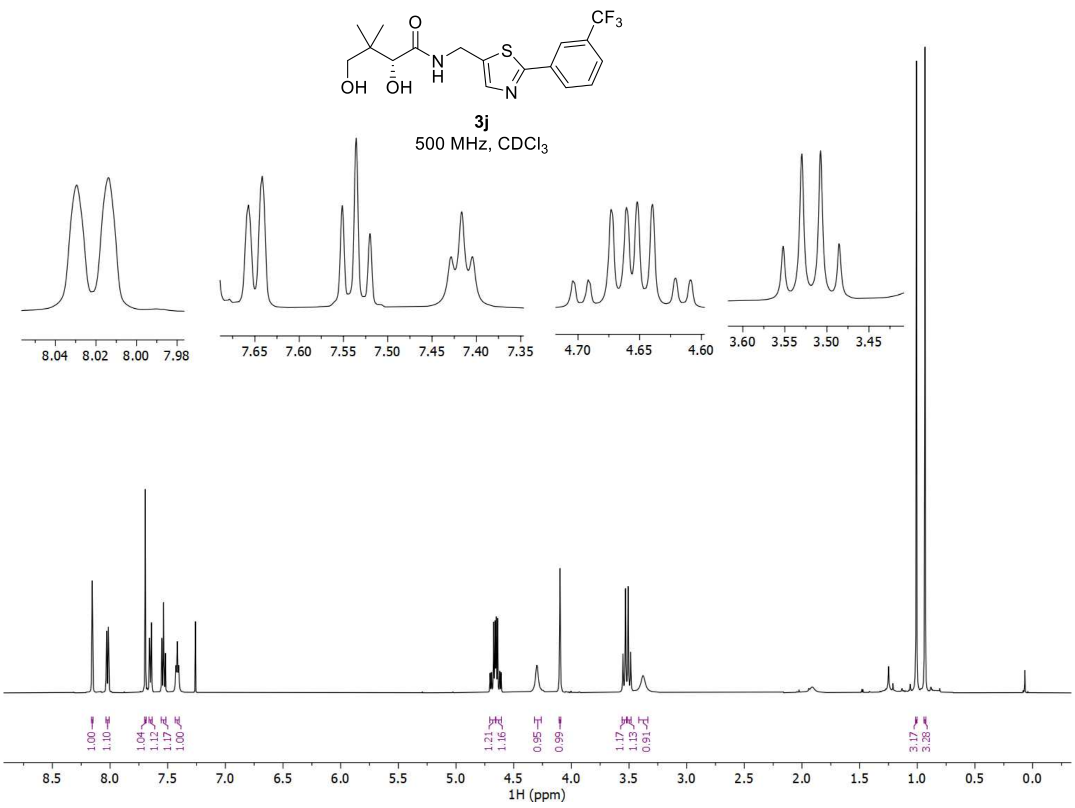


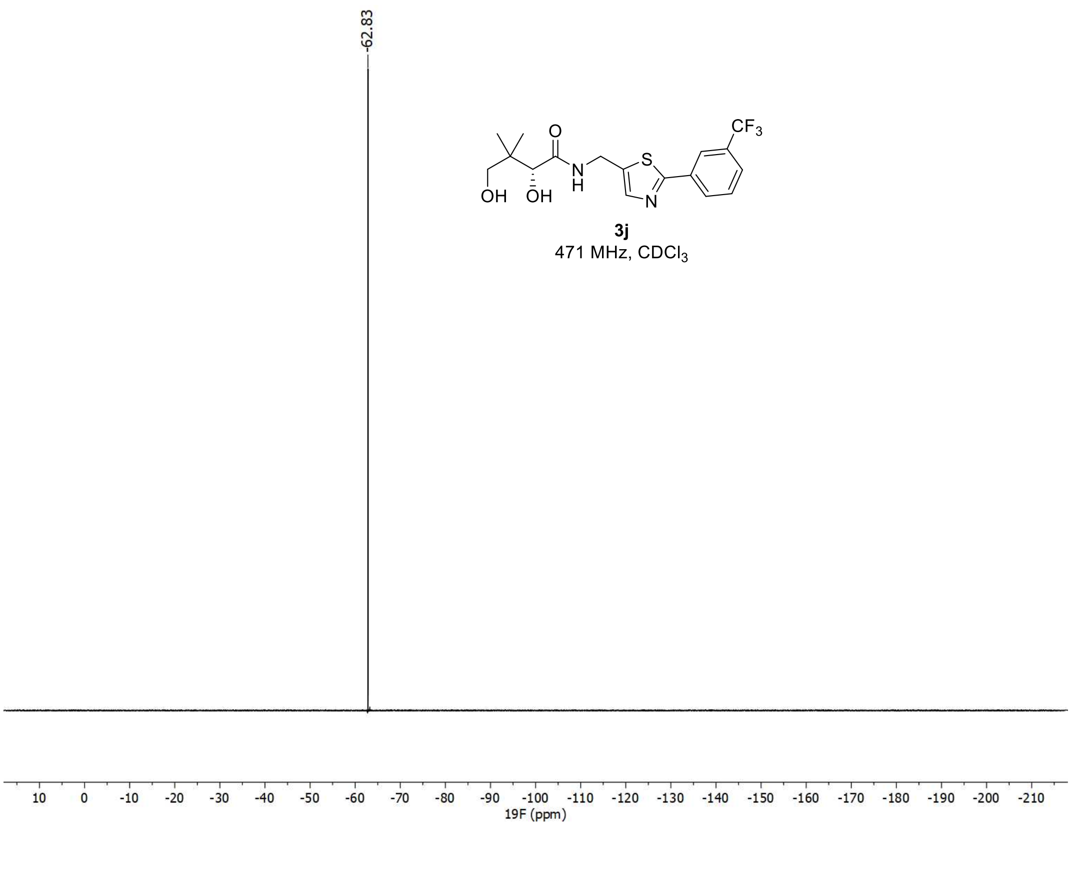

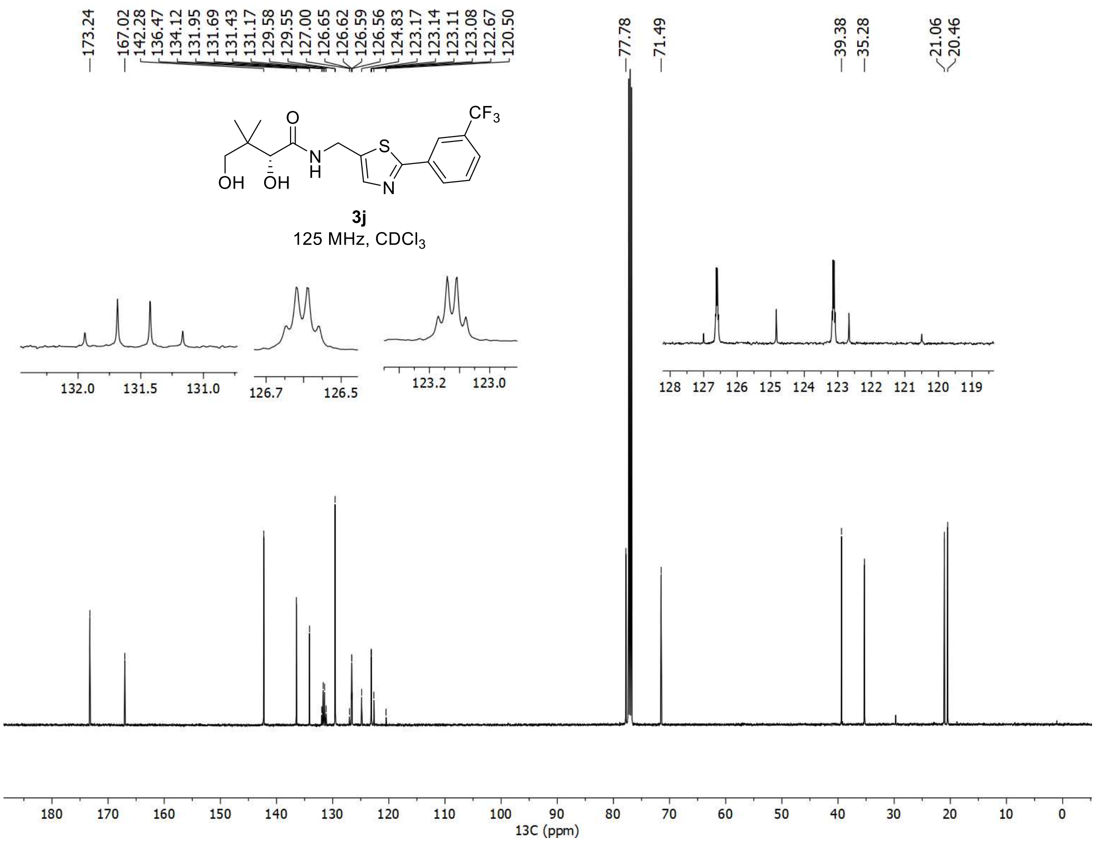
173.24
167.02
142.28
136.47
134.12
131.95
131.69
131.43
131.17
129.58
129.55
127.00
126.65
126.62
126.59
126.56
124.83
123.17
123.14
123.11
123.08
122.67
120.50
77.78
71.49
39.38
35.28
21.06
20.46
CF3
O
S
N
H
N
OH
OH
3j
125 MHz, CDCl3
13C (ppm)

#### 6. References


[1] Krasovskiy, A.; Knochel, P. Convenient Titration Method for Organometallic Zinc, Magnesium, and Lanthanide Reagents. *Synthesis* **2006**, *5*, 890-891.

[2] Lan, C. B.; Auclair, K. 1,5,7-Triazabicyclo[4.4.0]dec-5-ene: An Effective Catalyst for Amide Formation by Lactone Aminolysis. *J. Org. Chem*. **2023**, *88*, 10086-10095.

[3] Zhou, W.; Ni, S.; Mei, H.; Han, J. Pan, Y. Cyclization Reaction of *N*-Allylbenzothioamide for Direct Construction of Thiazole and Thiazoline. *Tetrahedron Lett.* **2015**, *56*, 4128-4130.

[4] Dexter, H. L.; Williams, H. E. L.; Lewis, W.; Moody, C. J. Total Synthesis of the Post-translationally Modified Polyazole Peptide Antibiotic Goadsporin. *Angew. Chem. Int. Ed.* **2017**, *129*, 3115-3119.

[5] Marsicano, V.; Arcadi, A.; Aschi, M.; Michelet, V. Experimental and Computational Evidence on Gold-Catalyzed Regioselective Hydration of Phthalimido-Protected Propargylamines: An Entry to β-Amino Ketones. *Org. Biomol. Chem.* **2020**, *18*, 9438-9447.

[6] Pace ,V.; Holzer, W. A Straightforward and General Access to α-Phthalimido-α'-Substituted Propan-2-ones. *Tetrahedron Lett.* **2012**, *53*, 5106-5109.

[7] Itoh, F.; Kimura, H.; Igata, H.; Kawamoto, T.; Sasaki, M.; Kitamura, S. JNK Inhibitor. EP 1 484 320 A1.

[8] Guo, J.; Hao, Y.; Ji, X.; Wang, Z.; Liu, Y.; Ma, D.; Li, Y.; Pang, H.; Ni, J.; Wang, Q. Optimization, Structure–Activity Relationship, and Mode of Action of Nortopsentin Analogues Containing Thiazole and Oxazole Moieties. *J. Agric. Food Chem*. **2019**, *67*, 10018-10031.

[9] Yadav, A. K.; Srivastava, V. P.; Yadav, L. D. S. Metal-Free, One-Pot Oxidative Conversion of Aldehydes to Primary Thioamides in Aqueous Media. *Synth. Commun*. **2014**, *44*, 408-416.

[10] Imaeda, Y.; Wakabayashi, T.; Kimura, E.; Tokumaru, K. Thiazole Derivative. EP 2 530 078 A1.

[11] Jin, H.; Ge. X.; Zhou, S. General Construction of Thioamides under Mild Conditions: A Stepwise Proton Transfer Process Mediated by EDTA. *Eur. J. Org. Chem*. **2021**, *2021*, 6015-6021.

[12] Li, H.; Wang, K.; Zhao, W.; Li, X.; Fu, Y.; Do, H.; An, J.; Hu, Z. Highly Chemoselective Synthesis of α, α-Dideuterio Amines by the Reductive Deuteration of Thioamides Using Mild $SmI_2$–$D_2O$. *Org. Lett.* **2024**, *26*, 9120-9125.

[13] Wang, C.; Han, C.; Yang, J.; Zhang, Z.; Zhao, J. Ynamide-Mediated Thioamide and Primary Thioamide Syntheses. *J. Org. Chem.* **2022**, *87*, 5617-5629.

[14] Orr, D.; Tolfrey, A.; Percy, J. M.; Frieman, J.; Harrison, Z. A.; Campbell-Crawford, M.; Patel, V. K. Single-Step Microwave-Mediated Synthesis of Oxazoles and Thiazoles from 3-Oxetanone: A Synthetic and Computational Study. *Chem. Eur. J.* **2013**, *19*, 9655-9662.

[15] Wei, J.; Li, Y.; Jiang, X. Aqueous Compatible Protocol to Both Alkyl and Aryl Thioamide Synthesis. *Org. Lett.* **2016**, *18*, 340-343.
